# Site Pattern Probabilities Under the Multispecies Coalescent and a Relaxed Molecular Clock: Theory and Applications

**DOI:** 10.1101/2021.08.11.455878

**Authors:** A. Richards, L. Kubatko

## Abstract

The first step in statistical inference of the evolutionary histories of species is developing a probability model that describes the mutation process as accurately and realistically as possible. A major complication of this inference is that different loci on the genome can have histories that diverge from the common species history and each other. The multispecies coalescent process is commonly used to model one source of this divergence, incomplete lineage sorting, or ILS. Chifman and Kubatko (2015) computed the site pattern probabilities for four taxa under a full probability model based on the Jukes-Cantor substitution model when the molecular clock holds. This paper generalizes that work to a relaxed clock model, allowing for mutation rates to differ among species. This will enable better phylogentic inference in cases where the molecular clock does not hold.

## 2 Introduction

We begin with a collection of species as demonstrated in figure 1. The thick outer lines represent the evolutionary history of a set of six mammalian species. We can see that the chimpanzee is our closest relative, the mouse is most closely related to the rabbit, and so on. The inner blue lines represents the history of one locus on the genomes. At this particular locus, the mouse is more closely related to the (chimp,human) pair than it is to the rabbit. Different lineage histories could be depicted for this set of six species for each common locus on their respective genomes.

**Figure 1:**
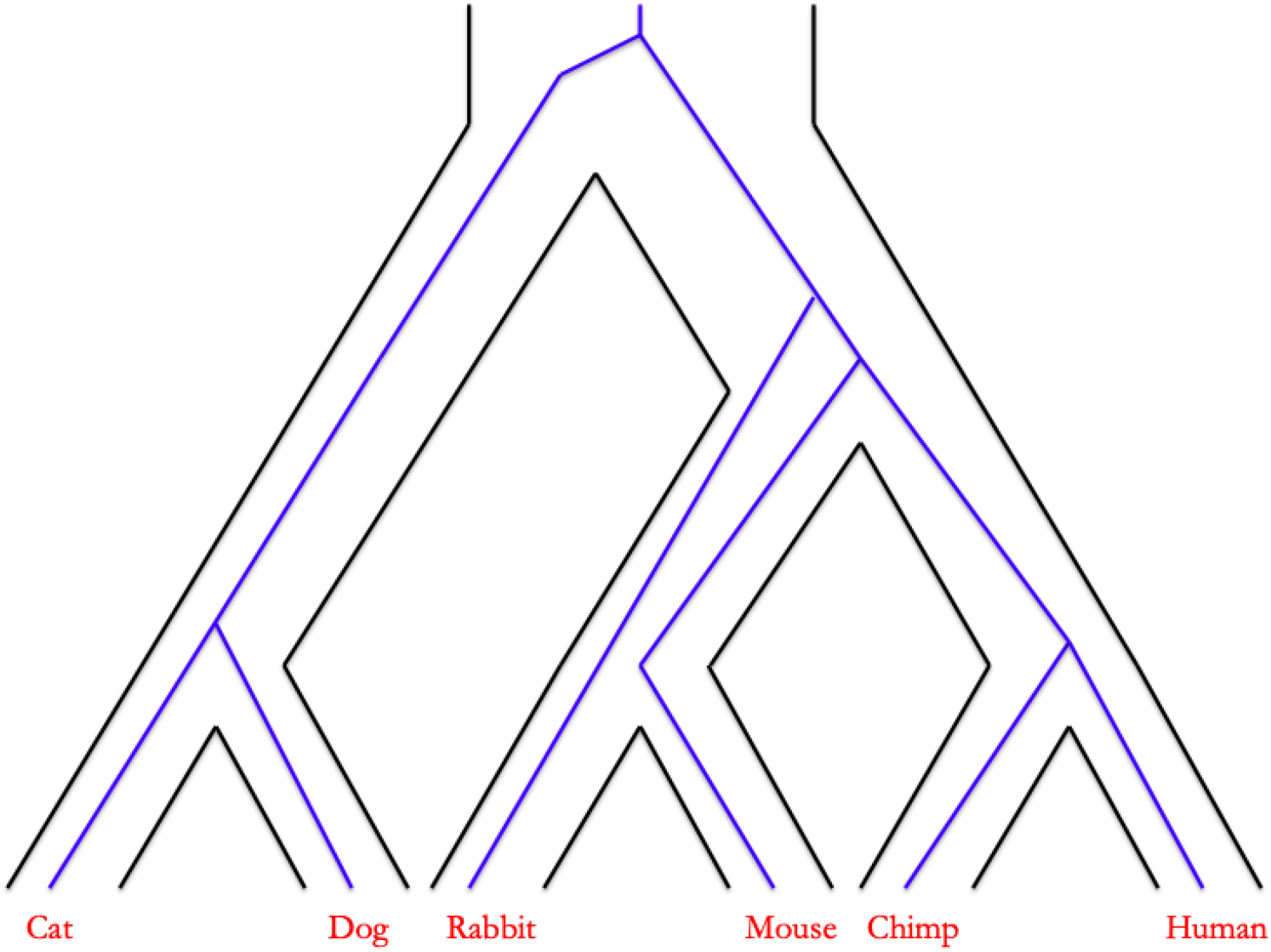
The multispecies coalescent problem: how to infer the species tree topology (black outer lines) when there may be mismatch with the gene tree (inner blue lines).

We describe the evolutionary histories of the species via a species tree:

### Definition 1

A *species tree* is an acyclic graph *S* = (*V* (*S*), *E*(*S*), ***τ*** _*S*_) where *V* (*S*) is the vertex set of *S, E*(*S*) is the edge set of *S*, and ***τ*** _*S*_ is a vector of node times.

The external nodes on the species tree represent extant species while the internal nodes represent speciation events. Internal branches represent ancestral species. If we know the common ancestor of all the species under consideration, then the tree is *rooted*, and the tree becomes a directed graph from the root outward. In the case that the location of the common ancestor is unknown, then we have an unrooted tree. Then we only know the direction of the branches that connect to external nodes. As we will see, an unrooted tree is sometimes also used when knowledge of the root location is unimportant in order to ease the computational burden.

Species tree inference is complicated by the possibility for divergence between the evolutionary history of species and individual elements of their genomes. Causes for this divergence include incomplete lineage sorting (ILS), gene duplication and loss (GDL) and horizontal gene transfer (HGT). ILS is commonly modeled by the coalescent process [8]. The history of individual loci on the genome is represented by a gene tree.

### Definition 2

A *gene tree* is a acyclic graph *G* = (*V* (*G*), *E*(*G*), ***t***_*G*_) where *V* (*G*) is the vertex set of *G, E*(*G*) is the edge set of *G*, and ***t***_*G*_ is a vector of node times. These node times are subject to the constraint that the nodes of the gene tree must must occur prior to the speciation time of the species in question.

In the sequel, we will assume there is one individual sampled from each species and there is no missing data. Then, both trees have the same leaf set *L*. When we need to distinguish the leaves of *G* from the leaves of *S* we use capital letters *A, B, C*, … for the gene tree and lower case for the species tree: *a, b, c* …. Internal nodes on the gene tree represent coalescent events, which identify the most recent common ancestor of two gene lineages. So the symmetric tree in figure 2a has a species tree ((*a, b*), (*c, d*)) and gene tree (*D*, (*A*, (*B, C*))).

**Figure 2:**
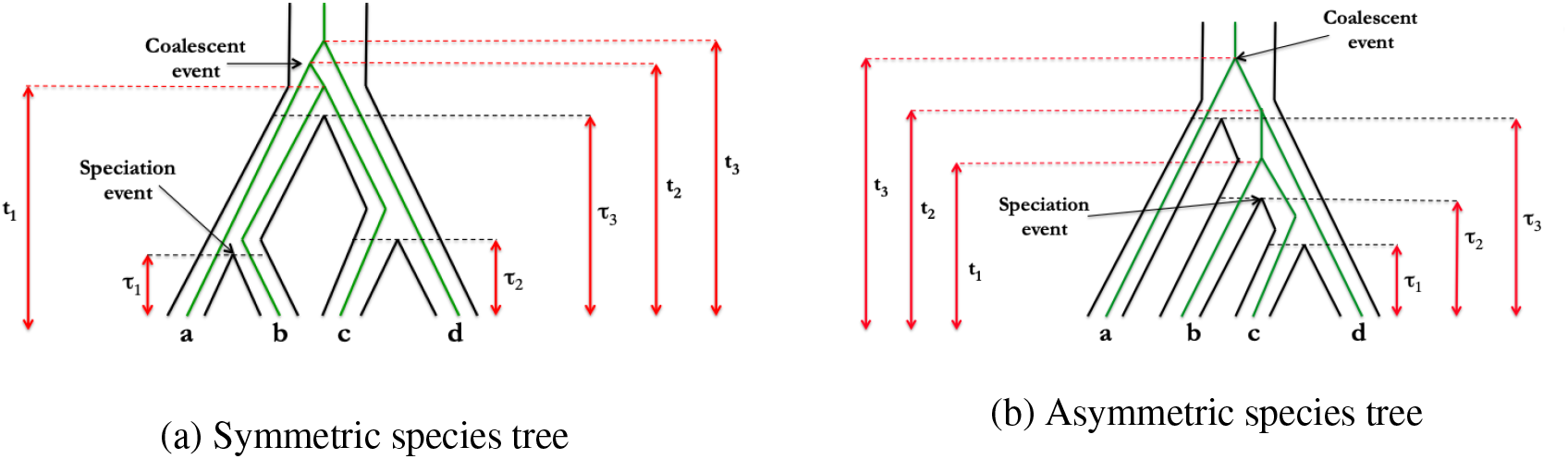
Species trees with four taxa under the coalescent process. The green lines show example gene trees evolving within the underlying species tree.

A number of different approaches have been taken with regard to species tree inference in the presence of ILS. The first is essentially to ignore the problem: perform gene tree inference on concatenated data using methods such as RAxML [28] or FastTree [16], treating all sites as if they share a single, common evolutionary history. This can be fast and accurate for estimating *S*. But, there are some concerns: concatenation has been shown to be statistically inconsistent for some values of (*S*, ***τ***) [4, 20], and speciation time estimates are biased since the coalescent event must naturally occur before the speciation time. Another approach is the use of summary statistics that first estimate the gene trees independently for each gene, and then use the gene tree estimates as inputs for species tree estimates. Examples of this approach include STEM [9], ASTRAL [13, 14, 36], and MP-EST [12]. These methods can be computationally efficient, however they depend on the accuracy of the gene tree estimates that are used as inputs as well as proper delineation of recombination-free segments of the genome [7, 27]. A third approach uses the full data to coestimate the species tree and each of the gene trees, generally using Markov chain Monte Carlo (MCMC) methods. Examples of this approach include BEST [11], *BEAST [5, 15], and BPP [35]. These methods can be quite accurate but are very computationally intensive when the number of loci and/or leaf set is large. Assessment of convergence is also a challenge, especially due to the multi-modal nature of the likelihood in the tree space [22].

A fourth approach, and the one we take in this paper, is to treat the gene trees as a nuisance parameter that can be integrated over. Previous examples of this approach include SVDQuartets [2] and Lily-Q [19]. SVDQuartets uses a methodology based in linear algebra to infer the proper unrooted species tree for each quartet of species under consideration, and then uses an assembly algorithm to infer the final n-taxon species tree estimate taking the set of unrooted quartet trees as input. Lily-Q instead takes the site pattern probabilities *P* (*D*|(*S*, ***τ***)) and then assumes a distribution on *τ* and *S* to find a posterior probability of *S* given the data *D*. This was shown to be fast and highly accurate, but the probabilities *P* (*D*|(*S*, ***τ***)) from [3] were calculated assuming a constant mutation rate over time (or more precisely, if *λ* represents the mutation rate there is a common *λ*(*t*) for all branches at time *t*), an assumption known as the *strict molecular clock*. Inference using Lily-Q was very sensitive to this assumption, prompting the need to generalize the work of [3] to cases for which the clock does not hold.

## 3 Model

Our model is as follows. We assume that there is no missing data and that each site is properly aligned across species and has an independent evolutionary history from every other site given the species tree. In other words, each site has its own gene tree which is a random draw from the distribution of gene trees induced by the coalescent process, *P* (*G* = *g*|*S*). We also assume neutral selection, which implies that the substitution process and coalescent process are independent [33]. Let 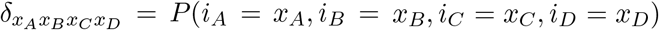, *X*_*j*_ ∈ {*A, C, G, T*} for *j* ∈ {*A, B, C, D*}. Then *δ*_*TCCG*_, for example, represents the probability that at a certain site the first species has T, the next two have C, and the fourth species has G, i.e., *δ*_*TCCG*_ = *P* (*i*_*A*_ = *T, i*_*B*_ = *C, i*_*C*_ = *C, i*_*D*_ = *G*) where *i*_*x*_ is the nucleotide for species *x* at any given site. Under the Jukes-Cantor substitution model, all four nucleotides have the same limiting probability of 1/4 and the same mutation rate between them, so *δ*_*TCCG*_ = *δ*_*AGGT*_ , for example, and we can instead refer to both of these probabilities as *δ*_*YXXZ*_. In fact, there are fifteen different site patterns under the Jukes-Cantor model, whose frequencies can be summarized by the 15 × 1 vector ***D*** = {*d*_*XXXX*_, *d*_*XXXY*_ , … *d*_*XYZW*_} which follows a multinomial distribution with probabilities ***δ*** = {*δ*_*XXXX*_, *δ*_*XXXY*_ , … *δ*_*XYZW*_} [19]. We parameterize the species and gene tree as follows. Let ***τ*** represent the speciation times (the times associated with the nodes of the species tree) looking back from the present. Similarly, ***t*** represents coalescent times measured back from the present. Time is measured in *mutation units*, so one unit of time is the expected time for one mutation in the common ancestor of all species (the mutation rate on the branch above the root of the tree).

Initially, we assume that there is a constant mutation rate on each branch of the tree, and parameterize these rates relative to the mutation rate above the root with the vector of relative rates given by ***γ***. For the symmetric tree, ***γ*** = {*γ*_*A*_, *γ*_*B*_, *γ*_*C*_, *γ*_*D*_, *γ*_*AB*_, *γ*_*CD*_} and for the asymmetric tree ***γ*** = {*γ*_*A*_, *γ*_*B*_, *γ*_*C*_, *γ*_*D*_, *γ*_*CD*_, *γ*_*BCD*_}, where the subscript indicates the set of species descending from the branch. Thus, we do not need to make an assumption about the absolute mutation rate to calculate probabilities, since we use mutation units for time and reference all relative rates to the rate above the root. For the pendant branches, whether or not there is a constant rate on the branch is irrelevant, but for internal branches we will later generalize our model in section 4 to allow for changing mutation rates over time.

The probability of the data is also dependent on the effective population size *N*_*e*_ through the population parameter *θ* = 4*N*_*e*_*µ*, where *µ* is the mutation rate per site per generation. So, our goal is to determine:

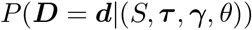

which due to the independence of the coalescent and substitution processes can be broken into

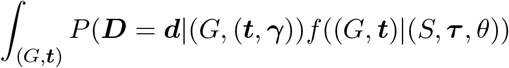

where the first term describes the substitution process and the second the coalescent process. Since we view the gene trees as a nuisance parameter, we integrate over all gene trees consistent with the species tree, 𝒢_*S*_:

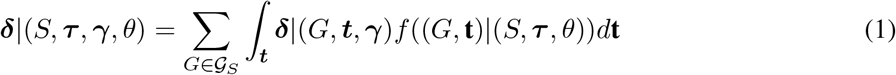

We will discuss the derivation of both of these terms beginning with the gene tree density given the species tree.

### 3.1 Gene tree densities given the species tree

#### 3.1.1 Background

Looking at figure 2, we can see that not all combinations (*G*, ***t***) are possible given (*S*, ***τ***). For example, if the species tree is ((*a, b*), (*c, d*)), then (*D*, (*A*, (*B, C*))) can only occur with the constraint that *t*_1_ *> τ*_3_ because *b* and *c* only share a common ancestor before *τ*_3_. In fact, 25 different gene tree histories are possible in the symmetric case and 34 are possible in the asymmetric case as laid out in tables 4 to 6.

The gene tree density for each of these histories is given by [18] and [3]. Given *j* lineages, the time to the next coalescent event *t*_*j*_ follows an exponential distribution with rate 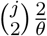. For any interior branch of the species tree, this process is censored, and so with *j* lineages entering a branch of length *τ*_*b*_ the probability of no coalescent events on the branch is given by 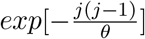. More generally, the density of the vector of coalescent times where *j* lineages enter an interior branch and *k* leave is given by:

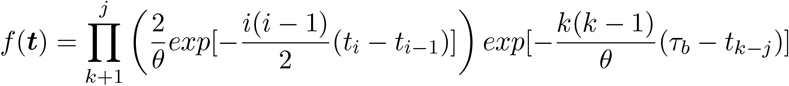

where *t*_0_ = 0 and time is measured from the start of the branch.

#### 3.1.2 Example

Figure 3 shows one example species tree, in this case the symmetric tree ((*a, b*), (*c, d*)) with the asymmetric gene tree (*A*, (*B*, (*C, D*))) embedded within. In this section, we will describe the derivation of that gene tree density, and in the next we will derive the mutation probabilities along the gene tree.

**Figure 3:**
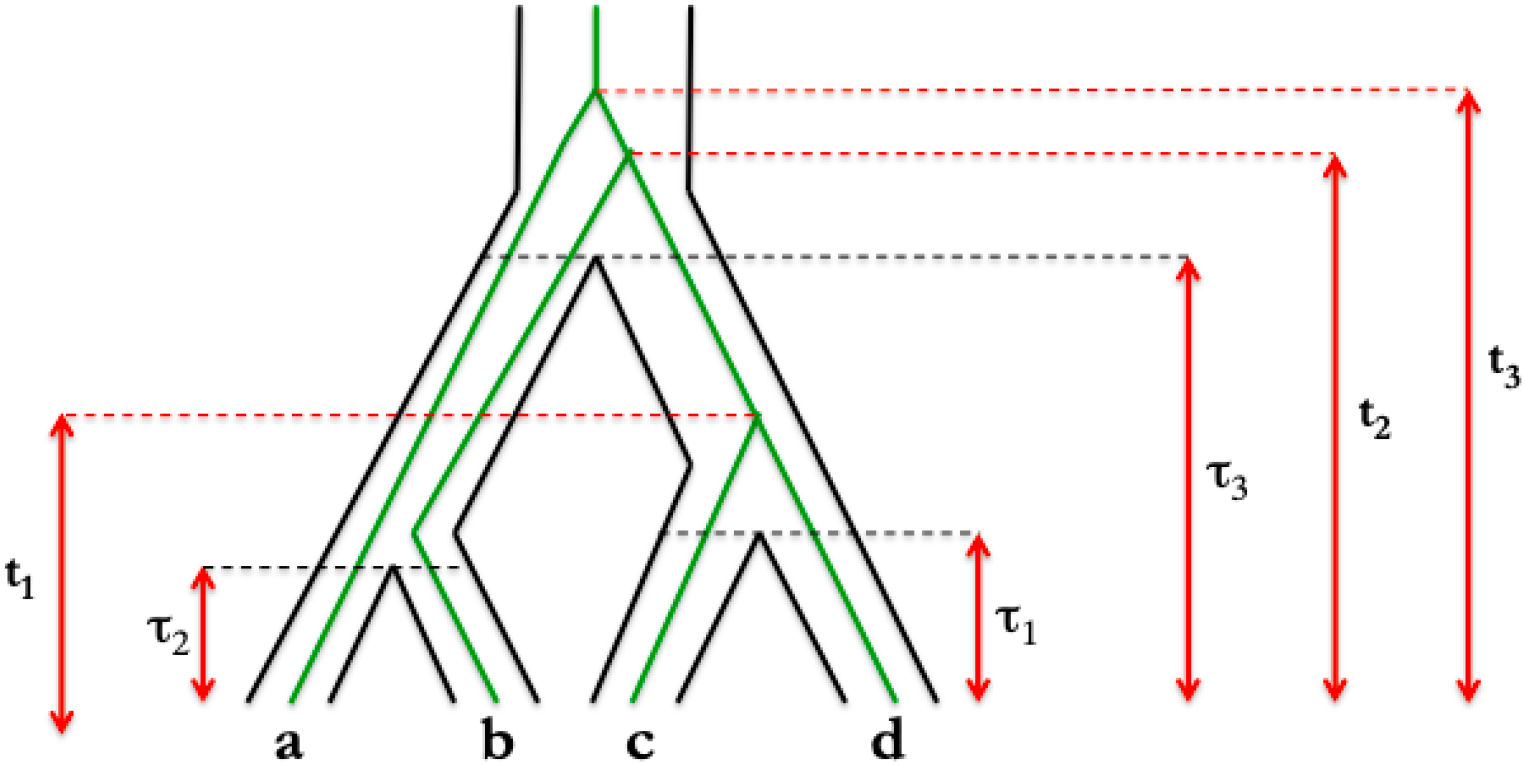
Example species tree and gene tree history to illustrate probability calculations

There is no coalescent event on the internal branch from the root to the divergence of (*a, b*). This occurs with probability 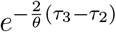. The coalescent event between *C* and *D* happens at rate 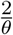, and can only occur between *τ*_1_ and *τ*_3_ in this gene tree history. The coalescent event between *B* and (*C, D*) occurs above the root and with a rate 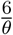, while the final coalescent event occurs after *t*_2_ and with a rate 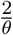. Finally, since there are three possibilities for the first coalescent event after the root, we divide the total probability by 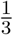, giving the density:

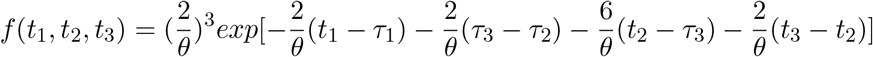

We will show in the next section that the site pattern probabilities given the gene tree are all linear combinations of terms that have the form 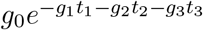. Then, we integrate over (*t*_1_, *t*_2_, *t*_3_) in the following manner:

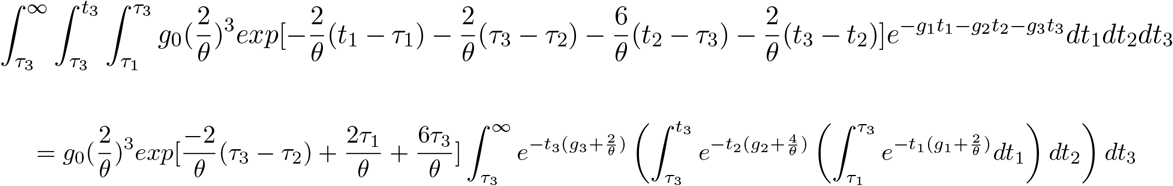

Let

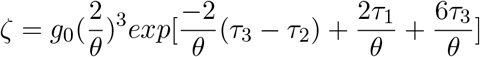

Then the above triple integral becomes:

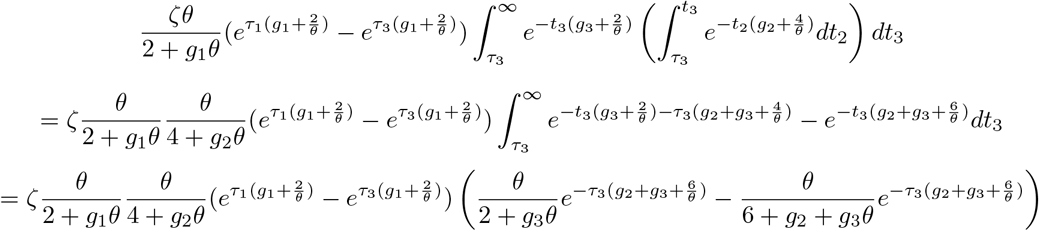

Canceling terms with *ζ* gives:

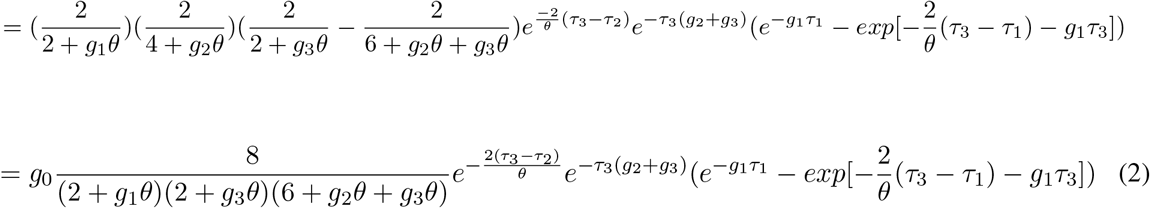

after algebraic simplification. This one example corresponds to history #10 from table 4 and integral form **B**. Similar calculations give the integral results for all 25 symmetric and 34 asymmetric gene tree histories, as listed in the appendix.

### 3.2 Site pattern probabilities given gene trees

We have from the Jukes-Cantor model that when time is measured in mutation units the transition probabilities on branch *l* with length *d*_*l*_ are given by:

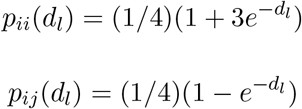

where *i, j* ∈ {*A, C, G, T*}.

Then, the probability of the observed data *i*_*A*_, *i*_*B*_, *i*_*C*_, *i*_*D*_ is given by:

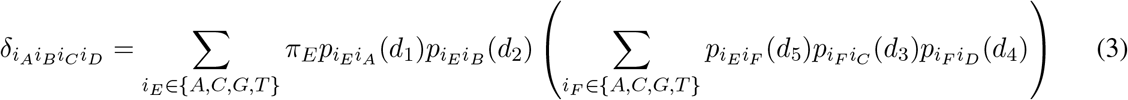

From the assumptions of the JC69 model, *π*_*E*_ = 1*/*4 for all nucleotides. We treat the gene tree as an unrooted tree in order to have one fewer branch substitution probability to concern ourselves with, and the nucleotide at the root can be safely ignored due to the Markov property. Although the Felsenstein pruning algorithm could be used here, the full enumeration of states is small enough to be accomplished readily by computer. The probability of any one particular state of the tree *i*_*A*_, *i*_*B*_, *i*_*C*_, *i*_*D*_, *i*_*E*_, *i*_*F*_ is given by:

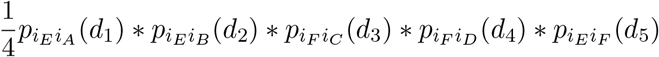

After the enumeration of states, the states that fit each site pattern were identified and common terms summed, with the results given both by [3] and repeated in the supplementary material. We now note that each of the terms 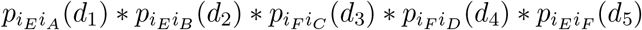 is a linear combination of 32 terms of the form:

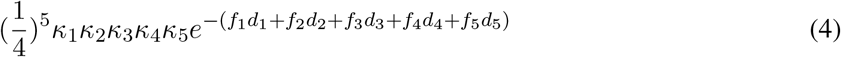

where each of the *κ*_*m*_ ∈ {−1, 3} and *f*_*m*_ ∈ {0, 1}.

Taken together, each of the fifteen different site pattern probabilities in the equations from the supplementary material is a linear combination of the terms in equation 4. Most of the terms vanish due to cancellations, with the result being another linear combination (where the coefficients are given by table 1):

**Table 1:**
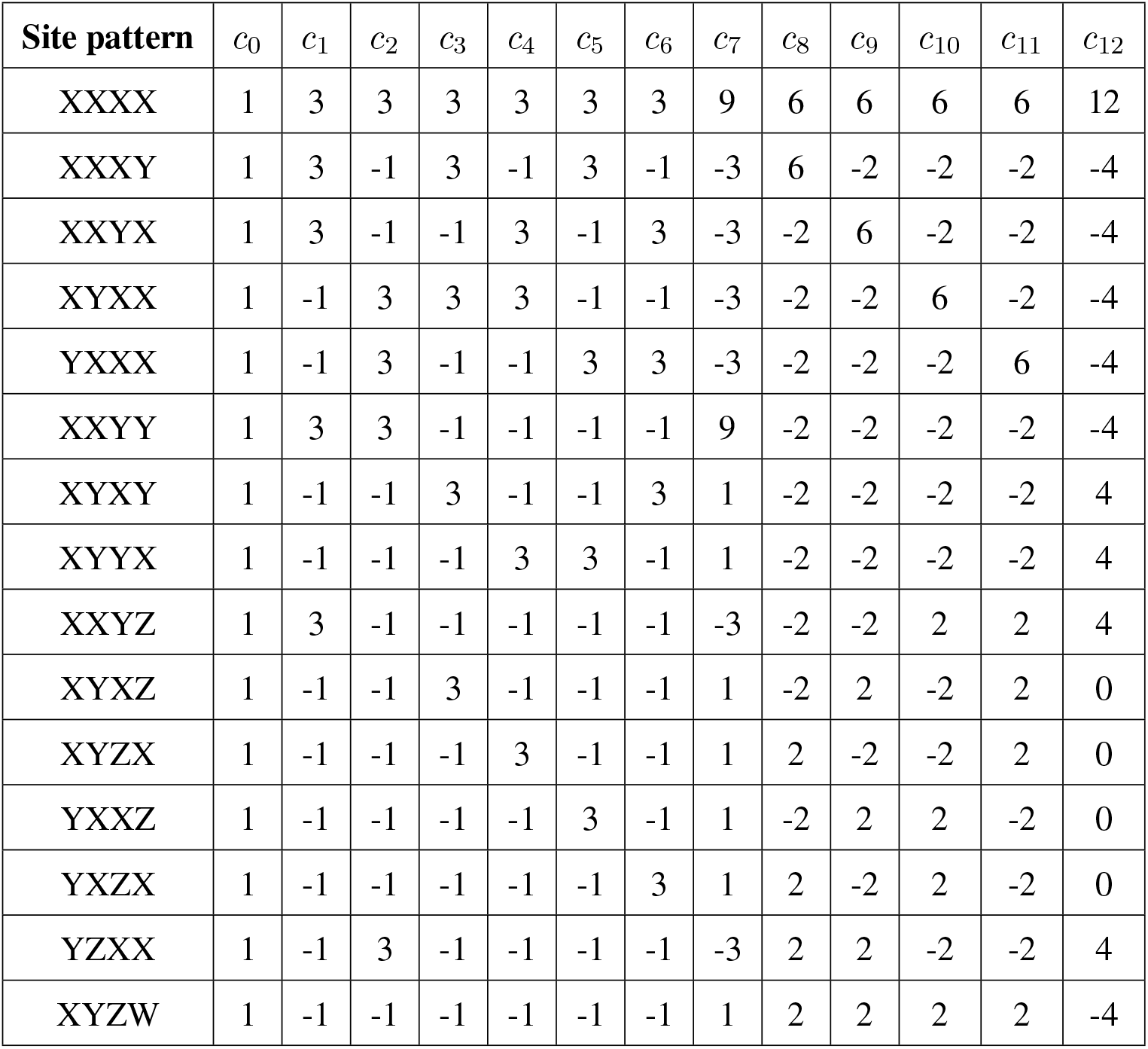
Coefficients for each gene tree site pattern probability

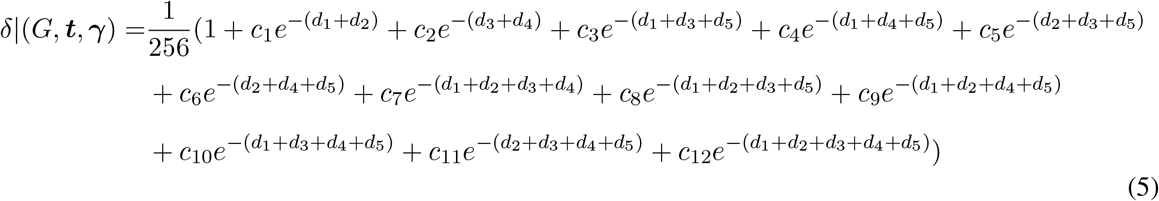

#### 3.2.1 Example

To find the branch lengths in the unrooted tree, we note that the branch lengths will be a function of the coalescent times and the mutation rate on each branch of the species tree. Refer back to our example in figure 3 and we can see how the gene tree in figure 4 arises within the species tree. Lineages *C* and *D* meet at time *t*_1_, but due to the different rates

**Figure 4:**
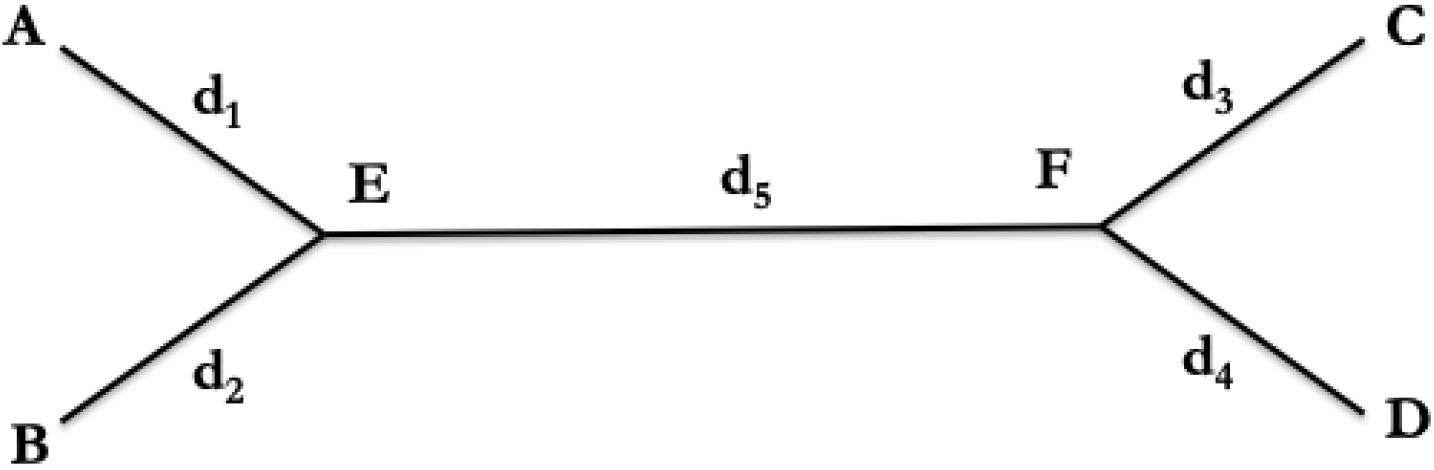
Branch length notation for 4-taxon (unrooted) gene tree

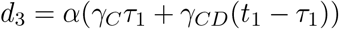

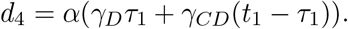

The term *α* arises from conversion to mutation units and for the Jukes-Cantor model is equal to 4*/*3. Likewise, it can be shown that:

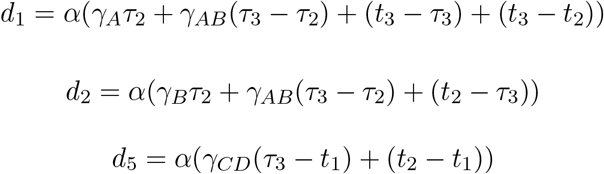

The additional term in *d*_1_ come from ignoring the root, so the path for *d*_1_ has to go “up” to the root and then back “down” to *t*_2_. Note we do not need a rate multiplier for the terms above the root of the species tree because we have defined the relative rates in reference to the rate of unity above the root. (Also, under the strict molecular clock all *γ*_*k*_ = 1 so *d*_1_ = 2*t*_3_ − *t*_2_, *d*_2_ = *t*_2_, *d*_3_ = *d*_4_ = *t*_1_, and *d*_5_ = *t*_2_ − *t*_1_.)

We will continue the example by considering one term of equation 5, showing how this term is found, and the results of entering it as an input into the integral result equation 2. The term we will expand is 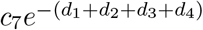. Then we have (ignoring the constant *c*_7_):

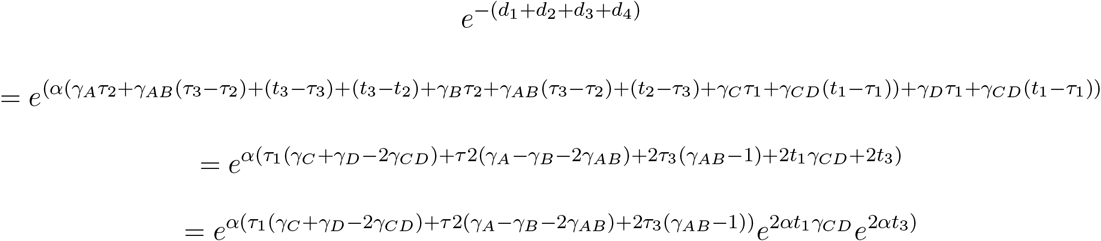

It can be seen then that the terms of equation 5 can be expressed by means of exponential functions of the coalescent times multiplied by a constant:

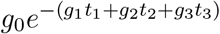

which can then be used in the integrals from the previous section to find the probability integrating over the gene tree histories. For the term in our example, 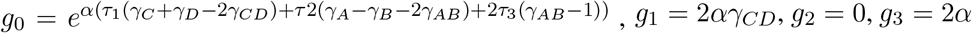. This can be repeated for all the terms of equation 5 for all histories, and the results are given in the supplementary material.

Plugging these results into equation 2 then gives the following:

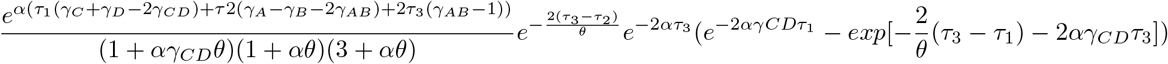

Repeating for all gene tree histories and terms in equation 5 gives the final results in the supplementary material. This example calculation corresponds to the *k*_114_ term, where we can see *k*_114_ = 2*c*_7_ because identical computations are performed for gene history 14.

If the unrooted gene tree is not of the topology (*A, B*|*C, D*) we need to permute the data. Figure 5 shows an example of how this can be done for one site pattern probability. Let *G*_*U*_ and *G*_*L*_ describe the upper and lower gene trees. Then given these phylogenies,

**Figure 5:**
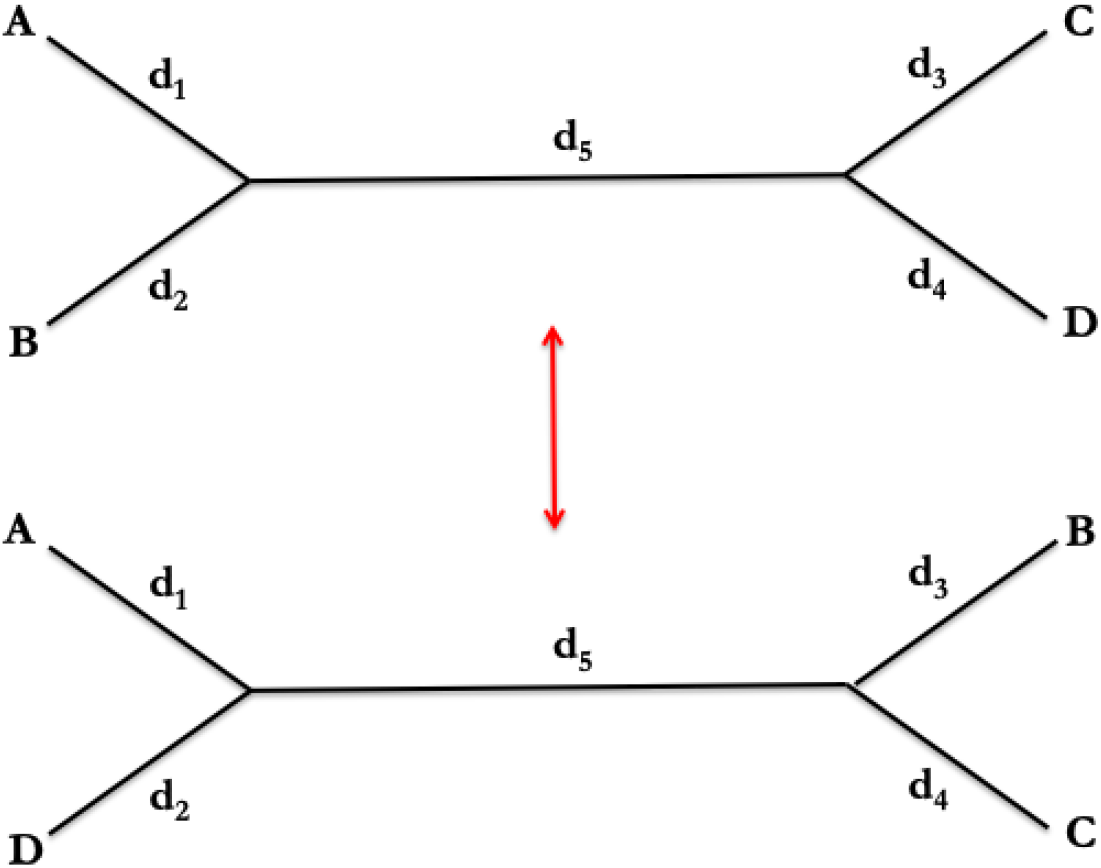
Site pattern probabilities given topologies other than (*A, B*|*C, D*) can be found by permuting as needed. *G*_*U*_ – upper tree. *G*_*L*_ – lower tree

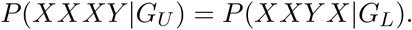

Table 2 gives the permutation matrix for when the gene tree is not (*A, B*|*C, D*). For example, let the gene tree of concern be (*C, A*|*B, D*) and we want to find the coefficients for equation 5 for site pattern XYYX. We go down the (*C, A*|*B, D*) column of table 2 until we find XYYX and then over to the (*A, B*|*C, D*) column and find this corresponds to XXYY. Then from table 1 we use the coefficients from the XXYY row in equation 5.

**Table 2:**
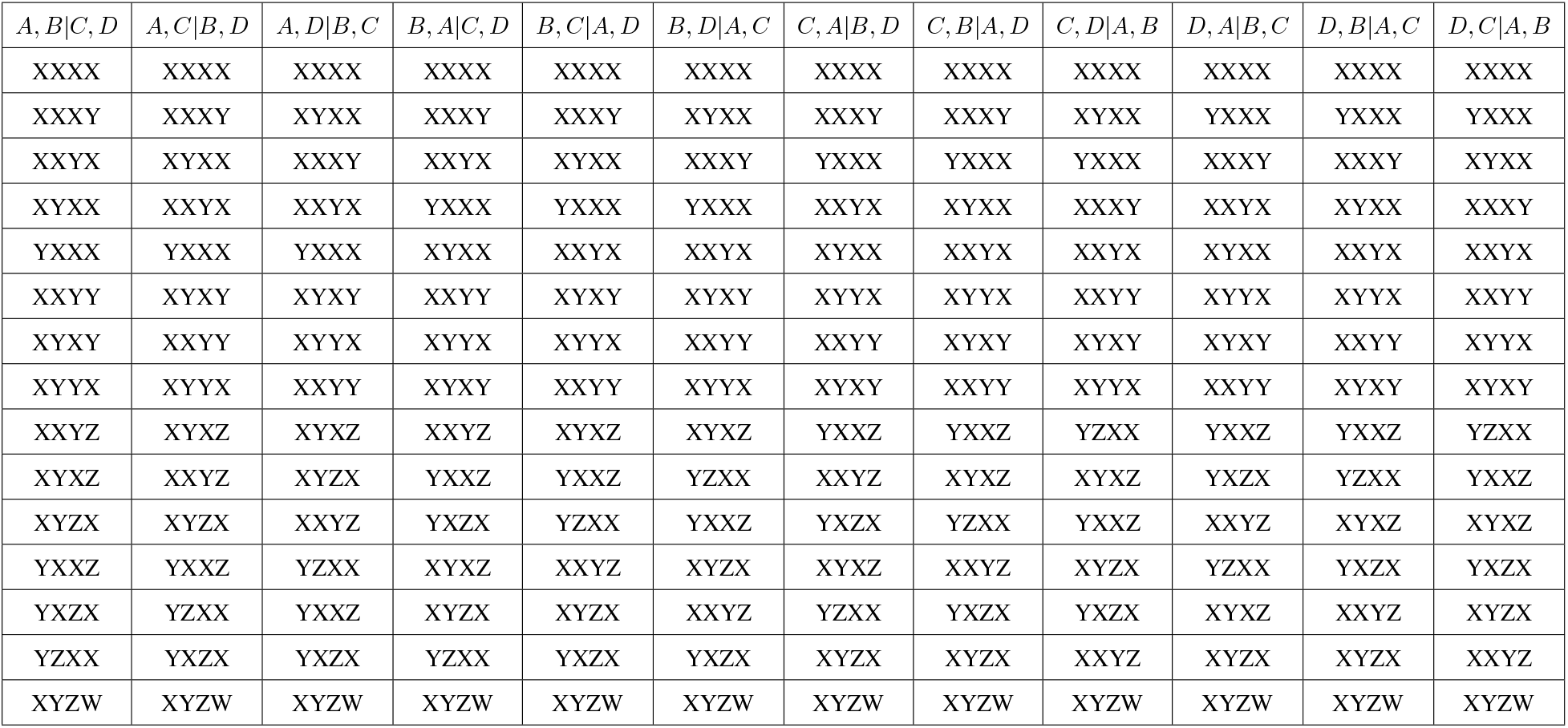
Site pattern permutation matrix

### 3.3 Implementation

Four programs, all written in C++ are provided at https://github.com/richards-1227/ noClock. The first performs the calculations that produced equation 3. The second conducts the linear algebra that results in table 4. The third produces the matrices of *k* values for the final result shown in the supplementary material. In addition, to prevent copying and editing errors, it outputs as strings the exact code used to instantiate the *k* matrix arrays used in the fourth program. The fourth program computes the final probabilities, and is included both as raw C++ code and a Mac and Unix executables. To run the program, enter:

~~~
./NoClock *τ*_1_ *τ*_2_ *τ*_3_ *γ*_*A*_ *γ*_*B*_ *γ*_*C*_ *γ*_*D*_ *γ*_*CD*_ *γ*_*AB*_ *θ*
~~~

for a symmetric tree and

~~~
./NoClock 0 *τ*_1_ *τ*_2_ *τ*_3_ *γ*_*A*_ *γ*_*B*_ *γ*_*C*_ *γ*_*D*_ *γ*_*CD*_ *γ*_*BCD*_ *θ*
~~~

for an asymmetric tree. Unlike the calculations in this paper, ***τ*** is parameterized in coalescent units, which are mutation units scaled by a factor of *θ/*2. The program outputs the vector of site pattern probabilities in the order given by table 4.

To check the accuracy of the implementation, we first generated site pattern probabilities for different branch lengths with ***γ*** equal to a vector of ones, representing a strict molecular clock. We compared that output to existing code which implements the results of [3] and confirmed that the results were identical.

Tian and Kubatko (2020) developed code that simulates the coalescent process for a 4-taxon species tree and then adjusts the gene tree lengths to allow for a relaxed clock. We simulated 1000 different species trees with various relative mutation rates, and then took the gene trees as inputs for simulating sequence evolution using seq-gen [17]. We then performed a Chi-squared goodness-of-fit test comparing the expected to actual counts of each site pattern and plotted the p-values (see figure 6). The results are consistent with having determined the correct site pattern probabilities, as it is improbable that both our implementation and the Tian and Kubatko implementation made simultaneous parallel errors resulting in the same probability distribution.

**Figure 6:**
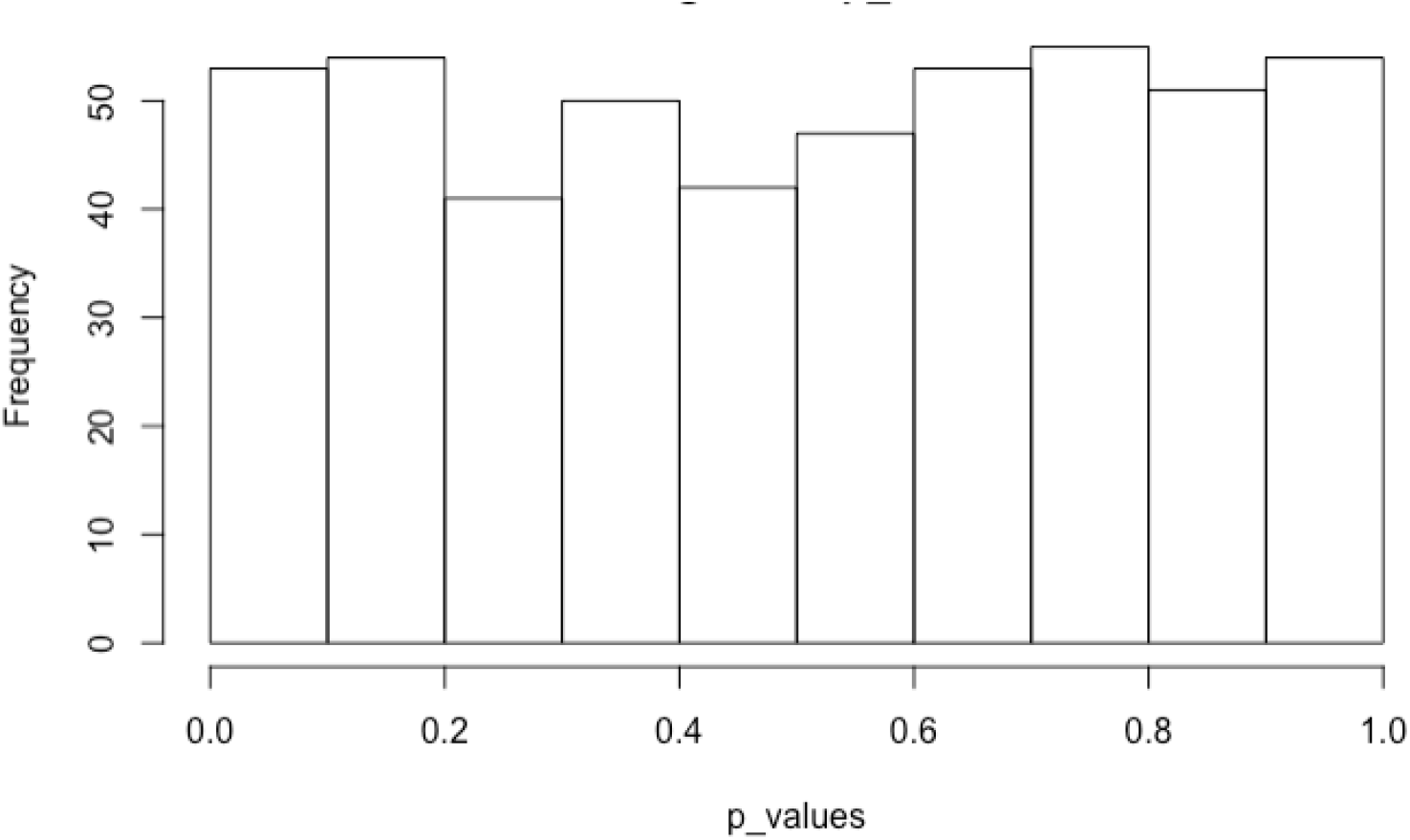
Distribution of 1000 goodness-of-fit test of our site pattern probabilities against simulated data generated by [32] and [17].

## 4 Varying mutation rates within branches

We next examine the case where the relative mutation rate, rather than being constant on a branch, is allowed to vary over time. For pendant branches, this distinction is unimportant, as coalescent events cannot happen on pendant branches of the species tree when there is only one individual per species. Thus, if the time-varying mutation rate on the pendant branch leading to species *d* is given by *γ*_*D*_(*u*), the total expected mutation is 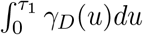, which leads to the same probability distribution as a constant rate of 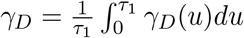.

For internal branches, it may be important to carefully consider differing relative rates at differing times. We could again try to replace the time-varying rates with an average rate as with pendant branches, for example 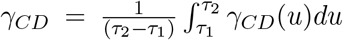. To see why this would fail to produce the correct site pattern probabilities, consider an asymmetric species tree (*a*, (*b*, (*c, d*))) where instead of a constant rate on the branch *cd*, the rate jumps halfway between *τ*_1_ and *τ*_2_ (see figure 7). We can let the two rates be *γ*_*CD*1_ and *γ*_*CD*2_ looking back from the speciation time between species *c* and *d* toward the root. Intuitively, if *γ*_*CD*1_ *> γ*_*CD*2_ there will be more mutation before the two lineages coalesce and *ceteris paribus* we will expect more XXXY and XXYX sites and fewer XXYY sites. Conversely, if *γ*_*CD*1_ *< γ*_*CD*2_ we have more XXYY and fewer XXXY and XXYX sites than if we treated the branch as having a single rate *γ*_*CD*_.

**Figure 7:**
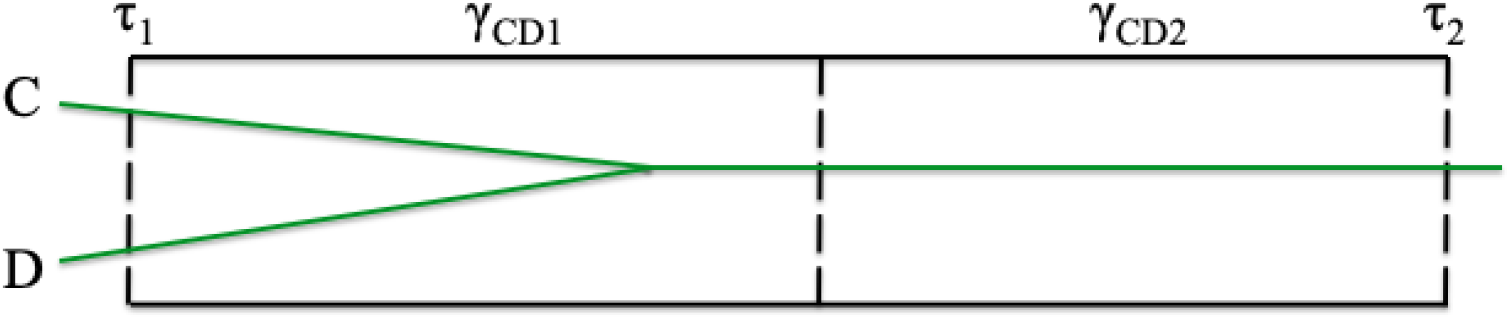
Effect of changing rates in internal branches. *γ*_*CD*1_ *> γ*_*CD*2_ will tend to produce more sites with the XXXY and XXYX patterns than taking a weighted average of *γ*_*CD*1_ and *γ*_*CD*2_. Conversely, *γ*_*CD*1_ *< γ*_*CD*2_ will tend to produce more XXYY sites.

Unfortunately, we cannot examine the full range of functional forms for *γ*_*CD*_(*u*) and other relative rates on the internal branches. Even a relatively simple linear form such as *γ*_*CD*_(*u*) = *u* − *τ*_1_ founders on the fact that after integrating over *u* we are left with a 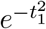 term, which famously has no known antiderivative. We thus limit our exploration of the possible relative rates on internal branches to step functions. This allows us to generalize our work from section 3 to find the site pattern probabilities, which also serve as a limiting approximation for any time-varying rate function that is continuous almost everywhere.

Unlike in the single rate case, we do not have a single algebraic formula that we can present for the site pattern probabilities. We will, however, show how the calculation steps of section 3 can be generalized, and discuss in detail our implementation program that calculates the site pattern probabilities given an input file of relative rates at different parts of the tree.

We use the following notation. Let ***ψ*** represent the series of breakpoints between different rates, where for example *ψ*_*CDi*_ is the i^th^ breakpoint on the branch above the speciation point between species *c* and *d*. Then, if there are *I* different rates on that branch, we have *ψ*_*CD*0_ = *τ*_1_, *ψ*_*CD*1_, … , *ψ*_*CDI*_ = *τ*_2_. Then *γ*_*j,I*_ describes the relative mutation rate on branch *j* between *ψ*_*j,i−*1_ and *ψ*_*j,i*_.

### 4.1 Generalizing the gene tree density integrals

Looking at the example in section 3 and the densities in the appendix, we see that each density is the product of two terms: a function of *θ* and ***τ*** that is a constant with respect to the coalescent times ***t***, and a function of ***t*** of the form *exp*[−(*a*_1_*t*_1_ + *a*_2_*t*_2_ + *a*_3_*t*_3_)]. Now, though, instead of 25 or 34 different gene tree histories (for symmetric and asymmetric species trees) we have additional histories depending on which breakpoints the coalescent events fall between. In the example in figure 3 if we had two different rates between *τ*_1_ and *τ*_3_ separated by *ψ*_*CD*1_, histories 1, 2 10, and 14 would need to be divided into separate cases for *t*_1_ before or after *ψ*_*CD*1_. Examining the limits of integration, we only need to consider four general cases (using ≐ to indicate two times fall in the same relative rate category and 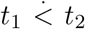 to indicate that a breakpoint falls between *t*_1_ and *t*_2_):

- 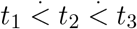
- 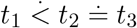
- 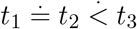
- *t*_1_ ≐ *t*_2_ ≐ *t*_3_

As before, we include the 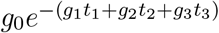 term from the the mutation process to get an integral of the form *∫ ∫ ∫ g*_0_*f* (*θ*, ***τ***)*exp*[−(*t*_1_(*a*_1_ + *g*_1_)) − (*t*_2_(*a*_2_ + *g*_2_)) − (*t*_3_(*a*_3_ + *g*_3_))]. Full details are given in the appendix.

As a final note, for a symmetric gene tree, we also need to consider cases when 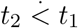 and when *t*_1_ ≐ *t*_2_ and *t*_2_ *< t*_1_. Since, by the assumptions of the coalescent model, the two cases are exchangeable and occur with equal probability, we can simply replace *t*_1_ with *t*_2_ and vice versa in the evaluation of the integral.

### 4.2 Generalizing the mutation probability term 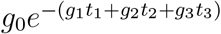

We now take a second look at the distance terms from the example in section 3. Taking a look at *d*_1_ we have

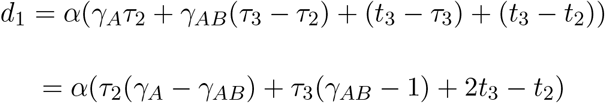

The first part of this result, *α*(*τ*_2_(*γ*_*A*_ − *γ*_*AB*_) + *τ*_3_(*γ*_*AB*_ − 1)), which is the contribution of this term to *g*_0_, is *α* times each of the break points times the difference in the relative mutation rates before and after the break point, or *ψ*_*j,i*_(*γ*_*j,i−*1_ − *γ*_*j,i*_). (In the event that *i* = 0 we can define *γ*_*j,i−*1_ to be the final rate of the branch below *j*.) The second part of the result, *α*(2*t*_3_ − *t*_2_), gives the contribution to *g*_2_ and *g*_3_ from the *d*_1_ term.

Taken together, we have

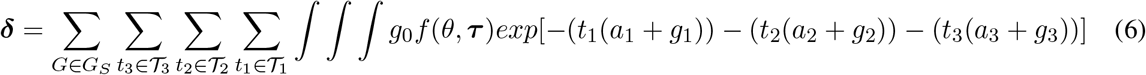

where 𝒯_*i*_ represents the sets of relative rate locations *t*_*i*_ can occur in (for example, in figure 3 *t*_1_ can occur in the *cd* branch or above the root, but not on the *ab* branch).

### 4.3 Implementing the step change model

We built a C++ program implementing equation 6 and located at https://github.com/richards-1227/noClock. We have included Mac and Unix executables as well as the C++ source code. Interested readers are encouraged to follow the source code along with the discussion in this section to see how the mathematical formulae are implemented.

Execution is via the command

~~~
. /NoClockSC symInd fileName *θ*
~~~

where symInd = 0 for an asymmetric tree and 1 for a symmetric tree. “fileName” is the name of a text file containing the sets of rates and breakpoints in the following format:

~~~
branch start_time end_time rate
~~~

For example, for a case where the *ab* branch has a single rate and the *cd* and root branches have two different rates each:

|  |  |  |  |
| --- | --- | --- | --- |
| A | 0 | $\tau_2$ | $\gamma_A$ |
| B | 0 | $\tau_2$ | $\gamma_B$ |
| C | 0 | $\tau_1$ | $\gamma_C$ |
| D | 0 | $\tau_1$ | $\gamma_D$ |
| AB | $\tau_2$ | $\tau_3$ | 1.0 |
| CD | $\tau_1$ | 2.5 | 2.0 |
| CD | 2.5 | $\tau_3$ | 1.5 |
| root | $\tau_3$ | 5.0 | 0.5 |
| root | 5.0 | 1000.0 | 1.0 |

The final time above the root must be chosen such that *e*^*ψ*^ ≈ 0. We have used 1000.0.

Pseudocode for implementing the calculations is shown below.

#### Algorithm 1

Site pattern probability calculation procedure

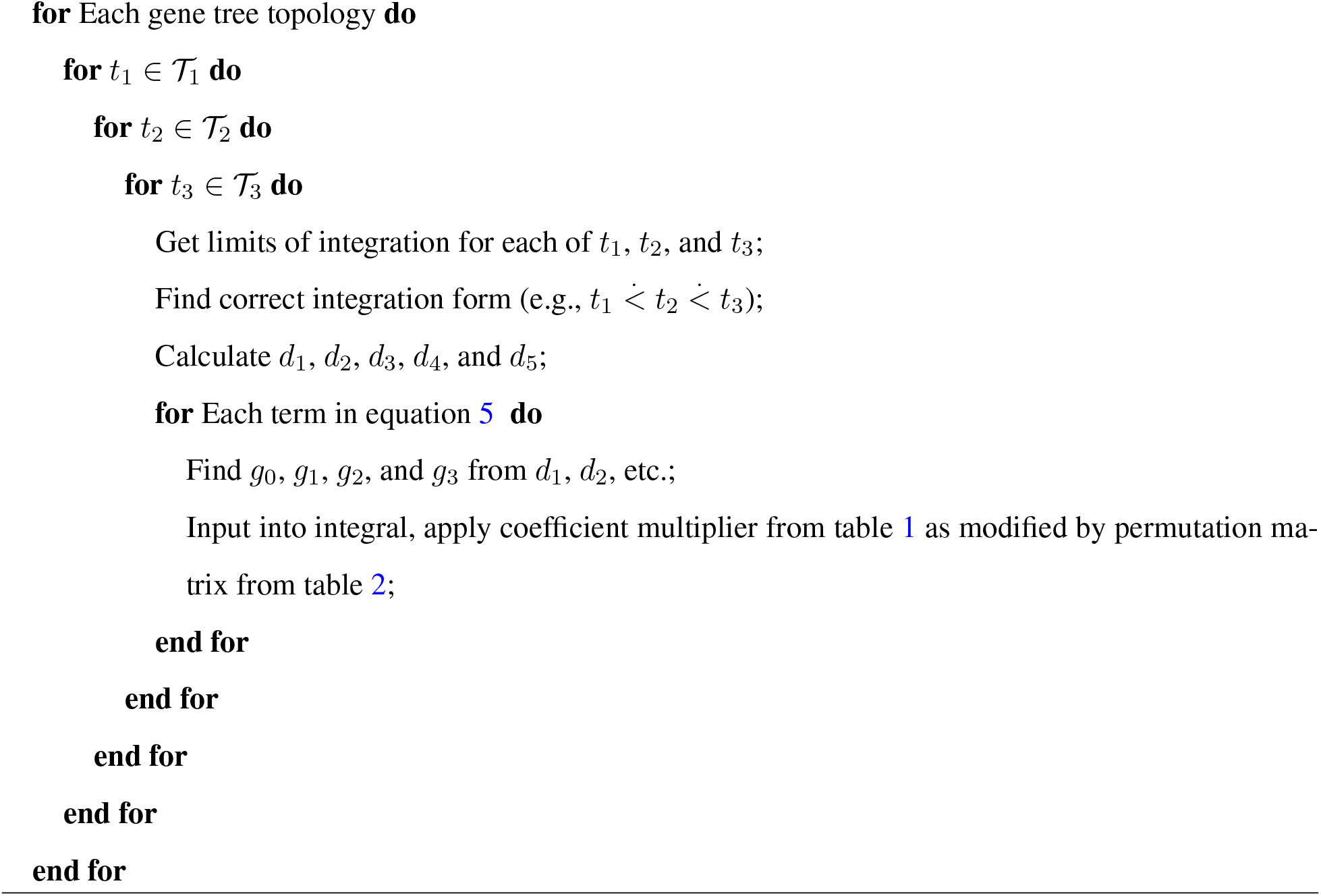

Again, we confirmed the accuracy of the code. First, we used a single rate category for each internal branch and verified that the program produced the same results as the program in section 3. Second, we modified the simulation code of Tian and Kubatko (2017) to have varying rates along the internal branches and performed goodness of fit tests to check that the simulation site pattern frequencies fit our calculated site pattern probabilities within the bounds of normal sampling error.

## 5 Quantifying impact of varying *γ* on site pattern probabilities

To begin to see the impact of different relative mutation rates, we start with the asymmetric species tree (*a* : 1.0, (*b* : 0.6, (*c* : 0.3, *d* : 0.3) : 0.3) : 0.4). We compare the site pattern probabilities under the strict molecular clock to the case where *γ*_*D*_ = 7.5. For simplicity, in this section, we will assume constant rates on internal branches as in section 3. The results can be seen in table 3. We can see that *δ*_*XXXX*_ decreases by −2.89*e* − 3 and *δ*_*XXXY*_ increases by the same amount. Similarly, the increase in *δ*_*XXYZ*_ is equal to the sum of decreases in *δ*_*XXYX*_ and *δ*_*XXYY*_ . These changes can be explained by an increase in the number of mutations of the nucleotide of species *d*, which makes sense given the increases in *γ*_*D*_ lengthens the pendant branch leading to species *d*. These changes in site pattern probabilities can be summarized by:

**Table 3:**
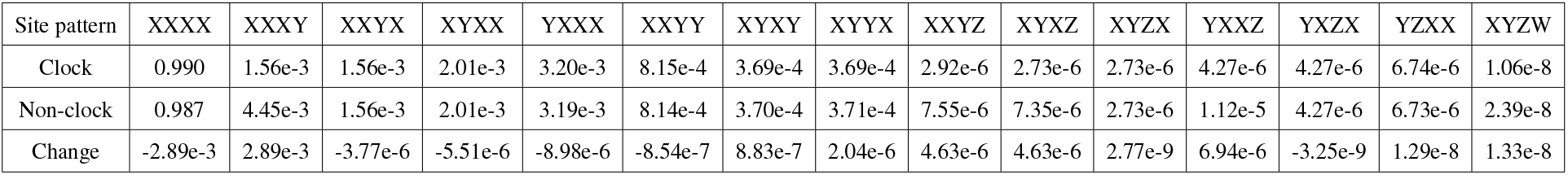
Effect of a change in *γ*_*D*_ from 1.0 to 7.5 on the site pattern probabilities for the tree (*a* : 1.0, (*b* : 0.6, (*c* : 0.3, *d* : 0.3) : 0.3) : 0.4).

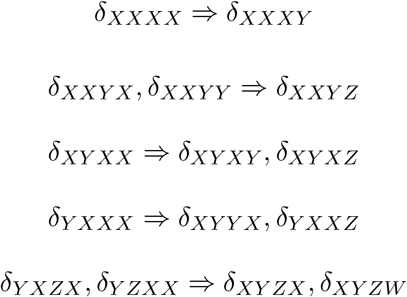

Next, we look at changes to *γ*_*D*_ along a spectrum of values for the species tree ((*a* : 0.6, *b* : 0.6) : 0.4, (*c* : 0.3, *d* : 0.3) : 0.7) as shown in figure 8. Again, we see less of the XXXX pattern and more of XYZW, which makes sense as increasing *γ*_*D*_ increases the total size of the tree. Patterns XXYY, XYXY, and XYYX appear to be largely unaffected. In the XXXY, XXYX, XYXX, and YXXX group, we see an increase to were XXXY overtakes XYXX/YXXX. We see a similar change in rank ordering in the other group of patterns. Because we only see a major change in two of the groupings, we focus on those in the remaining plots.

**Figure 8:**
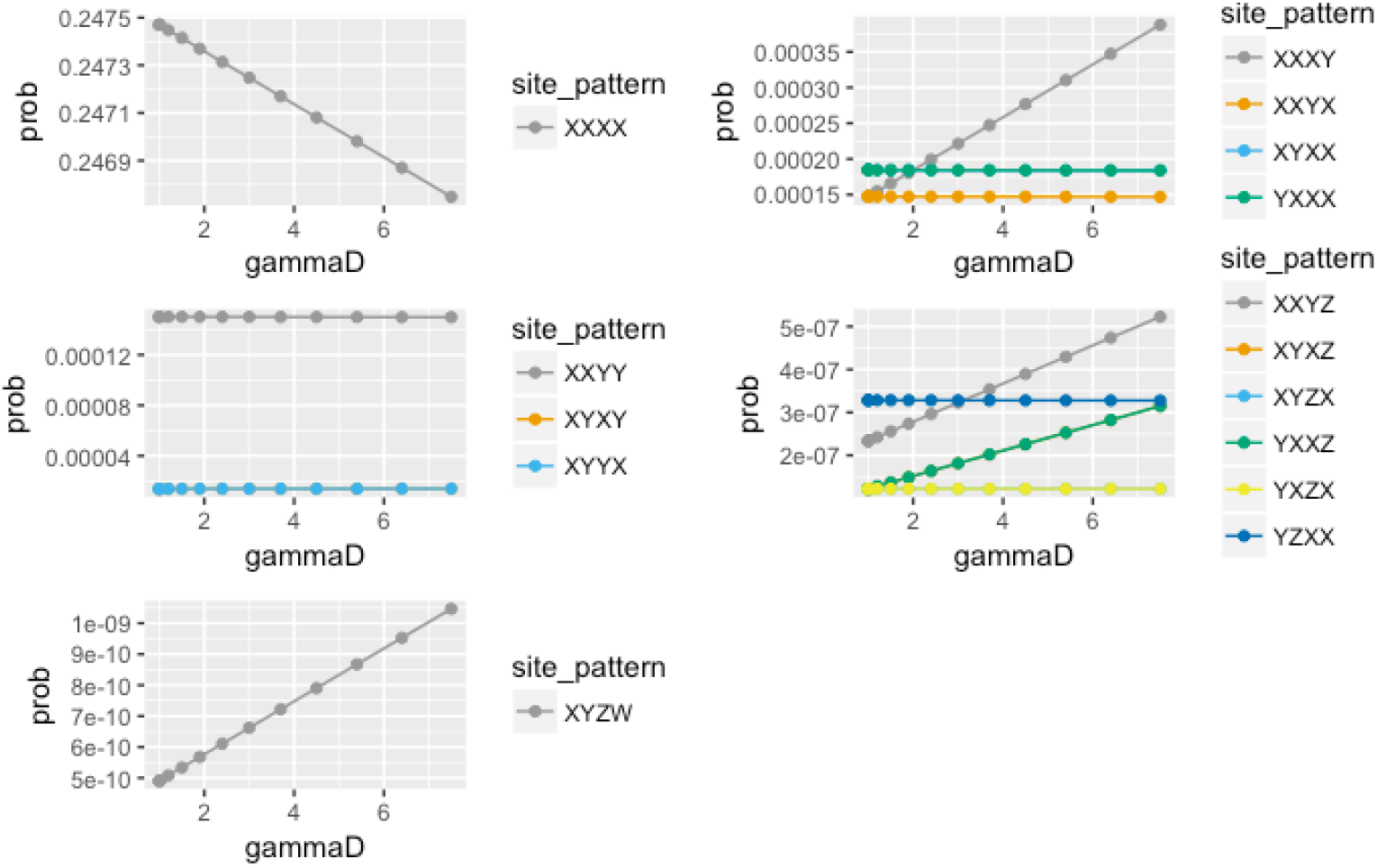
Effects of changing *γ*_*D*_ on species tree ((*a* : 0.6, *b* : 0.6) : 0.4, (*c* : 0.3, *d* : 0.3) : 0.7). Site patterns are grouped by similar magnitudes of the site pattern probabilities.

Figure 9 shows the effect of changing *γ*_*D*_ for different lengths of species *d*’s pendant branch. We see that the longer the pendant branch, the greater the impact of *γ*_*D*_ changes on the rank ordering of site pattern probabilities.

**Figure 9:**
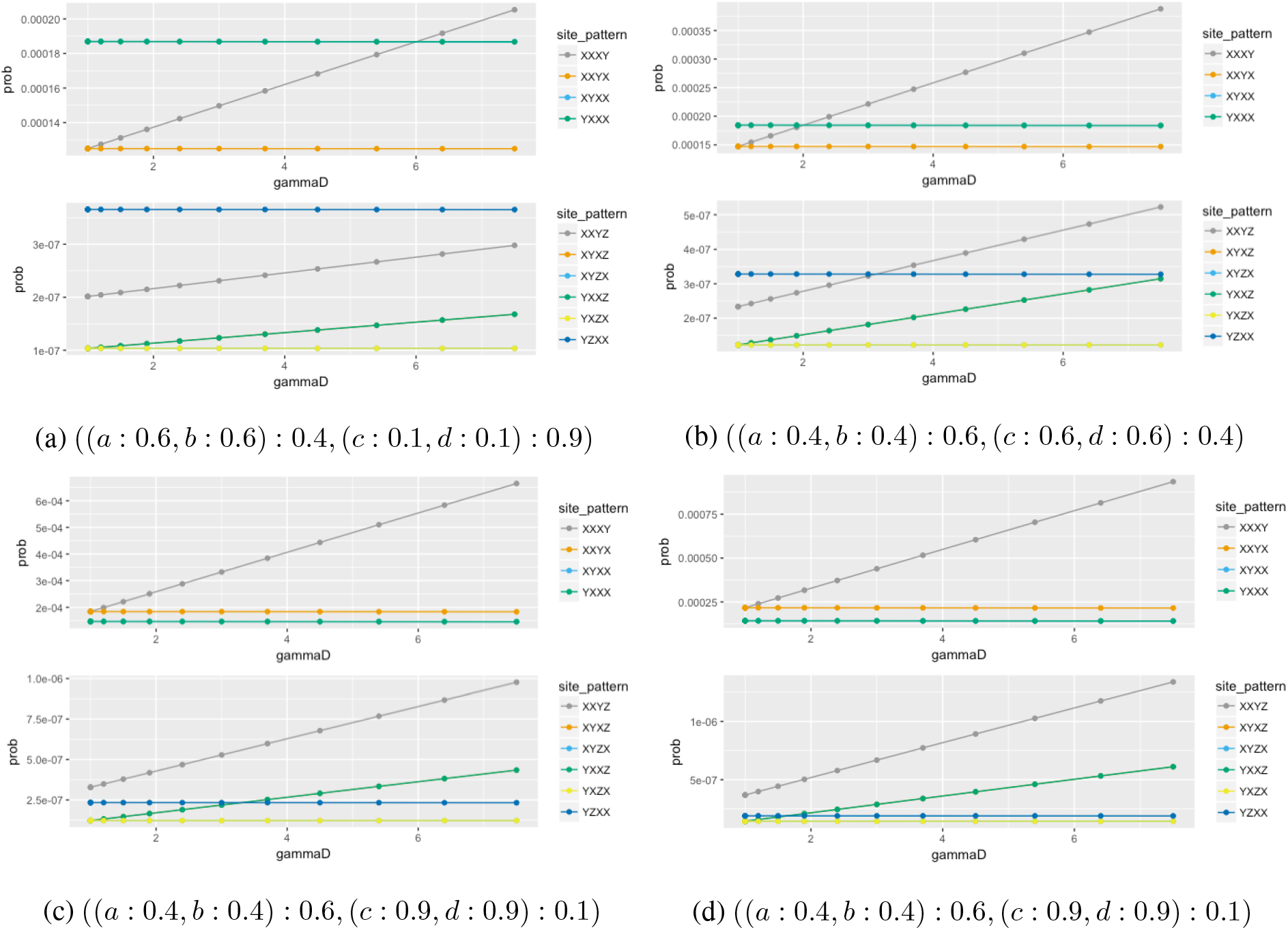
Interaction of branch length with *γ*_*D*_.

Figures 10 and 11 show the effect of changing the relative rates on different branches of the tree. For the symmetric tree ((*a* : 0.6, *b* : 0.6) : 0.4, (*c* : 0.3, *d* : 0.3) : 0.7), there is a slightly greater change to the site pattern probabilities for a change in *γ*_*CD*_ than for *γ*_*D*_. This tree does have a much longer internal vs. pendant branch (0.7 vs. 0.3 coalescent units) so that explains much of this difference. We also see that changing *γ*_*CD*_ increases both *δ*_*XXYX*_ and *δ*_*XXXY*_ whereas *γ*_*D*_ does not increase *δ*_*XXYX*_ . For the asymmetric tree (*a* : 1.0, (*b* : 0.6, (*c* : 0.3, *d* : 0.3) : 0.3) : 0.4), changing *γ*_*B*_ has a much greater impact on *δ*_*XYXX*_ than changing *γ*_*D*_ has on *δ*_*XXXY*_ . It is somewhat interesting that changing the branch on the internal rates *γ*_*CD*_ or *γ*_*BCD*_ causes as quadratic rather than a linear response on some of the site pattern probabilities such as *δ*_*XXYZ*_, *δ*_*YZXX*_. The deviation from linearity is small and we have no intuition for why the response is different, but it is worth noting.

**Figure 10:**
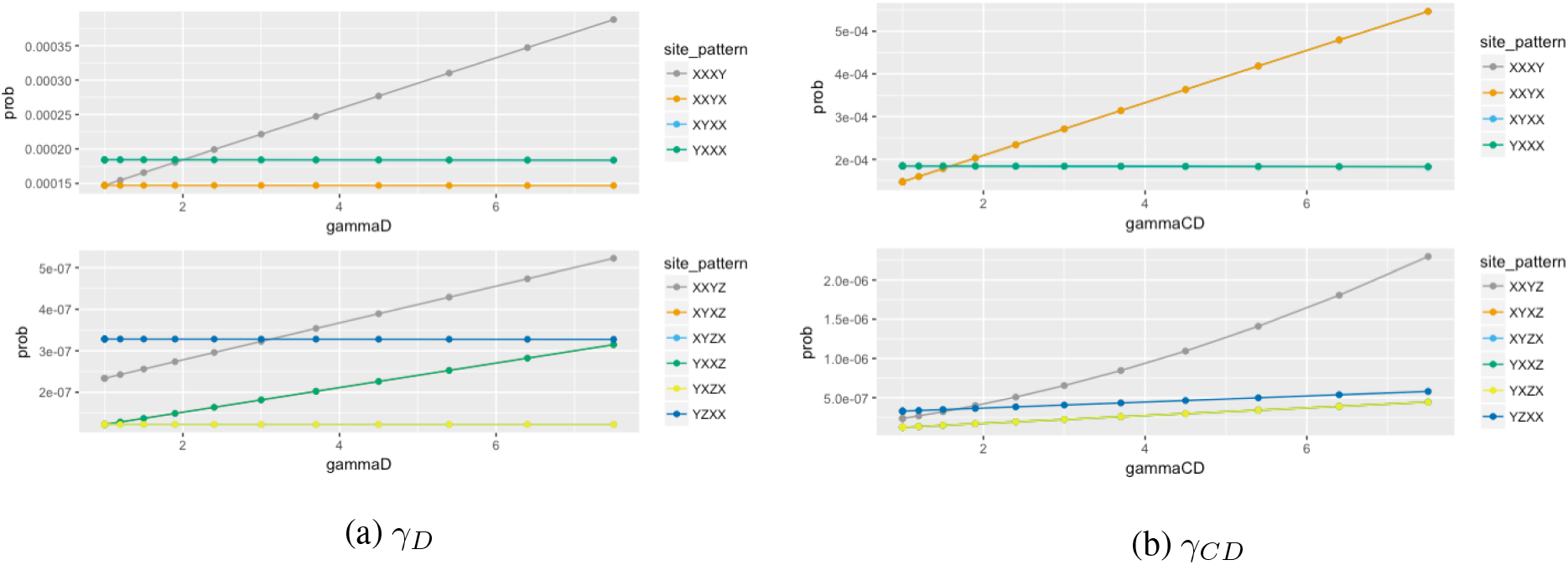
Effect of changing the relative rate on a pendant vs. an internal branch for the symmetric species tree: ((*a* : 0.6, *b* : 0.6) : 0.4, (*c* : 0.3, *d* : 0.3) : 0.7)

**Figure 11:**
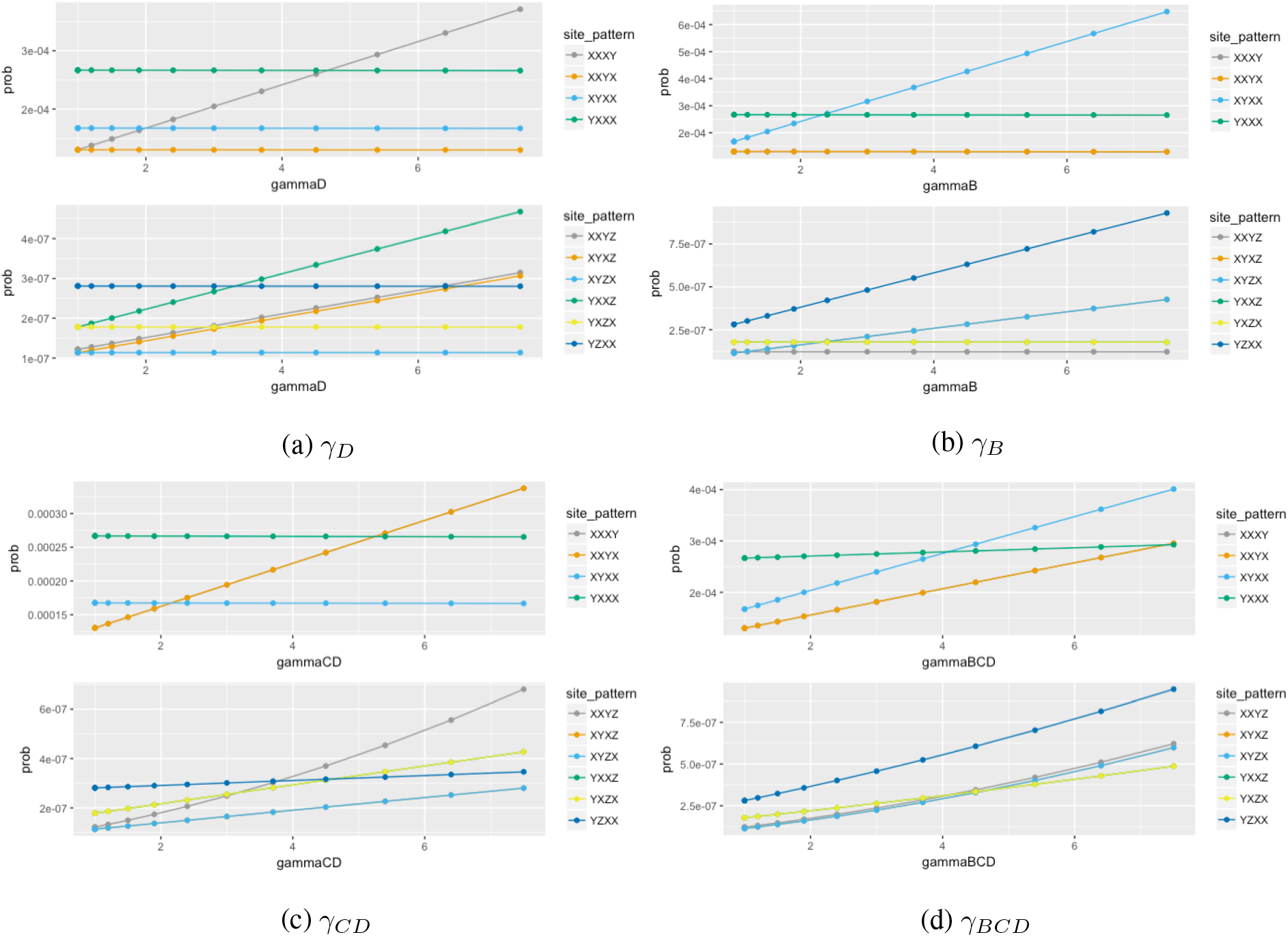
Effect of changing the relative rate on various branches for the asymmetric species tree: (*a* : 1.0, (*b* : 0.6, (*c* : 0.3, *d* : 0.3) : 0.3) : 0.4)

### 5.1 Applications

In Richards and Kubatko (2021), it was shown that their method, Lily-T, would systematically infer the incorrect tree if the relative rates a branch were too high, with greater sensitivity to changes on the pendant branches. We considered another way in which clock violations could interfere with inference. Methods such as ABBA-BABA [6] and HyDe [10] use the relative frequencies of the site patterns XXYY, XYXY, and XYYX to infer whether or not hybridization occurs. The tests compare the hypotheses

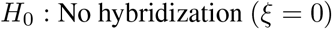

and

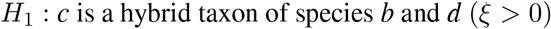

using test statistics that follow the standard normal distribution. Surprisingly, changing the relative rate of any branch had little effect on inference. Even with the tree (*a* : 1.0, (*b* : 0.6, (*c* : 0.5, *d* : 0.5) : 0.1) : 0.4), *γ*_*D*_ = 7.5 and 240,000 sites, the expected value of the test statistic only moved from 0 to 0.03. In fact, we had to change the tree to (*a* : 10.0, (*b* : 9.5, (*c* : 9.0, *d* : 9.0) : 0.5) : 0.5), *γ*_*D*_ = 10.0, and 2.4 million sites before we would expect to falsely reject the null hypothesis of no hybridization.

So why is the Lily procedure so sensitive to violations of the clock where the ABBA-BABA and HyDe tests are relatively robust? The difference lies in the fact that the relative frequencies of {*XXXY, XXYX, XYXX, YXXX*} have a large contribution to the likelihood calculations Lily uses to make species tree inference. By contrast, as we saw in figure 8 and table 3 the relative frequencies of {*XXYY, XYXY, XYYX*} are less sensitive to violations of the strict molecular clock, so methods that only depend on the relative frequencies of these three site patterns are more robust against clock violations.

## 6 Conclusions and future work

In this paper we provide complete site pattern probabilities under the multispecies coalescent model and the Jukes-Cantor substitution model, allowing for differing mutation rates in different branches of the species tree. Our work opens up two new avenues of research. The first is in expanding the probability calculations performed here to higher numbers of taxa, either with or without the molecular clock. Although the theoretical basis for this extension is well-known, the extensive computation required has prevented this in practice. Our framework for breaking down the site pattern probabilities arising from equation 3 can be easily expanded to higher numbers of taxa, and we expect that when the gene tree histories are enumerated we would again see a small enough number of integral forms and combinations of inputs as in table 4 to enable direct calculation of the probabilities.

**Table 4:** Symmetric species tree gene histories

| Index | Gene Tree History Topology | Time constraints | Integral Form |
| --- | --- | --- | --- |
| 1 | $((A, B), (C, D))$ | $t_1 < \tau_3, t_2 < \tau_3$ | A |
| 2 | $((A, B), (C, D))$ | $t_1 < \tau_3 < t_2$ | B |
| 3 | $((A, B), (C, D))$ | $t_2 < \tau_3 < t_1$ | C |
| 4 | $((A, B), (C, D))$ | $\tau_3 < t_1 < t_2$ | D |
| 5 | $((A, B), (C, D))$ | $\tau_3 < t_2 < t_1$ | E |
| 6 | $((A, C), (B, D))$ | $\tau_3 < t_1 < t_2$ | D |
| 7 | $((A, C), (B, D))$ | $\tau_3 < t_2 < t_1$ | E |
| 8 | $((A, D), (B, C))$ | $\tau_3 < t_1 < t_2$ | D |
| 9 | $((A, D), (B, C))$ | $\tau_3 < t_2 < t_1$ | E |
| 10 | $(A, (B, (C, D)))$ | $t_1 < \tau_3 < t_2$ | B |
| 11 | $(A, (B, (C, D)))$ | $\tau_3 < t_1 < t_2$ | D |
| 12 | $(A, (C, (B, D)))$ | $\tau_3 < t_1 < t_2$ | D |
| 13 | $(A, (D, (B, C)))$ | $\tau_3 < t_1 < t_2$ | D |
| 14 | $(B, (A, (C, D)))$ | $t_1 < \tau_3 < t_2$ | B |
| 15 | $(B, (A, (C, D)))$ | $\tau_3 < t_1 < t_2$ | D |
| 16 | $(B, (C, (A, D)))$ | $\tau_3 < t_1 < t_2$ | D |
| 17 | $(B, (D, (A, C)))$ | $\tau_3 < t_1 < t_2$ | D |
| 18 | $(C, (A, (B, D)))$ | $\tau_3 < t_1 < t_2$ | D |
| 19 | $(C, (B, (A, D)))$ | $\tau_3 < t_1 < t_2$ | D |
| 20 | $(C, (D, (A, B)))$ | $t_1 < \tau_3 < t_2$ | F |
| 21 | $(C, (D, (A, B)))$ | $\tau_3 < t_1 < t_2$ | D |
| 22 | $(D, (A, (B, C)))$ | $\tau_3 < t_1 < t_2$ | D |
| 23 | $(D, (B, (A, C)))$ | $\tau_3 < t_1 < t_2$ | D |
| 24 | $(D, (C, (A, B)))$ | $t_1 < \tau_3 < t_2$ | F |
| 25 | $(D, (C, (A, B)))$ | $\tau_3 < t_1 < t_2$ | D |

**Table 5:** Asymmetric species tree gene histories, pt.1

| Index | Gene History Topology | Time constraints | Integral Form |
| --- | --- | --- | --- |
| 1 | $((A, B), (C, D))$ | $t_1 < \tau_2 < \tau_3 < t_2$ | A |
| 2 | $((A, B), (C, D))$ | $\tau_2 < t_1 < \tau_3 < t_2$ | B |
| 3 | $((A, B), (C, D))$ | $\tau_3 < t_1 < t_2$ | C |
| 4 | $((A, B), (C, D))$ | $\tau_3 < t_2 < t_1$ | D |
| 5 | $((A, C), (B, D))$ | $\tau_2 < t_1 < \tau_3 < t_2$ | B |
| 6 | $((A, C), (B, D))$ | $\tau_3 < t_1 < t_2$ | C |
| 7 | $((A, C), (B, D))$ | $\tau_3 < t_2 < t_1$ | D |
| 8 | $((A, D), (B, C))$ | $\tau_2 < t_1 < \tau_3 < t_2$ | B |
| 9 | $((A, D), (B, C))$ | $\tau_3 < t_1 < t_2$ | C |
| 10 | $((A, D), (B, C))$ | $\tau_3 < t_2 < t_1$ | D |
| 11 | $(A, (B, (C, D)))$ | $t_1 < \tau_2 < t_2 < \tau_3$ | E |
| 12 | $(A, (B, (C, D)))$ | $t_1 < \tau_2 < \tau_3 < t_2$ | A |
| 13 | $(A, (B, (C, D)))$ | $\tau_2 < t_1 < t_2 < \tau_3$ | F |
| 14 | $(A, (B, (C, D)))$ | $\tau_2 < t_1 < \tau_3 < t_2$ | B |
| 15 | $(A, (B, (C, D)))$ | $\tau_3 < t_1$ | C |
| 16 | $(A, (C, (B, D)))$ | $\tau_2 < t_1 < t_2 < \tau_3$ | F |
| 17 | $(A, (C, (B, D)))$ | $\tau_2 < t_1 < \tau_3 < t_2$ | B |
| 18 | $(A, (C, (B, D)))$ | $\tau_3 < t_1$ | C |
| 19 | $(A, (D, (B, C)))$ | $\tau_2 < t_1 < t_2 < \tau_3$ | F |
| 20 | $(A, (D, (B, C)))$ | $\tau_2 < t_1 < \tau_3 < t_2$ | B |
| 21 | $(A, (D, (B, C)))$ | $\tau_3 < t_1$ | C |
| 22 | $(B, (A, (C, D)))$ | $t_1 < \tau_2 < \tau_3 < t_2$ | A |
| 23 | $(B, (A, (C, D)))$ | $\tau_2 < t_1 < \tau_3 < t_2$ | B |
| 24 | $(B, (A, (C, D)))$ | $\tau_3 < t_1$ | C |
| 25 | $(B, (C, (A, D)))$ | $\tau_3 < t_1$ | C |
| 26 | $(B, (D, (A, C)))$ | $\tau_3 < t_1$ | C |

**Table 6:** Asymmetric species tree gene histories, pt.2

| Index | Topology | Time constraints | Integral Form |
| --- | --- | --- | --- |
| 27 | $(C, (A, (B, D)))$ | $\tau_2 < t_1 < \tau_3 < t_2$ | B |
| 28 | $(C, (A, (B, D)))$ | $\tau_3 < t_1$ | C |
| 29 | $(C, (B, (A, D)))$ | $\tau_3 < t_1$ | C |
| 30 | $(C, (D, (A, B)))$ | $\tau_3 < t_1$ | C |
| 31 | $(D, (A, (B, C)))$ | $\tau_2 < t_1 < \tau_3 < t_2$ | B |
| 32 | $(D, (A, (B, C)))$ | $\tau_3 < t_1$ | C |
| 33 | $(D, (B, (A, C)))$ | $\tau_3 < t_1$ | C |
| 34 | $(D, (C, (A, B)))$ | $\tau_3 < t_1$ | C |

This would open up improvements on existing assembly methods for species tree inference. Assembly methods such as [1], [21], [24], [25], [29], [34], [23], [26], [31], and [30] begin with a triplet or quartet of species, infer the topology of these quartets, and use them as inputs to an assembly algorithm. These inputs are treated as independent, although for an n-taxon tree there is statistical dependence due to any branches the quartets share in common. Expanding the exact site pattern probabilities to 5 or 6 taxa would allow at least pairwise statistical dependence between triplets to be determined, which should improve both point estimation and uncertainty quantification.

It is worth noting that the invariants we see with gene trees do not occur with species trees under the multivariate coalescent. For example, for the gene tree ((*A, B*), (*C, D*)) the two sides of the tree are independent such that *P* (*i*_*A*_ = *i*_*B*_ ∩ *i*_*C*_ = *i*_*D*_) = *P* (*i*_*A*_ = *i*_*B*_) × *P* (*i*_*C*_ = *i*_*D*_). This does not hold for the species tree ((*a, b*), (*c, d*)) even under the molecular clock. (This can be shown from the site pattern probabilities in the supplemental material of [3] by marginalizing over the taxa). This is perhaps counterintuitive, but arises out of the fact that for any finite *τ*_3_, there is some possibility that the gene tree and species tree don’t match, and therefore there are some sites where the characters on either side of the root of the species tree are no longer independent (because they arise from gene trees whose tips are no longer independent).

The results of this paper should also enable improving quartet inference. Lily-Q [19] is a procedure that takes *P* (***D*** = ***d***|(*S*, ***τ***)) and assuming a distribution on (*S*, ***τ***), produces posterior probabilities on *S* for groups of 4 taxa. Given either a distribution on ***γ*** or a means of estimating 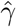, we should be able to develop an extension on Lily-Q to the relaxed-clock case, providing a means for fast, accurate species tree quartet inference in cases where the molecular clock does not hold.

## 7 Appendix

### 7.1 Gene tree histories and integrals – symmetric case

#### Form A

Density: 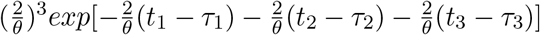

Integral: 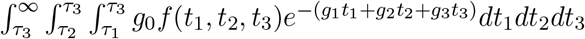

Result: 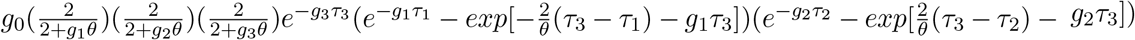

#### Form B

Density: 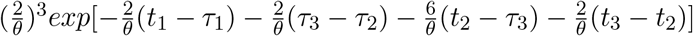

Integral: 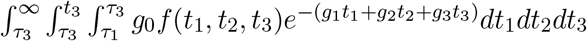

Result: 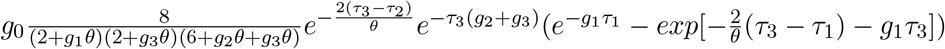

#### Form C

Density: 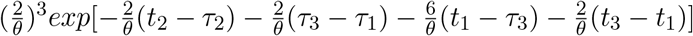

Integral: 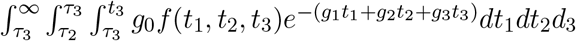

Result: 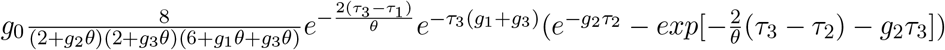

#### Form D

Density: 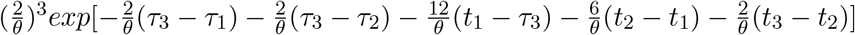

Integral: 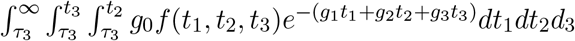

Result: 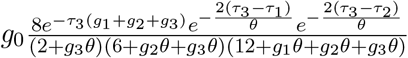

#### Form E

Density: 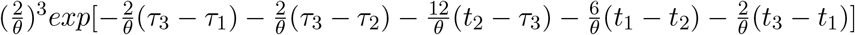

Integral: 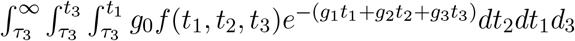

Result: 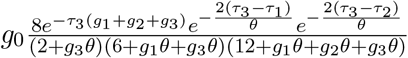

#### Form F

Density: 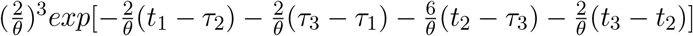

Integral: 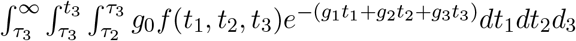

Result: 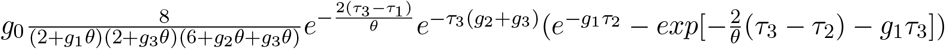

### 7.2 Gene tree histories and integrals – asymmetric case

#### Form A

Density: 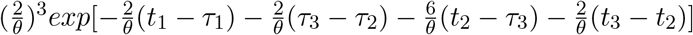

Integral: 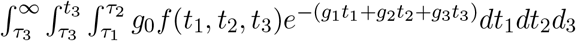

Result: 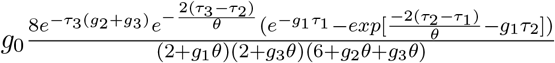

#### Form B

Density: 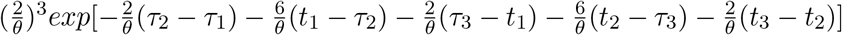

Integral: 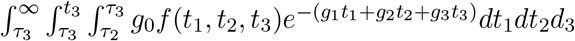

Result: 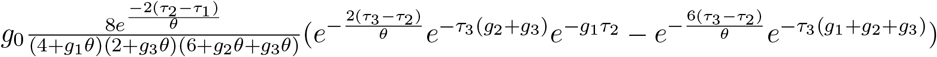

#### Form C

Density: 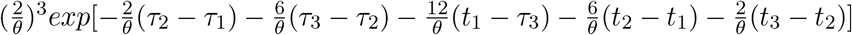

Integral: 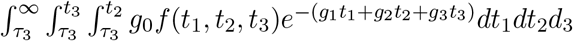

Result: 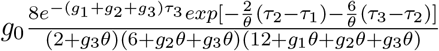

#### Form D

Density: 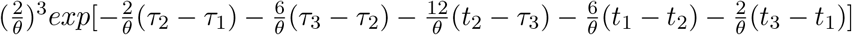

Integral: 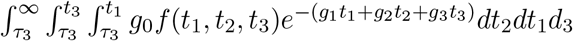

Result: 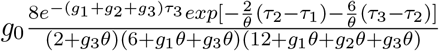

#### Form E

Density: 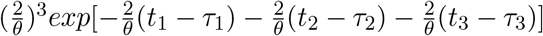

Integral: 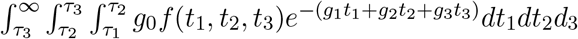

Result: 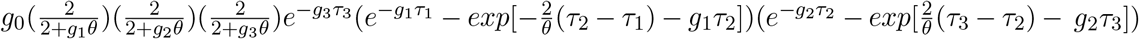

#### Form F

Density: 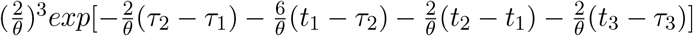

Integral: 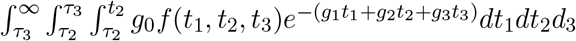

Result: 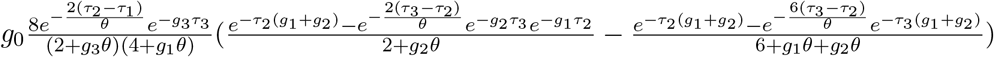

### 7.3 General gene density integrals

#### 7.3.1 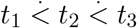

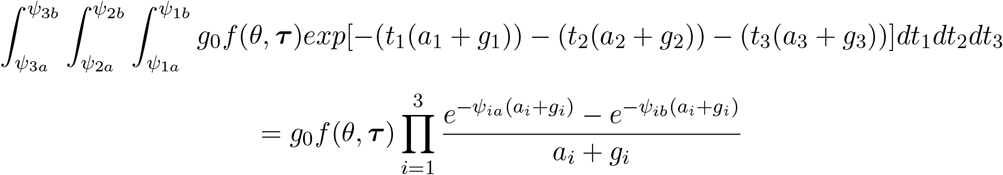

#### 7.3.2 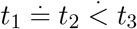

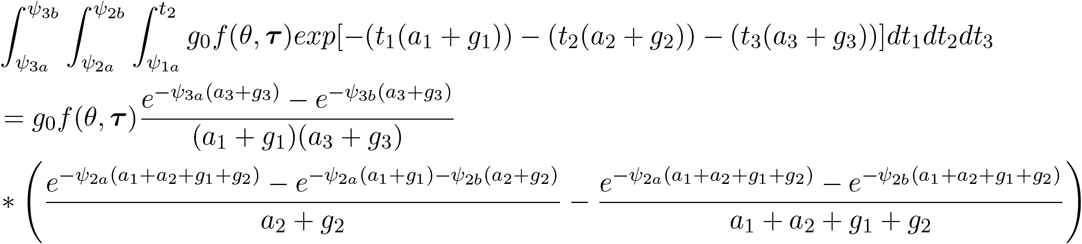

#### 7.3.3 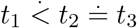

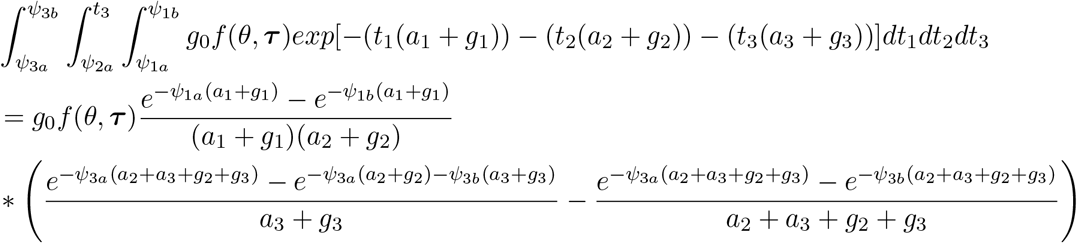

#### 7.3.4 *t*_1_ ≐ *t*_2_ ≐ *t*_3_

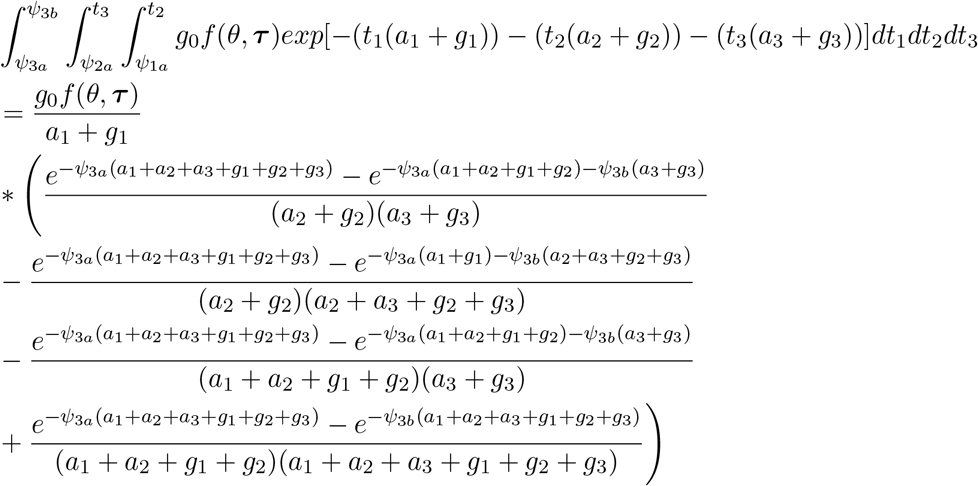

## Notes

### Competing Interest Statement

The authors have declared no competing interest.

