## Supplemental Material for "Site Pattern Probabilities Under the Multispecies Coalescent and a Relaxed Molecular Clock: Theory and Applications"

### 1 Final probability calculations

Upon reviewing the tables in the appendix, there are only a limited number of possible combinations of $g_1, g_2, g_3$. The base cases are given in table 1.

| **Case** | **Value** |
|---|---|
| 1 | $g_1 = 2\alpha$ |
| 2 | $g_2 = 2\alpha$ |
| 3 | $g_3 = 2\alpha$ |
| 4 | $g_1 = \alpha, g_2 = 2\alpha$ |
| 5 | $g_1 = \alpha, g_3 = 2\alpha$ |
| 6 | $g_2 = \alpha, g_3 = 2\alpha$ |
| 7 | $g_1 = \alpha, g_2 = \alpha, g_3 = 2\alpha$ |
| 8 | $g_1 = 2\alpha, g_2 = 2\alpha$ |
| 9 | $g_1 = 2\alpha, g_3 = 2\alpha$ |

Table 1: Possible $g$ values, base case

Adjustments to these general cases are made for certain gene tree histories. For example, in the symmetric case in all gene tree histories using integral form A, $\alpha$ is replaced with $\alpha\gamma_{CD}$ for $g_1$ and $\alpha$ is replaced with $\alpha\gamma_{AB}$ for $g_2$. The full set of adjustments is given in table 2. Also, note that not all of the base cases occur with all integral forms. For example, $g_1 = 2\alpha, g_2 = 2\alpha$ does not occur with integral form F in the symmetric case. The final output of the equation

$$\boldsymbol{\delta}|(S, \boldsymbol{\tau}, \boldsymbol{\gamma}, \theta) = \sum_{G \in \mathcal{G}_S} \int_{\boldsymbol{t}} \boldsymbol{\delta}|(G, \boldsymbol{t}, \boldsymbol{\gamma}) f((G, \mathbf{t})|(S, \boldsymbol{\tau}, \theta)) d\mathbf{t}$$

is then given below in order: case 1, integral form A; case 1, integral form B; all the way to case 9, integral form F. It is worth noting that if the molecular clock holds, all $h$ terms are unity, and the formulas below reduce, with much algebraic simplification, down to the formulae given in [1].

| Species tree topology | Integral Form | Adjustment |
|---|---|---|
| Symmetric | A | $g_1 : \alpha \to \alpha\gamma_{CD}, g_2 : \alpha \to \alpha\gamma_{AB}$ |
| Symmetric | B | $g_1 : \alpha \to \alpha\gamma_{CD}$ |
| Symmetric | C | $g_2 : \alpha \to \alpha\gamma_{AB}$ |
| Symmetric | F | $g_1 : \alpha \to \alpha\gamma_{AB}$ |
| Asymmetric | A | $g_1 : \alpha \to \alpha\gamma_{CD}$ |
| Asymmetric | B | $g_1 : \alpha \to \alpha\gamma_{BCD}$ |
| Asymmetric | E | $g_1 : \alpha \to \alpha\gamma_{CD}, g_2 : \alpha \to \alpha\gamma_{BCD}$ |
| Asymmetric | F | $g_1, g_2 : \alpha \to \alpha\gamma_{BCD}$ |

Table 2: $g$ values, adjustments

#### 1.1 Symmetric case

Let

$$x = exp[-\frac{2(\tau_3 - \tau_1)}{\theta}], y = exp[-\frac{2(\tau_3 - \tau_2)}{\theta}], a = e^{-\alpha\tau_1}, b = e^{-\alpha\tau_2}, c = e^{-\alpha\tau_3}$$

$$h_1 = e^{-\alpha\tau_1(\gamma_C - \gamma_{CD})}, h_2 = e^{-\alpha\tau_1(\gamma_D - \gamma_{CD})}, h_3 = e^{-\alpha\tau_2(\gamma_B - \gamma_{AB})}$$

$$h_4 = e^{-\alpha\tau_2(\gamma_B - \gamma_{AB})}, h_5 = e^{-\alpha\tau_3(\gamma_{AB} - 1)}, h_6 = e^{-\alpha\tau_3(\gamma_{CD} - 1)}$$

Then each site pattern is of the form:

$$\frac{1}{256}[1 + \frac{k_1 h_1 h_2}{1 + \alpha\gamma_{CD}\theta}(a^{2\gamma_{CD}} - c^{2\gamma_{CD}}x)(1-y) + \frac{k_2 h_1 h_2}{3(1 + \alpha\gamma_{CD}\theta)}y(a^{2\gamma_{CD}} - c^{2\gamma_{CD}}x) + \frac{k_3 h_1 h_6^2}{3 + \alpha\theta}c^2 x(1-y)$$

$$+\frac{k_4 h_1 h_2 h_6^2 + k_5 h_1 h_3 h_5 h_6 + k_6 h_1 h_4 h_5 h_6 + k_7 h_2 h_3 h_5 h_6 + k_8 h_2 h_4 h_5 h_6 + k_9 h_3 h_4 h_5^2}{3(6 + \alpha\theta)}c^2 xy$$

$$+\frac{k_{10} h_1 h_2 h_6^2 + k_{11} h_1 h_4 h_5 h_6 + k_{12} h_2 h_4 h_5 h_6}{(3 + \alpha\theta)(6 + \alpha\theta)}c^2 xy + \frac{k_{13} h_3 h_4}{3(1 + \alpha\gamma_{AB}\theta)}x(b^{2\gamma_{AB}} - c^{2\gamma_{AB}}y)$$

$$\frac{k_{14} h_3 h_4}{1 + \alpha\gamma_{AB}\theta}(1-x)(b^{2\gamma_{AB}} - c^{2\gamma_{AB}}y)$$

$$+\frac{k_{15} h_3 h_4 h_5^2 + k_{16} h_1 h_3 h_5 h_6 + k_{17} h_1 h_4 h_5 h_6 + k_{18} h_2 h_3 h_5 h_6 + k_{19} h_2 h_4 h_5 h_6}{3(1 + \alpha\theta)}c^2 y(1-x)$$

$$\frac{k_{20}h_3h_4}{3(1+\alpha\gamma_{AB}\theta)}x(b^{2\gamma_{AB}}-c^{2\gamma_{AB}}y)$$

$$+\frac{k_{21}h_1h_2h_6^2+k_{22}h_1h_3h_5h_6+k_{23}h_1h_4h_5h_6+k_{24}h_2h_3h_5h_6+k_{25}h_2h_4h_5h_6+k_{26}h_3h_4h_5^2}{(3+\alpha\theta)(6+\alpha\theta)}c^2xy$$

$$+\frac{k_{27}h_3h_4h_5^2+k_{28}h_1h_3h_5h_6+k_{29}h_2h_3h_5h_6}{3(6+\alpha\theta)}c^2xy$$

$$+\frac{k_{30}h_1h_3h_5h_6+k_{31}h_1h_4h_5h_6+k_{32}h_2h_3h_5h_6+k_{33}h_2h_4h_5h_6}{3+\alpha\theta}c^2x(1-y)$$

$$+\frac{k_{34}h_1h_3h_5h_6+k_{35}h_1h_4h_5h_6+k_{36}h_2h_3h_5h_6+k_{37}h_2h_4h_5h_6}{1+\alpha\theta}c^2(1-x)(1-y)$$

$$+\frac{k_{38}h_1h_3h_5h_6+k_{39}h_1h_4h_5h_6+k_{40}h_2h_3h_5h_6+k_{41}h_2h_4h_5h_6+k_{42}h_3h_4h_5^2}{(1+\alpha\theta)(3+\alpha\theta)}c^2y(1-x)$$

$$+\frac{k_{43}h_1h_3h_5h_6+k_{44}h_1h_4h_5h_6+k_{45}h_2h_3h_5h_6+k_{46}h_2h_4h_5h_6}{(1+\alpha\theta)(3+\alpha\theta)}c^2x(1-y)$$

$$+\frac{k_{47}h_1h_3h_5h_6+k_{48}h_1h_4h_5h_6+k_{49}h_2h_3h_5h_6+k_{50}h_2h_4h_5h_6+k_{51}h_1h_2h_6^2+k_{52}h_3h_4h_5^2}{(1+\alpha\theta)(3+\alpha\theta)(6+\alpha\theta)}c^2xy$$

$$+\frac{k_{53}h_1h_3h_5h_6+k_{54}h_1h_4h_5h_6+k_{55}h_2h_3h_5h_6+k_{56}h_2h_4h_5h_6+k_{57}h_1h_2h_6^2+k_{58}h_3h_4h_5^2}{(1+\alpha\theta)(3+\alpha\theta)(6+\alpha\theta)}c^2xy$$

$$+\frac{k_{59}h_1h_3h_5h_6+k_{60}h_1h_4h_5h_6+k_{61}h_2h_3h_5h_6+k_{62}h_2h_4h_5h_6+k_{63}h_1h_2h_6^2}{(1+\alpha\theta)(3+\alpha\theta)}c^2x(1-y)$$

$$+\frac{2(k_{64}h_1h_2h_3h_5h_6+k_{65}h_1h_2h_4h_5h_6)}{(2+\alpha\gamma_{CD}\theta)(3+\alpha\theta)}c^2y(a^{\gamma_{CD}}-c^{\gamma_{CD}}x)$$

$$+\frac{2(k_{66}h_1h_2h_3h_5h_6^2+k_{67}h_1h_2h_4h_5h_6^2+k_{68}h_1h_3h_4h_5^2h_6+k_{69}h_2h_3h_4h_5^2h_6)}{3(3+\alpha\theta)(4+\alpha\theta)}c^3xy$$

$$+\frac{2(k_{70}h_1h_3h_4h_5h_6+k_{71}h_2h_3h_4h_5h_6)}{(2+\alpha\gamma_{AB}\theta)(3+\alpha\theta)}c^2x(b^{\gamma_{AB}}-b^{\gamma_{AB}}y)$$

$$+\frac{2(k_{72}h_1h_2h_3h_5h_6+k_{73}h_1h_2h_4h_5h_6)}{(2+\alpha\gamma_{CD}\theta)(1+\alpha\theta)}c^2(a^{\gamma_{CD}}-c^{\gamma_{CD}}x)(1-y)$$

$$+\frac{2(k_{74}h_1h_2h_3h_5h_6+k_{75}h_1h_2h_4h_5h_6)}{(2+\alpha\gamma_{CD}\theta)(1+\alpha\theta)(3+\alpha\theta)}c^2y(a^{\gamma_{CD}}-c^{\gamma_{CD}}x)$$

$$+\frac{2(k_{76}h_1h_2h_3h_5h_6^2+k_{77}h_1h_2h_4h_5h_6^2)}{3(1+\alpha\theta)(2+\alpha\theta)}c^3x(1-y)$$

$$+\frac{2(k_{78}h_1h_2h_3h_5h_6^2+k_{79}h_1h_2h_4h_5h_6^2+k_{80}h_1h_3h_4h_5^2h_6+k_{81}h_2h_3h_4h_5^2h_6)}{3(1+\alpha\theta)(3+\alpha\theta)(4+\alpha\theta)}c^3xy$$

$$+\frac{4(k_{82}h_1h_2h_3h_5h_6^2+k_{83}h_1h_2h_4h_5h_6^2+k_{84}h_1h_3h_4h_5^2h_6+k_{85}h_2h_3h_4h_5^2h_6)}{9(1+\alpha\theta)(2+\alpha\theta)(4+\alpha\theta)}c^3xy$$

$$+\frac{2(k_{86}h_1h_3h_4h_5h_6+k_{87}h_2h_3h_4h_5h_6)}{(2+\alpha\gamma_{AB}\theta)(1+\alpha\theta)(3+\alpha\theta)}c^2x(b^{\gamma_{AB}}-c^{\gamma_{AB}}y)$$

$$+\frac{2(k_{88}h_1h_3h_4h_5h_6+k_{89}h_2h_3h_4h_5h_6)}{(2+\alpha\gamma_{AB}\theta)(1+\alpha\theta)}c^2(1-x)(b^{\gamma_{AB}}-c^{\gamma_{AB}}y)$$

$$+\frac{2(k_{90}h_1h_3h_4h_5^2h_6+k_{91}h_2h_3h_4h_5^2h_6)}{3(1+\alpha\theta)(2+\alpha\theta)}c^3y(1-x)$$

$$+\frac{2(k_{92}h_1h_3h_4h_5h_6+k_{93}h_2h_3h_4h_5h_6)}{(1+\alpha\theta)(2+\alpha\gamma_{AB}\theta)(3+\alpha\theta)}c^2x(b^{\gamma_{AB}}-c^{\gamma_{AB}}y)$$

$$+\frac{4(k_{94}h_1h_2h_3h_5h_6^2+k_{95}h_1h_2h_4h_5h_6^2+k_{96}h_1h_3h_4h_5^2h_6+k_{97}h_2h_3h_4h_5^2h_6)}{9(1+\alpha\theta)(2+\alpha\theta)(4+\alpha\theta)}c^3xy$$

$$+\frac{2(k_{98}h_1h_2h_3h_5h_6^2+k_{99}h_1h_3h_4h_5^2h_6+k_{100}h_2h_3h_4h_5^2h_6)}{3(1+\alpha\theta)(3+\alpha\theta)(4+\alpha\theta)}c^3xy$$

$$+\frac{2(k_{101}h_1h_2h_3h_5h_6^2+k_{102}h_1h_2h_4h_5h_6^2)}{3(1+\alpha\theta)(2+\alpha\theta)}c^3x(1-y)$$

$$+\frac{4(k_{103}h_1h_2h_3h_4h_5h_6)}{(1+\alpha\theta)(2+\alpha\gamma_{CD}\theta)(2+\alpha\gamma_{AB}\theta)}c^2(a^{\gamma_{CD}}-c^{\gamma_{CD}}x)(b^{\gamma_{AB}}-c^{\gamma_{AB}}y)$$

$$+\frac{4(k_{104}h_1h_2h_3h_4h_5^2h_6)}{3(1+\alpha\theta)(2+\alpha\gamma_{CD}\theta)(2+\alpha\theta)}c^3y(a^{\gamma_{CD}}-c^{\gamma_{CD}}x)$$

$$+\frac{4(k_{105}h_1h_2h_3h_4h_5h_6^2)}{3(1+\alpha\theta)(2+\alpha\gamma_{AB}\theta)(2+\alpha\theta)}c^3x(b^{\gamma_{AB}}-c^{\gamma_{AB}}y)$$

$$+\frac{k_{106}h_1h_2h_3h_4h_5^2h_6^2+k_{107}h_1h_2h_3h_4h_5^2h_6^2}{3(1+\alpha\theta)(2+\alpha\theta)(3+\alpha\theta)}c^4xy$$

$$+\frac{4(k_{108}h_1h_2h_3h_4h_5h_6^2)}{3(1+\alpha\theta)(2+\alpha\gamma_{AB}\theta)(2+\alpha\theta)}c^3x(b^{\gamma_{AB}}-c^{\gamma_{AB}}y)$$

$$+\frac{k_{109}h_1h_2h_3h_4}{(1+\alpha\gamma_{CD}\theta)(1+\alpha\gamma_{AB}\theta)}(a^{2\gamma_{CD}}-c^{2\gamma_{CD}}x)(b^{2\gamma_{AB}}-c^{2\gamma_{AB}}y)$$

$$+\frac{k_{110}h_1h_2h_3h_4h_5^2}{(1+\alpha\gamma_{CD}\theta)(3+\alpha\theta)}c^2y(a^{2\gamma_{CD}}-c^{2\gamma_{CD}}x)+\frac{k_{111}h_1h_2h_3h_4h_6^2}{(1+\alpha\gamma_{AB}\theta)(3+\alpha\theta)}c^2x(b^{2\gamma_{AB}}-c^{2\gamma_{AB}}y)$$

$$+\frac{k_{112}h_1h_2h_3h_4h_5^2h_6^2+k_{113}h_1h_2h_3h_4h_5^2h_6^2}{2(3+\alpha\theta)^2}c^4xy+\frac{k_{114}h_1h_2h_3h_4h_5^2}{(1+\alpha\theta)(1+\alpha\gamma_{CD}\theta)(3+\alpha\theta)}c^2y(a^{2\gamma_{CD}}-c^{2\gamma_{CD}}x)$$

$$+\frac{k_{115}h_1h_2h_3h_4h_5^2h_6^2}{2(1+\alpha\theta)(3+\alpha\theta)^2}c^4xy$$

$$+\frac{k_{116}h_1h_2h_3h_4h_6^2}{(1+\alpha\theta)(1+\alpha\gamma_{AB}\theta)(3+\alpha\theta)}c^2x(b^{2\gamma_{AB}}-c^{2\gamma_{AB}}y)]$$

The constants $k_1 \ldots k_{116}$ differ by site pattern and are given by the following table:

| **Site pattern** | XXXX | XXXY | XXYX | XYXX | YXXX | XXYY | XYXY | XYYX | XXYZ | XYXZ | XYZX | YXXZ | YXZX | YZXX | XYWW |
|---|---|---|---|---|---|---|---|---|---|---|---|---|---|---|---|
| $k_1$ | 3 | -1 | -1 | 3 | 3 | 3 | -1 | -1 | -1 | -1 | -1 | -1 | -1 | 3 | -1 |
| $k_2$ | 9 | -3 | -3 | 9 | 9 | 9 | -3 | -3 | -3 | -3 | -3 | -3 | -3 | 9 | -3 |
| $k_3$ | 3 | -1 | -1 | 3 | 3 | 3 | -1 | -1 | -1 | -1 | -1 | -1 | -1 | 3 | -1 |
| $k_4$ | 9 | -3 | -3 | 9 | 9 | 9 | -3 | -3 | -3 | -3 | -3 | -3 | -3 | 9 | -3 |
| $k_5$ | 6 | 6 | -2 | 6 | -2 | -2 | 6 | -2 | -2 | 6 | -2 | -2 | -2 | -2 | -2 |
| $k_6$ | 9 | 9 | -3 | -3 | 9 | -3 | -3 | 9 | -3 | -3 | -3 | 9 | -3 | -3 | -3 |
| $k_7$ | 6 | -2 | 6 | 6 | -2 | -2 | -2 | 6 | -2 | -2 | 6 | -2 | -2 | -2 | -2 |
| $k_8$ | 9 | -3 | 9 | -3 | 9 | -3 | 9 | -3 | -3 | -3 | -3 | -3 | 9 | -3 | -3 |
| $k_9$ | 6 | 6 | 6 | -2 | -2 | 6 | -2 | -2 | 6 | -2 | -2 | -2 | -2 | -2 | -2 |
| $k_{10}$ | 3 | -1 | -1 | 3 | 3 | 3 | -1 | -1 | -1 | -1 | -1 | -1 | -1 | 3 | -1 |
| $k_{11}$ | 3 | 3 | -1 | -1 | 3 | -1 | -1 | 3 | -1 | -1 | -1 | 3 | -1 | -1 | -1 |
| $k_{12}$ | 3 | -1 | 3 | -1 | 3 | -1 | 3 | -1 | -1 | -1 | -1 | -1 | 3 | -1 | -1 |
| $k_{13}$ | 6 | 6 | 6 | -2 | -2 | 6 | -2 | -2 | 6 | -2 | -2 | -2 | -2 | -2 | -2 |
| $k_{14}$ | 3 | 3 | 3 | -1 | -1 | 3 | -1 | -1 | 3 | -1 | -1 | -1 | -1 | -1 | -1 |
| $k_{15}$ | 3 | 3 | 3 | -1 | -1 | 3 | -1 | -1 | 3 | -1 | -1 | -1 | -1 | -1 | -1 |
| $k_{16}$ | 3 | 3 | -1 | 3 | -1 | -1 | 3 | -1 | -1 | 3 | -1 | -1 | -1 | -1 | -1 |
| $k_{17}$ | 3 | 3 | -1 | -1 | 3 | -1 | -1 | 3 | -1 | -1 | -1 | 3 | -1 | -1 | -1 |
| $k_{18}$ | 3 | -1 | 3 | 3 | -1 | -1 | -1 | 3 | -1 | -1 | 3 | -1 | -1 | -1 | -1 |
| $k_{19}$ | 3 | -1 | 3 | -1 | 3 | -1 | 3 | -1 | -1 | -1 | -1 | -1 | 3 | -1 | -1 |
| $k_{20}$ | 3 | 3 | 3 | -1 | -1 | 3 | -1 | -1 | 3 | -1 | -1 | -1 | -1 | -1 | -1 |
| $k_{21}$ | 12 | -4 | -4 | 12 | 12 | 12 | -4 | -4 | -4 | -4 | -4 | -4 | -4 | 12 | -4 |
| $k_{22}$ | 15 | 15 | -5 | 15 | -5 | -5 | 15 | -5 | -5 | 15 | -5 | -5 | -5 | -5 | -5 |
| $k_{23}$ | 12 | 12 | -4 | -4 | 12 | -4 | -4 | 12 | -4 | -4 | -4 | 12 | -4 | -4 | -4 |
| $k_{24}$ | 15 | -5 | 15 | 15 | -5 | -5 | -5 | 15 | -5 | -5 | 15 | -5 | -5 | -5 | -5 |
| $k_{25}$ | 12 | -4 | 12 | -4 | 12 | -4 | 12 | -4 | -4 | -4 | -4 | -4 | 12 | -4 | -4 |
| $k_{26}$ | 15 | 15 | 15 | -5 | -5 | 15 | -5 | -5 | 15 | -5 | -5 | -5 | -5 | -5 | -5 |
| $k_{27}$ | 3 | 3 | 3 | -1 | -1 | 3 | -1 | -1 | 3 | -1 | -1 | -1 | -1 | -1 | -1 |
| $k_{28}$ | 3 | 3 | -1 | 3 | -1 | -1 | 3 | -1 | -1 | 3 | -1 | -1 | -1 | -1 | -1 |
| $k_{29}$ | 3 | -1 | 3 | 3 | -1 | -1 | -1 | 3 | -1 | -1 | 3 | -1 | -1 | -1 | -1 |
| $k_{30}$ | 3 | 3 | -1 | 3 | -1 | -1 | 3 | -1 | -1 | 3 | -1 | -1 | -1 | -1 | -1 |
| $k_{31}$ | 3 | 3 | -1 | -1 | 3 | -1 | -1 | 3 | -1 | -1 | -1 | 3 | -1 | -1 | -1 |
| $k_{32}$ | 3 | -1 | 3 | 3 | -1 | -1 | -1 | 3 | -1 | -1 | 3 | -1 | -1 | -1 | -1 |
| $k_{33}$ | 3 | -1 | 3 | -1 | 3 | -1 | 3 | -1 | -1 | -1 | -1 | -1 | 3 | -1 | -1 |
| $k_{34}$ | 3 | 3 | -1 | 3 | -1 | -1 | 3 | -1 | -1 | 3 | -1 | -1 | -1 | -1 | -1 |
| $k_{35}$ | 3 | 3 | -1 | -1 | 3 | -1 | -1 | 3 | -1 | -1 | -1 | 3 | -1 | -1 | -1 |
| $k_{36}$ | 3 | -1 | 3 | 3 | -1 | -1 | -1 | 3 | -1 | -1 | 3 | -1 | -1 | -1 | -1 |
| $k_{37}$ | 3 | -1 | 3 | -1 | 3 | -1 | 3 | -1 | -1 | -1 | -1 | -1 | 3 | -1 | -1 |
| $k_{38}$ | 6 | 6 | -2 | 6 | -2 | -2 | 6 | -2 | -2 | 6 | -2 | -2 | -2 | -2 | -2 |
| $k_{39}$ | 6 | 6 | -2 | -2 | 6 | -2 | -2 | 6 | -2 | -2 | -2 | 6 | -2 | -2 | -2 |
| $k_{40}$ | 6 | -2 | 6 | 6 | -2 | -2 | -2 | 6 | -2 | -2 | 6 | -2 | -2 | -2 | -2 |

| Site pattern | XXXX | XXXY | XXYX | XYXX | YXXX | XXYY | XYXY | XYYX | XXYZ | XYXZ | XYZX | YXXZ | YXZX | YZXX | XYWW |
|---|---|---|---|---|---|---|---|---|---|---|---|---|---|---|---|
| $k_{41}$ | 6 | -2 | 6 | -2 | 6 | -2 | 6 | -2 | -2 | -2 | -2 | -2 | 6 | -2 | -2 |
| $k_{42}$ | 6 | 6 | 6 | -2 | -2 | 6 | -2 | -2 | 6 | -2 | -2 | -2 | -2 | -2 | -2 |
| $k_{43}$ | 3 | 3 | -1 | 3 | -1 | -1 | 3 | -1 | -1 | 3 | -1 | -1 | -1 | -1 | -1 |
| $k_{44}$ | 3 | 3 | -1 | -1 | 3 | -1 | -1 | 3 | -1 | -1 | -1 | 3 | -1 | -1 | -1 |
| $k_{45}$ | 3 | -1 | 3 | 3 | -1 | -1 | -1 | 3 | -1 | -1 | 3 | -1 | -1 | -1 | -1 |
| $k_{46}$ | 3 | -1 | 3 | -1 | 3 | -1 | 3 | -1 | -1 | -1 | -1 | -1 | 3 | -1 | -1 |
| $k_{47}$ | 24 | 24 | -8 | 24 | -8 | -8 | 24 | -8 | -8 | 24 | -8 | -8 | -8 | -8 | -8 |
| $k_{48}$ | 24 | 24 | -8 | -8 | 24 | -8 | -8 | 24 | -8 | -8 | -8 | 24 | -8 | -8 | -8 |
| $k_{49}$ | 24 | -8 | 24 | 24 | -8 | -8 | -8 | 24 | -8 | -8 | 24 | -8 | -8 | -8 | -8 |
| $k_{50}$ | 24 | -8 | 24 | -8 | 24 | -8 | 24 | -8 | -8 | -8 | -8 | -8 | 24 | -8 | -8 |
| $k_{51}$ | 24 | -8 | -8 | 24 | 24 | 24 | -8 | -8 | -8 | -8 | -8 | -8 | -8 | 24 | -8 |
| $k_{52}$ | 24 | 24 | 24 | -8 | -8 | 24 | -8 | -8 | 24 | -8 | -8 | -8 | -8 | -8 | -8 |
| $k_{53}$ | 6 | 6 | -2 | 6 | -2 | -2 | 6 | -2 | -2 | 6 | -2 | -2 | -2 | -2 | -2 |
| $k_{54}$ | 6 | 6 | -2 | -2 | 6 | -2 | -2 | 6 | -2 | -2 | -2 | 6 | -2 | -2 | -2 |
| $k_{55}$ | 6 | -2 | 6 | 6 | -2 | -2 | -2 | 6 | -2 | -2 | 6 | -2 | -2 | -2 | -2 |
| $k_{56}$ | 6 | -2 | 6 | -2 | 6 | -2 | 6 | -2 | -2 | -2 | -2 | -2 | 6 | -2 | -2 |
| $k_{57}$ | 6 | -2 | -2 | 6 | 6 | 6 | -2 | -2 | -2 | -2 | -2 | -2 | -2 | 6 | -2 |
| $k_{58}$ | 6 | 6 | 6 | -2 | -2 | 6 | -2 | -2 | 6 | -2 | -2 | -2 | -2 | -2 | -2 |
| $k_{59}$ | 3 | 3 | -1 | 3 | -1 | -1 | 3 | -1 | -1 | 3 | -1 | -1 | -1 | -1 | -1 |
| $k_{60}$ | 3 | 3 | -1 | -1 | 3 | -1 | -1 | 3 | -1 | -1 | -1 | 3 | -1 | -1 | -1 |
| $k_{61}$ | 3 | -1 | 3 | 3 | -1 | -1 | -1 | 3 | -1 | -1 | 3 | -1 | -1 | -1 | -1 |
| $k_{62}$ | 3 | -1 | 3 | -1 | 3 | -1 | 3 | -1 | -1 | -1 | -1 | -1 | 3 | -1 | -1 |
| $k_{63}$ | 6 | -2 | -2 | 6 | 6 | 6 | -2 | -2 | -2 | -2 | -2 | -2 | -2 | 6 | -2 |
| $k_{64}$ | 6 | -2 | -2 | 6 | -2 | -2 | -2 | -2 | 2 | -2 | -2 | 2 | 2 | -2 | 2 |
| $k_{65}$ | 6 | -2 | -2 | -2 | 6 | -2 | -2 | -2 | 2 | 2 | 2 | -2 | -2 | -2 | 2 |
| $k_{66}$ | 18 | -6 | -6 | 18 | -6 | -6 | -6 | -6 | 6 | -6 | -6 | 6 | 6 | -6 | 6 |
| $k_{67}$ | 18 | -6 | -6 | -6 | 18 | -6 | -6 | -6 | 6 | 6 | 6 | -6 | -6 | -6 | 6 |
| $k_{68}$ | 18 | 18 | -6 | -6 | -6 | -6 | -6 | -6 | -6 | -6 | 6 | -6 | 6 | 6 | 6 |
| $k_{69}$ | 18 | -6 | 18 | -6 | -6 | -6 | -6 | -6 | -6 | 6 | -6 | 6 | -6 | 6 | 6 |
| $k_{70}$ | 6 | 6 | -2 | -2 | -2 | -2 | -2 | -2 | -2 | -2 | 2 | -2 | 2 | 2 | 2 |
| $k_{71}$ | 6 | -2 | 6 | -2 | -2 | -2 | -2 | -2 | -2 | 2 | -2 | 2 | -2 | 2 | 2 |
| $k_{72}$ | 6 | -2 | -2 | 6 | -2 | -2 | -2 | -2 | 2 | -2 | -2 | 2 | 2 | -2 | 2 |
| $k_{73}$ | 6 | -2 | -2 | -2 | 6 | -2 | -2 | -2 | 2 | 2 | 2 | -2 | -2 | -2 | 2 |
| $k_{74}$ | 12 | -4 | -4 | 12 | -4 | -4 | -4 | -4 | 4 | -4 | -4 | 4 | 4 | -4 | 4 |
| $k_{75}$ | 12 | -4 | -4 | -4 | 12 | -4 | -4 | -4 | 4 | 4 | 4 | -4 | -4 | -4 | 4 |
| $k_{76}$ | 6 | -2 | -2 | 6 | -2 | -2 | -2 | -2 | 2 | -2 | -2 | 2 | 2 | -2 | 2 |
| $k_{77}$ | 6 | -2 | -2 | -2 | 6 | -2 | -2 | -2 | 2 | 2 | 2 | -2 | -2 | -2 | 2 |
| $k_{78}$ | 24 | -8 | -8 | 24 | -8 | -8 | -8 | -8 | 8 | -8 | -8 | 8 | 8 | -8 | 8 |
| $k_{79}$ | 36 | -12 | -12 | -12 | 36 | -12 | -12 | -12 | 12 | 12 | 12 | -12 | -12 | -12 | 12 |
| $k_{80}$ | 24 | 24 | -8 | -8 | -8 | -8 | -8 | -8 | -8 | -8 | 8 | -8 | 8 | 8 | 8 |

| **Site pattern** | XXXX | XXXY | XXYX | XYXX | YXXX | XXYY | XYXY | XYYX | XXYZ | XYXZ | XYZX | YXXZ | YXZX | YZXX | XYWW |
|---|---|---|---|---|---|---|---|---|---|---|---|---|---|---|---|
| $k_{81}$ | 24 | -8 | 24 | -8 | -8 | -8 | -8 | -8 | -8 | 8 | -8 | 8 | -8 | 8 | 8 |
| $k_{82}$ | 6 | -2 | -2 | 6 | -2 | -2 | -2 | -2 | 2 | -2 | -2 | 2 | 2 | -2 | 2 |
| $k_{83}$ | 18 | -6 | -6 | -6 | 18 | -6 | -6 | -6 | 6 | 6 | 6 | -6 | -6 | -6 | 6 |
| $k_{84}$ | 6 | 6 | -2 | -2 | -2 | -2 | -2 | -2 | -2 | -2 | 2 | -2 | 2 | 2 | 2 |
| $k_{85}$ | 6 | -2 | 6 | -2 | -2 | -2 | -2 | -2 | -2 | 2 | -2 | 2 | -2 | 2 | 2 |
| $k_{86}$ | 6 | 6 | -2 | -2 | -2 | -2 | -2 | -2 | -2 | -2 | 2 | -2 | 2 | 2 | 2 |
| $k_{87}$ | 6 | -2 | 6 | -2 | -2 | -2 | -2 | -2 | -2 | 2 | -2 | 2 | -2 | 2 | 2 |
| $k_{88}$ | 6 | 6 | -2 | -2 | -2 | -2 | -2 | -2 | -2 | -2 | 2 | -2 | 2 | 2 | 2 |
| $k_{89}$ | 6 | -2 | 6 | -2 | -2 | -2 | -2 | -2 | -2 | 2 | -2 | 2 | -2 | 2 | 2 |
| $k_{90}$ | 18 | 18 | -6 | -6 | -6 | -6 | -6 | -6 | -6 | -6 | 6 | -6 | 6 | 6 | 6 |
| $k_{91}$ | 18 | -6 | 18 | -6 | -6 | -6 | -6 | -6 | -6 | 6 | -6 | 6 | -6 | 6 | 6 |
| $k_{92}$ | 6 | 6 | -2 | -2 | -2 | -2 | -2 | -2 | -2 | -2 | 2 | -2 | 2 | 2 | 2 |
| $k_{93}$ | 6 | -2 | 6 | -2 | -2 | -2 | -2 | -2 | -2 | 2 | -2 | 2 | -2 | 2 | 2 |
| $k_{94}$ | 48 | -16 | -16 | 48 | -16 | -16 | -16 | -16 | 16 | -16 | -16 | 16 | 16 | -16 | 16 |
| $k_{95}$ | 36 | -12 | -12 | -12 | 36 | -12 | -12 | -12 | 12 | 12 | 12 | -12 | -12 | -12 | 12 |
| $k_{96}$ | 48 | 48 | -16 | -16 | -16 | -16 | -16 | -16 | -16 | -16 | 16 | -16 | 16 | 16 | 16 |
| $k_{97}$ | 48 | -16 | 48 | -16 | -16 | -16 | -16 | -16 | -16 | 16 | -16 | 16 | -16 | 16 | 16 |
| $k_{98}$ | 12 | -4 | -4 | 12 | -4 | -4 | -4 | -4 | 4 | -4 | -4 | 4 | 4 | -4 | 4 |
| $k_{99}$ | 12 | 12 | -4 | -4 | -4 | -4 | -4 | -4 | -4 | -4 | 4 | -4 | 4 | 4 | 4 |
| $k_{100}$ | 12 | -4 | 12 | -4 | -4 | -4 | -4 | -4 | -4 | 4 | -4 | 4 | -4 | 4 | 4 |
| $k_{101}$ | 12 | -4 | -4 | 12 | -4 | -4 | -4 | -4 | 4 | -4 | -4 | 4 | 4 | -4 | 4 |
| $k_{102}$ | 12 | -4 | -4 | -4 | 12 | -4 | -4 | -4 | 4 | 4 | 4 | -4 | -4 | -4 | 4 |
| $k_{103}$ | 12 | -4 | -4 | -4 | -4 | -4 | 4 | 4 | 4 | 0 | 0 | 0 | 0 | 4 | -4 |
| $k_{104}$ | 36 | -12 | -12 | -12 | -12 | -12 | 12 | 12 | 12 | 0 | 0 | 0 | 0 | 12 | -12 |
| $k_{105}$ | 12 | -4 | -4 | -4 | -4 | -4 | 4 | 4 | 4 | 0 | 0 | 0 | 0 | 4 | -4 |
| $k_{106}$ | 180 | -60 | -60 | -60 | -60 | 20 | 20 | 20 | 20 | 20 | 20 | 20 | 20 | 20 | -60 |
| $k_{107}$ | 36 | -12 | -12 | -12 | -12 | 4 | 4 | 4 | 4 | 4 | 4 | 4 | 4 | 4 | -12 |
| $k_{108}$ | 24 | -8 | -8 | -8 | -8 | -8 | 8 | 8 | 8 | 0 | 0 | 0 | 0 | 8 | -8 |
| $k_{109}$ | 9 | -3 | -3 | -3 | -3 | 9 | 1 | 1 | -3 | 1 | 1 | 1 | 1 | -3 | 1 |
| $k_{110}$ | 9 | -3 | -3 | -3 | -3 | 9 | 1 | 1 | -3 | 1 | 1 | 1 | 1 | -3 | 1 |
| $k_{111}$ | 9 | -3 | -3 | -3 | -3 | 9 | 1 | 1 | -3 | 1 | 1 | 1 | 1 | -3 | 1 |
| $k_{112}$ | 27 | -9 | -9 | -9 | -9 | 11 | 11 | 11 | -1 | -1 | -1 | -1 | -1 | -1 | 3 |
| $k_{113}$ | 27 | -9 | -9 | -9 | -9 | 11 | 11 | 11 | -1 | -1 | -1 | -1 | -1 | -1 | 3 |
| $k_{114}$ | 18 | -6 | -6 | -6 | -6 | 18 | 2 | 2 | -6 | 2 | 2 | 2 | 2 | -6 | 2 |
| $k_{115}$ | 108 | -36 | -36 | -36 | -36 | 44 | 44 | 44 | -4 | -4 | -4 | -4 | -4 | -4 | 12 |
| $k_{116}$ | 18 | -6 | -6 | -6 | -6 | 18 | 2 | 2 | -6 | 2 | 2 | 2 | 2 | -6 | 2 |

#### 1.2 Asymmetric case

Let

$$x = exp[-\frac{2(\tau_2 - \tau_1)}{\theta}], y = exp[-\frac{2(\tau_3 - \tau_2)}{\theta}], a = e^{-\alpha\tau_1}, b = e^{-\alpha\tau_2}, c = e^{-\alpha\tau_3}$$

$$h_1 = e^{-\alpha\tau_1(\gamma_C - \gamma_{CD})}, h_2 = e^{-\alpha\tau_1(\gamma_D - \gamma_{CD})}, h_3 = e^{-\alpha\tau_2(\gamma_B - \gamma_{BCD})}$$

$$h_4 = e^{-\alpha\tau_2(\gamma_{CD} - \gamma_{BCD})}, h_5 = e^{-\alpha\tau_3(\gamma_{AB} - 1)}, h_6 = e^{-\alpha\tau_3(\gamma_{BCD} - 1)}$$

Then each site pattern probability is of the form:

$$\frac{1}{256}[1 + \frac{k_1 h_1 h_2}{3(1+\alpha\gamma_{CD}\theta)} y(a^{2\gamma_{CD}} - b^{2\gamma_{CD}}x)$$

$$+\frac{k_2 h_1 h_2 h_4^2 + k_3 h_1 h_3 h_4 + k_4 h_2 h_3 h_4}{3(2+\alpha\gamma_{BCD}\theta)} x(b^{2\gamma_{BCD}}y - c^{2\gamma_{BCD}}y^3)$$

$$+\frac{k_5 h_1 h_2 h_4^2 h_6^2 + k_6 h_1 h_3 h_4 h_6^2 + k_7 h_1 h_4 h_5 h_6 + k_8 h_2 h_3 h_4 h_6^2 + k_9 h_2 h_4 h_5 h_6 + k_{10} h_3 h_5 h_6}{3(6+\alpha\theta)} c^2 x y^3$$

$$+\frac{k_{11} h_1 h_2 h_4^2 h_6^2 + k_{12} h_1 h_3 h_4 h_6^2 + k_{13} h_2 h_3 h_4 h_6^2}{(3+\alpha\theta)(6+\alpha\theta)} c^2 x y^3 + \frac{k_{14} h_1 h_2}{(1+\alpha\gamma_{CD}\theta)}(a^{2\gamma_{CD}} - b^{2\gamma_{CD}}x)(1-y)$$

$$+\frac{2(k_{15} h_1 h_2 h_4^2 + k_{16} h_1 h_3 h_4 + k_{17} h_2 h_3 h_4)}{2+\alpha\gamma_{BCD}\theta} x(\frac{b^{2\gamma_{BCD}}(1-y)}{2} - \frac{b^{2\gamma_{BCD}} - c^{2\gamma_{BCD}}y^3}{6+2\alpha\gamma_{BCD}\theta})$$

$$+\frac{k_{18} h_1 h_3 h_4 h_6^2 + k_{19} h_1 h_4 h_5 h_6 + k_{20} h_2 h_3 h_4 h_6^2 + k_{21} h_2 h_4 h_5 h_6 + k_{22} h_3 h_5 h_6}{3+\alpha\theta} c^2 y(1-x)$$

$$+\frac{k_{23} h_1 h_2 h_4^2 h_6^2 + k_{24} h_1 h_3 h_4 h_6^2 + k_{25} h_1 h_4 h_5 h_6 + k_{26} h_2 h_3 h_4 h_6^2 + k_{27} h_2 h_4 h_5 h_6 + k_{28} h_3 h_5 h_6}{2(3+\alpha\theta)} x(c^2 y - c^2 y^3)$$

$$+\frac{k_{29} h_1 h_2 h_4^2 h_6^2 + k_{30} h_1 h_3 h_4 h_6^2 + k_{31} h_1 h_4 h_5 h_6 + k_{32} h_2 h_3 h_4 h_6^2 + k_{33} h_2 h_4 h_5 h_6 + k_{34} h_3 h_5 h_6}{(3+\alpha\theta)(6+\alpha\theta)} c^2 x y^3$$

$$+\frac{k_{35} h_1 h_4 h_5 h_6 + k_{36} h_2 h_4 h_5 h_6 + k_{37} h_3 h_5 h_6}{3(6+\alpha\theta)} c^2 x y^3 + \frac{k_{38} h_1 h_3 h_4 + k_{39} h_2 h_3 h_4}{(1+\alpha\gamma_{BCD}\theta)}(1-x)(b^{2\gamma_{BCD}} - c^{2\gamma_{BCD}}y)$$

$$+(k_{40} h_1 h_2 h_4^2 + k_{41} h_1 h_3 h_4 + k_{42} h_2 h_3 h_4) x(\frac{b^{2\gamma_{BCD}} - c^{2\gamma_{BCD}}y}{2+2\alpha\gamma_{BCD}\theta} - \frac{b^{2\gamma_{BCD}} - c^{2\gamma_{BCD}}y^3}{6+2\alpha\gamma_{BCD}\theta})$$

$$+\frac{k_{43} h_1 h_3 h_4 h_6^2 + k_{44} h_1 h_4 h_5 h_6 + k_{45} h_2 h_3 h_4 h_6^2 + k_{46} h_2 h_4 h_5 h_6 + k_{47} h_3 h_5 h_6}{(1+\alpha\theta)(3+\alpha\theta)} c^2 y(1-x)$$

$$+\frac{k_{48} h_1 h_2 h_4^2 h_6^2 + k_{49} h_1 h_3 h_4 h_6^2 + k_{50} h_1 h_4 h_5 h_6 + k_{51} h_2 h_3 h_4 h_6^2 + k_{52} h_2 h_4 h_5 h_6 + k_{53} h_3 h_5 h_6}{2(1+\alpha\theta)(3+\alpha\theta)} x(c^2 y - c^2 y^3)$$

$$+\frac{k_{54}h_1h_2h_4^2h_6^2+k_{55}h_1h_3h_4h_6^2+k_{56}h_1h_4h_5h_6+k_{57}h_2h_3h_4h_6^2+k_{58}h_2h_4h_5h_6+k_{59}h_3h_5h_6}{(1+\alpha\theta)(3+\alpha\theta)(6+\alpha\theta)}c^2xy^3$$

$$+\frac{k_{60}h_1h_2h_4^2h_6^2+k_{61}h_1h_3h_4h_6^2+k_{62}h_1h_4h_5h_6+k_{63}h_2h_3h_4h_6^2+k_{64}h_2h_4h_5h_6+k_{65}h_3h_5h_6}{(1+\alpha\theta)(3+\alpha\theta)(6+\alpha\theta)}c^2xy^3$$

$$+\frac{k_{66}h_1h_4h_5h_6+k_{67}h_2h_4h_5h_6+k_{68}h_3h_5h_6}{(1+\alpha\theta)}c^2(1-x)(1-y)$$

$$+\frac{k_{69}h_1h_4h_5h_6+k_{70}h_2h_4h_5h_6+k_{71}h_3h_5h_6}{(1+\alpha\theta)}c^2x(\frac{(1-y)}{2}-\frac{1-y^3}{6})$$

$$+\frac{2(k_{72}h_1h_2h_3h_4h_6^2+k_{73}h_1h_2h_4h_5h_6)}{(2+\alpha\gamma_{CD}\theta)(3+\alpha\theta)}c^2y(a^{\gamma_{CD}}-b^{\gamma_{CD}}x)$$

$$+\frac{2(k_{74}h_1h_2h_3h_4^2h_6^2+k_{75}h_1h_2h_4^2h_5h_6+k_{76}h_1h_3h_4h_5h_6+k_{77}h_2h_3h_4h_5h_6)}{(3+\alpha\theta)(4+\alpha\gamma_{BCD}\theta)}x(b^{\gamma_{BCD}}c^2y-c^{2+\gamma_{BCD}}y^3)$$

$$+\frac{2(k_{78}h_1h_2h_3h_4^2h_6^3+k_{79}h_1h_2h_4^2h_5h_6^2+k_{80}h_1h_3h_4h_5h_6^2+k_{81}h_2h_3h_4h_5h_6^2)}{3(3+\alpha\theta)(4+\alpha\theta)}c^3xy^3$$

$$+\frac{2(k_{82}h_1h_2h_3h_4)}{(1+\alpha\gamma_{BCD}\theta)(2+\alpha\gamma_{CD}\theta)}(a^{\gamma_{CD}}-b^{\gamma_{CD}}x)(b^{2\gamma_{BCD}}-c^{2\gamma_{BCD}}y)$$

$$+\frac{4(k_{83}h_1h_2h_3h_4^2)}{4+\alpha\gamma_{BCD}\theta}x(\frac{b^{3\gamma_{BCD}}-b^{\gamma_{BCD}}c^{2\gamma_{BCD}}y}{2+2\alpha\gamma_{BCD}\theta}-\frac{b^{3\gamma_{BCD}}-c^{3\gamma_{BCD}}y^3}{6+3\alpha\gamma_{BCD}\theta})$$

$$+\frac{2(k_{84}h_1h_2h_3h_4h_6^2+k_{85}h_1h_2h_4h_5h_6)}{(1+\alpha\theta)(2+\alpha\gamma_{CD}\theta)(3+\alpha\theta)}c^2y(a^{\gamma_{CD}}-b^{\gamma_{CD}}x)$$

$$+\frac{2(k_{86}h_1h_2h_3h_4^2h_6^2+k_{87}h_1h_2h_4^2h_5h_6+k_{88}h_1h_3h_4h_5h_6+k_{89}h_2h_3h_4h_5h_6)}{(1+\alpha\theta)(3+\alpha\theta)(4+\alpha\gamma_{BCD}\theta)}x(b^{\gamma_{BCD}}c^2y-c^{2+\gamma_{BCD}}y^3)$$

$$+\frac{2(k_{90}h_1h_2h_3h_4^2h_6^3+k_{91}h_1h_2h_4^2h_5h_6^2+k_{92}h_1h_3h_4h_5h_6^2+k_{93}h_2h_3h_4h_5h_6^2)}{3(1+\alpha\theta)(3+\alpha\theta)(4+\alpha\theta)}c^3xy^3$$

$$+\frac{4(k_{94}h_1h_2h_3h_4^2h_6^3+k_{95}h_1h_2h_4^2h_5h_6^2+k_{96}h_1h_3h_4h_5h_6^2+k_{97}h_2h_3h_4h_5h_6^2)}{9(1+\alpha\theta)(2+\alpha\theta)(4+\alpha\theta)}c^3xy^3$$

$$+\frac{2(k_{98}h_1h_2h_4h_5h_6)}{(1+\alpha\theta)(2+\alpha\gamma_{CD}\theta)}c^2(a^{\gamma_{CD}}-b^{\gamma_{CD}}x)(1-y)$$

$$+\frac{4(k_{99}h_1h_2h_4^2h_5h_6+k_{100}h_1h_3h_4h_5h_6+k_{101}h_2h_3h_4h_5h_6)}{(1+\alpha\theta)(4+\alpha\gamma_{BCD}\theta)}c^2x(\frac{b^{\gamma_{BCD}}-b^{\gamma_{BCD}}y}{2}-\frac{b^{\gamma_{BCD}}-c^{\gamma_{BCD}}y^3}{6+\alpha\gamma_{BCD}\theta})$$

$$+\frac{2(k_{102}h_1h_3h_4h_5h_6^2+k_{103}h_2h_3h_4h_5h_6^2)}{3(1+\alpha\theta)(2+\alpha\theta)}c^3y(1-x)$$

$$+\frac{k_{104}h_1h_2h_4^2h_5h_6^2+k_{105}h_1h_3h_4h_5h_6^2+k_{106}h_2h_3h_4h_5h_6^2}{3(1+\alpha\theta)(2+\alpha\theta)}xc^3(y-y^3)$$

$$+\frac{4(k_{107}h_1h_2h_3h_4^2h_6^3+k_{108}h_1h_2h_4^2h_5h_6^2+k_{109}h_1h_3h_4h_5h_6^2+k_{110}h_2h_3h_4h_5h_6^2)}{9(1+\alpha\theta)(2+\alpha\theta)(4+\alpha\theta)}c^3xy^3$$

$$+\frac{2(k_{111}h_1h_2h_4^2h_5h_6^2+k_{112}h_1h_3h_4h_5h_6^2+k_{113}h_2h_3h_4h_5h_6^2)}{3(1+\alpha\theta)(3+\alpha\theta)(4+\alpha\theta)}c^3xy^3$$

$$+\frac{2(k_{114}h_1h_3h_4h_5h_6+k_{115}h_2h_3h_4h_5h_6)}{(1+\alpha\theta)(2+\alpha\gamma_{BCD}\theta)}c^2(1-x)(b^{\gamma_{BCD}}-c^{\gamma_{BCD}}y)$$

$$+\frac{k_{116}h_1h_2h_4^2h_5h_6+k_{117}h_1h_3h_4h_5h_6+k_{118}h_2h_3h_4h_5h_6}{(1+\alpha\theta)}c^2x(\frac{b^{\gamma_{BCD}}-c^{\gamma_{BCD}}y}{2+\alpha\gamma_{BCD}\theta}-\frac{b^{\gamma_{BCD}}-c^{\gamma_{BCD}}y^3}{6+\alpha\gamma_{BCD}\theta})$$

$$+\frac{4k_{119}h_1h_2h_3h_4h_5h_6^2}{3(1+\alpha\theta)(2+\alpha\gamma_{CD}\theta)(2+\alpha\theta)}c^3y(a^{\gamma_{CD}}-b^{\gamma_{CD}}x)$$

$$+\frac{4k_{120}h_1h_2h_3h_4^2h_5h_6^2}{3(1+\alpha\theta)(2+\alpha\theta)(4+\alpha\gamma_{BCD}\theta)}x(b^{\gamma_{BCD}}c^2y-c^{2+\gamma_{BCD}}y^3)$$

$$+\frac{k_{121}h_1h_2h_3h_4^2h_5h_6^3+k_{122}h_1h_2h_3h_4^2h_5h_6^3}{3(1+\alpha\theta)(2+\alpha\theta)(3+\alpha\theta)}c^4xy^3$$

$$+\frac{4(k_{123}h_1h_2h_3h_4h_5h_6)}{(1+\alpha\theta)(2+\alpha\gamma_{CD}\theta)(2+\alpha\gamma_{BCD}\theta)}c^2(a^{\gamma_{CD}}-b^{\gamma_{CD}}x)(b^{\gamma_{BCD}}-c^{\gamma_{BCD}}y)$$

$$+\frac{4(k_{124}h_1h_2h_3h_4^2h_5h_6)}{(1+\alpha\theta)(4+\alpha\gamma_{BCD}\theta)}c^2x(\frac{b^{2\gamma_{BCD}}-b^{\gamma_{BCD}}c^{\gamma_{BCD}}y}{2+\alpha\gamma_{BCD}\theta}-\frac{b^{2\gamma_{BCD}}-c^{2\gamma_{BCD}}y^3}{6+2\alpha\gamma_{BCD}\theta})$$

$$+\frac{k_{125}h_1h_2h_3h_5h_6}{(1+\alpha\gamma_{CD}\theta)(3+\alpha\theta)}c^2y(a^{2\gamma_{CD}}-b^{2\gamma_{CD}}x)$$

$$+\frac{k_{126}h_1h_2h_3h_4^2h_5h_6}{(2+\alpha\gamma_{BCD}\theta)(3+\alpha\theta)}x(b^{2\gamma_{BCD}}c^2y-c^{2+2\gamma_{BCD}}y^3)+\frac{k_{127}h_1h_2h_3h_4^2h_5h_6^3+k_{128}h_1h_2h_3h_4^2h_5h_6^3}{2(3+\alpha\theta)^2}c^4xy^3$$

$$+\frac{k_{129}h_1h_2h_3h_5h_6}{(1+\alpha\theta)(1+\alpha\gamma_{CD}\theta)(3+\alpha\theta)}c^2y(a^{2\gamma_{CD}}-b^{2\gamma_{CD}}x)$$

$$+\frac{k_{130}h_1h_2h_3h_4^2h_5h_6}{(1+\alpha\theta)(2+\alpha\gamma_{BCD}\theta)(3+\alpha\theta)}x(b^{2\gamma_{BCD}}c^2y-c^{2+2\gamma_{BCD}}y^3)$$

$$+\frac{k_{131}h_1h_2h_3h_4^2h_5h_6^3}{2(1+\alpha\theta)(3+\alpha\theta)^2}c^4xy^3+\frac{k_{132}h_1h_2h_3h_5h_6}{(1+\alpha\theta)(1+\alpha\gamma_{CD}\theta)}c^2(a^{2\gamma_{CD}}-b^{2\gamma_{CD}}x)(1-y)$$

$$+\frac{2(k_{133}h_1h_2h_3h_4^2h_5h_6)}{(1+\alpha\theta)(2+\alpha\gamma_{BCD}\theta)}c^2x(\frac{b^{2\gamma_{BCD}}-b^{2\gamma_{BCD}}y}{2}-\frac{b^{2\gamma_{BCD}}-c^{2\gamma_{BCD}}y^3}{6+2\alpha\gamma_{BCD}\theta})]$$

| **Site pattern** | XXXX | XXXY | XXYX | XYXX | YXXX | XXYY | XYXY | XYYX | XXYZ | XYXZ | XYZX | YXXZ | YXZX | YZXX | XYWW |
|---|---|---|---|---|---|---|---|---|---|---|---|---|---|---|---|
| $k_1$ | 9 | -3 | -3 | 9 | 9 | 9 | -3 | -3 | -3 | -3 | -3 | -3 | -3 | 9 | -3 |
| $k_2$ | 9 | -3 | -3 | 9 | 9 | 9 | -3 | -3 | -3 | -3 | -3 | -3 | -3 | 9 | -3 |
| $k_3$ | 9 | 9 | -3 | -3 | 9 | -3 | -3 | 9 | -3 | -3 | -3 | 9 | -3 | -3 | -3 |
| $k_4$ | 9 | -3 | 9 | -3 | 9 | -3 | 9 | -3 | -3 | -3 | -3 | -3 | 9 | -3 | -3 |
| $k_5$ | 9 | -3 | -3 | 9 | 9 | 9 | -3 | -3 | -3 | -3 | -3 | -3 | -3 | 9 | -3 |
| $k_6$ | 9 | 9 | -3 | -3 | 9 | -3 | -3 | 9 | -3 | -3 | -3 | 9 | -3 | -3 | -3 |
| $k_7$ | 6 | 6 | -2 | 6 | -2 | -2 | 6 | -2 | -2 | 6 | -2 | -2 | -2 | -2 | -2 |
| $k_8$ | 9 | -3 | 9 | -3 | 9 | -3 | 9 | -3 | -3 | -3 | -3 | -3 | 9 | -3 | -3 |
| $k_9$ | 6 | -2 | 6 | 6 | -2 | -2 | -2 | 6 | -2 | -2 | 6 | -2 | -2 | -2 | -2 |
| $k_{10}$ | 6 | 6 | 6 | -2 | -2 | 6 | -2 | -2 | 6 | -2 | -2 | -2 | -2 | -2 | -2 |
| $k_{11}$ | 3 | -1 | -1 | 3 | 3 | 3 | -1 | -1 | -1 | -1 | -1 | -1 | -1 | 3 | -1 |
| $k_{12}$ | 3 | 3 | -1 | -1 | 3 | -1 | -1 | 3 | -1 | -1 | -1 | 3 | -1 | -1 | -1 |
| $k_{13}$ | 3 | -1 | 3 | -1 | 3 | -1 | 3 | -1 | -1 | -1 | -1 | -1 | 3 | -1 | -1 |
| $k_{14}$ | 3 | -1 | -1 | 3 | 3 | 3 | -1 | -1 | -1 | -1 | -1 | -1 | -1 | 3 | -1 |
| $k_{15}$ | 3 | -1 | -1 | 3 | 3 | 3 | -1 | -1 | -1 | -1 | -1 | -1 | -1 | 3 | -1 |
| $k_{16}$ | 3 | 3 | -1 | -1 | 3 | -1 | -1 | 3 | -1 | -1 | -1 | 3 | -1 | -1 | -1 |
| $k_{17}$ | 3 | -1 | 3 | -1 | 3 | -1 | 3 | -1 | -1 | -1 | -1 | -1 | 3 | -1 | -1 |
| $k_{18}$ | 3 | 3 | -1 | -1 | 3 | -1 | -1 | 3 | -1 | -1 | -1 | 3 | -1 | -1 | -1 |
| $k_{19}$ | 3 | 3 | -1 | 3 | -1 | -1 | 3 | -1 | -1 | 3 | -1 | -1 | -1 | -1 | -1 |
| $k_{20}$ | 3 | -1 | 3 | -1 | 3 | -1 | 3 | -1 | -1 | -1 | -1 | -1 | 3 | -1 | -1 |
| $k_{21}$ | 3 | -1 | 3 | 3 | -1 | -1 | -1 | 3 | -1 | -1 | 3 | -1 | -1 | -1 | -1 |
| $k_{22}$ | 3 | 3 | 3 | -1 | -1 | 3 | -1 | -1 | 3 | -1 | -1 | -1 | -1 | -1 | -1 |
| $k_{23}$ | 6 | -2 | -2 | 6 | 6 | 6 | -2 | -2 | -2 | -2 | -2 | -2 | -2 | 6 | -2 |
| $k_{24}$ | 6 | 6 | -2 | -2 | 6 | -2 | -2 | 6 | -2 | -2 | -2 | 6 | -2 | -2 | -2 |
| $k_{25}$ | 9 | 9 | -3 | 9 | -3 | -3 | 9 | -3 | -3 | 9 | -3 | -3 | -3 | -3 | -3 |
| $k_{26}$ | 6 | -2 | 6 | -2 | 6 | -2 | 6 | -2 | -2 | -2 | -2 | -2 | 6 | -2 | -2 |
| $k_{27}$ | 9 | -3 | 9 | 9 | -3 | -3 | -3 | 9 | -3 | -3 | 9 | -3 | -3 | -3 | -3 |
| $k_{28}$ | 9 | 9 | 9 | -3 | -3 | 9 | -3 | -3 | 9 | -3 | -3 | -3 | -3 | -3 | -3 |
| $k_{29}$ | 12 | -4 | -4 | 12 | 12 | 12 | -4 | -4 | -4 | -4 | -4 | -4 | -4 | 12 | -4 |
| $k_{30}$ | 12 | 12 | -4 | -4 | 12 | -4 | -4 | 12 | -4 | -4 | -4 | 12 | -4 | -4 | -4 |
| $k_{31}$ | 15 | 15 | -5 | 15 | -5 | -5 | 15 | -5 | -5 | 15 | -5 | -5 | -5 | -5 | -5 |
| $k_{32}$ | 12 | -4 | 12 | -4 | 12 | -4 | 12 | -4 | -4 | -4 | -4 | -4 | 12 | -4 | -4 |
| $k_{33}$ | 15 | -5 | 15 | 15 | -5 | -5 | -5 | 15 | -5 | -5 | 15 | -5 | -5 | -5 | -5 |
| $k_{34}$ | 15 | 15 | 15 | -5 | -5 | 15 | -5 | -5 | 15 | -5 | -5 | -5 | -5 | -5 | -5 |
| $k_{35}$ | 3 | 3 | -1 | 3 | -1 | -1 | 3 | -1 | -1 | 3 | -1 | -1 | -1 | -1 | -1 |
| $k_{36}$ | 3 | -1 | 3 | 3 | -1 | -1 | -1 | 3 | -1 | -1 | 3 | -1 | -1 | -1 | -1 |
| $k_{37}$ | 3 | 3 | 3 | -1 | -1 | 3 | -1 | -1 | 3 | -1 | -1 | -1 | -1 | -1 | -1 |
| $k_{38}$ | 3 | 3 | -1 | -1 | 3 | -1 | -1 | 3 | -1 | -1 | -1 | 3 | -1 | -1 | -1 |
| $k_{39}$ | 3 | -1 | 3 | -1 | 3 | -1 | 3 | -1 | -1 | -1 | -1 | -1 | 3 | -1 | -1 |
| $k_{40}$ | 6 | -2 | -2 | 6 | 6 | 6 | -2 | -2 | -2 | -2 | -2 | -2 | -2 | 6 | -2 |
| $k_{41}$ | 6 | 6 | -2 | -2 | 6 | -2 | -2 | 6 | -2 | -2 | -2 | 6 | -2 | -2 | -2 |
| $k_{42}$ | 6 | -2 | 6 | -2 | 6 | -2 | 6 | -2 | -2 | -2 | -2 | -2 | 6 | -2 | -2 |
| $k_{43}$ | 6 | 6 | -2 | -2 | 6 | -2 | -2 | 6 | -2 | -2 | -2 | 6 | -2 | -2 | -2 |
| $k_{44}$ | 6 | 6 | -2 | 6 | -2 | -2 | 6 | -2 | -2 | 6 | -2 | -2 | -2 | -2 | -2 |

| **Site pattern** | XXXX | XXXY | XXYX | XYXX | YXXX | XXYY | XYXY | XYYX | XXYZ | XYXZ | XYZX | YXXZ | YXZX | YZXX | XYWW |
|---|---|---|---|---|---|---|---|---|---|---|---|---|---|---|---|
| $k_{45}$ | 6 | -2 | 6 | -2 | 6 | -2 | 6 | -2 | -2 | -2 | -2 | -2 | 6 | -2 | -2 |
| $k_{46}$ | 6 | -2 | 6 | 6 | -2 | -2 | -2 | 6 | -2 | -2 | 6 | -2 | -2 | -2 | -2 |
| $k_{47}$ | 6 | 6 | 6 | -2 | -2 | 6 | -2 | -2 | 6 | -2 | -2 | -2 | -2 | -2 | -2 |
| $k_{48}$ | 12 | -4 | -4 | 12 | 12 | 12 | -4 | -4 | -4 | -4 | -4 | -4 | -4 | 12 | -4 |
| $k_{49}$ | 12 | 12 | -4 | -4 | 12 | -4 | -4 | 12 | -4 | -4 | -4 | 12 | -4 | -4 | -4 |
| $k_{50}$ | 18 | 18 | -6 | 18 | -6 | -6 | 18 | -6 | -6 | 18 | -6 | -6 | -6 | -6 | -6 |
| $k_{51}$ | 12 | -4 | 12 | -4 | 12 | -4 | 12 | -4 | -4 | -4 | -4 | -4 | 12 | -4 | -4 |
| $k_{52}$ | 18 | -6 | 18 | 18 | -6 | -6 | -6 | 18 | -6 | -6 | 18 | -6 | -6 | -6 | -6 |
| $k_{53}$ | 18 | 18 | 18 | -6 | -6 | 18 | -6 | -6 | 18 | -6 | -6 | -6 | -6 | -6 | -6 |
| $k_{54}$ | 24 | -8 | -8 | 24 | 24 | 24 | -8 | -8 | -8 | -8 | -8 | -8 | -8 | 24 | -8 |
| $k_{55}$ | 24 | 24 | -8 | -8 | 24 | -8 | -8 | 24 | -8 | -8 | -8 | 24 | -8 | -8 | -8 |
| $k_{56}$ | 24 | 24 | -8 | 24 | -8 | -8 | 24 | -8 | -8 | 24 | -8 | -8 | -8 | -8 | -8 |
| $k_{57}$ | 24 | -8 | 24 | -8 | 24 | -8 | 24 | -8 | -8 | -8 | -8 | -8 | 24 | -8 | -8 |
| $k_{58}$ | 24 | -8 | 24 | 24 | -8 | -8 | -8 | 24 | -8 | -8 | 24 | -8 | -8 | -8 | -8 |
| $k_{59}$ | 24 | 24 | 24 | -8 | -8 | 24 | -8 | -8 | 24 | -8 | -8 | -8 | -8 | -8 | -8 |
| $k_{60}$ | 6 | -2 | -2 | 6 | 6 | 6 | -2 | -2 | -2 | -2 | -2 | -2 | -2 | 6 | -2 |
| $k_{61}$ | 6 | 6 | -2 | -2 | 6 | -2 | -2 | 6 | -2 | -2 | -2 | 6 | -2 | -2 | -2 |
| $k_{62}$ | 6 | 6 | -2 | 6 | -2 | -2 | 6 | -2 | -2 | 6 | -2 | -2 | -2 | -2 | -2 |
| $k_{63}$ | 6 | -2 | 6 | -2 | 6 | -2 | 6 | -2 | -2 | -2 | -2 | -2 | 6 | -2 | -2 |
| $k_{64}$ | 6 | -2 | 6 | 6 | -2 | -2 | -2 | 6 | -2 | -2 | 6 | -2 | -2 | -2 | -2 |
| $k_{65}$ | 6 | 6 | 6 | -2 | -2 | 6 | -2 | -2 | 6 | -2 | -2 | -2 | -2 | -2 | -2 |
| $k_{66}$ | 3 | 3 | -1 | 3 | -1 | -1 | 3 | -1 | -1 | 3 | -1 | -1 | -1 | -1 | -1 |
| $k_{67}$ | 3 | -1 | 3 | 3 | -1 | -1 | -1 | 3 | -1 | -1 | 3 | -1 | -1 | -1 | -1 |
| $k_{68}$ | 3 | 3 | 3 | -1 | -1 | 3 | -1 | -1 | 3 | -1 | -1 | -1 | -1 | -1 | -1 |
| $k_{69}$ | 9 | 9 | -3 | 9 | -3 | -3 | 9 | -3 | -3 | 9 | -3 | -3 | -3 | -3 | -3 |
| $k_{70}$ | 9 | -3 | 9 | 9 | -3 | -3 | -3 | 9 | -3 | -3 | 9 | -3 | -3 | -3 | -3 |
| $k_{71}$ | 9 | 9 | 9 | -3 | -3 | 9 | -3 | -3 | 9 | -3 | -3 | -3 | -3 | -3 | -3 |
| $k_{72}$ | 6 | -2 | -2 | -2 | 6 | -2 | -2 | -2 | 2 | 2 | 2 | -2 | -2 | -2 | 2 |
| $k_{73}$ | 6 | -2 | -2 | 6 | -2 | -2 | -2 | -2 | 2 | -2 | -2 | 2 | 2 | -2 | 2 |
| $k_{74}$ | 18 | -6 | -6 | -6 | 18 | -6 | -6 | -6 | 6 | 6 | 6 | -6 | -6 | -6 | 6 |
| $k_{75}$ | 6 | -2 | -2 | 6 | -2 | -2 | -2 | -2 | 2 | -2 | -2 | 2 | 2 | -2 | 2 |
| $k_{76}$ | 6 | 6 | -2 | -2 | -2 | -2 | -2 | -2 | -2 | -2 | 2 | -2 | 2 | 2 | 2 |
| $k_{77}$ | 6 | -2 | 6 | -2 | -2 | -2 | -2 | -2 | -2 | 2 | -2 | 2 | -2 | 2 | 2 |
| $k_{78}$ | 18 | -6 | -6 | -6 | 18 | -6 | -6 | -6 | 6 | 6 | 6 | -6 | -6 | -6 | 6 |
| $k_{79}$ | 18 | -6 | -6 | 18 | -6 | -6 | -6 | -6 | 6 | -6 | -6 | 6 | 6 | -6 | 6 |
| $k_{80}$ | 18 | 18 | -6 | -6 | -6 | -6 | -6 | -6 | -6 | -6 | 6 | -6 | 6 | 6 | 6 |
| $k_{81}$ | 18 | -6 | 18 | -6 | -6 | -6 | -6 | -6 | -6 | 6 | -6 | 6 | -6 | 6 | 6 |
| $k_{82}$ | 6 | -2 | -2 | -2 | 6 | -2 | -2 | -2 | 2 | 2 | 2 | -2 | -2 | -2 | 2 |
| $k_{83}$ | 18 | -6 | -6 | -6 | 18 | -6 | -6 | -6 | 6 | 6 | 6 | -6 | -6 | -6 | 6 |
| $k_{84}$ | 12 | -4 | -4 | -4 | 12 | -4 | -4 | -4 | 4 | 4 | 4 | -4 | -4 | -4 | 4 |
| $k_{85}$ | 12 | -4 | -4 | 12 | -4 | -4 | -4 | -4 | 4 | -4 | -4 | 4 | 4 | -4 | 4 |
| $k_{86}$ | 36 | -12 | -12 | -12 | 36 | -12 | -12 | -12 | 12 | 12 | 12 | -12 | -12 | -12 | 12 |
| $k_{87}$ | 12 | -4 | -4 | 12 | -4 | -4 | -4 | -4 | 4 | -4 | -4 | 4 | 4 | -4 | 4 |
| $k_{88}$ | 12 | 12 | -4 | -4 | -4 | -4 | -4 | -4 | -4 | -4 | 4 | -4 | 4 | 4 | 4 |

| **Site pattern** | XXXX | XXXY | XXYX | XYXX | YXXX | XXYY | XYXY | XYYX | XXYZ | XYXZ | XYZX | YXXZ | YXZX | YZXX | XYWW |
|---|---|---|---|---|---|---|---|---|---|---|---|---|---|---|---|
| $k_{89}$ | 12 | -4 | 12 | -4 | -4 | -4 | -4 | -4 | -4 | 4 | -4 | 4 | -4 | 4 | 4 |
| $k_{90}$ | 36 | -12 | -12 | -12 | 36 | -12 | -12 | -12 | 12 | 12 | 12 | -12 | -12 | -12 | 12 |
| $k_{91}$ | 24 | -8 | -8 | 24 | -8 | -8 | -8 | -8 | 8 | -8 | -8 | 8 | 8 | -8 | 8 |
| $k_{92}$ | 24 | 24 | -8 | -8 | -8 | -8 | -8 | -8 | -8 | -8 | 8 | -8 | 8 | 8 | 8 |
| $k_{93}$ | 24 | -8 | 24 | -8 | -8 | -8 | -8 | -8 | -8 | 8 | -8 | 8 | -8 | 8 | 8 |
| $k_{94}$ | 18 | -6 | -6 | -6 | 18 | -6 | -6 | -6 | 6 | 6 | 6 | -6 | -6 | -6 | 6 |
| $k_{95}$ | 6 | -2 | -2 | 6 | -2 | -2 | -2 | -2 | 2 | -2 | -2 | 2 | 2 | -2 | 2 |
| $k_{96}$ | 6 | 6 | -2 | -2 | -2 | -2 | -2 | -2 | -2 | -2 | 2 | -2 | 2 | 2 | 2 |
| $k_{97}$ | 6 | -2 | 6 | -2 | -2 | -2 | -2 | -2 | -2 | 2 | -2 | 2 | -2 | 2 | 2 |
| $k_{98}$ | 6 | -2 | -2 | 6 | -2 | -2 | -2 | -2 | 2 | -2 | -2 | 2 | 2 | -2 | 2 |
| $k_{99}$ | 6 | -2 | -2 | 6 | -2 | -2 | -2 | -2 | 2 | -2 | -2 | 2 | 2 | -2 | 2 |
| $k_{100}$ | 6 | 6 | -2 | -2 | -2 | -2 | -2 | -2 | -2 | -2 | 2 | -2 | 2 | 2 | 2 |
| $k_{101}$ | 6 | -2 | 6 | -2 | -2 | -2 | -2 | -2 | -2 | 2 | -2 | 2 | -2 | 2 | 2 |
| $k_{102}$ | 18 | 18 | -6 | -6 | -6 | -6 | -6 | -6 | -6 | -6 | 6 | -6 | 6 | 6 | 6 |
| $k_{103}$ | 18 | -6 | 18 | -6 | -6 | -6 | -6 | -6 | -6 | 6 | -6 | 6 | -6 | 6 | 6 |
| $k_{104}$ | 36 | -12 | -12 | 36 | -12 | -12 | -12 | -12 | 12 | -12 | -12 | 12 | 12 | -12 | 12 |
| $k_{105}$ | 36 | 36 | -12 | -12 | -12 | -12 | -12 | -12 | -12 | -12 | 12 | -12 | 12 | 12 | 12 |
| $k_{106}$ | 36 | -12 | 36 | -12 | -12 | -12 | -12 | -12 | -12 | 12 | -12 | 12 | -12 | 12 | 12 |
| $k_{107}$ | 36 | -12 | -12 | -12 | 36 | -12 | -12 | -12 | 12 | 12 | 12 | -12 | -12 | -12 | 12 |
| $k_{108}$ | 48 | -16 | -16 | 48 | -16 | -16 | -16 | -16 | 16 | -16 | -16 | 16 | 16 | -16 | 16 |
| $k_{109}$ | 48 | 48 | -16 | -16 | -16 | -16 | -16 | -16 | -16 | -16 | 16 | -16 | 16 | 16 | 16 |
| $k_{110}$ | 48 | -16 | 48 | -16 | -16 | -16 | -16 | -16 | -16 | 16 | -16 | 16 | -16 | 16 | 16 |
| $k_{111}$ | 12 | -4 | -4 | 12 | -4 | -4 | -4 | -4 | 4 | -4 | -4 | 4 | 4 | -4 | 4 |
| $k_{112}$ | 12 | 12 | -4 | -4 | -4 | -4 | -4 | -4 | -4 | -4 | 4 | -4 | 4 | 4 | 4 |
| $k_{113}$ | 12 | -4 | 12 | -4 | -4 | -4 | -4 | -4 | -4 | 4 | -4 | 4 | -4 | 4 | 4 |
| $k_{114}$ | 6 | 6 | -2 | -2 | -2 | -2 | -2 | -2 | -2 | -2 | 2 | -2 | 2 | 2 | 2 |
| $k_{115}$ | 6 | -2 | 6 | -2 | -2 | -2 | -2 | -2 | -2 | 2 | -2 | 2 | -2 | 2 | 2 |
| $k_{116}$ | 12 | -4 | -4 | 12 | -4 | -4 | -4 | -4 | 4 | -4 | -4 | 4 | 4 | -4 | 4 |
| $k_{117}$ | 12 | 12 | -4 | -4 | -4 | -4 | -4 | -4 | -4 | -4 | 4 | -4 | 4 | 4 | 4 |
| $k_{118}$ | 12 | -4 | 12 | -4 | -4 | -4 | -4 | -4 | -4 | 4 | -4 | 4 | -4 | 4 | 4 |
| $k_{119}$ | 36 | -12 | -12 | -12 | -12 | -12 | 12 | 12 | 12 | 0 | 0 | 0 | 0 | 12 | -12 |
| $k_{120}$ | 108 | -36 | -36 | -36 | -36 | 12 | 12 | 12 | 12 | 12 | 12 | 12 | 12 | 12 | -36 |
| $k_{121}$ | 180 | -60 | -60 | -60 | -60 | 20 | 20 | 20 | 20 | 20 | 20 | 20 | 20 | 20 | -60 |
| $k_{122}$ | 36 | -12 | -12 | -12 | -12 | 4 | 4 | 4 | 4 | 4 | 4 | 4 | 4 | 4 | -12 |
| $k_{123}$ | 12 | -4 | -4 | -4 | -4 | -4 | 4 | 4 | 4 | 0 | 0 | 0 | 0 | 4 | -4 |
| $k_{124}$ | 36 | -12 | -12 | -12 | -12 | 4 | 4 | 4 | 4 | 4 | 4 | 4 | 4 | 4 | -12 |
| $k_{125}$ | 9 | -3 | -3 | -3 | -3 | 9 | 1 | 1 | -3 | 1 | 1 | 1 | 1 | -3 | 1 |
| $k_{126}$ | 27 | -9 | -9 | -9 | -9 | 11 | 11 | 11 | -1 | -1 | -1 | -1 | -1 | -1 | 3 |
| $k_{127}$ | 27 | -9 | -9 | -9 | -9 | 11 | 11 | 11 | -1 | -1 | -1 | -1 | -1 | -1 | 3 |
| $k_{128}$ | 27 | -9 | -9 | -9 | -9 | 11 | 11 | 11 | -1 | -1 | -1 | -1 | -1 | -1 | 3 |
| $k_{129}$ | 18 | -6 | -6 | -6 | -6 | 18 | 2 | 2 | -6 | 2 | 2 | 2 | 2 | -6 | 2 |
| $k_{130}$ | 54 | -18 | -18 | -18 | -18 | 22 | 22 | 22 | -2 | -2 | -2 | -2 | -2 | -2 | 6 |
| $k_{131}$ | 108 | -36 | -36 | -36 | -36 | 44 | 44 | 44 | -4 | -4 | -4 | -4 | -4 | -4 | 12 |
| $k_{132}$ | 9 | -3 | -3 | -3 | -3 | 9 | 1 | 1 | -3 | 1 | 1 | 1 | 1 | -3 | 1 |
| $k_{133}$ | 27 | -9 | -9 | -9 | -9 | 11 | 11 | 11 | -1 | -1 | -1 | -1 | -1 | -1 | 3 |

#### 2 Site pattern probabilities given the gene tree

Here we have a slight change in notation from the main text. $p_{ik}^{(l)}$ is used as shorthand for $p_{ik}(d_l)$ throughout this section.

$$\begin{aligned} p_{xxxx} = \frac{1}{4}(& p_{ii}^{(1)}p_{ii}^{(2)}p_{ii}^{(3)}p_{ii}^{(4)}p_{ii}^{(5)} + 3p_{ii}^{(1)}p_{ii}^{(2)}p_{ij}^{(3)}p_{ij}^{(4)}p_{ij}^{(5)} + 3p_{ij}^{(1)}p_{ij}^{(2)}p_{ii}^{(3)}p_{ii}^{(4)}p_{ij}^{(5)} + 3p_{ij}^{(1)}p_{ij}^{(2)}p_{ij}^{(3)}p_{ij}^{(4)}p_{ii}^{(5)} \\ & + 6p_{ij}^{(1)}p_{ij}^{(2)}p_{ij}^{(3)}p_{ij}^{(4)}p_{ij}^{(5)}) \end{aligned} \tag{1}$$

$$\begin{aligned} p_{xxxy} = \frac{1}{4}(& p_{ii}^{(1)}p_{ii}^{(2)}p_{ii}^{(3)}p_{ij}^{(4)}p_{ii}^{(5)} + p_{ii}^{(1)}p_{ii}^{(2)}p_{ij}^{(3)}p_{ii}^{(4)}p_{ij}^{(5)} + p_{ij}^{(1)}p_{ij}^{(2)}p_{ij}^{(3)}p_{ii}^{(4)}p_{ii}^{(5)} + 2p_{ii}^{(1)}p_{ii}^{(2)}p_{ij}^{(3)}p_{ij}^{(4)}p_{ij}^{(5)} \\ & + 2p_{ij}^{(1)}p_{ij}^{(2)}p_{ij}^{(3)}p_{ij}^{(4)}p_{ii}^{(5)} + 2p_{ij}^{(1)}p_{ij}^{(2)}p_{ij}^{(3)}p_{ii}^{(4)}p_{ij}^{(5)} + 3p_{ij}^{(1)}p_{ij}^{(2)}p_{ii}^{(3)}p_{ij}^{(4)}p_{ij}^{(5)} + 4p_{ij}^{(1)}p_{ij}^{(2)}p_{ij}^{(3)}p_{ij}^{(4)}p_{ij}^{(5)}) \end{aligned} \tag{2}$$

$$\begin{aligned} p_{xxyx} = \frac{1}{4}(& p_{ii}^{(1)}p_{ij}^{(2)}p_{ij}^{(3)}p_{ii}^{(4)}p_{ii}^{(5)} + p_{ii}^{(1)}p_{ii}^{(2)}p_{ii}^{(3)}p_{ij}^{(4)}p_{ij}^{(5)} + p_{ij}^{(1)}p_{ij}^{(2)}p_{ii}^{(3)}p_{ij}^{(4)}p_{ii}^{(5)} + 2p_{ii}^{(1)}p_{ii}^{(2)}p_{ij}^{(3)}p_{ij}^{(4)}p_{ij}^{(5)} \\ & + 2p_{ij}^{(1)}p_{ij}^{(2)}p_{ij}^{(3)}p_{ij}^{(4)}p_{ii}^{(5)} + 2p_{ij}^{(1)}p_{ij}^{(2)}p_{ii}^{(3)}p_{ij}^{(4)}p_{ij}^{(5)} + 3p_{ij}^{(1)}p_{ij}^{(2)}p_{ij}^{(3)}p_{ii}^{(4)}p_{ij}^{(5)} + 4p_{ij}^{(1)}p_{ij}^{(2)}p_{ij}^{(3)}p_{ij}^{(4)}p_{ij}^{(5)}) \end{aligned} \tag{3}$$

$$\begin{aligned} p_{xyxx} = \frac{1}{4}(& p_{ii}^{(1)}p_{ij}^{(2)}p_{ii}^{(3)}p_{ii}^{(4)}p_{ii}^{(5)} + p_{ij}^{(1)}p_{ii}^{(2)}p_{ii}^{(3)}p_{ii}^{(4)}p_{ij}^{(5)} + p_{ij}^{(1)}p_{ii}^{(2)}p_{ij}^{(3)}p_{ij}^{(4)}p_{ii}^{(5)} + 2p_{ij}^{(1)}p_{ij}^{(2)}p_{ii}^{(3)}p_{ii}^{(4)}p_{ij}^{(5)} \\ & + 2p_{ij}^{(1)}p_{ij}^{(2)}p_{ij}^{(3)}p_{ij}^{(4)}p_{ii}^{(5)} + 2p_{ij}^{(1)}p_{ii}^{(2)}p_{ij}^{(3)}p_{ij}^{(4)}p_{ij}^{(5)} + 3p_{ii}^{(1)}p_{ij}^{(2)}p_{ij}^{(3)}p_{ij}^{(4)}p_{ij}^{(5)} + 4p_{ij}^{(1)}p_{ij}^{(2)}p_{ij}^{(3)}p_{ij}^{(4)}p_{ij}^{(5)}) \end{aligned} \tag{4}$$

$$\begin{aligned} p_{yxxx} = \frac{1}{4}(& p_{ii}^{(1)}p_{ij}^{(2)}p_{ij}^{(3)}p_{ij}^{(4)}p_{ii}^{(5)} + p_{ii}^{(1)}p_{ij}^{(2)}p_{ii}^{(3)}p_{ii}^{(4)}p_{ij}^{(5)} + p_{ij}^{(1)}p_{ii}^{(2)}p_{ii}^{(3)}p_{ii}^{(4)}p_{ii}^{(5)} + 2p_{ij}^{(1)}p_{ij}^{(2)}p_{ii}^{(3)}p_{ii}^{(4)}p_{ij}^{(5)} \\ & + 2p_{ij}^{(1)}p_{ij}^{(2)}p_{ij}^{(3)}p_{ij}^{(4)}p_{ii}^{(5)} + 2p_{ii}^{(1)}p_{ij}^{(2)}p_{ij}^{(3)}p_{ij}^{(4)}p_{ij}^{(5)} + 3p_{ij}^{(1)}p_{ii}^{(2)}p_{ij}^{(3)}p_{ij}^{(4)}p_{ij}^{(5)} + 4p_{ij}^{(1)}p_{ij}^{(2)}p_{ij}^{(3)}p_{ij}^{(4)}p_{ij}^{(5)}) \end{aligned} \tag{5}$$

$$\begin{aligned} p_{xxyy} = \frac{1}{4}(& p_{ii}^{(1)}p_{ii}^{(2)}p_{ij}^{(3)}p_{ij}^{(4)}p_{ii}^{(5)} + p_{ii}^{(1)}p_{ii}^{(2)}p_{ii}^{(3)}p_{ii}^{(4)}p_{ij}^{(5)} + p_{ij}^{(1)}p_{ij}^{(2)}p_{ii}^{(3)}p_{ii}^{(4)}p_{ii}^{(5)} + 2p_{ii}^{(1)}p_{ii}^{(2)}p_{ij}^{(3)}p_{ij}^{(4)}p_{ij}^{(5)} \\ & + 2p_{ij}^{(1)}p_{ij}^{(2)}p_{ii}^{(3)}p_{ii}^{(4)}p_{ij}^{(5)} + 2p_{ij}^{(1)}p_{ij}^{(2)}p_{ij}^{(3)}p_{ij}^{(4)}p_{ii}^{(5)} + 7p_{ij}^{(1)}p_{ij}^{(2)}p_{ij}^{(3)}p_{ij}^{(4)}p_{ij}^{(5)}) \end{aligned} \tag{6}$$

$$\begin{aligned} p_{xyxy} = \frac{1}{4}(& p_{ii}^{(1)}p_{ij}^{(2)}p_{ii}^{(3)}p_{ij}^{(4)}p_{ii}^{(5)} + p_{ii}^{(1)}p_{ij}^{(2)}p_{ij}^{(3)}p_{ii}^{(4)}p_{ij}^{(5)} + p_{ij}^{(1)}p_{ii}^{(2)}p_{ii}^{(3)}p_{ij}^{(4)}p_{ij}^{(5)} + p_{ij}^{(1)}p_{ii}^{(2)}p_{ij}^{(3)}p_{ii}^{(4)}p_{ii}^{(5)} \\ & + 2p_{ii}^{(1)}p_{ij}^{(2)}p_{ij}^{(3)}p_{ij}^{(4)}p_{ij}^{(5)} + 2p_{ij}^{(1)}p_{ii}^{(2)}p_{ij}^{(3)}p_{ij}^{(4)}p_{ij}^{(5)} + 2p_{ij}^{(1)}p_{ij}^{(2)}p_{ii}^{(3)}p_{ij}^{(4)}p_{ij}^{(5)} + 2p_{ij}^{(1)}p_{ij}^{(2)}p_{ij}^{(3)}p_{ii}^{(4)}p_{ij}^{(5)} \\ & + 2p_{ij}^{(1)}p_{ij}^{(2)}p_{ij}^{(3)}p_{ij}^{(4)}p_{ii}^{(5)} + 2p_{ij}^{(1)}p_{ij}^{(2)}p_{ij}^{(3)}p_{ij}^{(4)}p_{ij}^{(5)}) \end{aligned} \tag{7}$$

$$
\begin{aligned}
p_{xyyx} = \frac{1}{4}(&p_{ij}^{(1)} p_{ii}^{(2)} p_{ij}^{(3)} p_{ii}^{(4)} p_{ij}^{(5)} + p_{ii}^{(1)} p_{ij}^{(2)} p_{ij}^{(3)} p_{ii}^{(4)} p_{ii}^{(5)} + p_{ii}^{(1)} p_{ij}^{(2)} p_{ii}^{(3)} p_{ij}^{(4)} p_{ij}^{(5)} + p_{ij}^{(1)} p_{ii}^{(2)} p_{ii}^{(3)} p_{ij}^{(4)} p_{ii}^{(5)} \\
&+ 2p_{ii}^{(1)} p_{ij}^{(2)} p_{ij}^{(3)} p_{ij}^{(4)} p_{ij}^{(5)} + 2p_{ij}^{(1)} p_{ii}^{(2)} p_{ij}^{(3)} p_{ij}^{(4)} p_{ij}^{(5)} + 2p_{ij}^{(1)} p_{ij}^{(2)} p_{ii}^{(3)} p_{ij}^{(4)} p_{ij}^{(5)} + 2p_{ij}^{(1)} p_{ij}^{(2)} p_{ij}^{(3)} p_{ii}^{(4)} p_{ij}^{(5)} \\
&+ 2p_{ij}^{(1)} p_{ij}^{(2)} p_{ij}^{(3)} p_{ij}^{(4)} p_{ii}^{(5)} + 2p_{ij}^{(1)} p_{ij}^{(2)} p_{ij}^{(3)} p_{ij}^{(4)} p_{ij}^{(5)})
\end{aligned} \tag{8}
$$

$$
\begin{aligned}
p_{xxyz} = \frac{1}{4}(&p_{ii}^{(1)} p_{ii}^{(2)} p_{ij}^{(3)} p_{ij}^{(4)} p_{ij}^{(5)} + p_{ij}^{(1)} p_{ij}^{(2)} p_{ij}^{(3)} p_{ij}^{(4)} p_{ii}^{(5)} + p_{ii}^{(1)} p_{ii}^{(2)} p_{ij}^{(3)} p_{ii}^{(4)} p_{ij}^{(5)} + p_{ij}^{(1)} p_{ij}^{(2)} p_{ij}^{(3)} p_{ii}^{(4)} p_{ii}^{(5)} \\
&+ p_{ii}^{(1)} p_{ii}^{(2)} p_{ii}^{(3)} p_{ij}^{(4)} p_{ij}^{(5)} + p_{ij}^{(1)} p_{ij}^{(2)} p_{ii}^{(3)} p_{ij}^{(4)} p_{ii}^{(5)} + p_{ii}^{(1)} p_{ii}^{(2)} p_{ij}^{(3)} p_{ij}^{(4)} p_{ii}^{(5)} + 2p_{ij}^{(1)} p_{ij}^{(2)} p_{ii}^{(3)} p_{ij}^{(4)} p_{ij}^{(5)} \\
&+ 2p_{ij}^{(1)} p_{ij}^{(2)} p_{ij}^{(3)} p_{ii}^{(4)} p_{ij}^{(5)} + 5p_{ij}^{(1)} p_{ij}^{(2)} p_{ij}^{(3)} p_{ij}^{(4)} p_{ij}^{(5)})
\end{aligned} \tag{9}
$$

$$
\begin{aligned}
p_{xyxz} = \frac{1}{4}(&p_{ij}^{(1)} p_{ij}^{(2)} p_{ij}^{(3)} p_{ij}^{(4)} p_{ii}^{(5)} + p_{ij}^{(1)} p_{ij}^{(2)} p_{ij}^{(3)} p_{ii}^{(4)} p_{ii}^{(5)} + p_{ij}^{(1)} p_{ij}^{(2)} p_{ij}^{(3)} p_{ii}^{(4)} p_{ij}^{(5)} + p_{ij}^{(1)} p_{ii}^{(2)} p_{ij}^{(3)} p_{ij}^{(4)} p_{ii}^{(5)} \\
&+ p_{ij}^{(1)} p_{ii}^{(2)} p_{ij}^{(3)} p_{ij}^{(4)} p_{ij}^{(5)} + p_{ii}^{(1)} p_{ij}^{(2)} p_{ii}^{(3)} p_{ij}^{(4)} p_{ii}^{(5)} + p_{ii}^{(1)} p_{ij}^{(2)} p_{ij}^{(3)} p_{ii}^{(4)} p_{ij}^{(5)} + p_{ij}^{(1)} p_{ii}^{(2)} p_{ii}^{(3)} p_{ij}^{(4)} p_{ij}^{(5)} \\
&+ p_{ij}^{(1)} p_{ii}^{(2)} p_{ij}^{(3)} p_{ii}^{(4)} p_{ij}^{(5)} + 2p_{ij}^{(1)} p_{ij}^{(2)} p_{ii}^{(3)} p_{ij}^{(4)} p_{ij}^{(5)} + 2p_{ii}^{(1)} p_{ij}^{(2)} p_{ij}^{(3)} p_{ij}^{(4)} p_{ij}^{(5)} + 3p_{ij}^{(1)} p_{ij}^{(2)} p_{ij}^{(3)} p_{ij}^{(4)} p_{ij}^{(5)})
\end{aligned} \tag{10}
$$

$$
\begin{aligned}
p_{xyzx} = \frac{1}{4}(&p_{ij}^{(1)} p_{ij}^{(2)} p_{ij}^{(3)} p_{ij}^{(4)} p_{ii}^{(5)} + p_{ij}^{(1)} p_{ij}^{(2)} p_{ii}^{(3)} p_{ij}^{(4)} p_{ij}^{(5)} + p_{ij}^{(1)} p_{ij}^{(2)} p_{ii}^{(3)} p_{ij}^{(4)} p_{ii}^{(5)} + p_{ij}^{(1)} p_{ii}^{(2)} p_{ij}^{(3)} p_{ij}^{(4)} p_{ii}^{(5)} \\
&+ p_{ij}^{(1)} p_{ii}^{(2)} p_{ij}^{(3)} p_{ij}^{(4)} p_{ij}^{(5)} + p_{ij}^{(1)} p_{ii}^{(2)} p_{ii}^{(3)} p_{ij}^{(4)} p_{ij}^{(5)} + p_{ij}^{(1)} p_{ii}^{(2)} p_{ij}^{(3)} p_{ii}^{(4)} p_{ij}^{(5)} + p_{ii}^{(1)} p_{ij}^{(2)} p_{ij}^{(3)} p_{ii}^{(4)} p_{ii}^{(5)} \\
&+ p_{ii}^{(1)} p_{ij}^{(2)} p_{ii}^{(3)} p_{ij}^{(4)} p_{ij}^{(5)} + 2p_{ij}^{(1)} p_{ij}^{(2)} p_{ij}^{(3)} p_{ii}^{(4)} p_{ij}^{(5)} + 2p_{ii}^{(1)} p_{ij}^{(2)} p_{ij}^{(3)} p_{ij}^{(4)} p_{ij}^{(5)} + 3p_{ij}^{(1)} p_{ij}^{(2)} p_{ij}^{(3)} p_{ij}^{(4)} p_{ij}^{(5)})
\end{aligned} \tag{11}
$$

$$
\begin{aligned}
p_{yxxz} = \frac{1}{4}(&p_{ij}^{(1)} p_{ij}^{(2)} p_{ij}^{(3)} p_{ij}^{(4)} p_{ii}^{(5)} + p_{ij}^{(1)} p_{ij}^{(2)} p_{ij}^{(3)} p_{ii}^{(4)} p_{ii}^{(5)} + p_{ij}^{(1)} p_{ij}^{(2)} p_{ij}^{(3)} p_{ii}^{(4)} p_{ij}^{(5)} + p_{ii}^{(1)} p_{ij}^{(2)} p_{ij}^{(3)} p_{ij}^{(4)} p_{ij}^{(5)} \\
&+ p_{ii}^{(1)} p_{ij}^{(2)} p_{ij}^{(3)} p_{ii}^{(4)} p_{ij}^{(5)} + p_{ij}^{(1)} p_{ii}^{(2)} p_{ij}^{(3)} p_{ii}^{(4)} p_{ij}^{(5)} + p_{ii}^{(1)} p_{ij}^{(2)} p_{ii}^{(3)} p_{ij}^{(4)} p_{ij}^{(5)} + p_{ij}^{(1)} p_{ii}^{(2)} p_{ii}^{(3)} p_{ij}^{(4)} p_{ii}^{(5)} \\
&+ p_{ii}^{(1)} p_{ij}^{(2)} p_{ij}^{(3)} p_{ij}^{(4)} p_{ii}^{(5)} + 2p_{ij}^{(1)} p_{ii}^{(2)} p_{ij}^{(3)} p_{ij}^{(4)} p_{ij}^{(5)} + 2p_{ij}^{(1)} p_{ij}^{(2)} p_{ii}^{(3)} p_{ij}^{(4)} p_{ij}^{(5)} + 3p_{ij}^{(1)} p_{ij}^{(2)} p_{ij}^{(3)} p_{ij}^{(4)} p_{ij}^{(5)})
\end{aligned} \tag{12}
$$

$$
\begin{aligned}
p_{yxzx} = \frac{1}{4}(&p_{ij}^{(1)} p_{ij}^{(2)} p_{ij}^{(3)} p_{ij}^{(4)} p_{ii}^{(5)} + p_{ij}^{(1)} p_{ij}^{(2)} p_{ii}^{(3)} p_{ij}^{(4)} p_{ij}^{(5)} + p_{ij}^{(1)} p_{ij}^{(2)} p_{ii}^{(3)} p_{ij}^{(4)} p_{ii}^{(5)} + p_{ii}^{(1)} p_{ij}^{(2)} p_{ij}^{(3)} p_{ij}^{(4)} p_{ij}^{(5)} \\
&+ p_{ii}^{(1)} p_{ij}^{(2)} p_{ij}^{(3)} p_{ii}^{(4)} p_{ij}^{(5)} + p_{ij}^{(1)} p_{ii}^{(2)} p_{ii}^{(3)} p_{ij}^{(4)} p_{ij}^{(5)} + p_{ij}^{(1)} p_{ii}^{(2)} p_{ij}^{(3)} p_{ii}^{(4)} p_{ii}^{(5)} + p_{ii}^{(1)} p_{ij}^{(2)} p_{ii}^{(3)} p_{ij}^{(4)} p_{ij}^{(5)} \\
&+ p_{ii}^{(1)} p_{ij}^{(2)} p_{ij}^{(3)} p_{ij}^{(4)} p_{ii}^{(5)} + 2p_{ij}^{(1)} p_{ii}^{(2)} p_{ij}^{(3)} p_{ij}^{(4)} p_{ij}^{(5)} + 2p_{ij}^{(1)} p_{ij}^{(2)} p_{ij}^{(3)} p_{ii}^{(4)} p_{ij}^{(5)} + 3p_{ij}^{(1)} p_{ij}^{(2)} p_{ij}^{(3)} p_{ij}^{(4)} p_{ij}^{(5)})
\end{aligned} \tag{13}
$$

$$
\begin{aligned}
p_{yzxx} = \frac{1}{4}(&p_{ij}^{(1)} p_{ij}^{(2)} p_{ii}^{(3)} p_{ii}^{(4)} p_{ij}^{(5)} + p_{ij}^{(1)} p_{ij}^{(2)} p_{ij}^{(3)} p_{ij}^{(4)} p_{ii}^{(5)} + p_{ij}^{(1)} p_{ij}^{(2)} p_{ii}^{(3)} p_{ii}^{(4)} p_{ii}^{(5)} + p_{ij}^{(1)} p_{ii}^{(2)} p_{ii}^{(3)} p_{ii}^{(4)} p_{ij}^{(5)} \\
&+ p_{ij}^{(1)} p_{ii}^{(2)} p_{ij}^{(3)} p_{ij}^{(4)} p_{ii}^{(5)} + p_{ii}^{(1)} p_{ij}^{(2)} p_{ij}^{(3)} p_{ij}^{(4)} p_{ii}^{(5)} + p_{ii}^{(1)} p_{ij}^{(2)} p_{ii}^{(3)} p_{ii}^{(4)} p_{ij}^{(5)} + 2p_{ii}^{(1)} p_{ij}^{(2)} p_{ij}^{(3)} p_{ij}^{(4)} p_{ij}^{(5)} \\
&+ 2p_{ij}^{(1)} p_{ii}^{(2)} p_{ij}^{(3)} p_{ij}^{(4)} p_{ij}^{(5)} + 5p_{ij}^{(1)} p_{ij}^{(2)} p_{ij}^{(3)} p_{ij}^{(4)} p_{ij}^{(5)})
\end{aligned} \tag{14}
$$

$$
\begin{aligned}
p_{xyzw} = \frac{1}{4}(&p_{ij}^{(1)}p_{ij}^{(2)}p_{ii}^{(3)}p_{ij}^{(4)}p_{ij}^{(5)} + p_{ij}^{(1)}p_{ij}^{(2)}p_{ij}^{(3)}p_{ii}^{(4)}p_{ii}^{(5)} + p_{ij}^{(1)}p_{ij}^{(2)}p_{ij}^{(3)}p_{ii}^{(4)}p_{ij}^{(5)} + p_{ij}^{(1)}p_{ij}^{(2)}p_{ii}^{(3)}p_{ij}^{(4)}p_{ii}^{(5)} \\
&+ p_{ii}^{(1)}p_{ij}^{(2)}p_{ij}^{(3)}p_{ij}^{(4)}p_{ij}^{(5)} + p_{ij}^{(1)}p_{ii}^{(2)}p_{ij}^{(3)}p_{ij}^{(4)}p_{ii}^{(5)} + p_{ij}^{(1)}p_{ii}^{(2)}p_{ij}^{(3)}p_{ij}^{(4)}p_{ij}^{(5)} + p_{ii}^{(1)}p_{ij}^{(2)}p_{ij}^{(3)}p_{ii}^{(4)}p_{ij}^{(5)} \\
&+ p_{ij}^{(1)}p_{ii}^{(2)}p_{ii}^{(3)}p_{ij}^{(4)}p_{ij}^{(5)} + p_{ij}^{(1)}p_{ij}^{(2)}p_{ij}^{(3)}p_{ii}^{(4)}p_{ij}^{(5)} + p_{ii}^{(1)}p_{ij}^{(2)}p_{ii}^{(3)}p_{ij}^{(4)}p_{ij}^{(5)} + p_{ii}^{(1)}p_{ij}^{(2)}p_{ij}^{(3)}p_{ij}^{(4)}p_{ii}^{(5)} \\
&+ 4p_{ij}^{(1)}p_{ij}^{(2)}p_{ij}^{(3)}p_{ij}^{(4)}p_{ij}^{(5)})
\end{aligned} \tag{15}
$$

### 3 Table of $g_0$, $g_1$, $g_2$,and $g_3$ values.

#### 3.1 Symmetric species tree

| | **History 1** | | | |
|---|---|---|---|---|
| $d^{-(d_1+d_2)}$ | $g_0 = exp[-\alpha\tau_2(\gamma_A + \gamma_B - 2\gamma_{AB})]$ | $g_1 = 0$ | $g_2 = 2\alpha\gamma_{AB}$ | $g_3 = 0$ |
| $d^{-(d_3+d_4)}$ | $g_0 = exp[-\alpha\tau_1(\gamma_C + \gamma_D - 2\gamma_{CD})]$ | $g_1 = 2\alpha\gamma_{CD}$ | $g_2 = 0$ | $g_3 = 0$ |
| $d^{-(d_1+d_3+d_5)}$ | $g_0 = exp[-\alpha\tau_1(\gamma_C - \gamma_{CD}) - \alpha\tau_2(\gamma_A - \gamma_{AB}) - \alpha\tau_3(\gamma_{AB} + \gamma_{CD} - 2)]$ | $g_1 = 0$ | $g_2 = 0$ | $g_3 = 2\alpha$ |
| $d^{-(d_1+d_4+d_5)}$ | $g_0 = exp[-\alpha\tau_1(\gamma_D - \gamma_{CD}) - \alpha\tau_2(\gamma_A - \gamma_{AB}) - \alpha\tau_3(\gamma_{AB} + \gamma_{CD} - 2)]$ | $g_1 = 0$ | $g_2 = 0$ | $g_3 = 2\alpha$ |
| $d^{-(d_2+d_3+d_5)}$ | $g_0 = exp[-\alpha\tau_1(\gamma_C - \gamma_{CD}) - \alpha\tau_2(\gamma_B - \gamma_{AB}) - \alpha\tau_3(\gamma_{AB} + \gamma_{CD} - 2)]$ | $g_1 = 0$ | $g_2 = 0$ | $g_3 = 2\alpha$ |
| $d^{-(d_2+d_4+d_5)}$ | $g_0 = exp[-\alpha\tau_1(\gamma_D - \gamma_{CD}) - \alpha\tau_2(\gamma_B - \gamma_{AB}) - \alpha\tau_3(\gamma_{AB} + \gamma_{CD} - 2)]$ | $g_1 = 0$ | $g_2 = 0$ | $g_3 = 2\alpha$ |
| $d^{-(d_1+d_2+d_3+d_4)}$ | $g_0 = exp[-\alpha\tau_1(\gamma_C + \gamma_D - 2\gamma_{CD}) - \alpha\tau_2(\gamma_A + \gamma_B - 2\gamma_{AB})]$ | $g_1 = 2\alpha\gamma_{CD}$ | $g_2 = 2\alpha\gamma_{AB}$ | $g_3 = 0$ |
| $d^{-(d_1+d_2+d_3+d_5)}$ | $g_0 = exp[-\alpha\tau_1(\gamma_C - \gamma_{CD}) - \alpha\tau_2(\gamma_A + \gamma_B - 2\gamma_{AB}) - \alpha\tau_3(\gamma_{AB} + \gamma_{CD} - 2)]$ | $g_1 = 0$ | $g_2 = \alpha\gamma_{AB}$ | $g_3 = 2\alpha$ |
| $d^{-(d_1+d_2+d_4+d_5)}$ | $g_0 = exp[-\alpha\tau_1(\gamma_D - \gamma_{CD}) - \alpha\tau_2(\gamma_A + \gamma_B - 2\gamma_{AB}) - \alpha\tau_3(\gamma_{AB} + \gamma_{CD} - 2)]$ | $g_1 = 0$ | $g_2 = \alpha\gamma_{AB}$ | $g_3 = 2\alpha$ |
| $d^{-(d_1+d_3+d_4+d_5)}$ | $g_0 = exp[-\alpha\tau_1(\gamma_C + \gamma_D - 2\gamma_{CD}) - \alpha\tau_2(\gamma_A - \gamma_{AB}) - \alpha\tau_3(\gamma_{AB} + \gamma_{CD} - 2)]$ | $g_1 = \alpha\gamma_{CD}$ | $g_2 = 0$ | $g_3 = 2\alpha$ |
| $d^{-(d_2+d_3+d_4+d_5)}$ | $g_0 = exp[-\alpha\tau_1(\gamma_C + \gamma_D - 2\gamma_{CD}) - \alpha\tau_2(\gamma_B - \gamma_{AB}) - \alpha\tau_3(\gamma_{AB} + \gamma_{CD} - 2)]$ | $g_1 = \alpha\gamma_{CD}$ | $g_2 = 0$ | $g_3 = 2\alpha$ |
| $d^{-(d_1+d_2+d_3+d_4+d_5)}$ | $g_0 = exp[-\alpha\tau_1(\gamma_C + \gamma_D - 2\gamma_{CD}) - \alpha\tau_2(\gamma_A + \gamma_B - 2\gamma_{AB}) - \alpha\tau_3(\gamma_{AB} + \gamma_{CD} - 2)]$ | $g_1 = \alpha\gamma_{CD}$ | $g_2 = \alpha\gamma_{AB}$ | $g_3 = 2\alpha$ |

| | **History 2** | | | |
|---|---|---|---|---|
| $d^{-(d_1+d_2)}$ | $g_0 = exp[-\alpha\tau_2(\gamma_A + \gamma_B - 2\gamma_{AB}) - 2\alpha\tau_3(\gamma_{AB} - 1)]$ | $g_1 = 0$ | $g_2 = 2\alpha$ | $g_3 = 0$ |
| $d^{-(d_3+d_4)}$ | $g_0 = exp[-\alpha\tau_1(\gamma_C + \gamma_D - 2\gamma_{CD})]$ | $g_1 = 2\alpha\gamma_{CD}$ | $g_2 = 0$ | $g_3 = 0$ |
| $d^{-(d_1+d_3+d_5)}$ | $g_0 = exp[-\alpha\tau_1(\gamma_C - \gamma_{CD}) - \alpha\tau_2(\gamma_A - \gamma_{AB}) - \alpha\tau_3(\gamma_{AB} + \gamma_{CD} - 2)]$ | $g_1 = 0$ | $g_2 = 0$ | $g_3 = 2\alpha$ |
| $d^{-(d_1+d_4+d_5)}$ | $g_0 = exp[-\alpha\tau_1(\gamma_D - \gamma_{CD}) - \alpha\tau_2(\gamma_A - \gamma_{AB}) - \alpha\tau_3(\gamma_{AB} + \gamma_{CD} - 2)]$ | $g_1 = 0$ | $g_2 = 0$ | $g_3 = 2\alpha$ |
| $d^{-(d_2+d_3+d_5)}$ | $g_0 = exp[-\alpha\tau_1(\gamma_C - \gamma_{CD}) - \alpha\tau_2(\gamma_B - \gamma_{AB}) - \alpha\tau_3(\gamma_{AB} + \gamma_{CD} - 2)]$ | $g_1 = 0$ | $g_2 = 0$ | $g_3 = 2\alpha$ |
| $d^{-(d_2+d_4+d_5)}$ | $g_0 = exp[-\alpha\tau_1(\gamma_D - \gamma_{CD}) - \alpha\tau_2(\gamma_B - \gamma_{AB}) - \alpha\tau_3(\gamma_{AB} + \gamma_{CD} - 2)]$ | $g_1 = 0$ | $g_2 = 0$ | $g_3 = 2\alpha$ |
| $d^{-(d_1+d_2+d_3+d_4)}$ | $g_0 = exp[-\alpha\tau_1(\gamma_C + \gamma_D - 2\gamma_{CD}) - \alpha\tau_2(\gamma_A + \gamma_B - 2\gamma_{AB}) - 2\alpha\tau_3(\gamma_{AB} - 1)]$ | $g_1 = 2\alpha\gamma_{CD}$ | $g_2 = 2\alpha$ | $g_3 = 0$ |
| $d^{-(d_1+d_2+d_3+d_5)}$ | $g_0 = exp[-\alpha\tau_1(\gamma_C - \gamma_{CD}) - \alpha\tau_2(\gamma_A + \gamma_B - 2\gamma_{AB}) - \alpha\tau_3(2\gamma_{AB} + \gamma_{CD} - 3)]$ | $g_1 = 0$ | $g_2 = \alpha$ | $g_3 = 2\alpha$ |
| $d^{-(d_1+d_2+d_4+d_5)}$ | $g_0 = exp[-\alpha\tau_1(\gamma_D - \gamma_{CD}) - \alpha\tau_2(\gamma_A + \gamma_B - 2\gamma_{AB}) - \alpha\tau_3(2\gamma_{AB} + \gamma_{CD} - 3)]$ | $g_1 = 0$ | $g_2 = \alpha$ | $g_3 = 2\alpha$ |
| $d^{-(d_1+d_3+d_4+d_5)}$ | $g_0 = exp[-\alpha\tau_1(\gamma_C + \gamma_D - 2\gamma_{CD}) - \alpha\tau_2(\gamma_A - \gamma_{AB}) - \alpha\tau_3(\gamma_{AB} + \gamma_{CD} - 2)]$ | $g_1 = \alpha\gamma_{CD}$ | $g_2 = 0$ | $g_3 = 2\alpha$ |
| $d^{-(d_2+d_3+d_4+d_5)}$ | $g_0 = exp[-\alpha\tau_1(\gamma_C + \gamma_D - 2\gamma_{CD}) - \alpha\tau_2(\gamma_B - \gamma_{AB}) - \alpha\tau_3(\gamma_{AB} + \gamma_{CD} - 2)]$ | $g_1 = \alpha\gamma_{CD}$ | $g_2 = 0$ | $g_3 = 2\alpha$ |
| $d^{-(d_1+d_2+d_3+d_4+d_5)}$ | $g_0 = exp[-\alpha\tau_1(\gamma_C + \gamma_D - 2\gamma_{CD}) - \alpha\tau_2(\gamma_A + \gamma_B - 2\gamma_{AB}) - \alpha\tau_3(2\gamma_{AB} + \gamma_{CD} - 3)]$ | $g_1 = \alpha\gamma_{CD}$ | $g_2 = \alpha$ | $g_3 = 2\alpha$ |

| | **History 3** | | | |
|---|---|---|---|---|
| $d^{-(d_1+d_2)}$ | $g_0 = exp[-\alpha\tau_2(\gamma_A + \gamma_B - 2\gamma_{AB})]$ | $g_1 = 0$ | $g_2 = 2\alpha\gamma_{AB}$ | $g_3 = 0$ |
| $d^{-(d_3+d_4)}$ | $g_0 = exp[-\alpha\tau_1(\gamma_C + \gamma_D - 2\gamma_{CD}) - 2\alpha\tau_3(\gamma_{CD} - 1)]$ | $g_1 = 2\alpha$ | $g_2 = 0$ | $g_3 = 0$ |
| $d^{-(d_1+d_3+d_5)}$ | $g_0 = exp[-\alpha\tau_1(\gamma_C - \gamma_{CD}) - \alpha\tau_2(\gamma_A - \gamma_{AB}) - \alpha\tau_3(\gamma_{AB} + \gamma_{CD} - 2)]$ | $g_1 = 0$ | $g_2 = 0$ | $g_3 = 2\alpha$ |
| $d^{-(d_1+d_4+d_5)}$ | $g_0 = exp[-\alpha\tau_1(\gamma_D - \gamma_{CD}) - \alpha\tau_2(\gamma_A - \gamma_{AB}) - \alpha\tau_3(\gamma_{AB} + \gamma_{CD} - 2)]$ | $g_1 = 0$ | $g_2 = 0$ | $g_3 = 2\alpha$ |
| $d^{-(d_2+d_3+d_5)}$ | $g_0 = exp[-\alpha\tau_1(\gamma_C - \gamma_{CD}) - \alpha\tau_2(\gamma_B - \gamma_{AB}) - \alpha\tau_3(\gamma_{AB} + \gamma_{CD} - 2)]$ | $g_1 = 0$ | $g_2 = 0$ | $g_3 = 2\alpha$ |
| $d^{-(d_2+d_4+d_5)}$ | $g_0 = exp[-\alpha\tau_1(\gamma_D - \gamma_{CD}) - \alpha\tau_2(\gamma_B - \gamma_{AB}) - \alpha\tau_3(\gamma_{AB} + \gamma_{CD} - 2)]$ | $g_1 = 0$ | $g_2 = 0$ | $g_3 = 2\alpha$ |
| $d^{-(d_1+d_2+d_3+d_4)}$ | $g_0 = exp[-\alpha\tau_1(\gamma_C + \gamma_D - 2\gamma_{CD}) - \alpha\tau_2(\gamma_A + \gamma_B - 2\gamma_{AB}) - 2\alpha\tau_3(\gamma_{CD} - 1)]$ | $g_1 = 2\alpha$ | $g_2 = 2\alpha\gamma_{AB}$ | $g_3 = 0$ |
| $d^{-(d_1+d_2+d_3+d_5)}$ | $g_0 = exp[-\alpha\tau_1(\gamma_C - \gamma_{CD}) - \alpha\tau_2(\gamma_A + \gamma_B - 2\gamma_{AB}) - \alpha\tau_3(\gamma_{AB} + \gamma_{CD} - 2)]$ | $g_1 = 0$ | $g_2 = \alpha\gamma_{AB}$ | $g_3 = 2\alpha$ |
| $d^{-(d_1+d_2+d_4+d_5)}$ | $g_0 = exp[-\alpha\tau_1(\gamma_D - \gamma_{CD}) - \alpha\tau_2(\gamma_A + \gamma_B - 2\gamma_{AB}) - \alpha\tau_3(\gamma_{AB} + \gamma_{CD} - 2)]$ | $g_1 = 0$ | $g_2 = \alpha\gamma_{AB}$ | $g_3 = 2\alpha$ |
| $d^{-(d_1+d_3+d_4+d_5)}$ | $g_0 = exp[-\alpha\tau_1(\gamma_C + \gamma_D - 2\gamma_{CD}) - \alpha\tau_2(\gamma_A - \gamma_{AB}) - \alpha\tau_3(\gamma_{AB} + 2\gamma_{CD} - 3)]$ | $g_1 = \alpha$ | $g_2 = 0$ | $g_3 = 2\alpha$ |
| $d^{-(d_2+d_3+d_4+d_5)}$ | $g_0 = exp[-\alpha\tau_1(\gamma_C + \gamma_D - 2\gamma_{CD}) - \alpha\tau_2(\gamma_B - \gamma_{AB}) - \alpha\tau_3(\gamma_{AB} + 2\gamma_{CD} - 3)]$ | $g_1 = \alpha$ | $g_2 = 0$ | $g_3 = 2\alpha$ |
| $d^{-(d_1+d_2+d_3+d_4+d_5)}$ | $g_0 = exp[-\alpha\tau_1(\gamma_C + \gamma_D - 2\gamma_{CD}) - \alpha\tau_2(\gamma_A + \gamma_B - 2\gamma_{AB}) - \alpha\tau_3(\gamma_{AB} + 2\gamma_{CD} - 3)]$ | $g_1 = \alpha$ | $g_2 = \alpha\gamma_{AB}$ | $g_3 = 2\alpha$ |

| | **Histories 4 and 5** | | | |
|---|---|---|---|---|
| $d^{-(d_1+d_2)}$ | $g_0 = exp[-\alpha\tau_2(\gamma_A + \gamma_B - 2\gamma_{AB}) - 2\alpha\tau_3(\gamma_{AB} - 1)]$ | $g_1 = 0$ | $g_2 = 2\alpha$ | $g_3 = 0$ |
| $d^{-(d_3+d_4)}$ | $g_0 = exp[-\alpha\tau_1(\gamma_C + \gamma_D - 2\gamma_{CD}) - 2\alpha\tau_3(\gamma_{CD} - 1)]$ | $g_1 = 2\alpha$ | $g_2 = 0$ | $g_3 = 0$ |
| $d^{-(d_1+d_3+d_5)}$ | $g_0 = exp[-\alpha\tau_1(\gamma_C - \gamma_{CD}) - \alpha\tau_2(\gamma_A - \gamma_{AB}) - \alpha\tau_3(\gamma_{AB} + \gamma_{CD} - 2)]$ | $g_1 = 0$ | $g_2 = 0$ | $g_3 = 2\alpha$ |
| $d^{-(d_1+d_4+d_5)}$ | $g_0 = exp[-\alpha\tau_1(\gamma_D - \gamma_{CD}) - \alpha\tau_2(\gamma_A - \gamma_{AB}) - \alpha\tau_3(\gamma_{AB} + \gamma_{CD} - 2)]$ | $g_1 = 0$ | $g_2 = 0$ | $g_3 = 2\alpha$ |
| $d^{-(d_2+d_3+d_5)}$ | $g_0 = exp[-\alpha\tau_1(\gamma_C - \gamma_{CD}) - \alpha\tau_2(\gamma_B - \gamma_{AB}) - \alpha\tau_3(\gamma_{AB} + \gamma_{CD} - 2)]$ | $g_1 = 0$ | $g_2 = 0$ | $g_3 = 2\alpha$ |
| $d^{-(d_2+d_4+d_5)}$ | $g_0 = exp[-\alpha\tau_1(\gamma_D - \gamma_{CD}) - \alpha\tau_2(\gamma_B - \gamma_{AB}) - \alpha\tau_3(\gamma_{AB} + \gamma_{CD} - 2)]$ | $g_1 = 0$ | $g_2 = 0$ | $g_3 = 2\alpha$ |
| $d^{-(d_1+d_2+d_3+d_4)}$ | $g_0 = exp[-\alpha\tau_1(\gamma_C + \gamma_D - 2\gamma_{CD}) - \alpha\tau_2(\gamma_A + \gamma_B - 2\gamma_{AB}) - 2\alpha\tau_3(\gamma_{AB} + \gamma_{CD} - 2)]$ | $g_1 = 2\alpha$ | $g_2 = 2\alpha$ | $g_3 = 0$ |
| $d^{-(d_1+d_2+d_3+d_5)}$ | $g_0 = exp[-\alpha\tau_1(\gamma_C - \gamma_{CD}) - \alpha\tau_2(\gamma_A + \gamma_B - 2\gamma_{AB}) - \alpha\tau_3(2\gamma_{AB} + \gamma_{CD} - 3)]$ | $g_1 = 0$ | $g_2 = \alpha$ | $g_3 = 2\alpha$ |
| $d^{-(d_1+d_2+d_4+d_5)}$ | $g_0 = exp[-\alpha\tau_1(\gamma_D - \gamma_{CD}) - \alpha\tau_2(\gamma_A + \gamma_B - 2\gamma_{AB}) - \alpha\tau_3(2\gamma_{AB} + \gamma_{CD} - 3)]$ | $g_1 = 0$ | $g_2 = \alpha$ | $g_3 = 2\alpha$ |
| $d^{-(d_1+d_3+d_4+d_5)}$ | $g_0 = exp[-\alpha\tau_1(\gamma_C + \gamma_D - 2\gamma_{CD}) - \alpha\tau_2(\gamma_A - \gamma_{AB}) - \alpha\tau_3(\gamma_{AB} + 2\gamma_{CD} - 3)]$ | $g_1 = \alpha$ | $g_2 = 0$ | $g_3 = 2\alpha$ |
| $d^{-(d_2+d_3+d_4+d_5)}$ | $g_0 = exp[-\alpha\tau_1(\gamma_C + \gamma_D - 2\gamma_{CD}) - \alpha\tau_2(\gamma_B - \gamma_{AB}) - \alpha\tau_3(\gamma_{AB} + 2\gamma_{CD} - 3)]$ | $g_1 = \alpha$ | $g_2 = 0$ | $g_3 = 2\alpha$ |
| $d^{-(d_1+d_2+d_3+d_4+d_5)}$ | $g_0 = exp[-\alpha\tau_1(\gamma_C + \gamma_D - 2\gamma_{CD}) - \alpha\tau_2(\gamma_A + \gamma_B - 2\gamma_{AB}) - 2\alpha\tau_3(\gamma_{AB} + \gamma_{CD} - 2)]$ | $g_1 = \alpha$ | $g_2 = \alpha$ | $g_3 = 2\alpha$ |

| | **Histories 6 and 7** | | | |
|---|---|---|---|---|
| $d^{-(d_1+d_2)}$ | $g_0 = exp[-\alpha\tau_1(\gamma_C - \gamma_{CD}) - \alpha\tau_2(\gamma_A - \gamma_{AB}) - \alpha\tau_3(\gamma_{AB} + \gamma_{CD} - 2)]$ | $g_1 = 0$ | $g_2 = 2\alpha$ | $g_3 = 0$ |
| $d^{-(d_3+d_4)}$ | $g_0 = exp[-\alpha\tau_1(\gamma_D - 2\gamma_{CD}) - \alpha\tau_2(\gamma_B - \gamma_{AB}) - \alpha\tau_3(\gamma_{AB} + \gamma_{CD} - 2)]$ | $g_1 = 2\alpha$ | $g_2 = 0$ | $g_3 = 0$ |
| $d^{-(d_1+d_3+d_5)}$ | $g_0 = exp[-\alpha\tau_2(\gamma_A + \gamma_B - 2\gamma_{AB}) - 2\alpha\tau_3(\gamma_{AB} - 1)]$ | $g_1 = 0$ | $g_2 = 0$ | $g_3 = 2\alpha$ |
| $d^{-(d_1+d_4+d_5)}$ | $g_0 = exp[-\alpha\tau_1(\gamma_D - \gamma_{CD}) - \alpha\tau_2(\gamma_A - \gamma_{AB}) - \alpha\tau_3(\gamma_{AB} + \gamma_{CD} - 2)]$ | $g_1 = 0$ | $g_2 = 0$ | $g_3 = 2\alpha$ |
| $d^{-(d_2+d_3+d_5)}$ | $g_0 = exp[-\alpha\tau_1(\gamma_C - \gamma_{CD}) - \alpha\tau_2(\gamma_B - \gamma_{AB}) - \alpha\tau_3(\gamma_{AB} + \gamma_{CD} - 2)]$ | $g_1 = 0$ | $g_2 = 0$ | $g_3 = 2\alpha$ |
| $d^{-(d_2+d_4+d_5)}$ | $g_0 = exp[-\alpha\tau_1(\gamma_C + \gamma_D - 2\gamma_{CD}) - 2\alpha\tau_3(\gamma_{CD} - 1)]$ | $g_1 = 0$ | $g_2 = 0$ | $g_3 = 2\alpha$ |
| $d^{-(d_1+d_2+d_3+d_4)}$ | $g_0 = exp[-\alpha\tau_1(\gamma_C + \gamma_D - 2\gamma_{CD}) - \alpha\tau_2(\gamma_A + \gamma_B - 2\gamma_{AB}) - 2\alpha\tau_3(\gamma_{AB} + \gamma_{CD} - 2)]$ | $g_1 = 2\alpha$ | $g_2 = 2\alpha$ | $g_3 = 0$ |
| $d^{-(d_1+d_2+d_3+d_5)}$ | $g_0 = exp[-\alpha\tau_1(\gamma_C - \gamma_{CD}) - \alpha\tau_2(\gamma_A + \gamma_B - 2\gamma_{AB}) - \alpha\tau_3(2\gamma_{AB} + \gamma_{CD} - 3)]$ | $g_1 = 0$ | $g_2 = \alpha$ | $g_3 = 2\alpha$ |
| $d^{-(d_1+d_2+d_4+d_5)}$ | $g_0 = exp[-\alpha\tau_1(\gamma_C + \gamma_D - \gamma_{CD}) - \alpha\tau_2(\gamma_A - \gamma_{AB}) - \alpha\tau_3(\gamma_{AB} + 2\gamma_{CD} - 3)]$ | $g_1 = 0$ | $g_2 = \alpha$ | $g_3 = 2\alpha$ |
| $d^{-(d_1+d_3+d_4+d_5)}$ | $g_0 = exp[-\alpha\tau_1(\gamma_D - \gamma_{CD}) - \alpha\tau_2(\gamma_A + \gamma_B - 2\gamma_{AB}) - \alpha\tau_3(2\gamma_{AB} + \gamma_{CD} - 3)]$ | $g_1 = \alpha$ | $g_2 = 0$ | $g_3 = 2\alpha$ |
| $d^{-(d_2+d_3+d_4+d_5)}$ | $g_0 = exp[-\alpha\tau_1(\gamma_C + \gamma_D - 2\gamma_{CD}) - \alpha\tau_2(\gamma_B - \gamma_{AB}) - \alpha\tau_3(\gamma_{AB} + 2\gamma_{CD} - 3)]$ | $g_1 = \alpha$ | $g_2 = 0$ | $g_3 = 2\alpha$ |
| $d^{-(d_1+d_2+d_3+d_4+d_5)}$ | $g_0 = exp[-\alpha\tau_1(\gamma_C + \gamma_D - 2\gamma_{CD}) - \alpha\tau_2(\gamma_A + \gamma_B - 2\gamma_{AB}) - 2\alpha\tau_3(\gamma_{AB} + \gamma_{CD} - 2)]$ | $g_1 = \alpha$ | $g_2 = \alpha$ | $g_3 = 2\alpha$ |

| | **Histories 8 and 9** | | | |
|---|---|---|---|---|
| $d^{-(d_1+d_2)}$ | $g_0 = exp[-\alpha\tau_1(\gamma_D - \gamma_{CD}) - \alpha\tau_2(\gamma_A - \gamma_{AB}) - \alpha\tau_3(\gamma_{AB} + \gamma_{CD} - 2)]$ | $g_1 = 0$ | $g_2 = 2\alpha$ | $g_3 = 0$ |
| $d^{-(d_3+d_4)}$ | $g_0 = exp[-\alpha\tau_1(\gamma_C - \gamma_{CD}) - \alpha\tau_2(\gamma_B - \gamma_{AB}) - \alpha\tau_3(\gamma_{AB} + \gamma_{CD} - 2)]$ | $g_1 = 2\alpha$ | $g_2 = 0$ | $g_3 = 0$ |
| $d^{-(d_1+d_3+d_5)}$ | $g_0 = exp[-\alpha\tau_2(\gamma_A + \gamma_B - 2\gamma_{AB}) - 2\alpha\tau_3(\gamma_{AB} - 1)]$ | $g_1 = 0$ | $g_2 = 0$ | $g_3 = 2\alpha$ |
| $d^{-(d_1+d_4+d_5)}$ | $g_0 = exp[-\alpha\tau_1(\gamma_C - \gamma_{CD}) - \alpha\tau_2(\gamma_A - \gamma_{AB}) - \alpha\tau_3(\gamma_{AB} + \gamma_{CD} - 2)]$ | $g_1 = 0$ | $g_2 = 0$ | $g_3 = 2\alpha$ |
| $d^{-(d_2+d_3+d_5)}$ | $g_0 = exp[-\alpha\tau_1(\gamma_D - \gamma_{CD}) - \alpha\tau_2(\gamma_B - \gamma_{AB}) - \alpha\tau_3(\gamma_{AB} + \gamma_{CD} - 2)]$ | $g_1 = 0$ | $g_2 = 0$ | $g_3 = 2\alpha$ |
| $d^{-(d_2+d_4+d_5)}$ | $g_0 = exp[-\alpha\tau_1(\gamma_C + \gamma_D - 2\gamma_{CD}) - 2\alpha\tau_3(\gamma_{CD} - 1)]$ | $g_1 = 0$ | $g_2 = 0$ | $g_3 = 2\alpha$ |
| $d^{-(d_1+d_2+d_3+d_4)}$ | $g_0 = exp[-\alpha\tau_1(\gamma_C + \gamma_D - 2\gamma_{CD}) - \alpha\tau_2(\gamma_A + \gamma_B - 2\gamma_{AB}) - 2\alpha\tau_3(\gamma_{AB} + \gamma_{CD} - 2)]$ | $g_1 = 2\alpha$ | $g_2 = 2\alpha$ | $g_3 = 0$ |
| $d^{-(d_1+d_2+d_3+d_5)}$ | $g_0 = exp[-\alpha\tau_1(\gamma_D - \gamma_{CD}) - \alpha\tau_2(\gamma_A + \gamma_B - 2\gamma_{AB}) - \alpha\tau_3(2\gamma_{AB} + \gamma_{CD} - 3)]$ | $g_1 = 0$ | $g_2 = \alpha$ | $g_3 = 2\alpha$ |
| $d^{-(d_1+d_2+d_4+d_5)}$ | $g_0 = exp[-\alpha\tau_1(\gamma_C + \gamma_D - \gamma_{CD}) - \alpha\tau_2(\gamma_A - \gamma_{AB}) - \alpha\tau_3(\gamma_{AB} + 2\gamma_{CD} - 3)]$ | $g_1 = 0$ | $g_2 = \alpha$ | $g_3 = 2\alpha$ |
| $d^{-(d_1+d_3+d_4+d_5)}$ | $g_0 = exp[-\alpha\tau_1(\gamma_C - \gamma_{CD}) - \alpha\tau_2(\gamma_A + \gamma_B - 2\gamma_{AB}) - \alpha\tau_3(2\gamma_{AB} + \gamma_{CD} - 3)]$ | $g_1 = \alpha$ | $g_2 = 0$ | $g_3 = 2\alpha$ |
| $d^{-(d_2+d_3+d_4+d_5)}$ | $g_0 = exp[-\alpha\tau_1(\gamma_C + \gamma_D - 2\gamma_{CD}) - \alpha\tau_2(\gamma_B - \gamma_{AB}) - \alpha\tau_3(\gamma_{AB} + 2\gamma_{CD} - 3)]$ | $g_1 = \alpha$ | $g_2 = 0$ | $g_3 = 2\alpha$ |
| $d^{-(d_1+d_2+d_3+d_4+d_5)}$ | $g_0 = exp[-\alpha\tau_1(\gamma_C + \gamma_D - 2\gamma_{CD}) - \alpha\tau_2(\gamma_A + \gamma_B - 2\gamma_{AB}) - 2\alpha\tau_3(\gamma_{AB} + \gamma_{CD} - 2)]$ | $g_1 = \alpha$ | $g_2 = \alpha$ | $g_3 = 2\alpha$ |

| | **History 10** | | | |
|---|---|---|---|---|
| $d^{-(d_1+d_2)}$ | $g_0 = exp[-\alpha\tau_2(\gamma_A + \gamma_B - 2\gamma_{AB}) - 2\alpha\tau_3(\gamma_{AB} - 1)]$ | $g_1 = 0$ | $g_2 = 0$ | $g_3 = 2\alpha$ |
| $d^{-(d_3+d_4)}$ | $g_0 = exp[-\alpha\tau_1(\gamma_C + \gamma_D - 2\gamma_{CD})]$ | $g_1 = 2\alpha\gamma_{CD}$ | $g_2 = 0$ | $g_3 = 0$ |
| $d^{-(d_1+d_3+d_5)}$ | $g_0 = exp[-\alpha\tau_1(\gamma_C - \gamma_{CD}) - \alpha\tau_2(\gamma_A - \gamma_{AB}) - \alpha\tau_3(\gamma_{AB} + \gamma_{CD} - 2)]$ | $g_1 = 0$ | $g_2 = 0$ | $g_3 = 2\alpha$ |
| $d^{-(d_1+d_4+d_5)}$ | $g_0 = exp[-\alpha\tau_1(\gamma_D - \gamma_{CD}) - \alpha\tau_2(\gamma_A - \gamma_{AB}) - \alpha\tau_3(\gamma_{AB} + \gamma_{CD} - 2)]$ | $g_1 = 0$ | $g_2 = 0$ | $g_3 = 2\alpha$ |
| $d^{-(d_2+d_3+d_5)}$ | $g_0 = exp[-\alpha\tau_1(\gamma_C - \gamma_{CD}) - \alpha\tau_2(\gamma_B - \gamma_{AB}) - \alpha\tau_3(\gamma_{AB} + \gamma_{CD} - 2)]$ | $g_1 = 0$ | $g_2 = 2\alpha$ | $g_3 = 0$ |
| $d^{-(d_2+d_4+d_5)}$ | $g_0 = exp[-\alpha\tau_1(\gamma_D - \gamma_{CD}) - \alpha\tau_2(\gamma_B - \gamma_{AB}) - \alpha\tau_3(\gamma_{AB} + \gamma_{CD} - 2)]$ | $g_1 = 0$ | $g_2 = 2\alpha$ | $g_3 = 0$ |
| $d^{-(d_1+d_2+d_3+d_4)}$ | $g_0 = exp[-\alpha\tau_1(\gamma_C + \gamma_D - 2\gamma_{CD}) - \alpha\tau_2(\gamma_A + \gamma_B - 2\gamma_{AB}) - 2\alpha\tau_3(\gamma_{AB} - 1)]$ | $g_1 = 2\alpha\gamma_{CD}$ | $g_2 = 0$ | $g_3 = 2\alpha$ |
| $d^{-(d_1+d_2+d_3+d_5)}$ | $g_0 = exp[-\alpha\tau_1(\gamma_C - \gamma_{CD}) - \alpha\tau_2(\gamma_A + \gamma_B - 2\gamma_{AB}) - \alpha\tau_3(2\gamma_{AB} + \gamma_{CD} - 3)]$ | $g_1 = 0$ | $g_2 = \alpha$ | $g_3 = 2\alpha$ |
| $d^{-(d_1+d_2+d_4+d_5)}$ | $g_0 = exp[-\alpha\tau_1(\gamma_D - \gamma_{CD}) - \alpha\tau_2(\gamma_A + \gamma_B - 2\gamma_{AB}) - \alpha\tau_3(2\gamma_{AB} + \gamma_{CD} - 3)]$ | $g_1 = 0$ | $g_2 = \alpha$ | $g_3 = 2\alpha$ |
| $d^{-(d_1+d_3+d_4+d_5)}$ | $g_0 = exp[-\alpha\tau_1(\gamma_C + \gamma_D - 2\gamma_{CD}) - \alpha\tau_2(\gamma_A - \gamma_{AB}) - \alpha\tau_3(\gamma_{AB} + \gamma_{CD} - 2)]$ | $g_1 = \alpha\gamma_{CD}$ | $g_2 = 0$ | $g_3 = 2\alpha$ |
| $d^{-(d_2+d_3+d_4+d_5)}$ | $g_0 = exp[-\alpha\tau_1(\gamma_C + \gamma_D - 2\gamma_{CD}) - \alpha\tau_2(\gamma_B - \gamma_{AB}) - \alpha\tau_3(\gamma_{AB} + \gamma_{CD} - 2)]$ | $g_1 = \alpha\gamma_{CD}$ | $g_2 = 2\alpha$ | $g_3 = 0$ |
| $d^{-(d_1+d_2+d_3+d_4+d_5)}$ | $g_0 = exp[-\alpha\tau_1(\gamma_C + \gamma_D - 2\gamma_{CD}) - \alpha\tau_2(\gamma_A + \gamma_B - 2\gamma_{AB}) - \alpha\tau_3(2\gamma_{AB} + \gamma_{CD} - 3)]$ | $g_1 = \alpha\gamma_{CD}$ | $g_2 = \alpha$ | $g_3 = 2\alpha$ |

| | **History 11** | | | |
|---|---|---|---|---|
| $d^{-(d_1+d_2)}$ | $g_0 = exp[-\alpha\tau_2(\gamma_A + \gamma_B - 2\gamma_{AB}) - 2\alpha\tau_3(\gamma_{AB} - 1)]$ | $g_1 = 0$ | $g_2 = 0$ | $g_3 = 2\alpha$ |
| $d^{-(d_3+d_4)}$ | $g_0 = exp[-\alpha\tau_1(\gamma_C + \gamma_D - 2\gamma_{CD}) - 2\alpha\tau_3(\gamma_{CD} - 1)]$ | $g_1 = 2\alpha$ | $g_2 = 0$ | $g_3 = 0$ |
| $d^{-(d_1+d_3+d_5)}$ | $g_0 = exp[-\alpha\tau_1(\gamma_C - \gamma_{CD}) - \alpha\tau_2(\gamma_A - \gamma_{AB}) - \alpha\tau_3(\gamma_{AB} + \gamma_{CD} - 2)]$ | $g_1 = 0$ | $g_2 = 0$ | $g_3 = 2\alpha$ |
| $d^{-(d_1+d_4+d_5)}$ | $g_0 = exp[-\alpha\tau_1(\gamma_D - \gamma_{CD}) - \alpha\tau_2(\gamma_A - \gamma_{AB}) - \alpha\tau_3(\gamma_{AB} + \gamma_{CD} - 2)]$ | $g_1 = 0$ | $g_2 = 0$ | $g_3 = 2\alpha$ |
| $d^{-(d_2+d_3+d_5)}$ | $g_0 = exp[-\alpha\tau_1(\gamma_C - \gamma_{CD}) - \alpha\tau_2(\gamma_B - \gamma_{AB}) - \alpha\tau_3(\gamma_{AB} + \gamma_{CD} - 2)]$ | $g_1 = 0$ | $g_2 = 2\alpha$ | $g_3 = 0$ |
| $d^{-(d_2+d_4+d_5)}$ | $g_0 = exp[-\alpha\tau_1(\gamma_D - \gamma_{CD}) - \alpha\tau_2(\gamma_B - \gamma_{AB}) - \alpha\tau_3(\gamma_{AB} + \gamma_{CD} - 2)]$ | $g_1 = 0$ | $g_2 = 2\alpha$ | $g_3 = 0$ |
| $d^{-(d_1+d_2+d_3+d_4)}$ | $g_0 = exp[-\alpha\tau_1(\gamma_C + \gamma_D - 2\gamma_{CD}) - \alpha\tau_2(\gamma_A + \gamma_B - 2\gamma_{AB}) - 2\alpha\tau_3(\gamma_{AB} + \gamma_{CD} - 2)]$ | $g_1 = 2\alpha$ | $g_2 = 0$ | $g_3 = 2\alpha$ |
| $d^{-(d_1+d_2+d_3+d_5)}$ | $g_0 = exp[-\alpha\tau_1(\gamma_C - \gamma_{CD}) - \alpha\tau_2(\gamma_A + \gamma_B - 2\gamma_{AB}) - \alpha\tau_3(2\gamma_{AB} + \gamma_{CD} - 3)]$ | $g_1 = 0$ | $g_2 = \alpha$ | $g_3 = 2\alpha$ |
| $d^{-(d_1+d_2+d_4+d_5)}$ | $g_0 = exp[-\alpha\tau_1(\gamma_D - \gamma_{CD}) - \alpha\tau_2(\gamma_A + \gamma_B - 2\gamma_{AB}) - \alpha\tau_3(2\gamma_{AB} + \gamma_{CD} - 3)]$ | $g_1 = 0$ | $g_2 = \alpha$ | $g_3 = 2\alpha$ |
| $d^{-(d_1+d_3+d_4+d_5)}$ | $g_0 = exp[-\alpha\tau_1(\gamma_C + \gamma_D - 2\gamma_{CD}) - \alpha\tau_2(\gamma_A - \gamma_{AB}) - \alpha\tau_3(\gamma_{AB} + 2\gamma_{CD} - 3)]$ | $g_1 = \alpha$ | $g_2 = 0$ | $g_3 = 2\alpha$ |
| $d^{-(d_2+d_3+d_4+d_5)}$ | $g_0 = exp[-\alpha\tau_1(\gamma_C + \gamma_D - 2\gamma_{CD}) - \alpha\tau_2(\gamma_B - \gamma_{AB}) - \alpha\tau_3(\gamma_{AB} + 2\gamma_{CD} - 3)]$ | $g_1 = \alpha$ | $g_2 = 2\alpha$ | $g_3 = 0$ |
| $d^{-(d_1+d_2+d_3+d_4+d_5)}$ | $g_0 = exp[-\alpha\tau_1(\gamma_C + \gamma_D - 2\gamma_{CD}) - \alpha\tau_2(\gamma_A + \gamma_B - 2\gamma_{AB}) - 2\alpha\tau_3(\gamma_{AB} + \gamma_{CD} - 2)]$ | $g_1 = \alpha$ | $g_2 = \alpha$ | $g_3 = 2\alpha$ |

| | **History 12** | | | |
|---|---|---|---|---|
| $d^{-(d_1+d_2)}$ | $g_0 = exp[-\alpha\tau_1(\gamma_C - \gamma_{CD}) - \alpha\tau_2(\gamma_A - \gamma_{AB}) - \alpha\tau_3(\gamma_{AB} + \gamma_{CD} - 2)]$ | $g_1 = 0$ | $g_2 = 0$ | $g_3 = 2\alpha$ |
| $d^{-(d_3+d_4)}$ | $g_0 = exp[-\alpha\tau_1(\gamma_D - \gamma_{CD}) - \alpha\tau_2(\gamma_B - \gamma_{AB}) - \alpha\tau_3(\gamma_{AB} + \gamma_{CD} - 2)]$ | $g_1 = 2\alpha$ | $g_2 = 0$ | $g_3 = 0$ |
| $d^{-(d_1+d_3+d_5)}$ | $g_0 = exp[-\alpha\tau_1(\gamma_A + \gamma_B - 2\gamma_{AB}) - 2\alpha\tau_3(\gamma_{AB} - 1)]$ | $g_1 = 0$ | $g_2 = 0$ | $g_3 = 2\alpha$ |
| $d^{-(d_1+d_4+d_5)}$ | $g_0 = exp[-\alpha\tau_1(\gamma_D - \gamma_{CD}) - \alpha\tau_2(\gamma_A - \gamma_{AB}) - \alpha\tau_3(\gamma_{AB} + \gamma_{CD} - 2)]$ | $g_1 = 0$ | $g_2 = 0$ | $g_3 = 2\alpha$ |
| $d^{-(d_2+d_3+d_5)}$ | $g_0 = exp[-\alpha\tau_1(\gamma_C - \gamma_{CD}) - \alpha\tau_2(\gamma_B - \gamma_{AB}) - \alpha\tau_3(\gamma_{AB} + \gamma_{CD} - 2)]$ | $g_1 = 0$ | $g_2 = 2\alpha$ | $g_3 = 0$ |
| $d^{-(d_2+d_4+d_5)}$ | $g_0 = exp[-\alpha\tau_1(\gamma_C + \gamma_D - 2\gamma_{CD}) - 2\alpha\tau_3(\gamma_{CD} - 1)]$ | $g_1 = 0$ | $g_2 = 2\alpha$ | $g_3 = 0$ |
| $d^{-(d_1+d_2+d_3+d_4)}$ | $g_0 = exp[-\alpha\tau_1(\gamma_C + \gamma_D - 2\gamma_{CD}) - \alpha\tau_2(\gamma_A + \gamma_B - 2\gamma_{AB}) - 2\alpha\tau_3(\gamma_{AB} + \gamma_{CD} - 2)]$ | $g_1 = 2\alpha$ | $g_2 = 0$ | $g_3 = 2\alpha$ |
| $d^{-(d_1+d_2+d_3+d_5)}$ | $g_0 = exp[-\alpha\tau_1(\gamma_C - \gamma_{CD}) - \alpha\tau_2(\gamma_A + \gamma_B - 2\gamma_{AB}) - \alpha\tau_3(2\gamma_{AB} + \gamma_{CD} - 3)]$ | $g_1 = 0$ | $g_2 = \alpha$ | $g_3 = 2\alpha$ |
| $d^{-(d_1+d_2+d_4+d_5)}$ | $g_0 = exp[-\alpha\tau_1(\gamma_C + \gamma_D - 2\gamma_{CD}) - \alpha\tau_2(\gamma_A - \gamma_{AB}) - \alpha\tau_3(\gamma_{AB} + 2\gamma_{CD} - 3)]$ | $g_1 = 0$ | $g_2 = \alpha$ | $g_3 = 2\alpha$ |
| $d^{-(d_1+d_3+d_4+d_5)}$ | $g_0 = exp[-\alpha\tau_1(\gamma_D - \gamma_{CD}) - \alpha\tau_2(\gamma_A + \gamma_B - 2\gamma_{AB}) - \alpha\tau_3(2\gamma_{AB} + \gamma_{CD} - 3)]$ | $g_1 = \alpha$ | $g_2 = 0$ | $g_3 = 2\alpha$ |
| $d^{-(d_2+d_3+d_4+d_5)}$ | $g_0 = exp[-\alpha\tau_1(\gamma_C + \gamma_D - 2\gamma_{CD}) - \alpha\tau_2(\gamma_B - \gamma_{AB}) - \alpha\tau_3(\gamma_{AB} + 2\gamma_{CD} - 3)]$ | $g_1 = \alpha$ | $g_2 = 2\alpha$ | $g_3 = 0$ |
| $d^{-(d_1+d_2+d_3+d_4+d_5)}$ | $g_0 = exp[-\alpha\tau_1(\gamma_C + \gamma_D - 2\gamma_{CD}) - \alpha\tau_2(\gamma_A + \gamma_B - 2\gamma_{AB}) - 2\alpha\tau_3(\gamma_{AB} + \gamma_{CD} - 2)]$ | $g_1 = \alpha$ | $g_2 = \alpha$ | $g_3 = 2\alpha$ |

| | **History 13** | | | |
|---|---|---|---|---|
| $d^{-(d_1+d_2)}$ | $g_0 = exp[-\alpha\tau_1(\gamma_D - \gamma_{CD}) - \alpha\tau_2(\gamma_A - \gamma_{AB}) - \alpha\tau_3(\gamma_{AB} + \gamma_{CD} - 2)]$ | $g_1 = 0$ | $g_2 = 0$ | $g_3 = 2\alpha$ |
| $d^{-(d_3+d_4)}$ | $g_0 = exp[-\alpha\tau_1(\gamma_C - \gamma_{CD}) - \alpha\tau_2(\gamma_B - \gamma_{AB}) - \alpha\tau_3(\gamma_{AB} + \gamma_{CD} - 2)]$ | $g_1 = 2\alpha$ | $g_2 = 0$ | $g_3 = 0$ |
| $d^{-(d_1+d_3+d_5)}$ | $g_0 = exp[-\alpha\tau_1(\gamma_A + \gamma_B - 2\gamma_{AB}) - 2\alpha\tau_3(\gamma_{AB} - 1)]$ | $g_1 = 0$ | $g_2 = 0$ | $g_3 = 2\alpha$ |
| $d^{-(d_1+d_4+d_5)}$ | $g_0 = exp[-\alpha\tau_1(\gamma_C - \gamma_{CD}) - \alpha\tau_2(\gamma_A - \gamma_{AB}) - \alpha\tau_3(\gamma_{AB} + \gamma_{CD} - 2)]$ | $g_1 = 0$ | $g_2 = 0$ | $g_3 = 2\alpha$ |
| $d^{-(d_2+d_3+d_5)}$ | $g_0 = exp[-\alpha\tau_1(\gamma_D - \gamma_{CD}) - \alpha\tau_2(\gamma_B - \gamma_{AB}) - \alpha\tau_3(\gamma_{AB} + \gamma_{CD} - 2)]$ | $g_1 = 0$ | $g_2 = 2\alpha$ | $g_3 = 0$ |
| $d^{-(d_2+d_4+d_5)}$ | $g_0 = exp[-\alpha\tau_1(\gamma_C + \gamma_D - 2\gamma_{CD}) - 2\alpha\tau_3(\gamma_{CD} - 1)]$ | $g_1 = 0$ | $g_2 = 2\alpha$ | $g_3 = 0$ |
| $d^{-(d_1+d_2+d_3+d_4)}$ | $g_0 = exp[-\alpha\tau_1(\gamma_C + \gamma_D - 2\gamma_{CD}) - \alpha\tau_2(\gamma_A + \gamma_B - 2\gamma_{AB}) - 2\alpha\tau_3(\gamma_{AB} + \gamma_{CD} - 2)]$ | $g_1 = 2\alpha$ | $g_2 = 0$ | $g_3 = 2\alpha$ |
| $d^{-(d_1+d_2+d_3+d_5)}$ | $g_0 = exp[-\alpha\tau_1(\gamma_D - \gamma_{CD}) - \alpha\tau_2(\gamma_A + \gamma_B - 2\gamma_{AB}) - \alpha\tau_3(2\gamma_{AB} + \gamma_{CD} - 3)]$ | $g_1 = 0$ | $g_2 = \alpha$ | $g_3 = 2\alpha$ |
| $d^{-(d_1+d_2+d_4+d_5)}$ | $g_0 = exp[-\alpha\tau_1(\gamma_C + \gamma_D - 2\gamma_{CD}) - \alpha\tau_2(\gamma_A - \gamma_{AB}) - \alpha\tau_3(\gamma_{AB} + 2\gamma_{CD} - 3)]$ | $g_1 = 0$ | $g_2 = \alpha$ | $g_3 = 2\alpha$ |
| $d^{-(d_1+d_3+d_4+d_5)}$ | $g_0 = exp[-\alpha\tau_1(\gamma_C - \gamma_{CD}) - \alpha\tau_2(\gamma_A + \gamma_B - 2\gamma_{AB}) - \alpha\tau_3(2\gamma_{AB} + \gamma_{CD} - 3)]$ | $g_1 = \alpha$ | $g_2 = 0$ | $g_3 = 2\alpha$ |
| $d^{-(d_2+d_3+d_4+d_5)}$ | $g_0 = exp[-\alpha\tau_1(\gamma_C + \gamma_D - 2\gamma_{CD}) - \alpha\tau_2(\gamma_B - \gamma_{AB}) - \alpha\tau_3(\gamma_{AB} + 2\gamma_{CD} - 3)]$ | $g_1 = \alpha$ | $g_2 = 2\alpha$ | $g_3 = 0$ |
| $d^{-(d_1+d_2+d_3+d_4+d_5)}$ | $g_0 = exp[-\alpha\tau_1(\gamma_C + \gamma_D - 2\gamma_{CD}) - \alpha\tau_2(\gamma_A + \gamma_B - 2\gamma_{AB}) - 2\alpha\tau_3(\gamma_{AB} + \gamma_{CD} - 2)]$ | $g_1 = \alpha$ | $g_2 = \alpha$ | $g_3 = 2\alpha$ |

| | **History 14** | | | |
|---|---|---|---|---|
| $d^{-(d_1+d_2)}$ | $g_0 = exp[\alpha\tau_2(\gamma_A + \gamma_B - 2\gamma_{AB}) - 2\alpha\tau_3(\gamma_{AB} - 1)]$ | $g_1 = 0$ | $g_2 = 0$ | $g_3 = 2\alpha$ |
| $d^{-(d_3+d_4)}$ | $g_0 = exp[-\alpha\tau_1(\gamma_C + \gamma_D - 2\gamma_{CD})]$ | $g_1 = 2\alpha\gamma_{CD}$ | $g_2 = 0$ | $g_3 = 0$ |
| $d^{-(d_1+d_3+d_5)}$ | $g_0 = exp[-\alpha\tau_1(\gamma_C - \gamma_{CD}) - \alpha\tau_2(\gamma_B - \gamma_{AB}) - \alpha\tau_3(\gamma_{AB} + \gamma_{CD} - 2)]$ | $g_1 = 0$ | $g_2 = 0$ | $g_3 = 2\alpha$ |
| $d^{-(d_1+d_4+d_5)}$ | $g_0 = exp[-\alpha\tau_1(\gamma_D - \gamma_{CD}) - \alpha\tau_2(\gamma_B - \gamma_{AB}) - \alpha\tau_3(\gamma_{AB} + \gamma_{CD} - 2)]$ | $g_1 = 0$ | $g_2 = 0$ | $g_3 = 2\alpha$ |
| $d^{-(d_2+d_3+d_5)}$ | $g_0 = exp[-\alpha\tau_1(\gamma_C - \gamma_{CD}) - \alpha\tau_2(\gamma_A - \gamma_{AB}) - \alpha\tau_3(\gamma_{AB} + \gamma_{CD} - 2)]$ | $g_1 = 0$ | $g_2 = 2\alpha$ | $g_3 = 0$ |
| $d^{-(d_2+d_4+d_5)}$ | $g_0 = exp[-\alpha\tau_1(\gamma_D - \gamma_{CD}) - \alpha\tau_2(\gamma_A - \gamma_{AB}) - \alpha\tau_3(\gamma_{AB} + \gamma_{CD} - 2)]$ | $g_1 = 0$ | $g_2 = 2\alpha$ | $g_3 = 0$ |
| $d^{-(d_1+d_2+d_3+d_4)}$ | $g_0 = exp[-\alpha\tau_1(\gamma_C + \gamma_D - 2\gamma_{CD}) - \alpha\tau_2(\gamma_A + \gamma_B - 2\gamma_{AB}) - 2\alpha\tau_3(\gamma_{AB} - 1)]$ | $g_1 = 2\alpha\gamma_{CD}$ | $g_2 = 0$ | $g_3 = 2\alpha$ |
| $d^{-(d_1+d_2+d_3+d_5)}$ | $g_0 = exp[-\alpha\tau_1(\gamma_C - \gamma_{CD}) - \alpha\tau_2(\gamma_A + \gamma_B - 2\gamma_{AB}) - \alpha\tau_3(2\gamma_{AB} + \gamma_{CD} - 3)]$ | $g_1 = 0$ | $g_2 = \alpha$ | $g_3 = 2\alpha$ |
| $d^{-(d_1+d_2+d_4+d_5)}$ | $g_0 = exp[-\alpha\tau_1(\gamma_D - \gamma_{CD}) - \alpha\tau_2(\gamma_A + \gamma_B - 2\gamma_{AB}) - \alpha\tau_3(2\gamma_{AB} + \gamma_{CD} - 3)]$ | $g_1 = 0$ | $g_2 = \alpha$ | $g_3 = 2\alpha$ |
| $d^{-(d_1+d_3+d_4+d_5)}$ | $g_0 = exp[-\alpha\tau_1(\gamma_C + \gamma_D - 2\gamma_{CD}) - \alpha\tau_2(\gamma_B - \gamma_{AB}) - \alpha\tau_3(\gamma_{AB} + \gamma_{CD} - 2)]$ | $g_1 = \alpha\gamma_{CD}$ | $g_2 = 0$ | $g_3 = 2\alpha$ |
| $d^{-(d_2+d_3+d_4+d_5)}$ | $g_0 = exp[-\alpha\tau_1(\gamma_C + \gamma_D - 2\gamma_{CD}) - \alpha\tau_2(\gamma_A - \gamma_{AB}) - \alpha\tau_3(\gamma_{AB} + \gamma_{CD} - 2)]$ | $g_1 = \alpha\gamma_{CD}$ | $g_2 = 2\alpha$ | $g_3 = 0$ |
| $d^{-(d_1+d_2+d_3+d_4+d_5)}$ | $g_0 = exp[-\alpha\tau_1(\gamma_C + \gamma_D - 2\gamma_{CD}) - \alpha\tau_2(\gamma_A + \gamma_B - 2\gamma_{AB}) - \alpha\tau_3(2\gamma_{AB} + \gamma_{CD} - 3)]$ | $g_1 = \alpha\gamma_{CD}$ | $g_2 = \alpha$ | $g_3 = 2\alpha$ |

| | **History 15** | | | |
|---|---|---|---|---|
| $d^{-(d_1+d_2)}$ | $g_0 = exp[\alpha\tau_2(\gamma_A + \gamma_B - 2\gamma_{AB}) - 2\alpha\tau_3(\gamma_{AB} - 1)]$ | $g_1 = 0$ | $g_2 = 0$ | $g_3 = 2\alpha$ |
| $d^{-(d_3+d_4)}$ | $g_0 = exp[-\alpha\tau_1(\gamma_C + \gamma_D - 2\gamma_{CD}) - 2\alpha\tau_3(\gamma_{CD} - 1)]$ | $g_1 = 2\alpha$ | $g_2 = 0$ | $g_3 = 0$ |
| $d^{-(d_1+d_3+d_5)}$ | $g_0 = exp[-\alpha\tau_1(\gamma_C - \gamma_{CD}) - \alpha\tau_2(\gamma_B - \gamma_{AB}) - \alpha\tau_3(\gamma_{AB} + \gamma_{CD} - 2)]$ | $g_1 = 0$ | $g_2 = 0$ | $g_3 = 2\alpha$ |
| $d^{-(d_1+d_4+d_5)}$ | $g_0 = exp[-\alpha\tau_1(\gamma_D - \gamma_{CD}) - \alpha\tau_2(\gamma_B - \gamma_{AB}) - \alpha\tau_3(\gamma_{AB} + \gamma_{CD} - 2)]$ | $g_1 = 0$ | $g_2 = 0$ | $g_3 = 2\alpha$ |
| $d^{-(d_2+d_3+d_5)}$ | $g_0 = exp[-\alpha\tau_1(\gamma_C - \gamma_{CD}) - \alpha\tau_2(\gamma_A - \gamma_{AB}) - \alpha\tau_3(\gamma_{AB} + \gamma_{CD} - 2)]$ | $g_1 = 0$ | $g_2 = 2\alpha$ | $g_3 = 0$ |
| $d^{-(d_2+d_4+d_5)}$ | $g_0 = exp[-\alpha\tau_1(\gamma_D - \gamma_{CD}) - \alpha\tau_2(\gamma_A - \gamma_{AB}) - \alpha\tau_3(\gamma_{AB} + \gamma_{CD} - 2)]$ | $g_1 = 0$ | $g_2 = 2\alpha$ | $g_3 = 0$ |
| $d^{-(d_1+d_2+d_3+d_4)}$ | $g_0 = exp[-\alpha\tau_1(\gamma_C + \gamma_D - 2\gamma_{CD}) - \alpha\tau_2(\gamma_A + \gamma_B - 2\gamma_{AB}) - 2\alpha\tau_3(\gamma_{AB} + \gamma_{CD} - 2)]$ | $g_1 = 2\alpha$ | $g_2 = 0$ | $g_3 = 2\alpha$ |
| $d^{-(d_1+d_2+d_3+d_5)}$ | $g_0 = exp[-\alpha\tau_1(\gamma_C - \gamma_{CD}) - \alpha\tau_2(\gamma_A + \gamma_B - 2\gamma_{AB}) - \alpha\tau_3(2\gamma_{AB} + \gamma_{CD} - 3)]$ | $g_1 = 0$ | $g_2 = \alpha$ | $g_3 = 2\alpha$ |
| $d^{-(d_1+d_2+d_4+d_5)}$ | $g_0 = exp[-\alpha\tau_1(\gamma_D - \gamma_{CD}) - \alpha\tau_2(\gamma_A + \gamma_B - 2\gamma_{AB}) - \alpha\tau_3(2\gamma_{AB} + \gamma_{CD} - 3)]$ | $g_1 = 0$ | $g_2 = \alpha$ | $g_3 = 2\alpha$ |
| $d^{-(d_1+d_3+d_4+d_5)}$ | $g_0 = exp[-\alpha\tau_1(\gamma_C + \gamma_D - 2\gamma_{CD}) - \alpha\tau_2(\gamma_B - \gamma_{AB}) - \alpha\tau_3(\gamma_{AB} + 2\gamma_{CD} - 3)]$ | $g_1 = \alpha$ | $g_2 = 0$ | $g_3 = 2\alpha$ |
| $d^{-(d_2+d_3+d_4+d_5)}$ | $g_0 = exp[-\alpha\tau_1(\gamma_C + \gamma_D - 2\gamma_{CD}) - \alpha\tau_2(\gamma_A - \gamma_{AB}) - \alpha\tau_3(\gamma_{AB} + 2\gamma_{CD} - 3)]$ | $g_1 = \alpha$ | $g_2 = 2\alpha$ | $g_3 = 0$ |
| $d^{-(d_1+d_2+d_3+d_4+d_5)}$ | $g_0 = exp[-\alpha\tau_1(\gamma_C + \gamma_D - 2\gamma_{CD}) - \alpha\tau_2(\gamma_A + \gamma_B - 2\gamma_{AB}) - 2\alpha\tau_3(\gamma_{AB} + \gamma_{CD} - 2)]$ | $g_1 = \alpha$ | $g_2 = \alpha$ | $g_3 = 2\alpha$ |

| | **History 16** | | | |
|---|---|---|---|---|
| $d^{-(d_1+d_2)}$ | $g_0 = exp[-\alpha\tau_1(\gamma_C - \gamma_{CD}) - \alpha\tau_2(\gamma_B - \gamma_{AB}) - \alpha\tau_3(\gamma_{AB} + \gamma_{CD} - 2)]$ | $g_1 = 0$ | $g_2 = 0$ | $g_3 = 2\alpha$ |
| $d^{-(d_3+d_4)}$ | $g_0 = exp[-\alpha\tau_1(\gamma_D - \gamma_{CD}) - \alpha\tau_2(\gamma_A - \gamma_{AB}) - \alpha\tau_3(\gamma_{AB} + \gamma_{CD} - 2)]$ | $g_1 = 2\alpha$ | $g_2 = 0$ | $g_3 = 0$ |
| $d^{-(d_1+d_3+d_5)}$ | $g_0 = exp[-\alpha\tau_2(\gamma_A + \gamma_B - 2\gamma_{AB}) - 2\alpha\tau_3(\gamma_{AB} - 1)]$ | $g_1 = 0$ | $g_2 = 0$ | $g_3 = 2\alpha$ |
| $d^{-(d_1+d_4+d_5)}$ | $g_0 = exp[-\alpha\tau_1(\gamma_D - \gamma_{CD}) - \alpha\tau_2(\gamma_B - \gamma_{AB}) - \alpha\tau_3(\gamma_{AB} + \gamma_{CD} - 2)]$ | $g_1 = 0$ | $g_2 = 0$ | $g_3 = 2\alpha$ |
| $d^{-(d_2+d_3+d_5)}$ | $g_0 = exp[-\alpha\tau_1(\gamma_C - \gamma_{CD}) - \alpha\tau_2(\gamma_A - \gamma_{AB}) - \alpha\tau_3(\gamma_{AB} + \gamma_{CD} - 2)]$ | $g_1 = 0$ | $g_2 = 2\alpha$ | $g_3 = 0$ |
| $d^{-(d_2+d_4+d_5)}$ | $g_0 = exp[-\alpha\tau_1(\gamma_C + \gamma_D - 2\gamma_{CD}) - 2\alpha\tau_3(\gamma_{CD} - 1)]$ | $g_1 = 0$ | $g_2 = 2\alpha$ | $g_3 = 0$ |
| $d^{-(d_1+d_2+d_3+d_4)}$ | $g_0 = exp[-\alpha\tau_1(\gamma_C + \gamma_D - 2\gamma_{CD}) - \alpha\tau_2(\gamma_A + \gamma_B - 2\gamma_{AB}) - 2\alpha\tau_3(\gamma_{AB} + \gamma_{CD} - 2)]$ | $g_1 = 2\alpha$ | $g_2 = 0$ | $g_3 = 2\alpha$ |
| $d^{-(d_1+d_2+d_3+d_5)}$ | $g_0 = exp[-\alpha\tau_1(\gamma_C - \gamma_{CD}) - \alpha\tau_2(\gamma_A + \gamma_B - 2\gamma_{AB}) - \alpha\tau_3(2\gamma_{AB} + \gamma_{CD} - 3)]$ | $g_1 = 0$ | $g_2 = \alpha$ | $g_3 = 2\alpha$ |
| $d^{-(d_1+d_2+d_4+d_5)}$ | $g_0 = exp[-\alpha\tau_1(\gamma_C + \gamma_D - 2\gamma_{CD}) - \alpha\tau_2(\gamma_B - \gamma_{AB}) - \alpha\tau_3(\gamma_{AB} + 2\gamma_{CD} - 3)]$ | $g_1 = 0$ | $g_2 = \alpha$ | $g_3 = 2\alpha$ |
| $d^{-(d_1+d_3+d_4+d_5)}$ | $g_0 = exp[-\alpha\tau_1(\gamma_D - \gamma_{CD}) - \alpha\tau_2(\gamma_A + \gamma_B - 2\gamma_{AB}) - \alpha\tau_3(2\gamma_{AB} + \gamma_{CD} - 3)]$ | $g_1 = \alpha$ | $g_2 = 0$ | $g_3 = 2\alpha$ |
| $d^{-(d_2+d_3+d_4+d_5)}$ | $g_0 = exp[-\alpha\tau_1(\gamma_C + \gamma_D - 2\gamma_{CD}) - \alpha\tau_2(\gamma_A - \gamma_{AB}) - \alpha\tau_3(\gamma_{AB} + 2\gamma_{CD} - 3)]$ | $g_1 = \alpha$ | $g_2 = 2\alpha$ | $g_3 = 0$ |
| $d^{-(d_1+d_2+d_3+d_4+d_5)}$ | $g_0 = exp[-\alpha\tau_1(\gamma_C + \gamma_D - 2\gamma_{CD}) - \alpha\tau_2(\gamma_A + \gamma_B - 2\gamma_{AB}) - 2\alpha\tau_3(\gamma_{AB} + \gamma_{CD} - 2)]$ | $g_1 = \alpha$ | $g_2 = \alpha$ | $g_3 = 2\alpha$ |

| | **History 17** | | | |
|---|---|---|---|---|
| $d^{-(d_1+d_2)}$ | $g_0 = exp[-\alpha\tau_1(\gamma_D - \gamma_{CD}) - \alpha\tau_2(\gamma_B - \gamma_{AB}) - \alpha\tau_3(\gamma_{AB} + \gamma_{CD} - 2)]$ | $g_1 = 0$ | $g_2 = 0$ | $g_3 = 2\alpha$ |
| $d^{-(d_3+d_4)}$ | $g_0 = exp[-\alpha\tau_1(\gamma_C - \gamma_{CD}) - \alpha\tau_2(\gamma_A - \gamma_{AB}) - \alpha\tau_3(\gamma_{AB} + \gamma_{CD} - 2)]$ | $g_1 = 2\alpha$ | $g_2 = 0$ | $g_3 = 0$ |
| $d^{-(d_1+d_3+d_5)}$ | $g_0 = exp[-\alpha\tau_2(\gamma_A + \gamma_B - 2\gamma_{AB}) - 2\alpha\tau_3(\gamma_{AB} - 1)]$ | $g_1 = 0$ | $g_2 = 0$ | $g_3 = 2\alpha$ |
| $d^{-(d_1+d_4+d_5)}$ | $g_0 = exp[-\alpha\tau_1(\gamma_C - \gamma_{CD}) - \alpha\tau_2(\gamma_B - \gamma_{AB}) - \alpha\tau_3(\gamma_{AB} + \gamma_{CD} - 2)]$ | $g_1 = 0$ | $g_2 = 0$ | $g_3 = 2\alpha$ |
| $d^{-(d_2+d_3+d_5)}$ | $g_0 = exp[-\alpha\tau_1(\gamma_D - \gamma_{CD}) - \alpha\tau_2(\gamma_A - \gamma_{AB}) - \alpha\tau_3(\gamma_{AB} + \gamma_{CD} - 2)]$ | $g_1 = 0$ | $g_2 = 2\alpha$ | $g_3 = 0$ |
| $d^{-(d_2+d_4+d_5)}$ | $g_0 = exp[-\alpha\tau_1(\gamma_C + \gamma_D - 2\gamma_{CD}) - 2\alpha\tau_3(\gamma_{CD} - 1)]$ | $g_1 = 0$ | $g_2 = 2\alpha$ | $g_3 = 0$ |
| $d^{-(d_1+d_2+d_3+d_4)}$ | $g_0 = exp[-\alpha\tau_1(\gamma_C + \gamma_D - 2\gamma_{CD}) - \alpha\tau_2(\gamma_A + \gamma_B - 2\gamma_{AB}) - 2\alpha\tau_3(\gamma_{AB} + \gamma_{CD} - 2)]$ | $g_1 = 2\alpha$ | $g_2 = 0$ | $g_3 = 2\alpha$ |
| $d^{-(d_1+d_2+d_3+d_5)}$ | $g_0 = exp[-\alpha\tau_1(\gamma_D - \gamma_{CD}) - \alpha\tau_2(\gamma_A + \gamma_B - 2\gamma_{AB}) - \alpha\tau_3(2\gamma_{AB} + \gamma_{CD} - 3)]$ | $g_1 = 0$ | $g_2 = \alpha$ | $g_3 = 2\alpha$ |
| $d^{-(d_1+d_2+d_4+d_5)}$ | $g_0 = exp[-\alpha\tau_1(\gamma_C + \gamma_D - 2\gamma_{CD}) - \alpha\tau_2(\gamma_B - \gamma_{AB}) - \alpha\tau_3(\gamma_{AB} + 2\gamma_{CD} - 3)]$ | $g_1 = 0$ | $g_2 = \alpha$ | $g_3 = 2\alpha$ |
| $d^{-(d_1+d_3+d_4+d_5)}$ | $g_0 = exp[-\alpha\tau_1(\gamma_C - \gamma_{CD}) - \alpha\tau_2(\gamma_A + \gamma_B - 2\gamma_{AB}) - \alpha\tau_3(2\gamma_{AB} + \gamma_{CD} - 3)]$ | $g_1 = \alpha$ | $g_2 = 0$ | $g_3 = 2\alpha$ |
| $d^{-(d_2+d_3+d_4+d_5)}$ | $g_0 = exp[-\alpha\tau_1(\gamma_C + \gamma_D - 2\gamma_{CD}) - \alpha\tau_2(\gamma_A - \gamma_{AB}) - \alpha\tau_3(\gamma_{AB} + 2\gamma_{CD} - 3)]$ | $g_1 = \alpha$ | $g_2 = 2\alpha$ | $g_3 = 0$ |
| $d^{-(d_1+d_2+d_3+d_4+d_5)}$ | $g_0 = exp[-\alpha\tau_1(\gamma_C + \gamma_D - 2\gamma_{CD}) - \alpha\tau_2(\gamma_A + \gamma_B - 2\gamma_{AB}) - 2\alpha\tau_3(\gamma_{AB} + \gamma_{CD} - 2)]$ | $g_1 = \alpha$ | $g_2 = \alpha$ | $g_3 = 2\alpha$ |

| | **History 18** | | | |
|---|---|---|---|---|
| $d^{-(d_1+d_2)}$ | $g_0 = exp[-\alpha\tau_1(\gamma_C - \gamma_{CD}) - \alpha\tau_2(\gamma_A - \gamma_{AB}) - \alpha\tau_3(\gamma_{AB} + \gamma_{CD} - 2)]$ | $g_1 = 0$ | $g_2 = 0$ | $g_3 = 2\alpha$ |
| $d^{-(d_3+d_4)}$ | $g_0 = exp[-\alpha\tau_1(\gamma_D - \gamma_{CD}) - \alpha\tau_2(\gamma_B - \gamma_{AB}) - \alpha\tau_3(\gamma_{AB} + \gamma_{CD} - 2)]$ | $g_1 = 2\alpha$ | $g_2 = 0$ | $g_3 = 0$ |
| $d^{-(d_1+d_3+d_5)}$ | $g_0 = exp[-\alpha\tau_1(\gamma_C - \gamma_{CD}) - \alpha\tau_2(\gamma_B - \gamma_{AB}) - \alpha\tau_3(\gamma_{AB} + \gamma_{CD} - 2)]$ | $g_1 = 0$ | $g_2 = 0$ | $g_3 = 2\alpha$ |
| $d^{-(d_1+d_4+d_5)}$ | $g_0 = exp[-\alpha\tau_1(\gamma_C + \gamma_D - 2\gamma_{CD}) - 2\alpha\tau_3(\gamma_{CD} - 1)]$ | $g_1 = 0$ | $g_2 = 0$ | $g_3 = 2\alpha$ |
| $d^{-(d_2+d_3+d_5)}$ | $g_0 = exp[-\alpha\tau_2(\gamma_A + \gamma_B - 2\gamma_{AB}) - 2\alpha\tau_3(\gamma_{AB} - 1)]$ | $g_1 = 0$ | $g_2 = 2\alpha$ | $g_3 = 0$ |
| $d^{-(d_2+d_4+d_5)}$ | $g_0 = exp[-\alpha\tau_1(\gamma_D - \gamma_{CD}) - \alpha\tau_2(\gamma_A - \gamma_{AB}) - \alpha\tau_3(\gamma_{AB} + \gamma_{CD} - 2)]$ | $g_1 = 0$ | $g_2 = 2\alpha$ | $g_3 = 0$ |
| $d^{-(d_1+d_2+d_3+d_4)}$ | $g_0 = exp[-\alpha\tau_1(\gamma_C + \gamma_D - 2\gamma_{CD}) - \alpha\tau_2(\gamma_A + \gamma_B - 2\gamma_{AB}) - 2\alpha\tau_3(\gamma_{AB} + \gamma_{CD} - 2)]$ | $g_1 = 2\alpha$ | $g_2 = 0$ | $g_3 = 2\alpha$ |
| $d^{-(d_1+d_2+d_3+d_5)}$ | $g_0 = exp[-\alpha\tau_1(\gamma_C - \gamma_{CD}) - \alpha\tau_2(\gamma_A + \gamma_B - 2\gamma_{AB}) - \alpha\tau_3(2\gamma_{AB} + \gamma_{CD} - 3)]$ | $g_1 = 0$ | $g_2 = \alpha$ | $g_3 = 2\alpha$ |
| $d^{-(d_1+d_2+d_4+d_5)}$ | $g_0 = exp[-\alpha\tau_1(\gamma_C + \gamma_D - 2\gamma_{CD}) - \alpha\tau_2(\gamma_A - \gamma_{AB}) - \alpha\tau_3(\gamma_{AB} + 2\gamma_{CD} - 3)]$ | $g_1 = 0$ | $g_2 = \alpha$ | $g_3 = 2\alpha$ |
| $d^{-(d_1+d_3+d_4+d_5)}$ | $g_0 = exp[-\alpha\tau_1(\gamma_C + \gamma_D - 2\gamma_{CD}) - \alpha\tau_2(\gamma_B - \gamma_{AB}) - \alpha\tau_3(\gamma_{AB} + 2\gamma_{CD} - 3)]$ | $g_1 = \alpha$ | $g_2 = 0$ | $g_3 = 2\alpha$ |
| $d^{-(d_2+d_3+d_4+d_5)}$ | $g_0 = exp[-\alpha\tau_1(\gamma_D - \gamma_{CD}) - \alpha\tau_2(\gamma_A + \gamma_B - 2\gamma_{AB}) - \alpha\tau_3(2\gamma_{AB} + \gamma_{CD} - 3)]$ | $g_1 = \alpha$ | $g_2 = 2\alpha$ | $g_3 = 0$ |
| $d^{-(d_1+d_2+d_3+d_4+d_5)}$ | $g_0 = exp[-\alpha\tau_1(\gamma_C + \gamma_D - 2\gamma_{CD}) - \alpha\tau_2(\gamma_A + \gamma_B - 2\gamma_{AB}) - 2\alpha\tau_3(\gamma_{AB} + \gamma_{CD} - 2)]$ | $g_1 = \alpha$ | $g_2 = \alpha$ | $g_3 = 2\alpha$ |

| | **History 19** | | | |
|---|---|---|---|---|
| $d^{-(d_1+d_2)}$ | $g_0 = exp[-\alpha\tau_1(\gamma_C - \gamma_{CD}) - \alpha\tau_2(\gamma_B - \gamma_{AB}) - \alpha\tau_3(\gamma_{AB} + \gamma_{CD} - 2)]$ | $g_1 = 0$ | $g_2 = 0$ | $g_3 = 2\alpha$ |
| $d^{-(d_3+d_4)}$ | $g_0 = exp[-\alpha\tau_1(\gamma_D - \gamma_{CD}) - \alpha\tau_2(\gamma_A - \gamma_{AB}) - \alpha\tau_3(\gamma_{AB} + \gamma_{CD} - 2)]$ | $g_1 = 2\alpha$ | $g_2 = 0$ | $g_3 = 0$ |
| $d^{-(d_1+d_3+d_5)}$ | $g_0 = exp[-\alpha\tau_1(\gamma_C - \gamma_{CD}) - \alpha\tau_2(\gamma_A - \gamma_{AB}) - \alpha\tau_3(\gamma_{AB} + \gamma_{CD} - 2)]$ | $g_1 = 0$ | $g_2 = 0$ | $g_3 = 2\alpha$ |
| $d^{-(d_1+d_4+d_5)}$ | $g_0 = exp[-\alpha\tau_1(\gamma_C + \gamma_D - 2\gamma_{CD}) - 2\alpha\tau_3(\gamma_{CD} - 1)]$ | $g_1 = 0$ | $g_2 = 0$ | $g_3 = 2\alpha$ |
| $d^{-(d_2+d_3+d_5)}$ | $g_0 = exp[-\alpha\tau_2(\gamma_A + \gamma_B - 2\gamma_{AB}) - 2\alpha\tau_3(\gamma_{AB} - 1)]$ | $g_1 = 0$ | $g_2 = 2\alpha$ | $g_3 = 0$ |
| $d^{-(d_2+d_4+d_5)}$ | $g_0 = exp[-\alpha\tau_1(\gamma_D - \gamma_{CD}) - \alpha\tau_2(\gamma_B - \gamma_{AB}) - \alpha\tau_3(\gamma_{AB} + \gamma_{CD} - 2)]$ | $g_1 = 0$ | $g_2 = 2\alpha$ | $g_3 = 0$ |
| $d^{-(d_1+d_2+d_3+d_4)}$ | $g_0 = exp[-\alpha\tau_1(\gamma_C + \gamma_D - 2\gamma_{CD}) - \alpha\tau_2(\gamma_A + \gamma_B - 2\gamma_{AB}) - 2\alpha\tau_3(\gamma_{AB} + \gamma_{CD} - 2)]$ | $g_1 = 2\alpha$ | $g_2 = 0$ | $g_3 = 2\alpha$ |
| $d^{-(d_1+d_2+d_3+d_5)}$ | $g_0 = exp[-\alpha\tau_1(\gamma_C - \gamma_{CD}) - \alpha\tau_2(\gamma_A + \gamma_B - 2\gamma_{AB}) - \alpha\tau_3(2\gamma_{AB} + \gamma_{CD} - 3)]$ | $g_1 = 0$ | $g_2 = \alpha$ | $g_3 = 2\alpha$ |
| $d^{-(d_1+d_2+d_4+d_5)}$ | $g_0 = exp[-\alpha\tau_1(\gamma_C + \gamma_D - 2\gamma_{CD}) - \alpha\tau_2(\gamma_B - \gamma_{AB}) - \alpha\tau_3(\gamma_{AB} + 2\gamma_{CD} - 3)]$ | $g_1 = 0$ | $g_2 = \alpha$ | $g_3 = 2\alpha$ |
| $d^{-(d_1+d_3+d_4+d_5)}$ | $g_0 = exp[-\alpha\tau_1(\gamma_C + \gamma_D - 2\gamma_{CD}) - \alpha\tau_2(\gamma_A - \gamma_{AB}) - \alpha\tau_3(\gamma_{AB} + 2\gamma_{CD} - 3)]$ | $g_1 = \alpha$ | $g_2 = 0$ | $g_3 = 2\alpha$ |
| $d^{-(d_2+d_3+d_4+d_5)}$ | $g_0 = exp[-\alpha\tau_1(\gamma_D - \gamma_{CD}) - \alpha\tau_2(\gamma_A + \gamma_B - 2\gamma_{AB}) - \alpha\tau_3(2\gamma_{AB} + \gamma_{CD} - 3)]$ | $g_1 = \alpha$ | $g_2 = 2\alpha$ | $g_3 = 0$ |
| $d^{-(d_1+d_2+d_3+d_4+d_5)}$ | $g_0 = exp[-\alpha\tau_1(\gamma_C + \gamma_D - 2\gamma_{CD}) - \alpha\tau_2(\gamma_A + \gamma_B - 2\gamma_{AB}) - 2\alpha\tau_3(\gamma_{AB} + \gamma_{CD} - 2)]$ | $g_1 = \alpha$ | $g_2 = \alpha$ | $g_3 = 2\alpha$ |



| | **History 20** | | | |
|---|---|---|---|---|
| $d^{-(d_1+d_2)}$ | $g_0 = exp[-\alpha\tau_1(\gamma_C + \gamma_D - 2\gamma_{CD}) - 2\alpha\tau_3(\gamma_{CD} - 1)]$ | $g_1 = 0$ | $g_2 = 0$ | $g_3 = 2\alpha$ |
| $d^{-(d_3+d_4)}$ | $g_0 = exp[-\alpha\tau_2(\gamma_A + \gamma_B - 2\gamma_{AB})]$ | $g_1 = 2\alpha\gamma_{AB}$ | $g_2 = 0$ | $g_3 = 0$ |
| $d^{-(d_1+d_3+d_5)}$ | $g_0 = exp[-\alpha\tau_1(\gamma_C - \gamma_{CD}) - \alpha\tau_2(\gamma_A - \gamma_{AB}) - \alpha\tau_3(\gamma_{AB} + \gamma_{CD} - 2)]$ | $g_1 = 0$ | $g_2 = 0$ | $g_3 = 2\alpha$ |
| $d^{-(d_1+d_4+d_5)}$ | $g_0 = exp[-\alpha\tau_1(\gamma_C - \gamma_{CD}) - \alpha\tau_2(\gamma_B - \gamma_{AB}) - \alpha\tau_3(\gamma_{AB} - \gamma_{CD} - 2)]$ | $g_1 = 0$ | $g_2 = 0$ | $g_3 = 2\alpha$ |
| $d^{-(d_2+d_3+d_5)}$ | $g_0 = exp[-\alpha\tau_1(\gamma_D - \gamma_{CD}) - \alpha\tau_2(\gamma_A - \gamma_{AB}) - \alpha\tau_3(\gamma_{AB} - \gamma_{CD} - 2)]$ | $g_1 = 0$ | $g_2 = 2\alpha$ | $g_3 = 0$ |
| $d^{-(d_2+d_4+d_5)}$ | $g_0 = exp[-\alpha\tau_1(\gamma_D - \gamma_{CD}) - \alpha\tau_2(\gamma_B - \gamma_{AB}) - \alpha\tau_3(\gamma_{AB} + \gamma_{CD} - 2)]$ | $g_1 = 0$ | $g_2 = 2\alpha$ | $g_3 = 0$ |
| $d^{-(d_1+d_2+d_3+d_4)}$ | $g_0 = exp[-\alpha\tau_1(\gamma_C + \gamma_D - 2\gamma_{CD}) - \alpha\tau_2(\gamma_A + \gamma_B - 2\gamma_{AB}) - 2\alpha\tau_3(\gamma_{CD} - 1)]$ | $g_1 = 2\alpha\gamma_{AB}$ | $g_2 = 0$ | $g_3 = 2\alpha$ |
| $d^{-(d_1+d_2+d_3+d_5)}$ | $g_0 = exp[-\alpha\tau_1(\gamma_C + \gamma_D - 2\gamma_{CD}) - \alpha\tau_2(\gamma_A - \gamma_{AB}) - \alpha\tau_3(\gamma_{AB} + 2\gamma_{CD} - 3)]$ | $g_1 = 0$ | $g_2 = \alpha$ | $g_3 = 2\alpha$ |
| $d^{-(d_1+d_2+d_4+d_5)}$ | $g_0 = exp[-\alpha\tau_1(\gamma_C + \gamma_D - 2\gamma_{CD}) - \alpha\tau_2(\gamma_B - \gamma_{AB}) - \alpha\tau_3(\gamma_{AB} + 2\gamma_{CD} - 3)]$ | $g_1 = 0$ | $g_2 = \alpha$ | $g_3 = 2\alpha$ |
| $d^{-(d_1+d_3+d_4+d_5)}$ | $g_0 = exp[-\alpha\tau_1(\gamma_C - \gamma_{CD}) - \alpha\tau_2(\gamma_A + \gamma_B - 2\gamma_{AB}) - \alpha\tau_3(\gamma_{AB} + \gamma_{CD} - 2)]$ | $g_1 = \alpha\gamma_{AB}$ | $g_2 = 0$ | $g_3 = 2\alpha$ |
| $d^{-(d_2+d_3+d_4+d_5)}$ | $g_0 = exp[-\alpha\tau_1(\gamma_D - \gamma_{CD}) - \alpha\tau_2(\gamma_A + \gamma_B - 2\gamma_{AB}) - \alpha\tau_3(\gamma_{AB} + \gamma_{CD} - 2)]$ | $g_1 = \alpha\gamma_{AB}$ | $g_2 = 2\alpha$ | $g_3 = 0$ |
| $d^{-(d_1+d_2+d_3+d_4+d_5)}$ | $g_0 = exp[-\alpha\tau_1(\gamma_C + \gamma_D - 2\gamma_{CD}) - \alpha\tau_2(\gamma_A + \gamma_B - 2\gamma_{AB}) - \alpha\tau_3(\gamma_{AB} + 2\gamma_{CD} - 3)]$ | $g_1 = \alpha\gamma_{AB}$ | $g_2 = \alpha$ | $g_3 = 2\alpha$ |

| | **History 21** | | | |
|---|---|---|---|---|
| $d^{-(d_1+d_2)}$ | $g_0 = exp[-\alpha\tau_1(\gamma_C + \gamma_D - 2\gamma_{CD}) - 2\alpha\tau_3(\gamma_{CD} - 1)]$ | $g_1 = 0$ | $g_2 = 0$ | $g_3 = 2\alpha$ |
| $d^{-(d_3+d_4)}$ | $g_0 = exp[-\alpha\tau_2(\gamma_A + \gamma_B - 2\gamma_{AB}) - 2\alpha\tau_3(\gamma_{AB} - 1)]$ | $g_1 = 2\alpha$ | $g_2 = 0$ | $g_3 = 0$ |
| $d^{-(d_1+d_3+d_5)}$ | $g_0 = exp[-\alpha\tau_1(\gamma_C - \gamma_{CD}) - \alpha\tau_2(\gamma_A - \gamma_{AB}) - \alpha\tau_3(\gamma_{AB} + \gamma_{CD} - 2)]$ | $g_1 = 0$ | $g_2 = 0$ | $g_3 = 2\alpha$ |
| $d^{-(d_1+d_4+d_5)}$ | $g_0 = exp[-\alpha\tau_1(\gamma_C - \gamma_{CD}) - \alpha\tau_2(\gamma_B - \gamma_{AB}) - \alpha\tau_3(\gamma_{AB} - \gamma_{CD} - 2)]$ | $g_1 = 0$ | $g_2 = 0$ | $g_3 = 2\alpha$ |
| $d^{-(d_2+d_3+d_5)}$ | $g_0 = exp[-\alpha\tau_1(\gamma_D - \gamma_{CD}) - \alpha\tau_2(\gamma_A - \gamma_{AB}) - \alpha\tau_3(\gamma_{AB} - \gamma_{CD} - 2)]$ | $g_1 = 0$ | $g_2 = 2\alpha$ | $g_3 = 0$ |
| $d^{-(d_2+d_4+d_5)}$ | $g_0 = exp[-\alpha\tau_1(\gamma_D - \gamma_{CD}) - \alpha\tau_2(\gamma_B - \gamma_{AB}) - \alpha\tau_3(\gamma_{AB} + \gamma_{CD} - 2)]$ | $g_1 = 0$ | $g_2 = 2\alpha$ | $g_3 = 0$ |
| $d^{-(d_1+d_2+d_3+d_4)}$ | $g_0 = exp[-\alpha\tau_1(\gamma_C + \gamma_D - 2\gamma_{CD}) - \alpha\tau_2(\gamma_A + \gamma_B - 2\gamma_{AB}) - 2\alpha\tau_3(\gamma_{AB} + \gamma_{CD} - 2)]$ | $g_1 = 2\alpha$ | $g_2 = 0$ | $g_3 = 2\alpha$ |
| $d^{-(d_1+d_2+d_3+d_5)}$ | $g_0 = exp[-\alpha\tau_1(\gamma_C + \gamma_D - 2\gamma_{CD}) - \alpha\tau_2(\gamma_A - \gamma_{AB}) - \alpha\tau_3(\gamma_{AB} + 2\gamma_{CD} - 3)]$ | $g_1 = 0$ | $g_2 = \alpha$ | $g_3 = 2\alpha$ |
| $d^{-(d_1+d_2+d_4+d_5)}$ | $g_0 = exp[-\alpha\tau_1(\gamma_C + \gamma_D - 2\gamma_{CD}) - \alpha\tau_2(\gamma_B - \gamma_{AB}) - \alpha\tau_3(\gamma_{AB} + 2\gamma_{CD} - 3)]$ | $g_1 = 0$ | $g_2 = \alpha$ | $g_3 = 2\alpha$ |
| $d^{-(d_1+d_3+d_4+d_5)}$ | $g_0 = exp[-\alpha\tau_1(\gamma_C - \gamma_{CD}) - \alpha\tau_2(\gamma_A + \gamma_B - 2\gamma_{AB}) - \alpha\tau_3(2\gamma_{AB} + \gamma_{CD} - 3)]$ | $g_1 = \alpha$ | $g_2 = 0$ | $g_3 = 2\alpha$ |
| $d^{-(d_2+d_3+d_4+d_5)}$ | $g_0 = exp[-\alpha\tau_1(\gamma_D - \gamma_{CD}) - \alpha\tau_2(\gamma_A + \gamma_B - 2\gamma_{AB}) - \alpha\tau_3(2\gamma_{AB} + \gamma_{CD} - 3)]$ | $g_1 = \alpha$ | $g_2 = 2\alpha$ | $g_3 = 0$ |
| $d^{-(d_1+d_2+d_3+d_4+d_5)}$ | $g_0 = exp[-\alpha\tau_1(\gamma_C + \gamma_D - 2\gamma_{CD}) - \alpha\tau_2(\gamma_A + \gamma_B - 2\gamma_{AB}) - 2\alpha\tau_3(\gamma_{AB} + \gamma_{CD} - 2)]$ | $g_1 = \alpha$ | $g_2 = \alpha$ | $g_3 = 2\alpha$ |



| | **History 22** | | | |
|---|---|---|---|---|
| $d^{-(d_1+d_2)}$ | $g_0 = exp[-\alpha\tau_1(\gamma_D - \gamma_{CD}) - \alpha\tau_2(\gamma_A - \gamma_{AB}) - \alpha\tau_3(\gamma_{AB} + \gamma_{CD} - 2)]$ | $g_1 = 0$ | $g_2 = 0$ | $g_3 = 2\alpha$ |
| $d^{-(d_3+d_4)}$ | $g_0 = exp[-\alpha\tau_1(\gamma_C - \gamma_{CD}) - \alpha\tau_2(\gamma_B - \gamma_{AB}) - \alpha\tau_3(\gamma_{AB} + \gamma_{CD} - 2)]$ | $g_1 = 2\alpha$ | $g_2 = 0$ | $g_3 = 0$ |
| $d^{-(d_1+d_3+d_5)}$ | $g_0 = exp[-\alpha\tau_1(\gamma_D - \gamma_{CD}) - \alpha\tau_2(\gamma_B - \gamma_{AB}) - \alpha\tau_3(\gamma_{AB} + \gamma_{CD} - 2)]$ | $g_1 = 0$ | $g_2 = 0$ | $g_3 = 2\alpha$ |
| $d^{-(d_1+d_4+d_5)}$ | $g_0 = exp[-\alpha\tau_1(\gamma_C + \gamma_D - 2\gamma_{CD}) - 2\alpha\tau_3(\gamma_{CD} - 1)]$ | $g_1 = 0$ | $g_2 = 0$ | $g_3 = 2\alpha$ |
| $d^{-(d_2+d_3+d_5)}$ | $g_0 = exp[-\alpha\tau_2(\gamma_A + \gamma_B - 2\gamma_{AB}) - 2\alpha\tau_3(\gamma_{AB} - 1)]$ | $g_1 = 0$ | $g_2 = 2\alpha$ | $g_3 = 0$ |
| $d^{-(d_2+d_4+d_5)}$ | $g_0 = exp[-\alpha\tau_1(\gamma_C - \gamma_{CD}) - \alpha\tau_2(\gamma_A - \gamma_{AB}) - \alpha\tau_3(\gamma_{AB} + \gamma_{CD} - 2)]$ | $g_1 = 0$ | $g_2 = 2\alpha$ | $g_3 = 0$ |
| $d^{-(d_1+d_2+d_3+d_4)}$ | $g_0 = exp[-\alpha\tau_1(\gamma_C + \gamma_D - 2\gamma_{CD}) - \alpha\tau_2(\gamma_A + \gamma_B - 2\gamma_{AB}) - 2\alpha\tau_3(\gamma_{AB} + \gamma_{CD} - 2)]$ | $g_1 = 2\alpha$ | $g_2 = 0$ | $g_3 = 2\alpha$ |
| $d^{-(d_1+d_2+d_3+d_5)}$ | $g_0 = exp[-\alpha\tau_1(\gamma_D - \gamma_{CD}) - \alpha\tau_2(\gamma_A + \gamma_B - 2\gamma_{AB}) - \alpha\tau_3(2\gamma_{AB} + \gamma_{CD} - 3)]$ | $g_1 = 0$ | $g_2 = \alpha$ | $g_3 = 2\alpha$ |
| $d^{-(d_1+d_2+d_4+d_5)}$ | $g_0 = exp[-\alpha\tau_1(\gamma_C + \gamma_D - 2\gamma_{CD}) - \alpha\tau_2(\gamma_A - \gamma_{AB}) - \alpha\tau_3(\gamma_{AB} + 2\gamma_{CD} - 3)]$ | $g_1 = 0$ | $g_2 = \alpha$ | $g_3 = 2\alpha$ |
| $d^{-(d_1+d_3+d_4+d_5)}$ | $g_0 = exp[-\alpha\tau_1(\gamma_C + \gamma_D - 2\gamma_{CD}) - \alpha\tau_2(\gamma_B - \gamma_{AB}) - \alpha\tau_3(\gamma_{AB} + 2\gamma_{CD} - 3)]$ | $g_1 = \alpha$ | $g_2 = 0$ | $g_3 = 2\alpha$ |
| $d^{-(d_2+d_3+d_4+d_5)}$ | $g_0 = exp[-\alpha\tau_1(\gamma_C - \gamma_{CD}) - \alpha\tau_2(\gamma_A + \gamma_B - 2\gamma_{AB}) - \alpha\tau_3(2\gamma_{AB} + \gamma_{CD} - 3)]$ | $g_1 = \alpha$ | $g_2 = 2\alpha$ | $g_3 = 0$ |
| $d^{-(d_1+d_2+d_3+d_4+d_5)}$ | $g_0 = exp[-\alpha\tau_1(\gamma_C + \gamma_D - 2\gamma_{CD}) - \alpha\tau_2(\gamma_A + \gamma_B - 2\gamma_{AB}) - 2\alpha\tau_3(\gamma_{AB} + \gamma_{CD} - 2)]$ | $g_1 = \alpha$ | $g_2 = \alpha$ | $g_3 = 2\alpha$ |

| | **History 23** | | | |
|---|---|---|---|---|
| $d^{-(d_1+d_2)}$ | $g_0 = exp[-\alpha\tau_1(\gamma_D - \gamma_{CD}) - \alpha\tau_2(\gamma_B - \gamma_{AB}) - \alpha\tau_3(\gamma_{AB} + \gamma_{CD} - 2)]$ | $g_1 = 0$ | $g_2 = 0$ | $g_3 = 2\alpha$ |
| $d^{-(d_3+d_4)}$ | $g_0 = exp[-\alpha\tau_1(\gamma_C - \gamma_{CD}) - \alpha\tau_2(\gamma_A - \gamma_{AB}) - \alpha\tau_3(\gamma_{AB} + \gamma_{CD} - 2)]$ | $g_1 = 2\alpha$ | $g_2 = 0$ | $g_3 = 0$ |
| $d^{-(d_1+d_3+d_5)}$ | $g_0 = exp[-\alpha\tau_1(\gamma_D - \gamma_{CD}) - \alpha\tau_2(\gamma_A - \gamma_{AB}) - \alpha\tau_3(\gamma_{AB} + \gamma_{CD} - 2)]$ | $g_1 = 0$ | $g_2 = 0$ | $g_3 = 2\alpha$ |
| $d^{-(d_1+d_4+d_5)}$ | $g_0 = exp[-\alpha\tau_1(\gamma_C + \gamma_D - 2\gamma_{CD}) - 2\alpha\tau_3(\gamma_{CD} - 1)]$ | $g_1 = 0$ | $g_2 = 0$ | $g_3 = 2\alpha$ |
| $d^{-(d_2+d_3+d_5)}$ | $g_0 = exp[-\alpha\tau_2(\gamma_A + \gamma_B - 2\gamma_{AB}) - 2\alpha\tau_3(\gamma_{AB} - 1)]$ | $g_1 = 0$ | $g_2 = 2\alpha$ | $g_3 = 0$ |
| $d^{-(d_2+d_4+d_5)}$ | $g_0 = exp[-\alpha\tau_1(\gamma_C - \gamma_{CD}) - \alpha\tau_2(\gamma_B - \gamma_{AB}) - \alpha\tau_3(\gamma_{AB} + \gamma_{CD} - 2)]$ | $g_1 = 0$ | $g_2 = 2\alpha$ | $g_3 = 0$ |
| $d^{-(d_1+d_2+d_3+d_4)}$ | $g_0 = exp[-\alpha\tau_1(\gamma_C + \gamma_D - 2\gamma_{CD}) - \alpha\tau_2(\gamma_A + \gamma_B - 2\gamma_{AB}) - 2\alpha\tau_3(\gamma_{AB} + \gamma_{CD} - 2)]$ | $g_1 = 2\alpha$ | $g_2 = 0$ | $g_3 = 2\alpha$ |
| $d^{-(d_1+d_2+d_3+d_5)}$ | $g_0 = exp[-\alpha\tau_1(\gamma_D - \gamma_{CD}) - \alpha\tau_2(\gamma_A + \gamma_B - 2\gamma_{AB}) - \alpha\tau_3(2\gamma_{AB} + \gamma_{CD} - 3)]$ | $g_1 = 0$ | $g_2 = \alpha$ | $g_3 = 2\alpha$ |
| $d^{-(d_1+d_2+d_4+d_5)}$ | $g_0 = exp[-\alpha\tau_1(\gamma_C + \gamma_D - 2\gamma_{CD}) - \alpha\tau_2(\gamma_B - \gamma_{AB}) - \alpha\tau_3(\gamma_{AB} + 2\gamma_{CD} - 3)]$ | $g_1 = 0$ | $g_2 = \alpha$ | $g_3 = 2\alpha$ |
| $d^{-(d_1+d_3+d_4+d_5)}$ | $g_0 = exp[-\alpha\tau_1(\gamma_C + \gamma_D - 2\gamma_{CD}) - \alpha\tau_2(\gamma_A - \gamma_{AB}) - \alpha\tau_3(\gamma_{AB} + 2\gamma_{CD} - 3)]$ | $g_1 = \alpha$ | $g_2 = 0$ | $g_3 = 2\alpha$ |
| $d^{-(d_2+d_3+d_4+d_5)}$ | $g_0 = exp[-\alpha\tau_1(\gamma_C - \gamma_{CD}) - \alpha\tau_2(\gamma_A + \gamma_B - 2\gamma_{AB}) - \alpha\tau_3(2\gamma_{AB} + \gamma_{CD} - 3)]$ | $g_1 = \alpha$ | $g_2 = 2\alpha$ | $g_3 = 0$ |
| $d^{-(d_1+d_2+d_3+d_4+d_5)}$ | $g_0 = exp[-\alpha\tau_1(\gamma_C + \gamma_D - 2\gamma_{CD}) - \alpha\tau_2(\gamma_A + \gamma_B - 2\gamma_{AB}) - 2\alpha\tau_3(\gamma_{AB} + \gamma_{CD} - 2)]$ | $g_1 = \alpha$ | $g_2 = \alpha$ | $g_3 = 2\alpha$ |

| | **History 24** | | | |
|---|---|---|---|---|
| $d^{-(d_1+d_2)}$ | $g_0 = exp[-\alpha\tau_1(\gamma_C + \gamma_D - 2\gamma_{CD}) - 2\alpha\tau_3(\gamma_{CD} - 1)]$ | $g_1 = 0$ | $g_2 = 0$ | $g_3 = 2\alpha$ |
| $d^{-(d_3+d_4)}$ | $g_0 = exp[-\alpha\tau_2(\gamma_A + \gamma_B - 2\gamma_{AB})]$ | $g_1 = 2\alpha\gamma_{AB}$ | $g_2 = 0$ | $g_3 = 0$ |
| $d^{-(d_1+d_3+d_5)}$ | $g_0 = exp[-\alpha\tau_1(\gamma_D - \gamma_{CD}) - \alpha\tau_2(\gamma_A - \gamma_{AB}) - \alpha\tau_3(\gamma_{AB} + \gamma_{CD} - 2)]$ | $g_1 = 0$ | $g_2 = 0$ | $g_3 = 2\alpha$ |
| $d^{-(d_1+d_4+d_5)}$ | $g_0 = exp[-\alpha\tau_1(\gamma_D - \gamma_{CD}) - \alpha\tau_2(\gamma_B - \gamma_{AB}) - \alpha\tau_3(\gamma_{AB} + \gamma_{CD} - 2)]$ | $g_1 = 0$ | $g_2 = 0$ | $g_3 = 2\alpha$ |
| $d^{-(d_2+d_3+d_5)}$ | $g_0 = exp[-\alpha\tau_1(\gamma_C - \gamma_{CD}) - \alpha\tau_2(\gamma_A - \gamma_{AB}) - \alpha\tau_3(\gamma_{AB} + \gamma_{CD} - 2)]$ | $g_1 = 0$ | $g_2 = 2\alpha$ | $g_3 = 0$ |
| $d^{-(d_2+d_4+d_5)}$ | $g_0 = exp[-\alpha\tau_1(\gamma_C - \gamma_{CD}) - \alpha\tau_2(\gamma_B - \gamma_{AB}) - \alpha\tau_3(\gamma_{AB} + \gamma_{CD} - 2)]$ | $g_1 = 0$ | $g_2 = 2\alpha$ | $g_3 = 0$ |
| $d^{-(d_1+d_2+d_3+d_4)}$ | $g_0 = exp[-\alpha\tau_1(\gamma_C + \gamma_D - 2\gamma_{CD}) - \alpha\tau_2(\gamma_A + \gamma_B - 2\gamma_{AB}) - 2\alpha\tau_3(\gamma_{CD} - 1)]$ | $g_1 = 2\alpha\gamma_{AB}$ | $g_2 = 0$ | $g_3 = 2\alpha$ |
| $d^{-(d_1+d_2+d_3+d_5)}$ | $g_0 = exp[-\alpha\tau_1(\gamma_C + \gamma_D - 2\gamma_{CD}) - \alpha\tau_2(\gamma_A - \gamma_{AB}) - \alpha\tau_3(\gamma_{AB} + 2\gamma_{CD} - 3)]$ | $g_1 = 0$ | $g_2 = \alpha$ | $g_3 = 2\alpha$ |
| $d^{-(d_1+d_2+d_4+d_5)}$ | $g_0 = exp[-\alpha\tau_1(\gamma_C + \gamma_D - 2\gamma_{CD}) - \alpha\tau_2(\gamma_B - \gamma_{AB}) - \alpha\tau_3(\gamma_{AB} + 2\gamma_{CD} - 3)]$ | $g_1 = 0$ | $g_2 = \alpha$ | $g_3 = 2\alpha$ |
| $d^{-(d_1+d_3+d_4+d_5)}$ | $g_0 = exp[-\alpha\tau_1(\gamma_D - \gamma_{CD}) - \alpha\tau_2(\gamma_A + \gamma_B - 2\gamma_{AB}) - \alpha\tau_3(\gamma_{AB} + \gamma_{CD} - 2)]$ | $g_1 = \alpha\gamma_{AB}$ | $g_2 = 0$ | $g_3 = 2\alpha$ |
| $d^{-(d_2+d_3+d_4+d_5)}$ | $g_0 = exp[-\alpha\tau_1(\gamma_C - \gamma_{CD}) - \alpha\tau_2(\gamma_A + \gamma_B - 2\gamma_{AB}) - \alpha\tau_3(\gamma_{AB} + \gamma_{CD} - 2)]$ | $g_1 = \alpha\gamma_{AB}$ | $g_2 = 2\alpha$ | $g_3 = 0$ |
| $d^{-(d_1+d_2+d_3+d_4+d_5)}$ | $g_0 = exp[-\alpha\tau_1(\gamma_C + \gamma_D - 2\gamma_{CD}) - \alpha\tau_2(\gamma_A + \gamma_B - 2\gamma_{AB}) - \alpha\tau_3(\gamma_{AB} + 2\gamma_{CD} - 3)]$ | $g_1 = \alpha\gamma_{AB}$ | $g_2 = \alpha$ | $g_3 = 2\alpha$ |

| | **History 25** | | | |
|---|---|---|---|---|
| $d^{-(d_1+d_2)}$ | $g_0 = exp[-\alpha\tau_1(\gamma_C + \gamma_D - 2\gamma_{CD}) - 2\alpha\tau_3(\gamma_{CD} - 1)]$ | $g_1 = 0$ | $g_2 = 0$ | $g_3 = 2\alpha$ |
| $d^{-(d_3+d_4)}$ | $g_0 = exp[-\alpha\tau_2(\gamma_A + \gamma_B - 2\gamma_{AB}) - 2\alpha\tau_3(\gamma_{AB} - 1)]$ | $g_1 = 2\alpha$ | $g_2 = 0$ | $g_3 = 0$ |
| $d^{-(d_1+d_3+d_5)}$ | $g_0 = exp[-\alpha\tau_1(\gamma_D - \gamma_{CD}) - \alpha\tau_2(\gamma_A - \gamma_{AB}) - \alpha\tau_3(\gamma_{AB} + \gamma_{CD} - 2)]$ | $g_1 = 0$ | $g_2 = 0$ | $g_3 = 2\alpha$ |
| $d^{-(d_1+d_4+d_5)}$ | $g_0 = exp[-\alpha\tau_1(\gamma_D - \gamma_{CD}) - \alpha\tau_2(\gamma_B - \gamma_{AB}) - \alpha\tau_3(\gamma_{AB} + \gamma_{CD} - 2)]$ | $g_1 = 0$ | $g_2 = 0$ | $g_3 = 2\alpha$ |
| $d^{-(d_2+d_3+d_5)}$ | $g_0 = exp[-\alpha\tau_1(\gamma_C - \gamma_{CD}) - \alpha\tau_2(\gamma_A - \gamma_{AB}) - \alpha\tau_3(\gamma_{AB} + \gamma_{CD} - 2)]$ | $g_1 = 0$ | $g_2 = 2\alpha$ | $g_3 = 0$ |
| $d^{-(d_2+d_4+d_5)}$ | $g_0 = exp[-\alpha\tau_1(\gamma_C - \gamma_{CD}) - \alpha\tau_2(\gamma_B - \gamma_{AB}) - \alpha\tau_3(\gamma_{AB} + \gamma_{CD} - 2)]$ | $g_1 = 0$ | $g_2 = 2\alpha$ | $g_3 = 0$ |
| $d^{-(d_1+d_2+d_3+d_4)}$ | $g_0 = exp[-\alpha\tau_1(\gamma_C + \gamma_D - 2\gamma_{CD}) - \alpha\tau_2(\gamma_A + \gamma_B - 2\gamma_{AB}) - 2\alpha\tau_3(\gamma_{AB} + \gamma_{CD} - 2)]$ | $g_1 = 2\alpha$ | $g_2 = 0$ | $g_3 = 2\alpha$ |
| $d^{-(d_1+d_2+d_3+d_5)}$ | $g_0 = exp[-\alpha\tau_1(\gamma_C + \gamma_D - 2\gamma_{CD}) - \alpha\tau_2(\gamma_A - \gamma_{AB}) - \alpha\tau_3(\gamma_{AB} + 2\gamma_{CD} - 3)]$ | $g_1 = 0$ | $g_2 = \alpha$ | $g_3 = 2\alpha$ |
| $d^{-(d_1+d_2+d_4+d_5)}$ | $g_0 = exp[-\alpha\tau_1(\gamma_C + \gamma_D - 2\gamma_{CD}) - \alpha\tau_2(\gamma_B - \gamma_{AB}) - \alpha\tau_3(\gamma_{AB} + 2\gamma_{CD} - 3)]$ | $g_1 = 0$ | $g_2 = \alpha$ | $g_3 = 2\alpha$ |
| $d^{-(d_1+d_3+d_4+d_5)}$ | $g_0 = exp[-\alpha\tau_1(\gamma_D - \gamma_{CD}) - \alpha\tau_2(\gamma_A + \gamma_B - 2\gamma_{AB}) - \alpha\tau_3(2\gamma_{AB} + \gamma_{CD} - 3)]$ | $g_1 = \alpha$ | $g_2 = 0$ | $g_3 = 2\alpha$ |
| $d^{-(d_2+d_3+d_4+d_5)}$ | $g_0 = exp[-\alpha\tau_1(\gamma_C - \gamma_{CD}) - \alpha\tau_2(\gamma_A + \gamma_B - 2\gamma_{AB}) - \alpha\tau_3(2\gamma_{AB} + \gamma_{CD} - 3)]$ | $g_1 = \alpha$ | $g_2 = 2\alpha$ | $g_3 = 0$ |
| $d^{-(d_1+d_2+d_3+d_4+d_5)}$ | $g_0 = exp[-\alpha\tau_1(\gamma_C + \gamma_D - 2\gamma_{CD}) - \alpha\tau_2(\gamma_A + \gamma_B - 2\gamma_{AB}) - 2\alpha\tau_3(\gamma_{AB} + \gamma_{CD} - 2)]$ | $g_1 = \alpha$ | $g_2 = \alpha$ | $g_3 = 2\alpha$ |

#### 3.2 Asymmetric species tree

| | **History 1** | | | |
|---|---|---|---|---|
| $d^{-(d_1+d_2)}$ | $g_0 = exp[-\alpha\tau_2(\gamma_B - \gamma_{BCD}) - \alpha\tau_3(\gamma_A + \gamma_{BCD} - 2)]$ | $g_1 = 0$ | $g_2 = 2\alpha$ | $g_3 = 0$ |
| $d^{-(d_3+d_4)}$ | $g_0 = exp[-\alpha\tau_1(\gamma_C + \gamma_D - 2\gamma_{CD}))]$ | $g_1 = 2\alpha\gamma_{CD}$ | $g_2 = 0$ | $g_3 = 0$ |
| $d^{-(d_1+d_3+d_5)}$ | $g_0 = exp[-\alpha\tau_1(\gamma_C - \gamma_{CD}) - \alpha\tau_2(\gamma_{CD} - \gamma_{BCD}) - \alpha\tau_3(\gamma_A + \gamma_{BCD} - 2)]$ | $g_1 = 0$ | $g_2 = 0$ | $g_3 = 2\alpha$ |
| $d^{-(d_1+d_4+d_5)}$ | $g_0 = exp[-\alpha\tau_1(\gamma_D - \gamma_{CD}) - \alpha\tau_2(\gamma_{CD} - \gamma_{BCD}) - \alpha\tau_3(\gamma_A + \gamma_{BCD} - 2)]$ | $g_1 = 0$ | $g_2 = 0$ | $g_3 = 2\alpha$ |
| $d^{-(d_2+d_3+d_5)}$ | $g_0 = exp[-\alpha\tau_1(\gamma_C - \gamma_{CD}) - \alpha\tau_2(\gamma_B + \gamma_{CD} - 2\gamma_{BCD}) - 2\alpha\tau_3(\gamma_{BCD} - 1)]$ | $g_1 = 0$ | $g_2 = 0$ | $g_3 = 2\alpha$ |
| $d^{-(d_2+d_4+d_5)}$ | $g_0 = exp[-\alpha\tau_1(\gamma_D - \gamma_{CD}) - \alpha\tau_2(\gamma_B + \gamma_{CD} - 2\gamma_{BCD}) - 2\alpha\tau_3(\gamma_{BCD} - 1)]$ | $g_1 = 0$ | $g_2 = 0$ | $g_3 = 2\alpha$ |
| $d^{-(d_1+d_2+d_3+d_4)}$ | $g_0 = exp[-\alpha\tau_1(\gamma_C + \gamma_D - 2\gamma_{CD}) - \alpha\tau_2(\gamma_B - \gamma_{BCD}) - \alpha\tau_3(\gamma_A + \gamma_{BCD} - 2)]$ | $g_1 = 2\alpha\gamma_{CD}$ | $g_2 = 2\alpha$ | $g_3 = 0$ |
| $d^{-(d_1+d_2+d_3+d_5)}$ | $g_0 = exp[-\alpha\tau_1(\gamma_C - \gamma_{CD}) - \alpha\tau_2(\gamma_B + \gamma_{CD} - 2\gamma_{BCD}) - \alpha\tau_3(\gamma_A + 2\gamma_{BCD} - 3)]$ | $g_1 = 0$ | $g_2 = \alpha$ | $g_3 = 2\alpha$ |
| $d^{-(d_1+d_2+d_4+d_5)}$ | $g_0 = exp[-\alpha\tau_1(\gamma_D - \gamma_{CD}) - \alpha\tau_2(\gamma_B + \gamma_{CD} - 2\gamma_{BCD}) - \alpha\tau_3(\gamma_A + 2\gamma_{BCD} - 3)]$ | $g_1 = 0$ | $g_2 = \alpha$ | $g_3 = 2\alpha$ |
| $d^{-(d_1+d_3+d_4+d_5)}$ | $g_0 = exp[-\alpha\tau_1(\gamma_C + \gamma_D - 2\gamma_{CD}) - \alpha\tau_2(\gamma_{CD} - \gamma_{BCD}) - \alpha\tau_3(\gamma_A + \gamma_{BCD} - 2)]$ | $g_1 = \alpha\gamma_{CD}$ | $g_2 = 0$ | $g_3 = 2\alpha$ |
| $d^{-(d_2+d_3+d_4+d_5)}$ | $g_0 = exp[-\alpha\tau_1(\gamma_C + \gamma_D - 2\gamma_{CD}) - \alpha\tau_2(\gamma_B + \gamma_{CD} - 2\gamma_{BCD}) - 2\alpha\tau_3(\gamma_{BCD} - 1)]$ | $g_1 = \alpha\gamma_{CD}$ | $g_2 = 0$ | $g_3 = 2\alpha$ |
| $d^{-(d_1+d_2+d_3+d_4+d_5)}$ | $g_0 = exp[-\alpha\tau_1(\gamma_C + \gamma_D - 2\gamma_{CD}) - \alpha\tau_2(\gamma_B + \gamma_{CD} - 2\gamma_{BCD}) - \alpha\tau_3(\gamma_A + 2\gamma_{BCD} - 3)]$ | $g_1 = \alpha\gamma_{CD}$ | $g_2 = \alpha$ | $g_3 = 2\alpha$ |



| | **History 2** | | | |
|---|---|---|---|---|
| $d^{-(d_1+d_2)}$ | $g_0 = exp[-\alpha\tau_2(\gamma_B - \gamma_{BCD}) - \alpha\tau_3(\gamma_A + \gamma_{BCD} - 2)]$ | $g_1 = 0$ | $g_2 = 2\alpha$ | $g_3 = 0$ |
| $d^{-(d_3+d_4)}$ | $g_0 = exp[-\alpha\tau_1(\gamma_C + \gamma_D - 2\gamma_{CD}) - 2\alpha\tau_2(\gamma_{CD} - \gamma_{BCD})]$ | $g_1 = 2\alpha\gamma_{BCD}$ | $g_2 = 0$ | $g_3 = 0$ |
| $d^{-(d_1+d_3+d_5)}$ | $g_0 = exp[-\alpha\tau_1(\gamma_C - \gamma_{CD}) - \alpha\tau_2(\gamma_{CD} - \gamma_{BCD}) - \alpha\tau_3(\gamma_A + \gamma_{BCD} - 2)]$ | $g_1 = 0$ | $g_2 = 0$ | $g_3 = 2\alpha$ |
| $d^{-(d_1+d_4+d_5)}$ | $g_0 = exp[-\alpha\tau_1(\gamma_D - \gamma_{CD}) - \alpha\tau_2(\gamma_{CD} - \gamma_{BCD}) - \alpha\tau_3(\gamma_A + \gamma_{BCD} - 2)]$ | $g_1 = 0$ | $g_2 = 0$ | $g_3 = 2\alpha$ |
| $d^{-(d_2+d_3+d_5)}$ | $g_0 = exp[-\alpha\tau_1(\gamma_C - \gamma_{CD}) - \alpha\tau_2(\gamma_B + \gamma_{CD} - 2\gamma_{BCD}) - 2\alpha\tau_3(\gamma_{BCD} - 1)]$ | $g_1 = 0$ | $g_2 = 0$ | $g_3 = 2\alpha$ |
| $d^{-(d_2+d_4+d_5)}$ | $g_0 = exp[-\alpha\tau_1(\gamma_D - \gamma_{CD}) - \alpha\tau_2(\gamma_B + \gamma_{CD} - 2\gamma_{BCD}) - 2\alpha\tau_3(\gamma_{BCD} - 1)]$ | $g_1 = 0$ | $g_2 = 0$ | $g_3 = 2\alpha$ |
| $d^{-(d_1+d_2+d_3+d_4)}$ | $g_0 = exp[-\alpha\tau_1(\gamma_C + \gamma_D - 2\gamma_{CD}) - \alpha\tau_2(\gamma_B + 2\gamma_{CD} - 3\gamma_{BCD}) - \alpha\tau_3(\gamma_A + \gamma_{BCD} - 2)]$ | $g_1 = 2\alpha\gamma_{BCD}$ | $g_2 = 2\alpha$ | $g_3 = 0$ |
| $d^{-(d_1+d_2+d_3+d_5)}$ | $g_0 = exp[-\alpha\tau_1(\gamma_C - \gamma_{CD}) - \alpha\tau_2(\gamma_B + \gamma_{CD} - 2\gamma_{BCD}) - \alpha\tau_3(\gamma_A + 2\gamma_{BCD} - 3)]$ | $g_1 = 0$ | $g_2 = \alpha$ | $g_3 = 2\alpha$ |
| $d^{-(d_1+d_2+d_4+d_5)}$ | $g_0 = exp[-\alpha\tau_1(\gamma_D - \gamma_{CD}) - \alpha\tau_2(\gamma_B + \gamma_{CD} - 2\gamma_{BCD}) - \alpha\tau_3(\gamma_A + 2\gamma_{BCD} - 3)]$ | $g_1 = 0$ | $g_2 = \alpha$ | $g_3 = 2\alpha$ |
| $d^{-(d_1+d_3+d_4+d_5)}$ | $g_0 = exp[-\alpha\tau_1(\gamma_C + \gamma_D - 2\gamma_{CD}) - 2\alpha\tau_2(\gamma_{CD} - \gamma_{BCD}) - \alpha\tau_3(\gamma_A + \gamma_{BCD} - 2)]$ | $g_1 = \alpha\gamma_{BCD}$ | $g_2 = 0$ | $g_3 = 2\alpha$ |
| $d^{-(d_2+d_3+d_4+d_5)}$ | $g_0 = exp[-\alpha\tau_1(\gamma_C + \gamma_D - 2\gamma_{CD}) - \alpha\tau_2(\gamma_B + 2\gamma_{CD} - 3\gamma_{BCD}) - 2\alpha\tau_3(\gamma_{BCD} - 1)]$ | $g_1 = \alpha\gamma_{BCD}$ | $g_2 = 0$ | $g_3 = 2\alpha$ |
| $d^{-(d_1+d_2+d_3+d_4+d_5)}$ | $g_0 = exp[-\alpha\tau_1(\gamma_C + \gamma_D - 2\gamma_{CD}) - \alpha\tau_2(\gamma_B + 2\gamma_{CD} - 3\gamma_{BCD}) - \alpha\tau_3(\gamma_A + 2\gamma_{BCD} - 3)]$ | $g_1 = \alpha\gamma_{BCD}$ | $g_2 = \alpha$ | $g_3 = 2\alpha$ |

| | **Histories 3 and 4** | | | |
|---|---|---|---|---|
| $d^{-(d_1+d_2)}$ | $g_0 = exp[-\alpha\tau_2(\gamma_B - \gamma_{BCD}) - \alpha\tau_3(\gamma_A + \gamma_{BCD} - 2)]$ | $g_1 = 0$ | $g_2 = 2\alpha$ | $g_3 = 0$ |
| $d^{-(d_3+d_4)}$ | $g_0 = exp[-\alpha\tau_1(\gamma_C + \gamma_D - 2\gamma_{CD}) - 2\alpha\tau_2(\gamma_{CD} - \gamma_{BCD}) - 2\alpha\tau_3(\gamma_{BCD} - 1)]$ | $g_1 = 2\alpha$ | $g_2 = 0$ | $g_3 = 0$ |
| $d^{-(d_1+d_3+d_5)}$ | $g_0 = exp[-\alpha\tau_1(\gamma_C - \gamma_{CD}) - \alpha\tau_2(\gamma_{CD} - \gamma_{BCD}) - \alpha\tau_3(\gamma_A + \gamma_{BCD} - 2)]$ | $g_1 = 0$ | $g_2 = 0$ | $g_3 = 2\alpha$ |
| $d^{-(d_1+d_4+d_5)}$ | $g_0 = exp[-\alpha\tau_1(\gamma_D - \gamma_{CD}) - \alpha\tau_2(\gamma_{CD} - \gamma_{BCD}) - \alpha\tau_3(\gamma_A + \gamma_{BCD} - 2)]$ | $g_1 = 0$ | $g_2 = 0$ | $g_3 = 2\alpha$ |
| $d^{-(d_2+d_3+d_5)}$ | $g_0 = exp[-\alpha\tau_1(\gamma_C - \gamma_{CD}) - \alpha\tau_2(\gamma_B + \gamma_{CD} - 2\gamma_{BCD}) - 2\alpha\tau_3(\gamma_{BCD} - 1)]$ | $g_1 = 0$ | $g_2 = 0$ | $g_3 = 2\alpha$ |
| $d^{-(d_2+d_4+d_5)}$ | $g_0 = exp[-\alpha\tau_1(\gamma_D - \gamma_{CD}) - \alpha\tau_2(\gamma_B + \gamma_{CD} - 2\gamma_{BCD}) - 2\alpha\tau_3(\gamma_{BCD} - 1)]$ | $g_1 = 0$ | $g_2 = 0$ | $g_3 = 2\alpha$ |
| $d^{-(d_1+d_2+d_3+d_4)}$ | $g_0 = exp[-\alpha\tau_1(\gamma_C + \gamma_D - 2\gamma_{CD}) - \alpha\tau_2(\gamma_B + 2\gamma_{CD} - 3\gamma_{BCD}) - \alpha\tau_3(\gamma_A + 3\gamma_{BCD} - 4)]$ | $g_1 = 2\alpha$ | $g_2 = 2\alpha$ | $g_3 = 0$ |
| $d^{-(d_1+d_2+d_3+d_5)}$ | $g_0 = exp[-\alpha\tau_1(\gamma_C - \gamma_{CD}) - \alpha\tau_2(\gamma_B + \gamma_{CD} - 2\gamma_{BCD}) - \alpha\tau_3(\gamma_A + 2\gamma_{BCD} - 3)]$ | $g_1 = 0$ | $g_2 = \alpha$ | $g_3 = 2\alpha$ |
| $d^{-(d_1+d_2+d_4+d_5)}$ | $g_0 = exp[-\alpha\tau_1(\gamma_D - \gamma_{CD}) - \alpha\tau_2(\gamma_B + \gamma_{CD} - 2\gamma_{BCD}) - \alpha\tau_3(\gamma_A + 2\gamma_{BCD} - 3)]$ | $g_1 = 0$ | $g_2 = \alpha$ | $g_3 = 2\alpha$ |
| $d^{-(d_1+d_3+d_4+d_5)}$ | $g_0 = exp[-\alpha\tau_1(\gamma_C + \gamma_D - 2\gamma_{CD}) - 2\alpha\tau_2(\gamma_{CD} - \gamma_{BCD}) - \alpha\tau_3(\gamma_A + 2\gamma_{BCD} - 3)]$ | $g_1 = \alpha$ | $g_2 = 0$ | $g_3 = 2\alpha$ |
| $d^{-(d_2+d_3+d_4+d_5)}$ | $g_0 = exp[-\alpha\tau_1(\gamma_C + \gamma_D - 2\gamma_{CD}) - \alpha\tau_2(\gamma_B + 2\gamma_{CD} - 3\gamma_{BCD}) - 3\alpha\tau_3(\gamma_{BCD} - 1)]$ | $g_1 = \alpha$ | $g_2 = 0$ | $g_3 = 2\alpha$ |
| $d^{-(d_1+d_2+d_3+d_4+d_5)}$ | $g_0 = exp[-\alpha\tau_1(\gamma_C + \gamma_D - 2\gamma_{CD}) - \alpha\tau_2(\gamma_B + 2\gamma_{CD} - 3\gamma_{BCD}) - \alpha\tau_3(\gamma_A + 3\gamma_{BCD} - 4)]$ | $g_1 = \alpha$ | $g_2 = \alpha$ | $g_3 = 2\alpha$ |

| | **History 5** | | | |
|---|---|---|---|---|
| $d^{-(d_1+d_2)}$ | $g_0 = exp[-\alpha\tau_1(\gamma_C - \gamma_{CD}) - \alpha\tau_2(\gamma_{CD} - \gamma_{BCD}) - \alpha\tau_3(\gamma_A + \gamma_{BCD} - 2)]$ | $g_1 = 0$ | $g_2 = 2\alpha$ | $g_3 = 0$ |
| $d^{-(d_3+d_4)}$ | $g_0 = exp[-\alpha\tau_1(\gamma_D - \gamma_{CD}) - \alpha\tau_2(\gamma_B + \gamma_{CD} - 2\gamma_{BCD})]$ | $g_1 = 2\alpha\gamma_{BCD}$ | $g_2 = 0$ | $g_3 = 0$ |
| $d^{-(d_1+d_3+d_5)}$ | $g_0 = exp[-\alpha\tau_2(\gamma_B - \gamma_{BCD}) - \alpha\tau_3(\gamma_A + \gamma_{BCD} - 2)]$ | $g_1 = 0$ | $g_2 = 0$ | $g_3 = 2\alpha$ |
| $d^{-(d_1+d_4+d_5)}$ | $g_0 = exp[-\alpha\tau_1(\gamma_D - \gamma_{CD}) - \alpha\tau_2(\gamma_{CD} - \gamma_{BCD}) - \alpha\tau_3(\gamma_A + \gamma_{BCD} - 2)]$ | $g_1 = 0$ | $g_2 = 0$ | $g_3 = 2\alpha$ |
| $d^{-(d_2+d_3+d_5)}$ | $g_0 = exp[-\alpha\tau_1(\gamma_C - \gamma_{CD}) - \alpha\tau_2(\gamma_B + \gamma_{CD} - 2\gamma_{BCD}) - 2\alpha\tau_3(\gamma_{BCD} - 1)]$ | $g_1 = 0$ | $g_2 = 0$ | $g_3 = 2\alpha$ |
| $d^{-(d_2+d_4+d_5)}$ | $g_0 = exp[-\alpha\tau_1(\gamma_C + \gamma_D - 2\gamma_{CD}) - 2\alpha\tau_2(\gamma_{CD} - \gamma_{BCD}) - 2\alpha\tau_3(\gamma_{BCD} - 1)]$ | $g_1 = 0$ | $g_2 = 0$ | $g_3 = 2\alpha$ |
| $d^{-(d_1+d_2+d_3+d_4)}$ | $g_0 = exp[-\alpha\tau_1(\gamma_C + \gamma_D - 2\gamma_{CD}) - \alpha\tau_2(\gamma_B + 2\gamma_{CD} - 3\gamma_{BCD}) - \alpha\tau_3(\gamma_A + \gamma_{BCD} - 2)]$ | $g_1 = 2\alpha\gamma_{BCD}$ | $g_2 = 2\alpha$ | $g_3 = 0$ |
| $d^{-(d_1+d_2+d_3+d_5)}$ | $g_0 = exp[-\alpha\tau_1(\gamma_C - \gamma_{CD}) - \alpha\tau_2(\gamma_B + \gamma_{CD} - 2\gamma_{BCD}) - \alpha\tau_3(\gamma_A + 2\gamma_{BCD} - 3)]$ | $g_1 = 0$ | $g_2 = \alpha$ | $g_3 = 2\alpha$ |
| $d^{-(d_1+d_2+d_4+d_5)}$ | $g_0 = exp[-\alpha\tau_1(\gamma_C + \gamma_D - 2\gamma_{CD}) - 2\alpha\tau_2(\gamma_{CD} - \gamma_{BCD}) - \alpha\tau_3(\gamma_A + 2\gamma_{BCD} - 3)]$ | $g_1 = 0$ | $g_2 = \alpha$ | $g_3 = 2\alpha$ |
| $d^{-(d_1+d_3+d_4+d_5)}$ | $g_0 = exp[-\alpha\tau_1(\gamma_D - \gamma_{CD}) - \alpha\tau_2(\gamma_B + \gamma_{CD} - 2\gamma_{BCD}) - \alpha\tau_3(\gamma_A + \gamma_{BCD} - 2)]$ | $g_1 = \alpha\gamma_{BCD}$ | $g_2 = 0$ | $g_3 = 2\alpha$ |
| $d^{-(d_2+d_3+d_4+d_5)}$ | $g_0 = exp[-\alpha\tau_1(\gamma_C + \gamma_D - 2\gamma_{CD}) - \alpha\tau_2(\gamma_B + 2\gamma_{CD} - 3\gamma_{BCD}) - 2\alpha\tau_3(\gamma_{BCD} - 1)]$ | $g_1 = \alpha\gamma_{BCD}$ | $g_2 = 0$ | $g_3 = 2\alpha$ |
| $d^{-(d_1+d_2+d_3+d_4+d_5)}$ | $g_0 = exp[-\alpha\tau_1(\gamma_C + \gamma_D - 2\gamma_{CD}) - \alpha\tau_2(\gamma_B + 2\gamma_{CD} - 3\gamma_{BCD}) - \alpha\tau_3(\gamma_A + 2\gamma_{BCD} - 3)]$ | $g_1 = \alpha\gamma_{BCD}$ | $g_2 = \alpha$ | $g_3 = 2\alpha$ |

| | **Histories 6 and 7** | | | |
|---|---|---|---|---|
| $d^{-(d_1+d_2)}$ | $g_0 = exp[-\alpha\tau_1(\gamma_C - \gamma_{CD}) - \alpha\tau_2(\gamma_{CD} - \gamma_{BCD}) - \alpha\tau_3(\gamma_A + \gamma_{BCD} - 2)]$ | $g_1 = 0$ | $g_2 = 2\alpha$ | $g_3 = 0$ |
| $d^{-(d_3+d_4)}$ | $g_0 = exp[-\alpha\tau_1(\gamma_D - \gamma_{CD}) - \alpha\tau_2(\gamma_B + \gamma_{CD} - 2\gamma_{BCD}) - 2\alpha\tau_3(\gamma_{BCD} - 1)]$ | $g_1 = 2\alpha$ | $g_2 = 0$ | $g_3 = 0$ |
| $d^{-(d_1+d_3+d_5)}$ | $g_0 = exp[-\alpha\tau_2(\gamma_B - \gamma_{BCD}) - \alpha\tau_3(\gamma_A + \gamma_{BCD} - 2)]$ | $g_1 = 0$ | $g_2 = 0$ | $g_3 = 2\alpha$ |
| $d^{-(d_1+d_4+d_5)}$ | $g_0 = exp[-\alpha\tau_1(\gamma_D - \gamma_{CD}) - \alpha\tau_2(\gamma_{CD} - \gamma_{BCD}) - \alpha\tau_3(\gamma_A + \gamma_{BCD} - 2)]$ | $g_1 = 0$ | $g_2 = 0$ | $g_3 = 2\alpha$ |
| $d^{-(d_2+d_3+d_5)}$ | $g_0 = exp[-\alpha\tau_1(\gamma_C - \gamma_{CD}) - \alpha\tau_2(\gamma_B + \gamma_{CD} - 2\gamma_{BCD}) - 2\alpha\tau_3(\gamma_{BCD} - 1)]$ | $g_1 = 0$ | $g_2 = 0$ | $g_3 = 2\alpha$ |
| $d^{-(d_2+d_4+d_5)}$ | $g_0 = exp[-\alpha\tau_1(\gamma_C + \gamma_D - 2\gamma_{CD}) - 2\alpha\tau_2(\gamma_{CD} - \gamma_{BCD}) - 2\alpha\tau_3(\gamma_{BCD} - 1)]$ | $g_1 = 0$ | $g_2 = 0$ | $g_3 = 2\alpha$ |
| $d^{-(d_1+d_2+d_3+d_4)}$ | $g_0 = exp[-\alpha\tau_1(\gamma_C + \gamma_D - 2\gamma_{CD}) - \alpha\tau_2(\gamma_B + 2\gamma_{CD} - 3\gamma_{BCD}) - \alpha\tau_3(\gamma_A + 3\gamma_{BCD} - 4)]$ | $g_1 = 2\alpha$ | $g_2 = 2\alpha$ | $g_3 = 0$ |
| $d^{-(d_1+d_2+d_3+d_5)}$ | $g_0 = exp[-\alpha\tau_1(\gamma_C - \gamma_{CD}) - \alpha\tau_2(\gamma_B + \gamma_{CD} - 2\gamma_{BCD}) - \alpha\tau_3(\gamma_A + 2\gamma_{BCD} - 3)]$ | $g_1 = 0$ | $g_2 = \alpha$ | $g_3 = 2\alpha$ |
| $d^{-(d_1+d_2+d_4+d_5)}$ | $g_0 = exp[-\alpha\tau_1(\gamma_C + \gamma_D - 2\gamma_{CD}) - 2\alpha\tau_2(\gamma_{CD} - \gamma_{BCD}) - \alpha\tau_3(\gamma_A + 2\gamma_{BCD} - 3)]$ | $g_1 = 0$ | $g_2 = \alpha$ | $g_3 = 2\alpha$ |
| $d^{-(d_1+d_3+d_4+d_5)}$ | $g_0 = exp[-\alpha\tau_1(\gamma_D - \gamma_{CD}) - \alpha\tau_2(\gamma_B + \gamma_{CD} - 2\gamma_{BCD}) - \alpha\tau_3(\gamma_A + 2\gamma_{BCD} - 3)]$ | $g_1 = \alpha$ | $g_2 = 0$ | $g_3 = 2\alpha$ |
| $d^{-(d_2+d_3+d_4+d_5)}$ | $g_0 = exp[-\alpha\tau_1(\gamma_C + \gamma_D - 2\gamma_{CD}) - \alpha\tau_2(\gamma_B + 2\gamma_{CD} - 3\gamma_{BCD}) - 3\alpha\tau_3(\gamma_{BCD} - 1)]$ | $g_1 = \alpha$ | $g_2 = 0$ | $g_3 = 2\alpha$ |
| $d^{-(d_1+d_2+d_3+d_4+d_5)}$ | $g_0 = exp[-\alpha\tau_1(\gamma_C + \gamma_D - 2\gamma_{CD}) - \alpha\tau_2(\gamma_B + 2\gamma_{CD} - 3\gamma_{BCD}) - \alpha\tau_3(\gamma_A + 3\gamma_{BCD} - 4)]$ | $g_1 = \alpha$ | $g_2 = \alpha$ | $g_3 = 2\alpha$ |



| | **History 8** | | | |
|---|---|---|---|---|
| $d^{-(d_1+d_2)}$ | $g_0 = exp[-\alpha\tau_1(\gamma_D - \gamma_{CD}) - \alpha\tau_2(\gamma_{CD} - \gamma_{BCD}) - \alpha\tau_3(\gamma_A + \gamma_{BCD} - 2)]$ | $g_1 = 0$ | $g_2 = 2\alpha$ | $g_3 = 0$ |
| $d^{-(d_3+d_4)}$ | $g_0 = exp[-\alpha\tau_1(\gamma_C - \gamma_{CD}) - \alpha\tau_2(\gamma_B + \gamma_{CD} - 2\gamma_{BCD})]$ | $g_1 = 2\alpha\gamma_{BCD}$ | $g_2 = 0$ | $g_3 = 0$ |
| $d^{-(d_1+d_3+d_5)}$ | $g_0 = exp[-\alpha\tau_2(\gamma_B - \gamma_{BCD}) - \alpha\tau_3(\gamma_A + \gamma_{BCD} - 2)]$ | $g_1 = 0$ | $g_2 = 0$ | $g_3 = 2\alpha$ |
| $d^{-(d_1+d_4+d_5)}$ | $g_0 = exp[-\alpha\tau_1(\gamma_C - \gamma_{CD}) - \alpha\tau_2(\gamma_{CD} - \gamma_{BCD}) - \alpha\tau_3(\gamma_A + \gamma_{BCD} - 2)]$ | $g_1 = 0$ | $g_2 = 0$ | $g_3 = 2\alpha$ |
| $d^{-(d_2+d_3+d_5)}$ | $g_0 = exp[-\alpha\tau_1(\gamma_D - \gamma_{CD}) - \alpha\tau_2(\gamma_B + \gamma_{CD} - 2\gamma_{BCD}) - 2\alpha\tau_3(\gamma_{BCD} - 1)]$ | $g_1 = 0$ | $g_2 = 0$ | $g_3 = 2\alpha$ |
| $d^{-(d_2+d_4+d_5)}$ | $g_0 = exp[-\alpha\tau_1(\gamma_C + \gamma_D - 2\gamma_{CD}) - 2\alpha\tau_2(\gamma_{CD} - \gamma_{BCD}) - 2\alpha\tau_3(\gamma_{BCD} - 1)]$ | $g_1 = 0$ | $g_2 = 0$ | $g_3 = 2\alpha$ |
| $d^{-(d_1+d_2+d_3+d_4)}$ | $g_0 = exp[-\alpha\tau_1(\gamma_C + \gamma_D - 2\gamma_{CD}) - \alpha\tau_2(\gamma_B + 2\gamma_{CD} - 3\gamma_{BCD}) - \alpha\tau_3(\gamma_A + \gamma_{BCD} - 2)]$ | $g_1 = 2\alpha\gamma_{BCD}$ | $g_2 = 2\alpha$ | $g_3 = 0$ |
| $d^{-(d_1+d_2+d_3+d_5)}$ | $g_0 = exp[-\alpha\tau_1(\gamma_D - \gamma_{CD}) - \alpha\tau_2(\gamma_B + \gamma_{CD} - 2\gamma_{BCD}) - \alpha\tau_3(\gamma_A + 2\gamma_{BCD} - 3)]$ | $g_1 = 0$ | $g_2 = \alpha$ | $g_3 = 2\alpha$ |
| $d^{-(d_1+d_2+d_4+d_5)}$ | $g_0 = exp[-\alpha\tau_1(\gamma_C + \gamma_D - 2\gamma_{CD}) - 2\alpha\tau_2(\gamma_{CD} - \gamma_{BCD}) - \alpha\tau_3(\gamma_A + 2\gamma_{BCD} - 3)]$ | $g_1 = 0$ | $g_2 = \alpha$ | $g_3 = 2\alpha$ |
| $d^{-(d_1+d_3+d_4+d_5)}$ | $g_0 = exp[-\alpha\tau_1(\gamma_C - \gamma_{CD}) - \alpha\tau_2(\gamma_B + \gamma_{CD} - 2\gamma_{BCD}) - \alpha\tau_3(\gamma_A + \gamma_{BCD} - 2)]$ | $g_1 = \alpha\gamma_{BCD}$ | $g_2 = 0$ | $g_3 = 2\alpha$ |
| $d^{-(d_2+d_3+d_4+d_5)}$ | $g_0 = exp[-\alpha\tau_1(\gamma_C + \gamma_D - 2\gamma_{CD}) - \alpha\tau_2(\gamma_B + 2\gamma_{CD} - 3\gamma_{BCD}) - 2\alpha\tau_3(\gamma_{BCD} - 1)]$ | $g_1 = \alpha\gamma_{BCD}$ | $g_2 = 0$ | $g_3 = 2\alpha$ |
| $d^{-(d_1+d_2+d_3+d_4+d_5)}$ | $g_0 = exp[-\alpha\tau_1(\gamma_C + \gamma_D - 2\gamma_{CD}) - \alpha\tau_2(\gamma_B + 2\gamma_{CD} - 3\gamma_{BCD}) - \alpha\tau_3(\gamma_A + 2\gamma_{BCD} - 2)]$ | $g_1 = \alpha\gamma_{BCD}$ | $g_2 = \alpha$ | $g_3 = 2\alpha$ |

| | **Histories 9 and 10** | | | |
|---|---|---|---|---|
| $d^{-(d_1+d_2)}$ | $g_0 = exp[-\alpha\tau_1(\gamma_D - \gamma_{CD}) - \alpha\tau_2(\gamma_{CD} - \gamma_{BCD}) - \alpha\tau_3(\gamma_A + \gamma_{BCD} - 2)]$ | $g_1 = 0$ | $g_2 = 2\alpha$ | $g_3 = 0$ |
| $d^{-(d_3+d_4)}$ | $g_0 = exp[-\alpha\tau_1(\gamma_C - \gamma_{CD}) - \alpha\tau 2(\gamma_B + \gamma_{CD} - 2\gamma_{BCD}) - 2\alpha\tau_3(\gamma_{BCD} - 1)]$ | $g_1 = 2\alpha$ | $g_2 = 0$ | $g_3 = 0$ |
| $d^{-(d_1+d_3+d_5)}$ | $g_0 = exp[-\alpha\tau_2(\gamma_B - \gamma_{BCD}) - \alpha\tau_3(\gamma_A + \gamma_{BCD} - 2)]$ | $g_1 = 0$ | $g_2 = 0$ | $g_3 = 2\alpha$ |
| $d^{-(d_1+d_4+d_5)}$ | $g_0 = exp[-\alpha\tau_1(\gamma_C - \gamma_{CD}) - \alpha\tau_2(\gamma_{CD} - \gamma_{BCD}) - \alpha\tau_3(\gamma_A + \gamma_{BCD} - 2)]$ | $g_1 = 0$ | $g_2 = 0$ | $g_3 = 2\alpha$ |
| $d^{-(d_2+d_3+d_5)}$ | $g_0 = exp[-\alpha\tau_1(\gamma_D - \gamma_{CD}) - \alpha\tau_2(\gamma_B + \gamma_{CD} - 2\gamma_{BCD}) - 2\alpha\tau_3(\gamma_{BCD} - 1)]$ | $g_1 = 0$ | $g_2 = 0$ | $g_3 = 2\alpha$ |
| $d^{-(d_2+d_4+d_5)}$ | $g_0 = exp[-\alpha\tau_1(\gamma_C + \gamma_D - 2\gamma_{CD}) - 2\alpha\tau_2(\gamma_{CD} - \gamma_{BCD}) - 2\alpha\tau_3(\gamma_{BCD} - 1)]$ | $g_1 = 0$ | $g_2 = 0$ | $g_3 = 2\alpha$ |
| $d^{-(d_1+d_2+d_3+d_4)}$ | $g_0 = exp[-\alpha\tau_1(\gamma_C + \gamma_D - 2\gamma_{CD}) - \alpha\tau_2(\gamma_B + 2\gamma_{CD} - 3\gamma_{BCD}) - \alpha\tau_3(\gamma_A + 3\gamma_{BCD} - 4)]$ | $g_1 = 2\alpha$ | $g_2 = 2\alpha$ | $g_3 = 0$ |
| $d^{-(d_1+d_2+d_3+d_5)}$ | $g_0 = exp[-\alpha\tau_1(\gamma_D - \gamma_{CD}) - \alpha\tau_2(\gamma_B + \gamma_{CD} - 2\gamma_{BCD}) - \alpha\tau_3(\gamma_A + 2\gamma_{BCD} - 3)]$ | $g_1 = 0$ | $g_2 = \alpha$ | $g_3 = 2\alpha$ |
| $d^{-(d_1+d_2+d_4+d_5)}$ | $g_0 = exp[-\alpha\tau_1(\gamma_C + \gamma_D - 2\gamma_{CD}) - 2\alpha\tau_2(\gamma_{CD} - \gamma_{BCD}) - \alpha\tau_3(\gamma_A + 2\gamma_{BCD} - 3)]$ | $g_1 = 0$ | $g_2 = \alpha$ | $g_3 = 2\alpha$ |
| $d^{-(d_1+d_3+d_4+d_5)}$ | $g_0 = exp[-\alpha\tau_1(\gamma_C - \gamma_{CD}) - \alpha\tau_2(\gamma_B + \gamma_{CD} - 2\gamma_{BCD}) - \alpha\tau_3(\gamma_A + 2\gamma_{BCD} - 3)]$ | $g_1 = \alpha$ | $g_2 = 0$ | $g_3 = 2\alpha$ |
| $d^{-(d_2+d_3+d_4+d_5)}$ | $g_0 = exp[-\alpha\tau_1(\gamma_C + \gamma_D - 2\gamma_{CD}) - \alpha\tau_2(\gamma_B + 2\gamma_{CD} - 3\gamma_{BCD}) - 3\alpha\tau_3(\gamma_{BCD} - 1)]$ | $g_1 = \alpha$ | $g_2 = 0$ | $g_3 = 2\alpha$ |
| $d^{-(d_1+d_2+d_3+d_4+d_5)}$ | $g_0 = exp[-\alpha\tau_1(\gamma_C + \gamma_D - 2\gamma_{CD}) - \alpha\tau_2(\gamma_B + 2\gamma_{CD} - 3\gamma_{BCD}) - \alpha\tau_3(\gamma_A + 3\gamma_{BCD} - 4)]$ | $g_1 = \alpha$ | $g_2 = \alpha$ | $g_3 = 2\alpha$ |

| | **History 11** | | | |
|---|---|---|---|---|
| $d^{-(d_1+d_2)}$ | $g_0 = exp[-\alpha\tau_2(\gamma_B - \gamma_{BCD}) - \alpha\tau_3(\gamma_A + \gamma_{BCD} - 2)]$ | $g_1 = 0$ | $g_2 = 0$ | $g_3 = 2\alpha$ |
| $d^{-(d_3+d_4)}$ | $g_0 = exp[-\alpha\tau_1(\gamma_C + \gamma_D - 2\gamma_{CD})]$ | $g_1 = 2\alpha\gamma_{CD}$ | $g_2 = 0$ | $g_3 = 0$ |
| $d^{-(d_1+d_3+d_5)}$ | $g_0 = exp[-\alpha\tau_1(\gamma_C - \gamma_{CD}) - \alpha\tau_2(\gamma_{CD} - \gamma_{BCD}) - \alpha\tau_3(\gamma_A + \gamma_{BCD} - 2)]$ | $g_1 = 0$ | $g_2 = 0$ | $g_3 = 2\alpha$ |
| $d^{-(d_1+d_4+d_5)}$ | $g_0 = exp[-\alpha\tau_1(\gamma_D - \gamma_{CD}) - \alpha\tau_2(\gamma_{CD} - \gamma_{BCD}) - \alpha\tau_3(\gamma_A + \gamma_{BCD} - 2)]$ | $g_1 = 0$ | $g_2 = 0$ | $g_3 = 2\alpha$ |
| $d^{-(d_2+d_3+d_5)}$ | $g_0 = exp[-\alpha\tau_1(\gamma_C - \gamma_{CD}) - \alpha\tau_2(\gamma_B + \gamma_{CD} - 2\gamma_{BCD})]$ | $g_1 = 0$ | $g_2 = 2\alpha\gamma_{BCD}$ | $g_3 = 0$ |
| $d^{-(d_2+d_4+d_5)}$ | $g_0 = exp[-\alpha\tau_1(\gamma_D - \gamma_{CD}) - \alpha\tau_2(\gamma_B + \gamma_{CD} - 2\gamma_{BCD})]$ | $g_1 = 0$ | $g_2 = 2\alpha\gamma_{BCD}$ | $g_3 = 0$ |
| $d^{-(d_1+d_2+d_3+d_4)}$ | $g_0 = exp[-\alpha\tau_1(\gamma_C + \gamma_D - 2\gamma_{CD}) - \alpha\tau_2(\gamma_B - \gamma_{BCD}) - \alpha\tau_3(\gamma_A + \gamma_{BCD} - 2)]$ | $g_1 = 2\alpha\gamma_{CD}$ | $g_2 = 0$ | $g_3 = 2\alpha$ |
| $d^{-(d_1+d_2+d_3+d_5)}$ | $g_0 = exp[-\alpha\tau_1(\gamma_C - \gamma_{CD}) - \alpha\tau_2(\gamma_B + \gamma_{CD} - 2\gamma_{BCD}) - \alpha\tau_3(\gamma_A + \gamma_{BCD} - 2)]$ | $g_1 = 0$ | $g_2 = \alpha\gamma_{BCD}$ | $g_3 = 2\alpha$ |
| $d^{-(d_1+d_2+d_4+d_5)}$ | $g_0 = exp[-\alpha\tau_1(\gamma_D - \gamma_{CD}) - \alpha\tau_2(\gamma_B + \gamma_{CD} - 2\gamma_{BCD}) - \alpha\tau_3(\gamma_A + \gamma_{BCD} - 2)]$ | $g_1 = 0$ | $g_2 = \alpha\gamma_{BCD}$ | $g_3 = 2\alpha$ |
| $d^{-(d_1+d_3+d_4+d_5)}$ | $g_0 = exp[-\alpha\tau_1(\gamma_C + \gamma_D - 2\gamma_{CD}) - \alpha\tau_2(\gamma_{CD} - \gamma_{BCD}) - \alpha\tau_3(\gamma_A + \gamma_{BCD} - 2)]$ | $g_1 = \alpha\gamma_{CD}$ | $g_2 = 0$ | $g_3 = 2\alpha$ |
| $d^{-(d_2+d_3+d_4+d_5)}$ | $g_0 = exp[-\alpha\tau_1(\gamma_C + \gamma_D - 2\gamma_{CD}) - \alpha\tau_2(\gamma_B + \gamma_{CD} - 2\gamma_{BCD})]$ | $g_1 = \alpha\gamma_{CD}$ | $g_2 = 2\alpha\gamma_{BCD}$ | $g_3 = 0$ |
| $d^{-(d_1+d_2+d_3+d_4+d_5)}$ | $g_0 = exp[-\alpha\tau_1(\gamma_C + \gamma_D - 2\gamma_{CD}) - \alpha\tau_2(\gamma_B + \gamma_{CD} - 2\gamma_{BCD}) - \alpha\tau_3(\gamma_A + \gamma_{BCD} - 2)]$ | $g_1 = \alpha\gamma_{CD}$ | $g_2 = \alpha\gamma_{BCD}$ | $g_3 = 2\alpha$ |

| | **History 12** | | | |
|---|---|---|---|---|
| $d^{-(d_1+d_2)}$ | $g_0 = exp[-\alpha\tau_2(\gamma_B - \gamma_{BCD}) - \alpha\tau_3(\gamma_A + \gamma_{BCD} - 2)]$ | $g_1 = 0$ | $g_2 = 0$ | $g_3 = 2\alpha$ |
| $d^{-(d_3+d_4)}$ | $g_0 = exp[-\alpha\tau_1(\gamma_C + \gamma_D - 2\gamma_{CD})]$ | $g_1 = 2\alpha\gamma_{CD}$ | $g_2 = 0$ | $g_3 = 0$ |
| $d^{-(d_1+d_3+d_5)}$ | $g_0 = exp[-\alpha\tau_1(\gamma_C - \gamma_{CD}) - \alpha\tau_2(\gamma_{CD} - \gamma_{BCD}) - \alpha\tau_3(\gamma_A + \gamma_{BCD} - 2)]$ | $g_1 = 0$ | $g_2 = 0$ | $g_3 = 2\alpha$ |
| $d^{-(d_1+d_4+d_5)}$ | $g_0 = exp[-\alpha\tau_1(\gamma_D - \gamma_{CD}) - \alpha\tau_2(\gamma_{CD} - \gamma_{BCD}) - \alpha\tau_3(\gamma_A + \gamma_{BCD} - 2)]$ | $g_1 = 0$ | $g_2 = 0$ | $g_3 = 2\alpha$ |
| $d^{-(d_2+d_3+d_5)}$ | $g_0 = exp[-\alpha\tau_1(\gamma_C - \gamma_{CD}) - \alpha\tau_2(\gamma_B + \gamma_{CD} - 2\gamma_{BCD}) - 2\alpha\tau_3(\gamma_{BCD} - 1)]$ | $g_1 = 0$ | $g_2 = 2\alpha$ | $g_3 = 0$ |
| $d^{-(d_2+d_4+d_5)}$ | $g_0 = exp[-\alpha\tau_1(\gamma_D - \gamma_{CD}) - \alpha\tau_2(\gamma_B + \gamma_{CD} - 2\gamma_{BCD}) - 2\alpha\tau_3(\gamma_{BCD} - 1)]$ | $g_1 = 0$ | $g_2 = 2\alpha$ | $g_3 = 0$ |
| $d^{-(d_1+d_2+d_3+d_4)}$ | $g_0 = exp[-\alpha\tau_1(\gamma_C + \gamma_D - 2\gamma_{CD}) - \alpha\tau_2(\gamma_B - \gamma_{BCD}) - \alpha\tau_3(\gamma_A + \gamma_{BCD} - 2)]$ | $g_1 = 2\alpha\gamma_{CD}$ | $g_2 = 0$ | $g_3 = 2\alpha$ |
| $d^{-(d_1+d_2+d_3+d_5)}$ | $g_0 = exp[-\alpha\tau_1(\gamma_C - \gamma_{CD}) - \alpha\tau_2(\gamma_B + \gamma_{CD} - 2\gamma_{BCD}) - \alpha\tau_3(\gamma_A + 2\gamma_{BCD} - 3)]$ | $g_1 = 0$ | $g_2 = \alpha$ | $g_3 = 2\alpha$ |
| $d^{-(d_1+d_2+d_4+d_5)}$ | $g_0 = exp[-\alpha\tau_1(\gamma_D - \gamma_{CD}) - \alpha\tau_2(\gamma_B + \gamma_{CD} - 2\gamma_{BCD}) - \alpha\tau_3(\gamma_A + 2\gamma_{BCD} - 3)]$ | $g_1 = 0$ | $g_2 = \alpha$ | $g_3 = 2\alpha$ |
| $d^{-(d_1+d_3+d_4+d_5)}$ | $g_0 = exp[-\alpha\tau_1(\gamma_C + \gamma_D - 2\gamma_{CD}) - \alpha\tau_2(\gamma_{CD} - \gamma_{BCD}) - \alpha\tau_3(\gamma_A + \gamma_{BCD} - 2)]$ | $g_1 = \alpha\gamma_{CD}$ | $g_2 = 0$ | $g_3 = 2\alpha$ |
| $d^{-(d_2+d_3+d_4+d_5)}$ | $g_0 = exp[-\alpha\tau_1(\gamma_C + \gamma_D - 2\gamma_{CD}) - \alpha\tau_2(\gamma_B + \gamma_{CD} - 2\gamma_{BCD}) - 2\alpha\tau_3(\gamma_{BCD} - 1)]$ | $g_1 = \alpha\gamma_{CD}$ | $g_2 = 2\alpha$ | $g_3 = 0$ |
| $d^{-(d_1+d_2+d_3+d_4+d_5)}$ | $g_0 = exp[-\alpha\tau_1(\gamma_C + \gamma_D - 2\gamma_{CD}) - \alpha\tau_2(\gamma_B + \gamma_{CD} - 2\gamma_{BCD}) - \alpha\tau_3(\gamma_A + 2\gamma_{BCD} - 3)]$ | $g_1 = \alpha\gamma_{CD}$ | $g_2 = \alpha$ | $g_3 = 2\alpha$ |

| | **History 13** | | | |
|---|---|---|---|---|
| $d^{-(d_1+d_2)}$ | $g_0 = exp[-\alpha\tau_2(\gamma_B - \gamma_{BCD}) - \alpha\tau_3(\gamma_A + \gamma_{BCD} - 2)]$ | $g_1 = 0$ | $g_2 = 0$ | $g_3 = 2\alpha$ |
| $d^{-(d_3+d_4)}$ | $g_0 = exp[-\alpha\tau_1(\gamma_C + \gamma_D - 2\gamma_{CD}) - 2\alpha\tau_2(\gamma_{CD} - \gamma_{BCD})]$ | $g_1 = 2\alpha\gamma_{BCD}$ | $g_2 = 0$ | $g_3 = 0$ |
| $d^{-(d_1+d_3+d_5)}$ | $g_0 = exp[-\alpha\tau_1(\gamma_C - \gamma_{CD}) - \alpha\tau_2(\gamma_{CD} - \gamma_{BCD}) - \alpha\tau_3(\gamma_A + \gamma_{BCD} - 2)]$ | $g_1 = 0$ | $g_2 = 0$ | $g_3 = 2\alpha$ |
| $d^{-(d_1+d_4+d_5)}$ | $g_0 = exp[-\alpha\tau_1(\gamma_D - \gamma_{CD}) - \alpha\tau_2(\gamma_{CD} - \gamma_{BCD}) - \alpha\tau_3(\gamma_A + \gamma_{BCD} - 2)]$ | $g_1 = 0$ | $g_2 = 0$ | $g_3 = 2\alpha$ |
| $d^{-(d_2+d_3+d_5)}$ | $g_0 = exp[-\alpha\tau_1(\gamma_C - \gamma_{CD}) - \alpha\tau_2(\gamma_B + \gamma_{CD} - 2\gamma_{BCD})]$ | $g_1 = 0$ | $g_2 = 2\alpha\gamma_{BCD}$ | $g_3 = 0$ |
| $d^{-(d_2+d_4+d_5)}$ | $g_0 = exp[-\alpha\tau_1(\gamma_D - \gamma_{CD}) - \alpha\tau_2(\gamma_B + \gamma_{CD} - 2\gamma_{BCD})]$ | $g_1 = 0$ | $g_2 = 2\alpha\gamma_{BCD}$ | $g_3 = 0$ |
| $d^{-(d_1+d_2+d_3+d_4)}$ | $g_0 = exp[-\alpha\tau_1(\gamma_C + \gamma_D - 2\gamma_{CD}) - \alpha\tau_2(\gamma_B + 2\gamma_{CD} - 3\gamma_{BCD}) - \alpha\tau_3(\gamma_A + \gamma_{BCD} - 2)]$ | $g_1 = 2\alpha\gamma_{BCD}$ | $g_2 = 0$ | $g_3 = 2\alpha$ |
| $d^{-(d_1+d_2+d_3+d_5)}$ | $g_0 = exp[-\alpha\tau_1(\gamma_C - \gamma_{CD}) - \alpha\tau_2(\gamma_B + \gamma_{CD} - 2\gamma_{BCD}) - \alpha\tau_3(\gamma_A + \gamma_{BCD} - 2)]$ | $g_1 = 0$ | $g_2 = \alpha\gamma_{BCD}$ | $g_3 = 2\alpha$ |
| $d^{-(d_1+d_2+d_4+d_5)}$ | $g_0 = exp[-\alpha\tau_1(\gamma_D - \gamma_{CD}) - \alpha\tau_2(\gamma_B + \gamma_{CD} - 2\gamma_{BCD}) - \alpha\tau_3(\gamma_A + \gamma_{BCD} - 2)]$ | $g_1 = 0$ | $g_2 = \alpha\gamma_{BCD}$ | $g_3 = 2\alpha$ |
| $d^{-(d_1+d_3+d_4+d_5)}$ | $g_0 = exp[-\alpha\tau_1(\gamma_C + \gamma_D - 2\gamma_{CD}) - 2\alpha\tau_2(\gamma_{CD} - \gamma_{BCD}) - \alpha\tau_3(\gamma_A + \gamma_{BCD} - 2)]$ | $g_1 = \alpha\gamma_{BCD}$ | $g_2 = 0$ | $g_3 = 2\alpha$ |
| $d^{-(d_2+d_3+d_4+d_5)}$ | $g_0 = exp[-\alpha\tau_1(\gamma_C + \gamma_D - 2\gamma_{CD}) - \alpha\tau_2(\gamma_B + 2\gamma_{CD} - 3\gamma_{BCD})]$ | $g_1 = \alpha\gamma_{BCD}$ | $g_2 = 2\alpha\gamma_{BCD}$ | $g_3 = 0$ |
| $d^{-(d_1+d_2+d_3+d_4+d_5)}$ | $g_0 = exp[-\alpha\tau_1(\gamma_C + \gamma_D - 2\gamma_{CD}) - \alpha\tau_2(\gamma_B + 2\gamma_{CD} - 3\gamma_{BCD}) - \alpha\tau_3(\gamma_A + \gamma_{BCD} - 2)]$ | $g_1 = \alpha\gamma_{BCD}$ | $g_2 = \alpha\gamma_{BCD}$ | $g_3 = 2\alpha$ |

| | **History 14** | | | |
|---|---|---|---|---|
| $d^{-(d_1+d_2)}$ | $g_0 = exp[-\alpha\tau_2(\gamma_B - \gamma_{BCD}) - \alpha\tau_3(\gamma_A + \gamma_{BCD} - 2)]$ | $g_1 = 0$ | $g_2 = 0$ | $g_3 = 2\alpha$ |
| $d^{-(d_3+d_4)}$ | $g_0 = exp[-\alpha\tau_1(\gamma_C + \gamma_D - 2\gamma_{CD}) - 2\alpha\tau_2(\gamma_{CD} - \gamma_{BCD})]$ | $g_1 = 2\alpha\gamma_{BCD}$ | $g_2 = 0$ | $g_3 = 0$ |
| $d^{-(d_1+d_3+d_5)}$ | $g_0 = exp[-\alpha\tau_1(\gamma_C - \gamma_{CD}) - \alpha\tau_2(\gamma_{CD} - \gamma_{BCD}) - \alpha\tau_3(\gamma_A + \gamma_{BCD} - 2)]$ | $g_1 = 0$ | $g_2 = 0$ | $g_3 = 2\alpha$ |
| $d^{-(d_1+d_4+d_5)}$ | $g_0 = exp[-\alpha\tau_1(\gamma_D - \gamma_{CD}) - \alpha\tau_2(\gamma_{CD} - \gamma_{BCD}) - \alpha\tau_3(\gamma_A + \gamma_{BCD} - 2)]$ | $g_1 = 0$ | $g_2 = 0$ | $g_3 = 2\alpha$ |
| $d^{-(d_2+d_3+d_5)}$ | $g_0 = exp[-\alpha\tau_1(\gamma_C - \gamma_{CD}) - \alpha\tau_2(\gamma_B + \gamma_{CD} - 2\gamma_{BCD}) - 2\alpha\tau_3(\gamma_{BCD} - 1)]$ | $g_1 = 0$ | $g_2 = 2\alpha$ | $g_3 = 0$ |
| $d^{-(d_2+d_4+d_5)}$ | $g_0 = exp[-\alpha\tau_1(\gamma_D - \gamma_{CD}) - \alpha\tau_2(\gamma_B + \gamma_{CD} - 2\gamma_{BCD}) - 2\alpha\tau_3(\gamma_{BCD} - 1)]$ | $g_1 = 0$ | $g_2 = 2\alpha$ | $g_3 = 0$ |
| $d^{-(d_1+d_2+d_3+d_4)}$ | $g_0 = exp[-\alpha\tau_1(\gamma_C + \gamma_D - 2\gamma_{CD}) - \alpha\tau_2(\gamma_B + 2\gamma_{CD} - 3\gamma_{BCD}) - \alpha\tau_3(\gamma_A + \gamma_{BCD} - 2)]$ | $g_1 = 2\alpha\gamma_{BCD}$ | $g_2 = 0$ | $g_3 = 2\alpha$ |
| $d^{-(d_1+d_2+d_3+d_5)}$ | $g_0 = exp[-\alpha\tau_1(\gamma_C - \gamma_{CD}) - \alpha\tau_2(\gamma_B + \gamma_{CD} - 2\gamma_{BCD}) - \alpha\tau_3(\gamma_A + 2\gamma_{BCD} - 3)]$ | $g_1 = 0$ | $g_2 = \alpha$ | $g_3 = 2\alpha$ |
| $d^{-(d_1+d_2+d_4+d_5)}$ | $g_0 = exp[-\alpha\tau_1(\gamma_D - \gamma_{CD}) - \alpha\tau_2(\gamma_B + \gamma_{CD} - 2\gamma_{BCD}) - \alpha\tau_3(\gamma_A + 2\gamma_{BCD} - 3)]$ | $g_1 = 0$ | $g_2 = \alpha$ | $g_3 = 2\alpha$ |
| $d^{-(d_1+d_3+d_4+d_5)}$ | $g_0 = exp[-\alpha\tau_1(\gamma_C + \gamma_D - 2\gamma_{CD}) - 2\alpha\tau_2(\gamma_{CD} - \gamma_{BCD}) - \alpha\tau_3(\gamma_A + \gamma_{BCD} - 2)]$ | $g_1 = \alpha\gamma_{BCD}$ | $g_2 = 0$ | $g_3 = 2\alpha$ |
| $d^{-(d_2+d_3+d_4+d_5)}$ | $g_0 = exp[-\alpha\tau_1(\gamma_C + \gamma_D - 2\gamma_{CD}) - \alpha\tau_2(\gamma_B + 2\gamma_{CD} - 3\gamma_{BCD}) - 2\alpha\tau_3(\gamma_{BCD} - 1)]$ | $g_1 = \alpha\gamma_{BCD}$ | $g_2 = 2\alpha$ | $g_3 = 0$ |
| $d^{-(d_1+d_2+d_3+d_4+d_5)}$ | $g_0 = exp[-\alpha\tau_1(\gamma_C + \gamma_D - 2\gamma_{CD}) - \alpha\tau_2(\gamma_B + 2\gamma_{CD} - 3\gamma_{BCD}) - \alpha\tau_3(\gamma_A + 2\gamma_{BCD} - 3)]$ | $g_1 = \alpha\gamma_{BCD}$ | $g_2 = \alpha$ | $g_3 = 2\alpha$ |

| | **History 15** | | | |
|---|---|---|---|---|
| $d^{-(d_1+d_2)}$ | $g_0 = exp[-\alpha\tau_2(\gamma_B - \gamma_{BCD}) - \alpha\tau_3(\gamma_A + \gamma_{BCD} - 2)]$ | $g_1 = 0$ | $g_2 = 0$ | $g_3 = 2\alpha$ |
| $d^{-(d_3+d_4)}$ | $g_0 = exp[-\alpha\tau_1(\gamma_C + \gamma_D - 2\gamma_{CD}) - 2\alpha\tau_2(\gamma_{CD} - \gamma_{BCD}) - 2\alpha\tau_3[\gamma_{BCD} - 1]$ | $g_1 = 2\alpha$ | $g_2 = 0$ | $g_3 = 0$ |
| $d^{-(d_1+d_3+d_5)}$ | $g_0 = exp[-\alpha\tau_1(\gamma_C - \gamma_{CD}) - \alpha\tau_2(\gamma_{CD} - \gamma_{BCD}) - \alpha\tau_3(\gamma_A + \gamma_{BCD} - 2)]$ | $g_1 = 0$ | $g_2 = 0$ | $g_3 = 2\alpha$ |
| $d^{-(d_1+d_4+d_5)}$ | $g_0 = exp[-\alpha\tau_1(\gamma_D - \gamma_{CD}) - \alpha\tau_2(\gamma_{CD} - \gamma_{BCD}) - \alpha\tau_3(\gamma_A + \gamma_{BCD} - 2)]$ | $g_1 = 0$ | $g_2 = 0$ | $g_3 = 2\alpha$ |
| $d^{-(d_2+d_3+d_5)}$ | $g_0 = exp[-\alpha\tau_1(\gamma_C - \gamma_{CD}) - \alpha\tau_2(\gamma_B + \gamma_{CD} - 2\gamma_{BCD}) - 2\alpha\tau_3(\gamma_{BCD} - 1)]$ | $g_1 = 0$ | $g_2 = 2\alpha$ | $g_3 = 0$ |
| $d^{-(d_2+d_4+d_5)}$ | $g_0 = exp[-\alpha\tau_1(\gamma_D - \gamma_{CD}) - \alpha\tau_2(\gamma_B + \gamma_{CD} - 2\gamma_{BCD}) - 2\alpha\tau_3(\gamma_{BCD} - 1)]$ | $g_1 = 0$ | $g_2 = 2\alpha$ | $g_3 = 0$ |
| $d^{-(d_1+d_2+d_3+d_4)}$ | $g_0 = exp[-\alpha\tau_1(\gamma_C + \gamma_D - 2\gamma_{CD}) - \alpha\tau_2(\gamma_B + 2\gamma_{CD} - 3\gamma_{BCD}) - \alpha\tau_3(\gamma_A + 3\gamma_{BCD} - 4)]$ | $g_1 = 2\alpha$ | $g_2 = 0$ | $g_3 = 2\alpha$ |
| $d^{-(d_1+d_2+d_3+d_5)}$ | $g_0 = exp[-\alpha\tau_1(\gamma_C - \gamma_{CD}) - \alpha\tau_2(\gamma_B + \gamma_{CD} - 2\gamma_{BCD}) - \alpha\tau_3(\gamma_A + 2\gamma_{BCD} - 3)]$ | $g_1 = 0$ | $g_2 = \alpha$ | $g_3 = 2\alpha$ |
| $d^{-(d_1+d_2+d_4+d_5)}$ | $g_0 = exp[-\alpha\tau_1(\gamma_D - \gamma_{CD}) - \alpha\tau_2(\gamma_B + \gamma_{CD} - 2\gamma_{BCD}) - \alpha\tau_3(\gamma_A + 2\gamma_{BCD} - 3)]$ | $g_1 = 0$ | $g_2 = \alpha$ | $g_3 = 2\alpha$ |
| $d^{-(d_1+d_3+d_4+d_5)}$ | $g_0 = exp[-\alpha\tau_1(\gamma_C + \gamma_D - 2\gamma_{CD}) - 2\alpha\tau_2(\gamma_{CD} - \gamma_{BCD}) - \alpha\tau_3(\gamma_A + 2\gamma_{BCD} - 3)]$ | $g_1 = \alpha$ | $g_2 = 0$ | $g_3 = 2\alpha$ |
| $d^{-(d_2+d_3+d_4+d_5)}$ | $g_0 = exp[-\alpha\tau_1(\gamma_C + \gamma_D - 2\gamma_{CD}) - \alpha\tau_2(\gamma_B + 2\gamma_{CD} - 3\gamma_{BCD}) - 3\alpha\tau_3(\gamma_{BCD} - 1)]$ | $g_1 = \alpha$ | $g_2 = 2\alpha$ | $g_3 = 0$ |
| $d^{-(d_1+d_2+d_3+d_4+d_5)}$ | $g_0 = exp[-\alpha\tau_1(\gamma_C + \gamma_D - 2\gamma_{CD}) - \alpha\tau_2(\gamma_B + 2\gamma_{CD} - 3\gamma_{BCD}) - \alpha\tau_3(\gamma_A + 3\gamma_{BCD} - 4)]$ | $g_1 = \alpha$ | $g_2 = \alpha$ | $g_3 = 2\alpha$ |

| | **History 16** | | | |
|---|---|---|---|---|
| $d^{-(d_1+d_2)}$ | $g_0 = exp[-\alpha\tau_1(\gamma_C - \gamma_{CD}) - \alpha\tau_2(\gamma_{CD} - \gamma_{BCD}) - \alpha\tau_3(\gamma_A + \gamma_{BCD} - 2)]$ | $g_1 = 0$ | $g_2 = 0$ | $g_3 = 2\alpha$ |
| $d^{-(d_3+d_4)}$ | $g_0 = exp[-\alpha\tau_1(\gamma_D - \gamma_{CD}) - \alpha\tau_2(\gamma_B + \gamma_{CD} - 2\gamma_{BCD})]$ | $g_1 = 2\alpha\gamma_{BCD}$ | $g_2 = 0$ | $g_3 = 0$ |
| $d^{-(d_1+d_3+d_5)}$ | $g_0 = exp[-\alpha\tau_2(\gamma_B - \gamma_{BCD}) - \alpha\tau_3(\gamma_A + \gamma_{BCD} - 2)]$ | $g_1 = 0$ | $g_2 = 0$ | $g_3 = 2\alpha$ |
| $d^{-(d_1+d_4+d_5)}$ | $g_0 = exp[-\alpha\tau_1(\gamma_D - \gamma_{CD}) - \alpha\tau_2(\gamma_{CD} - \gamma_{BCD}) - \alpha\tau_3(\gamma_A + \gamma_{BCD} - 2)]$ | $g_1 = 0$ | $g_2 = 0$ | $g_3 = 2\alpha$ |
| $d^{-(d_2+d_3+d_5)}$ | $g_0 = exp[-\alpha\tau_1(\gamma_C - \gamma_{CD}) - \alpha\tau_2(\gamma_B + \gamma_{CD} - 2\gamma_{BCD})]$ | $g_1 = 0$ | $g_2 = 2\alpha\gamma_{BCD}$ | $g_3 = 0$ |
| $d^{-(d_2+d_4+d_5)}$ | $g_0 = exp[-\alpha\tau_1(\gamma_C + \gamma_D - 2\gamma_{CD}) - 2\alpha\tau_2(\gamma_{CD} - \gamma_{BCD})]$ | $g_1 = 0$ | $g_2 = 2\alpha\gamma_{BCD}$ | $g_3 = 0$ |
| $d^{-(d_1+d_2+d_3+d_4)}$ | $g_0 = exp[-\alpha\tau_1(\gamma_C + \gamma_D - 2\gamma_{CD}) - \alpha\tau_2(\gamma_B + 2\gamma_{CD} - 3\gamma_{BCD}) - \alpha\tau_3(\gamma_A + \gamma_{BCD} - 2)]$ | $g_1 = 2\alpha\gamma_{BCD}$ | $g_2 = 0$ | $g_3 = 2\alpha$ |
| $d^{-(d_1+d_2+d_3+d_5)}$ | $g_0 = exp[-\alpha\tau_1(\gamma_C - \gamma_{CD}) - \alpha\tau_2(\gamma_B + \gamma_{CD} - 2\gamma_{BCD}) - \alpha\tau_3(\gamma_A + \gamma_{BCD} - 2)]$ | $g_1 = 0$ | $g_2 = \alpha\gamma_{BCD}$ | $g_3 = 2\alpha$ |
| $d^{-(d_1+d_2+d_4+d_5)}$ | $g_0 = exp[-\alpha\tau_1(\gamma_C + \gamma_D - 2\gamma_{CD}) - 2\alpha\tau_2(\gamma_{CD} - \gamma_{BCD}) - \alpha\tau_3(\gamma_A + \gamma_{BCD} - 2)]$ | $g_1 = 0$ | $g_2 = \alpha\gamma_{BCD}$ | $g_3 = 2\alpha$ |
| $d^{-(d_1+d_3+d_4+d_5)}$ | $g_0 = exp[-\alpha\tau_1(\gamma_D - \gamma_{CD}) - \alpha\tau_2(\gamma_B + \gamma_{CD} - 2\gamma_{BCD}) - \alpha\tau_3(\gamma_A + \gamma_{BCD} - 2)]$ | $g_1 = \alpha\gamma_{BCD}$ | $g_2 = 0$ | $g_3 = 2\alpha$ |
| $d^{-(d_2+d_3+d_4+d_5)}$ | $g_0 = exp[-\alpha\tau_1(\gamma_C + \gamma_D - 2\gamma_{CD}) - \alpha\tau_2(\gamma_B + 2\gamma_{CD} - 3\gamma_{BCD})]$ | $g_1 = \alpha\gamma_{BCD}$ | $g_2 = 2\alpha\gamma_{BCD}$ | $g_3 = 0$ |
| $d^{-(d_1+d_2+d_3+d_4+d_5)}$ | $g_0 = exp[-\alpha\tau_1(\gamma_C + \gamma_D - 2\gamma_{CD}) - \alpha\tau_2(\gamma_B + 2\gamma_{CD} - 3\gamma_{BCD}) - \alpha\tau_3(\gamma_A + \gamma_{BCD} - 2)]$ | $g_1 = \alpha\gamma_{BCD}$ | $g_2 = \alpha\gamma_{BCD}$ | $g_3 = 2\alpha$ |

| | **History 17** | | | |
|---|---|---|---|---|
| $d^{-(d_1+d_2)}$ | $g_0 = exp[-\alpha\tau_1(\gamma_C - \gamma_{CD}) - \alpha\tau_2(\gamma_{CD} - \gamma_{BCD}) - \alpha\tau_3(\gamma_A + \gamma_{BCD} - 2)]$ | $g_1 = 0$ | $g_2 = 0$ | $g_3 = 2\alpha$ |
| $d^{-(d_3+d_4)}$ | $g_0 = exp[-\alpha\tau_1(\gamma_D - \gamma_{CD}) - \alpha\tau_2(\gamma_B + \gamma_{CD} - 2\gamma_{BCD})]$ | $g_1 = 2\alpha\gamma_{BCD}$ | $g_2 = 0$ | $g_3 = 0$ |
| $d^{-(d_1+d_3+d_5)}$ | $g_0 = exp[-\alpha\tau_2(\gamma_B - \gamma_{BCD}) - \alpha\tau_3(\gamma_A + \gamma_{BCD} - 2)]$ | $g_1 = 0$ | $g_2 = 0$ | $g_3 = 2\alpha$ |
| $d^{-(d_1+d_4+d_5)}$ | $g_0 = exp[-\alpha\tau_1(\gamma_D - \gamma_{CD}) - \alpha\tau_2(\gamma_{CD} - \gamma_{BCD}) - \alpha\tau_3(\gamma_A + \gamma_{BCD} - 2)]$ | $g_1 = 0$ | $g_2 = 0$ | $g_3 = 2\alpha$ |
| $d^{-(d_2+d_3+d_5)}$ | $g_0 = exp[-\alpha\tau_1(\gamma_C - \gamma_{CD}) - \alpha\tau_2(\gamma_B + \gamma_{CD} - 2\gamma_{BCD}) - 2\alpha\tau_3(\gamma_{BCD} - 1)]$ | $g_1 = 0$ | $g_2 = 2\alpha$ | $g_3 = 0$ |
| $d^{-(d_2+d_4+d_5)}$ | $g_0 = exp[-\alpha\tau_1(\gamma_C + \gamma_D - 2\gamma_{CD}) - 2\alpha\tau_2(\gamma_{CD} - \gamma_{BCD}) - 2\alpha\tau_3(\gamma_{BCD} - 1)]$ | $g_1 = 0$ | $g_2 = 2\alpha$ | $g_3 = 0$ |
| $d^{-(d_1+d_2+d_3+d_4)}$ | $g_0 = exp[-\alpha\tau_1(\gamma_C + \gamma_D - 2\gamma_{CD}) - \alpha\tau_2(\gamma_B + 2\gamma_{CD} - 3\gamma_{BCD}) - \alpha\tau_3(\gamma_A + \gamma_{BCD} - 2)]$ | $g_1 = 2\alpha\gamma_{BCD}$ | $g_2 = 0$ | $g_3 = 2\alpha$ |
| $d^{-(d_1+d_2+d_3+d_5)}$ | $g_0 = exp[-\alpha\tau_1(\gamma_C - \gamma_{CD}) - \alpha\tau_2(\gamma_B + \gamma_{CD} - 2\gamma_{BCD}) - \alpha\tau_3(\gamma_A + 2\gamma_{BCD} - 3)]$ | $g_1 = 0$ | $g_2 = \alpha$ | $g_3 = 2\alpha$ |
| $d^{-(d_1+d_2+d_4+d_5)}$ | $g_0 = exp[-\alpha\tau_1(\gamma_C + \gamma_D - 2\gamma_{CD}) - 2\alpha\tau_2(\gamma_{CD} - \gamma_{BCD}) - \alpha\tau_3(\gamma_A + 2\gamma_{BCD} - 3)]$ | $g_1 = 0$ | $g_2 = \alpha$ | $g_3 = 2\alpha$ |
| $d^{-(d_1+d_3+d_4+d_5)}$ | $g_0 = exp[-\alpha\tau_1(\gamma_D - \gamma_{CD}) - \alpha\tau_2(\gamma_B + \gamma_{CD} - 2\gamma_{BCD}) - \alpha\tau_3(\gamma_A + \gamma_{BCD} - 2)]$ | $g_1 = \alpha\gamma_{BCD}$ | $g_2 = 0$ | $g_3 = 2\alpha$ |
| $d^{-(d_2+d_3+d_4+d_5)}$ | $g_0 = exp[-\alpha\tau_1(\gamma_C + \gamma_D - 2\gamma_{CD}) - \alpha\tau_2(\gamma_B + 2\gamma_{CD} - 3\gamma_{BCD}) - 2\alpha\tau_3(\gamma_{BCD} - 1)]$ | $g_1 = \alpha\gamma_{BCD}$ | $g_2 = 2\alpha$ | $g_3 = 0$ |
| $d^{-(d_1+d_2+d_3+d_4+d_5)}$ | $g_0 = exp[-\alpha\tau_1(\gamma_C + \gamma_D - 2\gamma_{CD}) - \alpha\tau_2(\gamma_B + 2\gamma_{CD} - 3\gamma_{BCD}) - \alpha\tau_3(\gamma_A + 2\gamma_{BCD} - 3)]$ | $g_1 = \alpha\gamma_{BCD}$ | $g_2 = \alpha$ | $g_3 = 2\alpha$ |

| | **History 18** | | | |
|---|---|---|---|---|
| $d^{-(d_1+d_2)}$ | $g_0 = exp[-\alpha\tau_1(\gamma_C - \gamma_{CD}) - \alpha\tau_2(\gamma_{CD} - \gamma_{BCD}) - \alpha\tau_3(\gamma_A + \gamma_{BCD} - 2)]$ | $g_1 = 0$ | $g_2 = 0$ | $g_3 = 2\alpha$ |
| $d^{-(d_3+d_4)}$ | $g_0 = exp[-\alpha\tau_1(\gamma_D - \gamma_{CD}) - \alpha\tau_2(\gamma_B + \gamma_{CD} - 2\gamma_{BCD}) - 2\alpha\tau_3(\gamma_{BCD} - 1]$ | $g_1 = 2\alpha$ | $g_2 = 0$ | $g_3 = 0$ |
| $d^{-(d_1+d_3+d_5)}$ | $g_0 = exp[-\alpha\tau_2(\gamma_B - \gamma_{BCD}) - \alpha\tau_3(\gamma_A + \gamma_{BCD} - 2)]$ | $g_1 = 0$ | $g_2 = 0$ | $g_3 = 2\alpha$ |
| $d^{-(d_1+d_4+d_5)}$ | $g_0 = exp[-\alpha\tau_1(\gamma_D - \gamma_{CD}) - \alpha\tau_2(\gamma_{CD} - \gamma_{BCD}) - \alpha\tau_3(\gamma_A + \gamma_{BCD} - 2)]$ | $g_1 = 0$ | $g_2 = 0$ | $g_3 = 2\alpha$ |
| $d^{-(d_2+d_3+d_5)}$ | $g_0 = exp[-\alpha\tau_1(\gamma_C - \gamma_{CD}) - \alpha\tau_2(\gamma_B + \gamma_{CD} - 2\gamma_{BCD}) - 2\alpha\tau_3(\gamma_{BCD} - 1)]$ | $g_1 = 0$ | $g_2 = 2\alpha$ | $g_3 = 0$ |
| $d^{-(d_2+d_4+d_5)}$ | $g_0 = exp[-\alpha\tau_1(\gamma_C + \gamma_D - 2\gamma_{CD}) - 2\alpha\tau_2(\gamma_{CD} - \gamma_{BCD}) - 2\alpha\tau_3(\gamma_{BCD} - 1)]$ | $g_1 = 0$ | $g_2 = 2\alpha$ | $g_3 = 0$ |
| $d^{-(d_1+d_2+d_3+d_4)}$ | $g_0 = exp[-\alpha\tau_1(\gamma_C + \gamma_D - 2\gamma_{CD}) - \alpha\tau_2(\gamma_B + 2\gamma_{CD} - 3\gamma_{BCD}) - \alpha\tau_3(\gamma_A + 3\gamma_{BCD} - 4)]$ | $g_1 = 2\alpha$ | $g_2 = 0$ | $g_3 = 2\alpha$ |
| $d^{-(d_1+d_2+d_3+d_5)}$ | $g_0 = exp[-\alpha\tau_1(\gamma_C - \gamma_{CD}) - \alpha\tau_2(\gamma_B + \gamma_{CD} - 2\gamma_{BCD}) - \alpha\tau_3(\gamma_A + 2\gamma_{BCD} - 3)]$ | $g_1 = 0$ | $g_2 = \alpha$ | $g_3 = 2\alpha$ |
| $d^{-(d_1+d_2+d_4+d_5)}$ | $g_0 = exp[-\alpha\tau_1(\gamma_C + \gamma_D - 2\gamma_{CD}) - 2\alpha\tau_2(\gamma_{CD} - \gamma_{BCD}) - \alpha\tau_3(\gamma_A + 2\gamma_{BCD} - 3)]$ | $g_1 = 0$ | $g_2 = \alpha$ | $g_3 = 2\alpha$ |
| $d^{-(d_1+d_3+d_4+d_5)}$ | $g_0 = exp[-\alpha\tau_1(\gamma_D - \gamma_{CD}) - \alpha\tau_2(\gamma_B + \gamma_{CD} - 2\gamma_{BCD}) - \alpha\tau_3(\gamma_A + 2\gamma_{BCD} - 3)]$ | $g_1 = \alpha$ | $g_2 = 0$ | $g_3 = 2\alpha$ |
| $d^{-(d_2+d_3+d_4+d_5)}$ | $g_0 = exp[-\alpha\tau_1(\gamma_C + \gamma_D - 2\gamma_{CD}) - \alpha\tau_2(\gamma_B + 2\gamma_{CD} - 3\gamma_{BCD}) - 3\alpha\tau_3(\gamma_{BCD} - 1)]$ | $g_1 = \alpha$ | $g_2 = 2\alpha$ | $g_3 = 0$ |
| $d^{-(d_1+d_2+d_3+d_4+d_5)}$ | $g_0 = exp[-\alpha\tau_1(\gamma_C + \gamma_D - 2\gamma_{CD}) - \alpha\tau_2(\gamma_B + 2\gamma_{CD} - 3\gamma_{BCD}) - \alpha\tau_3(\gamma_A + 3\gamma_{BCD} - 4)]$ | $g_1 = \alpha$ | $g_2 = \alpha$ | $g_3 = 2\alpha$ |



| | **History 19** | | | |
|---|---|---|---|---|
| $d^{-(d_1+d_2)}$ | $g_0 = exp[-\alpha\tau_1(\gamma_D - \gamma_{CD}) - \alpha\tau_2(\gamma_{CD} - \gamma_{BCD}) - \alpha\tau_3(\gamma_A + \gamma_{BCD} - 2)]$ | $g_1 = 0$ | $g_2 = 0$ | $g_3 = 2\alpha$ |
| $d^{-(d_3+d_4)}$ | $g_0 = exp[-\alpha\tau_1(\gamma_C - \gamma_{CD}) - \alpha\tau_2(\gamma_B + \gamma_{CD} - 2\gamma_{BCD})]$ | $g_1 = 2\alpha\gamma_{BCD}$ | $g_2 = 0$ | $g_3 = 0$ |
| $d^{-(d_1+d_3+d_5)}$ | $g_0 = exp[-\alpha\tau_2(\gamma_B - \gamma_{BCD}) - \alpha\tau_3(\gamma_A + \gamma_{BCD} - 2)]$ | $g_1 = 0$ | $g_2 = 0$ | $g_3 = 2\alpha$ |
| $d^{-(d_1+d_4+d_5)}$ | $g_0 = exp[-\alpha\tau_1(\gamma_C - \gamma_{CD}) - \alpha\tau_2(\gamma_{CD} - \gamma_{BCD}) - \alpha\tau_3(\gamma_A + \gamma_{BCD} - 2)]$ | $g_1 = 0$ | $g_2 = 0$ | $g_3 = 2\alpha$ |
| $d^{-(d_2+d_3+d_5)}$ | $g_0 = exp[-\alpha\tau_1(\gamma_D - \gamma_{CD}) - \alpha\tau_2(\gamma_B + \gamma_{CD} - 2\gamma_{BCD})]$ | $g_1 = 0$ | $g_2 = 2\alpha\gamma_{BCD}$ | $g_3 = 0$ |
| $d^{-(d_2+d_4+d_5)}$ | $g_0 = exp[-\alpha\tau_1(\gamma_C + \gamma_D - 2\gamma_{CD}) - 2\alpha\tau_2(\gamma_{CD} - \gamma_{BCD})]$ | $g_1 = 0$ | $g_2 = 2\alpha\gamma_{BCD}$ | $g_3 = 0$ |
| $d^{-(d_1+d_2+d_3+d_4)}$ | $g_0 = exp[-\alpha\tau_1(\gamma_C + \gamma_D - 2\gamma_{CD}) - \alpha\tau_2(\gamma_B + 2\gamma_{CD} - 3\gamma_{BCD}) - \alpha\tau_3(\gamma_A + \gamma_{BCD} - 2)]$ | $g_1 = 2\alpha\gamma_{BCD}$ | $g_2 = 0$ | $g_3 = 2\alpha$ |
| $d^{-(d_1+d_2+d_3+d_5)}$ | $g_0 = exp[-\alpha\tau_1(\gamma_D - \gamma_{CD}) - \alpha\tau_2(\gamma_B + \gamma_{CD} - 2\gamma_{BCD}) - \alpha\tau_3(\gamma_A + \gamma_{BCD} - 2)]$ | $g_1 = 0$ | $g_2 = \alpha\gamma_{BCD}$ | $g_3 = 2\alpha$ |
| $d^{-(d_1+d_2+d_4+d_5)}$ | $g_0 = exp[-\alpha\tau_1(\gamma_C + \gamma_D - 2\gamma_{CD}) - 2\alpha\tau_2(\gamma_{CD} - \gamma_{BCD}) - \alpha\tau_3(\gamma_A + \gamma_{BCD} - 2)]$ | $g_1 = 0$ | $g_2 = \alpha\gamma_{BCD}$ | $g_3 = 2\alpha$ |
| $d^{-(d_1+d_3+d_4+d_5)}$ | $g_0 = exp[-\alpha\tau_1(\gamma_C - \gamma_{CD}) - \alpha\tau_2(\gamma_B + \gamma_{CD} - 2\gamma_{BCD}) - \alpha\tau_3(\gamma_A + \gamma_{BCD} - 2)]$ | $g_1 = \alpha\gamma_{BCD}$ | $g_2 = 0$ | $g_3 = 2\alpha$ |
| $d^{-(d_2+d_3+d_4+d_5)}$ | $g_0 = exp[-\alpha\tau_1(\gamma_C + \gamma_D - 2\gamma_{CD}) - \alpha\tau_2(\gamma_B + 2\gamma_{CD} - 3\gamma_{BCD})]$ | $g_1 = \alpha\gamma_{BCD}$ | $g_2 = 2\alpha\gamma_{BCD}$ | $g_3 = 0$ |
| $d^{-(d_1+d_2+d_3+d_4+d_5)}$ | $g_0 = exp[-\alpha\tau_1(\gamma_C + \gamma_D - 2\gamma_{CD}) - \alpha\tau_2(\gamma_B + 2\gamma_{CD} - 3\gamma_{BCD}) - \alpha\tau_3(\gamma_A + \gamma_{BCD} - 2)]$ | $g_1 = \alpha\gamma_{BCD}$ | $g_2 = \alpha\gamma_{BCD}$ | $g_3 = 2\alpha$ |

| | **History 20** | | | |
|---|---|---|---|---|
| $d^{-(d_1+d_2)}$ | $g_0 = exp[-\alpha\tau_1(\gamma_D - \gamma_{CD}) - \alpha\tau_2(\gamma_{CD} - \gamma_{BCD}) - \alpha\tau_3(\gamma_A + \gamma_{BCD} - 2)]$ | $g_1 = 0$ | $g_2 = 0$ | $g_3 = 2\alpha$ |
| $d^{-(d_3+d_4)}$ | $g_0 = exp[-\alpha\tau_1(\gamma_C - \gamma_{CD}) - \alpha\tau_2(\gamma_B + \gamma_{CD} - 2\gamma_{BCD})]$ | $g_1 = 2\alpha\gamma_{BCD}$ | $g_2 = 0$ | $g_3 = 0$ |
| $d^{-(d_1+d_3+d_5)}$ | $g_0 = exp[-\alpha\tau_2(\gamma_B - \gamma_{BCD}) - \alpha\tau_3(\gamma_A + \gamma_{BCD} - 2)]$ | $g_1 = 0$ | $g_2 = 0$ | $g_3 = 2\alpha$ |
| $d^{-(d_1+d_4+d_5)}$ | $g_0 = exp[-\alpha\tau_1(\gamma_C - \gamma_{CD}) - \alpha\tau_2(\gamma_{CD} - \gamma_{BCD}) - \alpha\tau_3(\gamma_A + \gamma_{BCD} - 2)]$ | $g_1 = 0$ | $g_2 = 0$ | $g_3 = 2\alpha$ |
| $d^{-(d_2+d_3+d_5)}$ | $g_0 = exp[-\alpha\tau_1(\gamma_D - \gamma_{CD}) - \alpha\tau_2(\gamma_B + \gamma_{CD} - 2\gamma_{BCD}) - 2\alpha\tau_3(\gamma_{BCD} - 1)]$ | $g_1 = 0$ | $g_2 = 2\alpha$ | $g_3 = 0$ |
| $d^{-(d_2+d_4+d_5)}$ | $g_0 = exp[-\alpha\tau_1(\gamma_C + \gamma_D - 2\gamma_{CD}) - 2\alpha\tau_2(\gamma_{CD} - \gamma_{BCD}) - 2\alpha\tau_3(\gamma_{BCD} - 1)]$ | $g_1 = 0$ | $g_2 = 2\alpha$ | $g_3 = 0$ |
| $d^{-(d_1+d_2+d_3+d_4)}$ | $g_0 = exp[-\alpha\tau_1(\gamma_C + \gamma_D - 2\gamma_{CD}) - \alpha\tau_2(\gamma_B + 2\gamma_{CD} - 3\gamma_{BCD}) - \alpha\tau_3(\gamma_A + \gamma_{BCD} - 2)]$ | $g_1 = 2\alpha\gamma_{BCD}$ | $g_2 = 0$ | $g_3 = 2\alpha$ |
| $d^{-(d_1+d_2+d_3+d_5)}$ | $g_0 = exp[-\alpha\tau_1(\gamma_D - \gamma_{CD}) - \alpha\tau_2(\gamma_B + \gamma_{CD} - 2\gamma_{BCD}) - \alpha\tau_3(\gamma_A + 2\gamma_{BCD} - 3)]$ | $g_1 = 0$ | $g_2 = \alpha$ | $g_3 = 2\alpha$ |
| $d^{-(d_1+d_2+d_4+d_5)}$ | $g_0 = exp[-\alpha\tau_1(\gamma_C + \gamma_D - 2\gamma_{CD}) - 2\alpha\tau_2(\gamma_{CD} - \gamma_{BCD}) - \alpha\tau_3(\gamma_A + 2\gamma_{BCD} - 3)]$ | $g_1 = 0$ | $g_2 = \alpha$ | $g_3 = 2\alpha$ |
| $d^{-(d_1+d_3+d_4+d_5)}$ | $g_0 = exp[-\alpha\tau_1(\gamma_C - \gamma_{CD}) - \alpha\tau_2(\gamma_B + \gamma_{CD} - 2\gamma_{BCD}) - \alpha\tau_3(\gamma_A + \gamma_{BCD} - 2)]$ | $g_1 = \alpha\gamma_{BCD}$ | $g_2 = 0$ | $g_3 = 2\alpha$ |
| $d^{-(d_2+d_3+d_4+d_5)}$ | $g_0 = exp[-\alpha\tau_1(\gamma_C + \gamma_D - 2\gamma_{CD}) - \alpha\tau_2(\gamma_B + 2\gamma_{CD} - 3\gamma_{BCD}) - 2\alpha\tau_3(\gamma_{BCD} - 1)]$ | $g_1 = \alpha\gamma_{BCD}$ | $g_2 = 2\alpha$ | $g_3 = 0$ |
| $d^{-(d_1+d_2+d_3+d_4+d_5)}$ | $g_0 = exp[-\alpha\tau_1(\gamma_C + \gamma_D - 2\gamma_{CD}) - \alpha\tau_2(\gamma_B + 2\gamma_{CD} - 3\gamma_{BCD}) - \alpha\tau_3(\gamma_A + 2\gamma_{BCD} - 3)]$ | $g_1 = \alpha\gamma_{BCD}$ | $g_2 = \alpha$ | $g_3 = 2\alpha$ |

| | **History 21** | | | |
|---|---|---|---|---|
| $d^{-(d_1+d_2)}$ | $g_0 = exp[-\alpha\tau_1(\gamma_D - \gamma_{CD}) - \alpha\tau_2(\gamma_{CD} - \gamma_{BCD}) - \alpha\tau_3(\gamma_A + \gamma_{BCD} - 2)]$ | $g_1 = 0$ | $g_2 = 0$ | $g_3 = 2\alpha$ |
| $d^{-(d_3+d_4)}$ | $g_0 = exp[-\alpha\tau_1(\gamma_C - \gamma_{CD}) - \alpha\tau_2(\gamma_B + \gamma_{CD} - 2\gamma_{BCD}) - 2\alpha\tau_3(\gamma_{BCD} - 1)]$ | $g_1 = 2\alpha$ | $g_2 = 0$ | $g_3 = 0$ |
| $d^{-(d_1+d_3+d_5)}$ | $g_0 = exp[-\alpha\tau_2(\gamma_B - \gamma_{BCD}) - \alpha\tau_3(\gamma_A + \gamma_{BCD} - 2)]$ | $g_1 = 0$ | $g_2 = 0$ | $g_3 = 2\alpha$ |
| $d^{-(d_1+d_4+d_5)}$ | $g_0 = exp[-\alpha\tau_1(\gamma_C - \gamma_{CD}) - \alpha\tau_2(\gamma_{CD} - \gamma_{BCD}) - \alpha\tau_3(\gamma_A + \gamma_{BCD} - 2)]$ | $g_1 = 0$ | $g_2 = 0$ | $g_3 = 2\alpha$ |
| $d^{-(d_2+d_3+d_5)}$ | $g_0 = exp[-\alpha\tau_1(\gamma_D - \gamma_{CD}) - \alpha\tau_2(\gamma_B + \gamma_{CD} - 2\gamma_{BCD}) - 2\alpha\tau_3(\gamma_{BCD} - 1)]$ | $g_1 = 0$ | $g_2 = 2\alpha$ | $g_3 = 0$ |
| $d^{-(d_2+d_4+d_5)}$ | $g_0 = exp[-\alpha\tau_1(\gamma_C + \gamma_D - 2\gamma_{CD}) - 2\alpha\tau_2(\gamma_{CD} - \gamma_{BCD}) - 2\alpha\tau_3(\gamma_{BCD} - 1)]$ | $g_1 = 0$ | $g_2 = 2\alpha$ | $g_3 = 0$ |
| $d^{-(d_1+d_2+d_3+d_4)}$ | $g_0 = exp[-\alpha\tau_1(\gamma_C + \gamma_D - 2\gamma_{CD}) - \alpha\tau_2(\gamma_B + 2\gamma_{CD} - 3\gamma_{BCD}) - \alpha\tau_3(\gamma_A + 3\gamma_{BCD} - 4)]$ | $g_1 = 2\alpha$ | $g_2 = 0$ | $g_3 = 2\alpha$ |
| $d^{-(d_1+d_2+d_3+d_5)}$ | $g_0 = exp[-\alpha\tau_1(\gamma_D - \gamma_{CD}) - \alpha\tau_2(\gamma_B + \gamma_{CD} - 2\gamma_{BCD}) - \alpha\tau_3(\gamma_A + 2\gamma_{BCD} - 3)]$ | $g_1 = 0$ | $g_2 = \alpha$ | $g_3 = 2\alpha$ |
| $d^{-(d_1+d_2+d_4+d_5)}$ | $g_0 = exp[-\alpha\tau_1(\gamma_C + \gamma_D - 2\gamma_{CD}) - 2\alpha\tau_2(\gamma_{CD} - \gamma_{BCD}) - \alpha\tau_3(\gamma_A + 2\gamma_{BCD} - 3)]$ | $g_1 = 0$ | $g_2 = \alpha$ | $g_3 = 2\alpha$ |
| $d^{-(d_1+d_3+d_4+d_5)}$ | $g_0 = exp[-\alpha\tau_1(\gamma_C - \gamma_{CD}) - \alpha\tau_2(\gamma_B + \gamma_{CD} - 2\gamma_{BCD}) - \alpha\tau_3(\gamma_A + 2\gamma_{BCD} - 3)]$ | $g_1 = \alpha$ | $g_2 = 0$ | $g_3 = 2\alpha$ |
| $d^{-(d_2+d_3+d_4+d_5)}$ | $g_0 = exp[-\alpha\tau_1(\gamma_C + \gamma_D - 2\gamma_{CD}) - \alpha\tau_2(\gamma_B + 2\gamma_{CD} - 3\gamma_{BCD}) - 3\alpha\tau_3(\gamma_{BCD} - 1)]$ | $g_1 = \alpha$ | $g_2 = 2\alpha$ | $g_3 = 0$ |
| $d^{-(d_1+d_2+d_3+d_4+d_5)}$ | $g_0 = exp[-\alpha\tau_1(\gamma_C + \gamma_D - 2\gamma_{CD}) - \alpha\tau_2(\gamma_B + 2\gamma_{CD} - 3\gamma_{BCD}) - \alpha\tau_3(\gamma_A + 3\gamma_{BCD} - 4)]$ | $g_1 = \alpha$ | $g_2 = \alpha$ | $g_3 = 2\alpha$ |

| | **History 22** | | | |
|---|---|---|---|---|
| $d^{-(d_1+d_2)}$ | $g_0 = exp[-\alpha\tau_2(\gamma_B - \gamma_{BCD}) - \alpha\tau_3(\gamma_A + \gamma_{BCD} - 2)]$ | $g_1 = 0$ | $g_2 = 0$ | $g_3 = 2\alpha$ |
| $d^{-(d_3+d_4)}$ | $g_0 = exp[-\alpha\tau_1(\gamma_C + \gamma_D - 2\gamma_{CD})]$ | $g_1 = 2\alpha\gamma_{CD}$ | $g_2 = 0$ | $g_3 = 0$ |
| $d^{-(d_1+d_3+d_5)}$ | $g_0 = exp[-\alpha\tau_1(\gamma_C - \gamma_{CD}) - \alpha\tau_2(\gamma_B + \gamma_{CD} - 2\gamma_{BCD}) - 2\alpha\tau_3(\gamma_{BCD} - 1)]$ | $g_1 = 0$ | $g_2 = 0$ | $g_3 = 2\alpha$ |
| $d^{-(d_1+d_4+d_5)}$ | $g_0 = exp[-\alpha\tau_1(\gamma_D - \gamma_{CD}) - \alpha\tau_2(\gamma_B + \gamma_{CD} - 2\gamma_{BCD}) - 2\alpha\tau_3(\gamma_{BCD} - 1)]$ | $g_1 = 0$ | $g_2 = 0$ | $g_3 = 2\alpha$ |
| $d^{-(d_2+d_3+d_5)}$ | $g_0 = exp[-\alpha\tau_1(\gamma_C - \gamma_{CD}) - \alpha\tau_2(\gamma_{CD} - \gamma_{BCD}) - \alpha\tau_3(\gamma_A + \gamma_{BCD} - 2)]$ | $g_1 = 0$ | $g_2 = 2\alpha$ | $g_3 = 0$ |
| $d^{-(d_2+d_4+d_5)}$ | $g_0 = exp[-\alpha\tau_1(\gamma_D - \gamma_{CD}) - \alpha\tau_2(\gamma_{CD} - \gamma_{BCD}) - \alpha\tau_3(\gamma_A + \gamma_{BCD} - 2))]$ | $g_1 = 0$ | $g_2 = 2\alpha$ | $g_3 = 0$ |
| $d^{-(d_1+d_2+d_3+d_4)}$ | $g_0 = exp[-\alpha\tau_1(\gamma_C + \gamma_D - 2\gamma_{CD}) - \alpha\tau_2(\gamma_B - \gamma_{BCD}) - \alpha\tau_3(\gamma_A + \gamma_{BCD} - 2)]$ | $g_1 = 2\alpha\gamma_{CD}$ | $g_2 = 0$ | $g_3 = 2\alpha$ |
| $d^{-(d_1+d_2+d_3+d_5)}$ | $g_0 = exp[-\alpha\tau_1(\gamma_C - \gamma_{CD}) - \alpha\tau_2(\gamma_B + \gamma_{CD} - 2\gamma_{BCD}) - \alpha\tau_3(\gamma_A + 2\gamma_{BCD} - 3)]$ | $g_1 = 0$ | $g_2 = \alpha$ | $g_3 = 2\alpha$ |
| $d^{-(d_1+d_2+d_4+d_5)}$ | $g_0 = exp[-\alpha\tau_1(\gamma_D - \gamma_{CD}) - \alpha\tau_2(\gamma_B + \gamma_{CD} - 2\gamma_{BCD}) - \alpha\tau_3(\gamma_A + 2\gamma_{BCD} - 3)]$ | $g_1 = 0$ | $g_2 = \alpha$ | $g_3 = 2\alpha$ |
| $d^{-(d_1+d_3+d_4+d_5)}$ | $g_0 = exp[-\alpha\tau_1(\gamma_C + \gamma_D - 2\gamma_{CD}) - \alpha\tau_2(\gamma_B + \gamma_{CD} - 2\gamma_{BCD}) - 2\alpha\tau_3(\gamma_{BCD} - 1)]$ | $g_1 = \alpha\gamma_{CD}$ | $g_2 = 0$ | $g_3 = 2\alpha$ |
| $d^{-(d_2+d_3+d_4+d_5)}$ | $g_0 = exp[-\alpha\tau_1(\gamma_C + \gamma_D - 2\gamma_{CD}) - \alpha\tau_2(\gamma_{CD} - \gamma_{BCD}) - \alpha\tau_3(\gamma_A + \gamma_{BCD} - 2)]$ | $g_1 = \alpha\gamma_{CD}$ | $g_2 = 2\alpha$ | $g_3 = 0$ |
| $d^{-(d_1+d_2+d_3+d_4+d_5)}$ | $g_0 = exp[-\alpha\tau_1(\gamma_C + \gamma_D - 2\gamma_{CD}) - \alpha\tau_2(\gamma_B + \gamma_{CD} - 2\gamma_{BCD}) - \alpha\tau_3(\gamma_A + 2\gamma_{BCD} - 3)]$ | $g_1 = \alpha\gamma_{CD}$ | $g_2 = \alpha$ | $g_3 = 2\alpha$ |

| | **History 23** | | | |
|---|---|---|---|---|
| $d^{-(d_1+d_2)}$ | $g_0 = exp[-\alpha\tau_2(\gamma_B - \gamma_{BCD}) - \alpha\tau_3(\gamma_A + \gamma_{BCD} - 2)]$ | $g_1 = 0$ | $g_2 = 0$ | $g_3 = 2\alpha$ |
| $d^{-(d_3+d_4)}$ | $g_0 = exp[-\alpha\tau_1(\gamma_C + \gamma_D - 2\gamma_{CD}) - 2\alpha\tau_2(\gamma_{CD} - \gamma_{BCD})]$ | $g_1 = 2\alpha\gamma_{BCD}$ | $g_2 = 0$ | $g_3 = 0$ |
| $d^{-(d_1+d_3+d_5)}$ | $g_0 = exp[-\alpha\tau_1(\gamma_C - \gamma_{CD}) - \alpha\tau_2(\gamma_B + \gamma_{CD} - 2\gamma_{BCD}) - 2\alpha\tau_3(\gamma_{BCD} - 1)]$ | $g_1 = 0$ | $g_2 = 0$ | $g_3 = 2\alpha$ |
| $d^{-(d_1+d_4+d_5)}$ | $g_0 = exp[-\alpha\tau_1(\gamma_D - \gamma_{CD}) - \alpha\tau_2(\gamma_B + \gamma_{CD} - 2\gamma_{BCD}) - 2\alpha\tau_3(\gamma_{BCD} - 1)]$ | $g_1 = 0$ | $g_2 = 0$ | $g_3 = 2\alpha$ |
| $d^{-(d_2+d_3+d_5)}$ | $g_0 = exp[-\alpha\tau_1(\gamma_C - \gamma_{CD}) - \alpha\tau_2(\gamma_{CD} - \gamma_{BCD}) - \alpha\tau_3(\gamma_A + \gamma_{BCD} - 2)]$ | $g_1 = 0$ | $g_2 = 2\alpha$ | $g_3 = 0$ |
| $d^{-(d_2+d_4+d_5)}$ | $g_0 = exp[-\alpha\tau_1(\gamma_D - \gamma_{CD}) - \alpha\tau_2(\gamma_{CD} - \gamma_{BCD}) - \alpha\tau_3(\gamma_A + \gamma_{BCD} - 2))]$ | $g_1 = 0$ | $g_2 = 2\alpha$ | $g_3 = 0$ |
| $d^{-(d_1+d_2+d_3+d_4)}$ | $g_0 = exp[-\alpha\tau_1(\gamma_C + \gamma_D - 2\gamma_{CD}) - \alpha\tau_2(\gamma_B + 2\gamma_{CD} - 3\gamma_{BCD}) - \alpha\tau_3(\gamma_A + \gamma_{BCD} - 2)]$ | $g_1 = 2\alpha\gamma_{BCD}$ | $g_2 = 0$ | $g_3 = 2\alpha$ |
| $d^{-(d_1+d_2+d_3+d_5)}$ | $g_0 = exp[-\alpha\tau_1(\gamma_C - \gamma_{CD}) - \alpha\tau_2(\gamma_B + \gamma_{CD} - 2\gamma_{BCD}) - \alpha\tau_3(\gamma_A + 2\gamma_{BCD} - 3)]$ | $g_1 = 0$ | $g_2 = \alpha$ | $g_3 = 2\alpha$ |
| $d^{-(d_1+d_2+d_4+d_5)}$ | $g_0 = exp[-\alpha\tau_1(\gamma_D - \gamma_{CD}) - \alpha\tau_2(\gamma_B + \gamma_{CD} - 2\gamma_{BCD}) - \alpha\tau_3(\gamma_A + 2\gamma_{BCD} - 3)]$ | $g_1 = 0$ | $g_2 = \alpha$ | $g_3 = 2\alpha$ |
| $d^{-(d_1+d_3+d_4+d_5)}$ | $g_0 = exp[-\alpha\tau_1(\gamma_C + \gamma_D - 2\gamma_{CD}) - \alpha\tau_2(\gamma_B + 2\gamma_{CD} - 3\gamma_{BCD}) - 2\alpha\tau_3(\gamma_{BCD} - 1)]$ | $g_1 = \alpha\gamma_{BCD}$ | $g_2 = 0$ | $g_3 = 2\alpha$ |
| $d^{-(d_2+d_3+d_4+d_5)}$ | $g_0 = exp[-\alpha\tau_1(\gamma_C + \gamma_D - 2\gamma_{CD}) - 2\alpha\tau_2(\gamma_{CD} - \gamma_{BCD}) - \alpha\tau_3(\gamma_A + \gamma_{BCD} - 2)]$ | $g_1 = \alpha\gamma_{BCD}$ | $g_2 = 2\alpha$ | $g_3 = 0$ |
| $d^{-(d_1+d_2+d_3+d_4+d_5)}$ | $g_0 = exp[-\alpha\tau_1(\gamma_C + \gamma_D - 2\gamma_{CD}) - \alpha\tau_2(\gamma_B + 2\gamma_{CD} - 3\gamma_{BCD}) - \alpha\tau_3(\gamma_A + 2\gamma_{BCD} - 3)]$ | $g_1 = \alpha\gamma_{BCD}$ | $g_2 = \alpha$ | $g_3 = 2\alpha$ |

| | **History 24** | | | |
|---|---|---|---|---|
| $d^{-(d_1+d_2)}$ | $g_0 = exp[-\alpha\tau_2(\gamma_B - \gamma_{BCD}) - \alpha\tau_3(\gamma_A + \gamma_{BCD} - 2)]$ | $g_1 = 0$ | $g_2 = 0$ | $g_3 = 2\alpha$ |
| $d^{-(d_3+d_4)}$ | $g_0 = exp[-\alpha\tau_1(\gamma_C + \gamma_D - 2\gamma_{CD}) - 2\alpha\tau_2(\gamma_{CD} - \gamma_{BCD}) - 2\alpha\tau_3(\gamma_{BCD} - 1)]$ | $g_1 = 2\alpha$ | $g_2 = 0$ | $g_3 = 0$ |
| $d^{-(d_1+d_3+d_5)}$ | $g_0 = exp[-\alpha\tau_1(\gamma_C - \gamma_{CD}) - \alpha\tau_2(\gamma_B + \gamma_{CD} - 2\gamma_{BCD}) - 2\alpha\tau_3(\gamma_{BCD} - 1)]$ | $g_1 = 0$ | $g_2 = 0$ | $g_3 = 2\alpha$ |
| $d^{-(d_1+d_4+d_5)}$ | $g_0 = exp[-\alpha\tau_1(\gamma_D - \gamma_{CD}) - \alpha\tau_2(\gamma_B + \gamma_{CD} - 2\gamma_{BCD}) - 2\alpha\tau_3(\gamma_{BCD} - 1)]$ | $g_1 = 0$ | $g_2 = 0$ | $g_3 = 2\alpha$ |
| $d^{-(d_2+d_3+d_5)}$ | $g_0 = exp[-\alpha\tau_1(\gamma_C - \gamma_{CD}) - \alpha\tau_2(\gamma_{CD} - \gamma_{BCD}) - \alpha\tau_3(\gamma_A + \gamma_{BCD} - 2)]$ | $g_1 = 0$ | $g_2 = 2\alpha$ | $g_3 = 0$ |
| $d^{-(d_2+d_4+d_5)}$ | $g_0 = exp[-\alpha\tau_1(\gamma_D - \gamma_{CD}) - \alpha\tau_2(\gamma_{CD} - \gamma_{BCD}) - \alpha\tau_3(\gamma_A + \gamma_{BCD} - 2))]$ | $g_1 = 0$ | $g_2 = 2\alpha$ | $g_3 = 0$ |
| $d^{-(d_1+d_2+d_3+d_4)}$ | $g_0 = exp[-\alpha\tau_1(\gamma_C + \gamma_D - 2\gamma_{CD}) - \alpha\tau_2(\gamma_B + 2\gamma_{CD} - 3\gamma_{BCD}) - \alpha\tau_3(\gamma_A + 3\gamma_{BCD} - 4)]$ | $g_1 = 2\alpha$ | $g_2 = 0$ | $g_3 = 2\alpha$ |
| $d^{-(d_1+d_2+d_3+d_5)}$ | $g_0 = exp[-\alpha\tau_1(\gamma_C - \gamma_{CD}) - \alpha\tau_2(\gamma_B + \gamma_{CD} - 2\gamma_{BCD}) - \alpha\tau_3(\gamma_A + 2\gamma_{BCD} - 3)]$ | $g_1 = 0$ | $g_2 = \alpha$ | $g_3 = 2\alpha$ |
| $d^{-(d_1+d_2+d_4+d_5)}$ | $g_0 = exp[-\alpha\tau_1(\gamma_D - \gamma_{CD}) - \alpha\tau_2(\gamma_B + \gamma_{CD} - 2\gamma_{BCD}) - \alpha\tau_3(\gamma_A + 2\gamma_{BCD} - 3)]$ | $g_1 = 0$ | $g_2 = \alpha$ | $g_3 = 2\alpha$ |
| $d^{-(d_1+d_3+d_4+d_5)}$ | $g_0 = exp[-\alpha\tau_1(\gamma_C + \gamma_D - 2\gamma_{CD}) - \alpha\tau_2(\gamma_B + 2\gamma_{CD} - 3\gamma_{BCD}) - 3\alpha\tau_3(\gamma_{BCD} - 1)]$ | $g_1 = \alpha$ | $g_2 = 0$ | $g_3 = 2\alpha$ |
| $d^{-(d_2+d_3+d_4+d_5)}$ | $g_0 = exp[-\alpha\tau_1(\gamma_C + \gamma_D - 2\gamma_{CD}) - 2\alpha\tau_2(\gamma_{CD} - \gamma_{BCD}) - \alpha\tau_3(\gamma_A + 2\gamma_{BCD} - 3)]$ | $g_1 = \alpha$ | $g_2 = 2\alpha$ | $g_3 = 0$ |
| $d^{-(d_1+d_2+d_3+d_4+d_5)}$ | $g_0 = exp[-\alpha\tau_1(\gamma_C + \gamma_D - 2\gamma_{CD}) - \alpha\tau_2(\gamma_B + 2\gamma_{CD} - 3\gamma_{BCD}) - \alpha\tau_3(\gamma_A + 3\gamma_{BCD} - 4)]$ | $g_1 = \alpha$ | $g_2 = \alpha$ | $g_3 = 2\alpha$ |

| | **History 25** | | | |
|---|---|---|---|---|
| $d^{-(d_1+d_2)}$ | $g_0 = exp[-\alpha\tau_1(\gamma_C - \gamma_{CD}) - \alpha\tau_2(\gamma_B + \gamma_{CD} - 2\gamma_{BCD}) - 2\alpha\tau_3(\gamma_{BCD} - 1)]$ | $g_1 = 0$ | $g_2 = 0$ | $g_3 = 2\alpha$ |
| $d^{-(d_3+d_4)}$ | $g_0 = exp[-\alpha\tau_1(\gamma_D - \gamma_{CD}) - \alpha\tau_2(\gamma_{CD} - \gamma_{BCD}) - \alpha\tau_3(\gamma_A + \gamma_{BCD} - 2)]$ | $g_1 = 2\alpha$ | $g_2 = 0$ | $g_3 = 0$ |
| $d^{-(d_1+d_3+d_5)}$ | $g_0 = exp[-\alpha\tau_2(\gamma_B - \gamma_{BCD}) - \alpha\tau_3(\gamma_A + \gamma_{BCD} - 2)]$ | $g_1 = 0$ | $g_2 = 0$ | $g_3 = 2\alpha$ |
| $d^{-(d_1+d_4+d_5)}$ | $g_0 = exp[-\alpha\tau_1(\gamma_D - \gamma_{CD}) - \alpha\tau_2(\gamma_B + \gamma_{CD} - 2\gamma_{BCD}) - 2\alpha\tau_3(\gamma_{BCD} - 1)]$ | $g_1 = 0$ | $g_2 = 0$ | $g_3 = 2\alpha$ |
| $d^{-(d_2+d_3+d_5)}$ | $g_0 = exp[-\alpha\tau_1(\gamma_C - \gamma_{CD}) - \alpha\tau_2(\gamma_{CD} - \gamma_{BCD}) - \alpha\tau_3(\gamma_A + \gamma_{BCD} - 2)]$ | $g_1 = 0$ | $g_2 = 2\alpha$ | $g_3 = 0$ |
| $d^{-(d_2+d_4+d_5)}$ | $g_0 = exp[-\alpha\tau_1(\gamma_C + \gamma_D - 2\gamma_{CD}) - 2\alpha\tau_2(\gamma_{CD} - \gamma_{BCD}) - 2\alpha\tau_3(\gamma_{BCD} - 1))]$ | $g_1 = 0$ | $g_2 = 2\alpha$ | $g_3 = 0$ |
| $d^{-(d_1+d_2+d_3+d_4)}$ | $g_0 = exp[-\alpha\tau_1(\gamma_C + \gamma_D - 2\gamma_{CD}) - \alpha\tau_2(\gamma_B + 2\gamma_{CD} - 3\gamma_{BCD}) - \alpha\tau_3(\gamma_A + 3\gamma_{BCD} - 4)]$ | $g_1 = 2\alpha$ | $g_2 = 0$ | $g_3 = 2\alpha$ |
| $d^{-(d_1+d_2+d_3+d_5)}$ | $g_0 = exp[-\alpha\tau_1(\gamma_C - \gamma_{CD}) - \alpha\tau_2(\gamma_B + \gamma_{CD} - 2\gamma_{BCD}) - \alpha\tau_3(\gamma_A + 2\gamma_{BCD} - 3)]$ | $g_1 = 0$ | $g_2 = \alpha$ | $g_3 = 2\alpha$ |
| $d^{-(d_1+d_2+d_4+d_5)}$ | $g_0 = exp[-\alpha\tau_1(\gamma_C + \gamma_D - 2\gamma_{CD}) - \alpha\tau_2(\gamma_B + 2\gamma_{CD} - 3\gamma_{BCD}) - 3\alpha\tau_3(\gamma_{BCD} - 1)]$ | $g_1 = 0$ | $g_2 = \alpha$ | $g_3 = 2\alpha$ |
| $d^{-(d_1+d_3+d_4+d_5)}$ | $g_0 = exp[-\alpha\tau_1(\gamma_D - \gamma_{CD}) - \alpha\tau_2(\gamma_B + \gamma_{CD} - 2\gamma_{BCD}) - \alpha\tau_3(\gamma_A + 2\gamma_{BCD} - 3)]$ | $g_1 = \alpha$ | $g_2 = 0$ | $g_3 = 2\alpha$ |
| $d^{-(d_2+d_3+d_4+d_5)}$ | $g_0 = exp[-\alpha\tau_1(\gamma_C + \gamma_D - 2\gamma_{CD}) - 2\alpha\tau_2(\gamma_{CD} - \gamma_{BCD}) - \alpha\tau_3(\gamma_A + 2\gamma_{BCD} - 3)]$ | $g_1 = \alpha$ | $g_2 = 2\alpha$ | $g_3 = 0$ |
| $d^{-(d_1+d_2+d_3+d_4+d_5)}$ | $g_0 = exp[-\alpha\tau_1(\gamma_C + \gamma_D - 2\gamma_{CD}) - \alpha\tau_2(\gamma_B + 2\gamma_{CD} - 3\gamma_{BCD}) - \alpha\tau_3(\gamma_A + 3\gamma_{BCD} - 4)]$ | $g_1 = \alpha$ | $g_2 = \alpha$ | $g_3 = 2\alpha$ |

| | **History 26** | | | |
|---|---|---|---|---|
| $d^{-(d_1+d_2)}$ | $g_0 = exp[-\alpha\tau_1(\gamma_D - \gamma_{CD}) - \alpha\tau_2(\gamma_B + \gamma_{CD} - 2\gamma_{BCD}) - 2\alpha\tau_3(\gamma_{BCD} - 1)]$ | $g_1 = 0$ | $g_2 = 0$ | $g_3 = 2\alpha$ |
| $d^{-(d_3+d_4)}$ | $g_0 = exp[-\alpha\tau_1(\gamma_C - \gamma_{CD}) - \alpha\tau_2(\gamma_{CD} - \gamma_{BCD}) - \alpha\tau_3(\gamma_A + \gamma_{BCD} - 2)]$ | $g_1 = 2\alpha$ | $g_2 = 0$ | $g_3 = 0$ |
| $d^{-(d_1+d_3+d_5)}$ | $g_0 = exp[-\alpha\tau_2(\gamma_B - \gamma_{BCD}) - \alpha\tau_3(\gamma_A + \gamma_{BCD} - 2)]$ | $g_1 = 0$ | $g_2 = 0$ | $g_3 = 2\alpha$ |
| $d^{-(d_1+d_4+d_5)}$ | $g_0 = exp[-\alpha\tau_1(\gamma_C - \gamma_{CD}) - \alpha\tau_2(\gamma_B + \gamma_{CD} - 2\gamma_{BCD}) - 2\alpha\tau_3(\gamma_{BCD} - 1)]$ | $g_1 = 0$ | $g_2 = 0$ | $g_3 = 2\alpha$ |
| $d^{-(d_2+d_3+d_5)}$ | $g_0 = exp[-\alpha\tau_1(\gamma_D - \gamma_{CD}) - \alpha\tau_2(\gamma_{CD} - \gamma_{BCD}) - \alpha\tau_3(\gamma_A + \gamma_{BCD} - 2)]$ | $g_1 = 0$ | $g_2 = 2\alpha$ | $g_3 = 0$ |
| $d^{-(d_2+d_4+d_5)}$ | $g_0 = exp[-\alpha\tau_1(\gamma_C + \gamma_D - 2\gamma_{CD}) - 2\alpha\tau_2(\gamma_{CD} - \gamma_{BCD}) - 2\alpha\tau_3(\gamma_{BCD} - 1))]$ | $g_1 = 0$ | $g_2 = 2\alpha$ | $g_3 = 0$ |
| $d^{-(d_1+d_2+d_3+d_4)}$ | $g_0 = exp[-\alpha\tau_1(\gamma_C + \gamma_D - 2\gamma_{CD}) - \alpha\tau_2(\gamma_B + 2\gamma_{CD} - 3\gamma_{BCD}) - \alpha\tau_3(\gamma_A + 3\gamma_{BCD} - 4)]$ | $g_1 = 2\alpha$ | $g_2 = 0$ | $g_3 = 2\alpha$ |
| $d^{-(d_1+d_2+d_3+d_5)}$ | $g_0 = exp[-\alpha\tau_1(\gamma_D - \gamma_{CD}) - \alpha\tau_2(\gamma_B + \gamma_{CD} - 2\gamma_{BCD}) - \alpha\tau_3(\gamma_A + 2\gamma_{BCD} - 3)]$ | $g_1 = 0$ | $g_2 = \alpha$ | $g_3 = 2\alpha$ |
| $d^{-(d_1+d_2+d_4+d_5)}$ | $g_0 = exp[-\alpha\tau_1(\gamma_C + \gamma_D - 2\gamma_{CD}) - \alpha\tau_2(\gamma_B + 2\gamma_{CD} - 3\gamma_{BCD}) - 3\alpha\tau_3(\gamma_{BCD} - 1)]$ | $g_1 = 0$ | $g_2 = \alpha$ | $g_3 = 2\alpha$ |
| $d^{-(d_1+d_3+d_4+d_5)}$ | $g_0 = exp[-\alpha\tau_1(\gamma_C - \gamma_{CD}) - \alpha\tau_2(\gamma_B + \gamma_{CD} - 2\gamma_{BCD}) - \alpha\tau_3(\gamma_A + 2\gamma_{BCD} - 3)]$ | $g_1 = \alpha$ | $g_2 = 0$ | $g_3 = 2\alpha$ |
| $d^{-(d_2+d_3+d_4+d_5)}$ | $g_0 = exp[-\alpha\tau_1(\gamma_C + \gamma_D - 2\gamma_{CD}) - 2\alpha\tau_2(\gamma_{CD} - \gamma_{BCD}) - \alpha\tau_3(\gamma_A + 2\gamma_{BCD} - 3)]$ | $g_1 = \alpha$ | $g_2 = 2\alpha$ | $g_3 = 0$ |
| $d^{-(d_1+d_2+d_3+d_4+d_5)}$ | $g_0 = exp[-\alpha\tau_1(\gamma_C + \gamma_D - 2\gamma_{CD}) - \alpha\tau_2(\gamma_B + 2\gamma_{CD} - 3\gamma_{BCD}) - \alpha\tau_3(\gamma_A + 3\gamma_{BCD} - 4)]$ | $g_1 = \alpha$ | $g_2 = \alpha$ | $g_3 = 2\alpha$ |



| | **History 27** | | | |
|---|---|---|---|---|
| $d^{-(d_1+d_2)}$ | $g_0 = exp[-\alpha\tau_1(\gamma_C - \gamma_{CD}) - \alpha\tau_2(\gamma_{CD} - \gamma_{BCD}) - \alpha\tau_3(\gamma_A + \gamma_{BCD} - 2)]$ | $g_1 = 0$ | $g_2 = 0$ | $g_3 = 2\alpha$ |
| $d^{-(d_3+d_4)}$ | $g_0 = exp[-\alpha\tau_1(\gamma_D - 2\gamma_{CD}) - \alpha\tau_2(\gamma_B + \gamma_{CD} - 2\gamma_{BCD})]$ | $g_1 = 2\alpha\gamma_{BCD}$ | $g_2 = 0$ | $g_3 = 0$ |
| $d^{-(d_1+d_3+d_5)}$ | $g_0 = exp[-\alpha\tau_1(\gamma_C - \gamma_{CD}) - \alpha\tau_2(\gamma_B + \gamma_{CD} - 2\gamma_{BCD}) - 2\alpha\tau_3(\gamma_{BCD} - 1)]$ | $g_1 = 0$ | $g_2 = 0$ | $g_3 = 2\alpha$ |
| $d^{-(d_1+d_4+d_5)}$ | $g_0 = exp[-\alpha\tau_1(\gamma_C + \gamma_D - 2\gamma_{CD}) - 2\alpha\tau_2(\gamma_{CD} - \gamma_{BCD}) - 2\alpha\tau_3(\gamma_{BCD} - 1)]$ | $g_1 = 0$ | $g_2 = 0$ | $g_3 = 2\alpha$ |
| $d^{-(d_2+d_3+d_5)}$ | $g_0 = exp[-\alpha\tau_2(\gamma_B - \gamma_{BCD}) - \alpha\tau_3(\gamma_A + \gamma_{BCD} - 2)]$ | $g_1 = 0$ | $g_2 = 2\alpha$ | $g_3 = 0$ |
| $d^{-(d_2+d_4+d_5)}$ | $g_0 = exp[-\alpha\tau_1(\gamma_D - \gamma_{CD}) - \alpha\tau_2(\gamma_{CD} - \gamma_{BCD}) - \alpha\tau_3(\gamma_A + \gamma_{BCD} - 2))]$ | $g_1 = 0$ | $g_2 = 2\alpha$ | $g_3 = 0$ |
| $d^{-(d_1+d_2+d_3+d_4)}$ | $g_0 = exp[-\alpha\tau_1(\gamma_C + \gamma_D - 2\gamma_{CD}) - \alpha\tau_2(\gamma_B + 2\gamma_{CD} - 3\gamma_{BCD}) - \alpha\tau_3(\gamma_A + \gamma_{BCD} - 2)]$ | $g_1 = 2\alpha\gamma_{BCD}$ | $g_2 = 0$ | $g_3 = 2\alpha$ |
| $d^{-(d_1+d_2+d_3+d_5)}$ | $g_0 = exp[-\alpha\tau_1(\gamma_C - \gamma_{CD}) - \alpha\tau_2(\gamma_B + \gamma_{CD} - 2\gamma_{BCD}) - \alpha\tau_3(\gamma_A + 2\gamma_{BCD} - 3)]$ | $g_1 = 0$ | $g_2 = \alpha$ | $g_3 = 2\alpha$ |
| $d^{-(d_1+d_2+d_4+d_5)}$ | $g_0 = exp[-\alpha\tau_1(\gamma_C + \gamma_D - 2\gamma_{CD}) - 2\alpha\tau_2(\gamma_{CD} - \gamma_{BCD}) - \alpha\tau_3(\gamma_A + 2\gamma_{BCD} - 3)]$ | $g_1 = 0$ | $g_2 = \alpha$ | $g_3 = 2\alpha$ |
| $d^{-(d_1+d_3+d_4+d_5)}$ | $g_0 = exp[-\alpha\tau_1(\gamma_C + \gamma_D - 2\gamma_{CD}) - \alpha\tau_2(\gamma_B + 2\gamma_{CD} - 3\gamma_{BCD}) - 2\alpha\tau_3(\gamma_{BCD} - 1)]$ | $g_1 = \alpha\gamma_{BCD}$ | $g_2 = 0$ | $g_3 = 2\alpha$ |
| $d^{-(d_2+d_3+d_4+d_5)}$ | $g_0 = exp[-\alpha\tau_1(\gamma_D - \gamma_{CD}) - \alpha\tau_2(\gamma_B + \gamma_{CD} - 2\gamma_{BCD}) - \alpha\tau_3(\gamma_A + \gamma_{BCD} - 2)]$ | $g_1 = \alpha\gamma_{BCD}$ | $g_2 = 2\alpha$ | $g_3 = 0$ |
| $d^{-(d_1+d_2+d_3+d_4+d_5)}$ | $g_0 = exp[-\alpha\tau_1(\gamma_C + \gamma_D - 2\gamma_{CD}) - \alpha\tau_2(\gamma_B + 2\gamma_{CD} - 3\gamma_{BCD}) - \alpha\tau_3(\gamma_A + 2\gamma_{BCD} - 3)]$ | $g_1 = \alpha\gamma_{BCD}$ | $g_2 = \alpha$ | $g_3 = 2\alpha$ |

| | **History 28** | | | |
|---|---|---|---|---|
| $d^{-(d_1+d_2)}$ | $g_0 = exp[-\alpha\tau_1(\gamma_C - \gamma_{CD}) - \alpha\tau_2(\gamma_{CD} - \gamma_{BCD}) - \alpha\tau_3(\gamma_A + \gamma_{BCD} - 2)]$ | $g_1 = 0$ | $g_2 = 0$ | $g_3 = 2\alpha$ |
| $d^{-(d_3+d_4)}$ | $g_0 = exp[-\alpha\tau_1(\gamma_D - \gamma_{CD}) - \alpha\tau_2(\gamma_B + \gamma_{CD} - 2\gamma_{BCD}) - 2\alpha\tau_3(\gamma_{BCD} - 1)]$ | $g_1 = 2\alpha$ | $g_2 = 0$ | $g_3 = 0$ |
| $d^{-(d_1+d_3+d_5)}$ | $g_0 = exp[-\alpha\tau_1(\gamma_C - \gamma_{CD}) - \alpha\tau_2(\gamma_B + \gamma_{CD} - 2\gamma_{BCD}) - 2\alpha\tau_3(\gamma_{BCD} - 1)]$ | $g_1 = 0$ | $g_2 = 0$ | $g_3 = 2\alpha$ |
| $d^{-(d_1+d_4+d_5)}$ | $g_0 = exp[-\alpha\tau_1(\gamma_C + \gamma_D - 2\gamma_{CD}) - 2\alpha\tau_2(\gamma_{CD} - \gamma_{BCD}) - 2\alpha\tau_3(\gamma_{BCD} - 1)]$ | $g_1 = 0$ | $g_2 = 0$ | $g_3 = 2\alpha$ |
| $d^{-(d_2+d_3+d_5)}$ | $g_0 = exp[-\alpha\tau_2(\gamma_B - \gamma_{BCD}) - \alpha\tau_3(\gamma_A + \gamma_{BCD} - 2)]$ | $g_1 = 0$ | $g_2 = 2\alpha$ | $g_3 = 0$ |
| $d^{-(d_2+d_4+d_5)}$ | $g_0 = exp[-\alpha\tau_1(\gamma_D - \gamma_{CD}) - \alpha\tau_2(\gamma_{CD} - \gamma_{BCD}) - \alpha\tau_3(\gamma_A + \gamma_{BCD} - 2))]$ | $g_1 = 0$ | $g_2 = 2\alpha$ | $g_3 = 0$ |
| $d^{-(d_1+d_2+d_3+d_4)}$ | $g_0 = exp[-\alpha\tau_1(\gamma_C + \gamma_D - 2\gamma_{CD}) - \alpha\tau_2(\gamma_B + 2\gamma_{CD} - 3\gamma_{BCD}) - \alpha\tau_3(\gamma_A + 3\gamma_{BCD} - 4)]$ | $g_1 = 2\alpha$ | $g_2 = 0$ | $g_3 = 2\alpha$ |
| $d^{-(d_1+d_2+d_3+d_5)}$ | $g_0 = exp[-\alpha\tau_1(\gamma_C - \gamma_{CD}) - \alpha\tau_2(\gamma_B + \gamma_{CD} - 2\gamma_{BCD}) - \alpha\tau_3(\gamma_A + 2\gamma_{BCD} - 3)]$ | $g_1 = 0$ | $g_2 = \alpha$ | $g_3 = 2\alpha$ |
| $d^{-(d_1+d_2+d_4+d_5)}$ | $g_0 = exp[-\alpha\tau_1(\gamma_C + \gamma_D - 2\gamma_{CD}) - 2\alpha\tau_2(\gamma_{CD} - \gamma_{BCD}) - \alpha\tau_3(\gamma_A + 2\gamma_{BCD} - 3)]$ | $g_1 = 0$ | $g_2 = \alpha$ | $g_3 = 2\alpha$ |
| $d^{-(d_1+d_3+d_4+d_5)}$ | $g_0 = exp[-\alpha\tau_1(\gamma_C + \gamma_D - 2\gamma_{CD}) - \alpha\tau_2(\gamma_B + 2\gamma_{CD} - 3\gamma_{BCD}) - 3\alpha\tau_3(\gamma_{BCD} - 1)]$ | $g_1 = \alpha$ | $g_2 = 0$ | $g_3 = 2\alpha$ |
| $d^{-(d_2+d_3+d_4+d_5)}$ | $g_0 = exp[-\alpha\tau_1(\gamma_D - \gamma_{CD}) - \alpha\tau_2(\gamma_B + \gamma_{CD} - 2\gamma_{BCD}) - \alpha\tau_3(\gamma_A + 2\gamma_{BCD} - 3)]$ | $g_1 = \alpha$ | $g_2 = 2\alpha$ | $g_3 = 0$ |
| $d^{-(d_1+d_2+d_3+d_4+d_5)}$ | $g_0 = exp[-\alpha\tau_1(\gamma_C + \gamma_D - 2\gamma_{CD}) - \alpha\tau_2(\gamma_B + 2\gamma_{CD} - 3\gamma_{BCD}) - \alpha\tau_3(\gamma_A + 3\gamma_{BCD} - 4)]$ | $g_1 = \alpha$ | $g_2 = \alpha$ | $g_3 = 2\alpha$ |

| | **History 29** | | | |
|---|---|---|---|---|
| $d^{-(d_1+d_2)}$ | $g_0 = exp[-\alpha\tau_1(\gamma_C - \gamma_{CD}) - \alpha\tau_2(\gamma_B + \gamma_{CD} - 2\gamma_{BCD}) - 2\alpha\tau_3(\gamma_{BCD} - 1)]$ | $g_1 = 0$ | $g_2 = 0$ | $g_3 = 2\alpha$ |
| $d^{-(d_3+d_4)}$ | $g_0 = exp[-\alpha\tau_1(\gamma_D - \gamma_{CD}) - \alpha\tau_2(\gamma_{CD} - \gamma_{BCD}) - \alpha\tau_3(\gamma_A + \gamma_{BCD} - 2)]$ | $g_1 = 2\alpha$ | $g_2 = 0$ | $g_3 = 0$ |
| $d^{-(d_1+d_3+d_5)}$ | $g_0 = exp[-\alpha\tau_1(\gamma_C - \gamma_{CD}) - \alpha\tau_2(\gamma_{CD} - \gamma_{BCD}) - \alpha\tau_3(\gamma_A + \gamma_{BCD} - 2)]$ | $g_1 = 0$ | $g_2 = 0$ | $g_3 = 2\alpha$ |
| $d^{-(d_1+d_4+d_5)}$ | $g_0 = exp[-\alpha\tau_1(\gamma_C + \gamma_D - 2\gamma_{CD}) - 2\alpha\tau_2(\gamma_{CD} - \gamma_{BCD}) - 2\alpha\tau_3(\gamma_{BCD} - 1)]$ | $g_1 = 0$ | $g_2 = 0$ | $g_3 = 2\alpha$ |
| $d^{-(d_2+d_3+d_5)}$ | $g_0 = exp[-\alpha\tau_2(\gamma_B - \gamma_{BCD}) - \alpha\tau_3(\gamma_A + \gamma_{BCD} - 2)]$ | $g_1 = 0$ | $g_2 = 2\alpha$ | $g_3 = 0$ |
| $d^{-(d_2+d_4+d_5)}$ | $g_0 = exp[-\alpha\tau_1(\gamma_D - \gamma_{CD}) - \alpha\tau_2(\gamma_B + \gamma_{CD} - 2\gamma_{BCD}) - 2\alpha\tau_3(\gamma_{BCD} - 1))]$ | $g_1 = 0$ | $g_2 = 2\alpha$ | $g_3 = 0$ |
| $d^{-(d_1+d_2+d_3+d_4)}$ | $g_0 = exp[-\alpha\tau_1(\gamma_C + \gamma_D - 2\gamma_{CD}) - \alpha\tau_2(\gamma_B + 2\gamma_{CD} - 3\gamma_{BCD}) - \alpha\tau_3(\gamma_A + 3\gamma_{BCD} - 4)]$ | $g_1 = 2\alpha$ | $g_2 = 0$ | $g_3 = 2\alpha$ |
| $d^{-(d_1+d_2+d_3+d_5)}$ | $g_0 = exp[-\alpha\tau_1(\gamma_C - \gamma_{CD}) - \alpha\tau_2(\gamma_B + \gamma_{CD} - 2\gamma_{BCD}) - \alpha\tau_3(\gamma_A + 2\gamma_{BCD} - 3)]$ | $g_1 = 0$ | $g_2 = \alpha$ | $g_3 = 2\alpha$ |
| $d^{-(d_1+d_2+d_4+d_5)}$ | $g_0 = exp[-\alpha\tau_1(\gamma_C + \gamma_D - 2\gamma_{CD}) - \alpha\tau_2(\gamma_B + 2\gamma_{CD} - 3\gamma_{BCD}) - 3\alpha\tau_3(\gamma_{BCD} - 1)]$ | $g_1 = 0$ | $g_2 = \alpha$ | $g_3 = 2\alpha$ |
| $d^{-(d_1+d_3+d_4+d_5)}$ | $g_0 = exp[-\alpha\tau_1(\gamma_C + \gamma_D - 2\gamma_{CD}) - 2\alpha\tau_2(\gamma_{CD} - \gamma_{BCD}) - \alpha\tau_3(\gamma_A + 2\gamma_{BCD} - 3)]$ | $g_1 = \alpha$ | $g_2 = 0$ | $g_3 = 2\alpha$ |
| $d^{-(d_2+d_3+d_4+d_5)}$ | $g_0 = exp[-\alpha\tau_1(\gamma_D - \gamma_{CD}) - \alpha\tau_2(\gamma_B + \gamma_{CD} - 2\gamma_{BCD}) - \alpha\tau_3(\gamma_A + 2\gamma_{BCD} - 3)]$ | $g_1 = \alpha$ | $g_2 = 2\alpha$ | $g_3 = 0$ |
| $d^{-(d_1+d_2+d_3+d_4+d_5)}$ | $g_0 = exp[-\alpha\tau_1(\gamma_C + \gamma_D - 2\gamma_{CD}) - \alpha\tau_2(\gamma_B + 2\gamma_{CD} - 3\gamma_{BCD}) - \alpha\tau_3(\gamma_A + 3\gamma_{BCD} - 4)]$ | $g_1 = \alpha$ | $g_2 = \alpha$ | $g_3 = 2\alpha$ |

| | **History 30** | | | |
|---|---|---|---|---|
| $d^{-(d_1+d_2)}$ | $g_0 = exp[-\alpha\tau_1(\gamma_C + \gamma_D - 2\gamma_{CD}) - 2\alpha\tau_2(\gamma_{CD} - \gamma_{BCD}) - 2\alpha\tau_3(\gamma_{BCD} - 1)]$ | $g_1 = 0$ | $g_2 = 0$ | $g_3 = 2\alpha$ |
| $d^{-(d_3+d_4)}$ | $g_0 = exp[-\alpha\tau_2(\gamma_B - \gamma_{BCD}) - \alpha\tau_3(\gamma_A + \gamma_{BCD} - 2)]$ | $g_1 = 2\alpha$ | $g_2 = 0$ | $g_3 = 0$ |
| $d^{-(d_1+d_3+d_5)}$ | $g_0 = exp[-\alpha\tau_1(\gamma_C - \gamma_{CD}) - \alpha\tau_2(\gamma_{CD} - \gamma_{BCD}) - \alpha\tau_3(\gamma_A + \gamma_{BCD} - 2)]$ | $g_1 = 0$ | $g_2 = 0$ | $g_3 = 2\alpha$ |
| $d^{-(d_1+d_4+d_5)}$ | $g_0 = exp[-\alpha\tau_1(\gamma_C - \gamma_{CD}) - \alpha\tau_2(\gamma_B + \gamma_{CD} - 2\gamma_{BCD}) - 2\alpha\tau_3(\gamma_{BCD} - 1)]$ | $g_1 = 0$ | $g_2 = 0$ | $g_3 = 2\alpha$ |
| $d^{-(d_2+d_3+d_5)}$ | $g_0 = exp[-\alpha\tau_1(\gamma_D - \gamma_{CD}) - \alpha\tau_2(\gamma_{CD} - \gamma_{BCD}) - \alpha\tau_3(\gamma_A + \gamma_{BCD} - 2)]$ | $g_1 = 0$ | $g_2 = 2\alpha$ | $g_3 = 0$ |
| $d^{-(d_2+d_4+d_5)}$ | $g_0 = exp[-\alpha\tau_1(\gamma_D - \gamma_{CD}) - \alpha\tau_2(\gamma_B + \gamma_{CD} - 2\gamma_{BCD}) - 2\alpha\tau_3(\gamma_{BCD} - 1))]$ | $g_1 = 0$ | $g_2 = 2\alpha$ | $g_3 = 0$ |
| $d^{-(d_1+d_2+d_3+d_4)}$ | $g_0 = exp[-\alpha\tau_1(\gamma_C + \gamma_D - 2\gamma_{CD}) - \alpha\tau_2(\gamma_B + 2\gamma_{CD} - 3\gamma_{BCD}) - \alpha\tau_3(\gamma_A + 3\gamma_{BCD} - 4)]$ | $g_1 = 2\alpha$ | $g_2 = 0$ | $g_3 = 2\alpha$ |
| $d^{-(d_1+d_2+d_3+d_5)}$ | $g_0 = exp[-\alpha\tau_1(\gamma_C + \gamma_D - 2\gamma_{CD}) - 2\alpha\tau_2(\gamma_{CD} - \gamma_{BCD}) - \alpha\tau_3(\gamma_A + 2\gamma_{BCD} - 3)]$ | $g_1 = 0$ | $g_2 = \alpha$ | $g_3 = 2\alpha$ |
| $d^{-(d_1+d_2+d_4+d_5)}$ | $g_0 = exp[-\alpha\tau_1(\gamma_C + \gamma_D - 2\gamma_{CD}) - \alpha\tau_2(\gamma_B + 2\gamma_{CD} - 3\gamma_{BCD}) - 3\alpha\tau_3(\gamma_{BCD} - 1)]$ | $g_1 = 0$ | $g_2 = \alpha$ | $g_3 = 2\alpha$ |
| $d^{-(d_1+d_3+d_4+d_5)}$ | $g_0 = exp[-\alpha\tau_1(\gamma_C - \gamma_{CD}) - \alpha\tau_2(\gamma_B + \gamma_{CD} - 2\gamma_{BCD}) - \alpha\tau_3(\gamma_A + 2\gamma_{BCD} - 3)]$ | $g_1 = \alpha$ | $g_2 = 0$ | $g_3 = 2\alpha$ |
| $d^{-(d_2+d_3+d_4+d_5)}$ | $g_0 = exp[-\alpha\tau_1(\gamma_D - \gamma_{CD}) - \alpha\tau_2(\gamma_B + \gamma_{CD} - 2\gamma_{BCD}) - \alpha\tau_3(\gamma_A + 2\gamma_{BCD} - 3)]$ | $g_1 = \alpha$ | $g_2 = 2\alpha$ | $g_3 = 0$ |
| $d^{-(d_1+d_2+d_3+d_4+d_5)}$ | $g_0 = exp[-\alpha\tau_1(\gamma_C + \gamma_D - 2\gamma_{CD}) - \alpha\tau_2(\gamma_B + 2\gamma_{CD} - 3\gamma_{BCD}) - \alpha\tau_3(\gamma_A + 3\gamma_{BCD} - 4)]$ | $g_1 = \alpha$ | $g_2 = \alpha$ | $g_3 = 2\alpha$ |

| | **History 31** | | | |
|---|---|---|---|---|
| $d^{-(d_1+d_2)}$ | $g_0 = exp[-\alpha\tau_1(\gamma_D - \gamma_{CD}) - \alpha\tau_2(\gamma_{CD} - \gamma_{BCD}) - \alpha\tau_3(\gamma_A + \gamma_{BCD} - 2)]$ | $g_1 = 0$ | $g_2 = 0$ | $g_3 = 2\alpha$ |
| $d^{-(d_3+d_4)}$ | $g_0 = exp[-\alpha\tau_1(\gamma_C - 2\gamma_{CD}) - \alpha\tau_2(\gamma_B + \gamma_{CD} - 2\gamma_{BCD})]$ | $g_1 = 2\alpha\gamma_{BCD}$ | $g_2 = 0$ | $g_3 = 0$ |
| $d^{-(d_1+d_3+d_5)}$ | $g_0 = exp[-\alpha\tau_1(\gamma_D - \gamma_{CD}) - \alpha\tau_2(\gamma_B + \gamma_{CD} - 2\gamma_{BCD}) - 2\alpha\tau_3(\gamma_{BCD} - 1)]$ | $g_1 = 0$ | $g_2 = 0$ | $g_3 = 2\alpha$ |
| $d^{-(d_1+d_4+d_5)}$ | $g_0 = exp[-\alpha\tau_1(\gamma_C + \gamma_D - 2\gamma_{CD}) - 2\alpha\tau_2(\gamma_{CD} - \gamma_{BCD}) - 2\alpha\tau_3(\gamma_{BCD} - 1)]$ | $g_1 = 0$ | $g_2 = 0$ | $g_3 = 2\alpha$ |
| $d^{-(d_2+d_3+d_5)}$ | $g_0 = exp[-\alpha\tau_2(\gamma_B - \gamma_{BCD}) - \alpha\tau_3(\gamma_A + \gamma_{BCD} - 2)]$ | $g_1 = 0$ | $g_2 = 2\alpha$ | $g_3 = 0$ |
| $d^{-(d_2+d_4+d_5)}$ | $g_0 = exp[-\alpha\tau_1(\gamma_C - \gamma_{CD}) - \alpha\tau_2(\gamma_{CD} - \gamma_{BCD}) - \alpha\tau_3(\gamma_A + \gamma_{BCD} - 2))]$ | $g_1 = 0$ | $g_2 = 2\alpha$ | $g_3 = 0$ |
| $d^{-(d_1+d_2+d_3+d_4)}$ | $g_0 = exp[-\alpha\tau_1(\gamma_C + \gamma_D - 2\gamma_{CD}) - \alpha\tau_2(\gamma_B + 2\gamma_{CD} - 3\gamma_{BCD}) - \alpha\tau_3(\gamma_A + \gamma_{BCD} - 2)]$ | $g_1 = 2\alpha\gamma_{BCD}$ | $g_2 = 0$ | $g_3 = 2\alpha$ |
| $d^{-(d_1+d_2+d_3+d_5)}$ | $g_0 = exp[-\alpha\tau_1(\gamma_D - \gamma_{CD}) - \alpha\tau_2(\gamma_B + \gamma_{CD} - 2\gamma_{BCD}) - \alpha\tau_3(\gamma_A + 2\gamma_{BCD} - 3)]$ | $g_1 = 0$ | $g_2 = \alpha$ | $g_3 = 2\alpha$ |
| $d^{-(d_1+d_2+d_4+d_5)}$ | $g_0 = exp[-\alpha\tau_1(\gamma_C + \gamma_D - 2\gamma_{CD}) - 2\alpha\tau_2(\gamma_{CD} - \gamma_{BCD}) - \alpha\tau_3(\gamma_A + 2\gamma_{BCD} - 3)]$ | $g_1 = 0$ | $g_2 = \alpha$ | $g_3 = 2\alpha$ |
| $d^{-(d_1+d_3+d_4+d_5)}$ | $g_0 = exp[-\alpha\tau_1(\gamma_C + \gamma_D - 2\gamma_{CD}) - \alpha\tau_2(\gamma_B + 2\gamma_{CD} - 3\gamma_{BCD}) - 2\alpha\tau_3(\gamma_{BCD} - 1)]$ | $g_1 = \alpha\gamma_{BCD}$ | $g_2 = 0$ | $g_3 = 2\alpha$ |
| $d^{-(d_2+d_3+d_4+d_5)}$ | $g_0 = exp[-\alpha\tau_1(\gamma_C - \gamma_{CD}) - \alpha\tau_2(\gamma_B + \gamma_{CD} - 2\gamma_{BCD}) - \alpha\tau_3(\gamma_A + \gamma_{BCD} - 2)]$ | $g_1 = \alpha\gamma_{BCD}$ | $g_2 = 2\alpha$ | $g_3 = 0$ |
| $d^{-(d_1+d_2+d_3+d_4+d_5)}$ | $g_0 = exp[-\alpha\tau_1(\gamma_C + \gamma_D - 2\gamma_{CD}) - \alpha\tau_2(\gamma_B + 2\gamma_{CD} - 3\gamma_{BCD}) - \alpha\tau_3(\gamma_A + 2\gamma_{BCD} - 3)]$ | $g_1 = \alpha\gamma_{BCD}$ | $g_2 = \alpha$ | $g_3 = 2\alpha$ |

| | **History 32** | | | |
|---|---|---|---|---|
| $d^{-(d_1+d_2)}$ | $g_0 = exp[-\alpha\tau_1(\gamma_D - \gamma_{CD}) - \alpha\tau_2(\gamma_{CD} - \gamma_{BCD}) - \alpha\tau_3(\gamma_A + \gamma_{BCD} - 2)]$ | $g_1 = 0$ | $g_2 = 0$ | $g_3 = 2\alpha$ |
| $d^{-(d_3+d_4)}$ | $g_0 = exp[-\alpha\tau_1(\gamma_C - \gamma_{CD}) - \alpha\tau_2(\gamma_B + \gamma_{CD} - 2\gamma_{BCD}) - 2\alpha\tau_3(\gamma_{BCD} - 1)]$ | $g_1 = 2\alpha$ | $g_2 = 0$ | $g_3 = 0$ |
| $d^{-(d_1+d_3+d_5)}$ | $g_0 = exp[-\alpha\tau_1(\gamma_D - \gamma_{CD}) - \alpha\tau_2(\gamma_B + \gamma_{CD} - 2\gamma_{BCD}) - 2\alpha\tau_3(\gamma_{BCD} - 1)]$ | $g_1 = 0$ | $g_2 = 0$ | $g_3 = 2\alpha$ |
| $d^{-(d_1+d_4+d_5)}$ | $g_0 = exp[-\alpha\tau_1(\gamma_C + \gamma_D - 2\gamma_{CD}) - 2\alpha\tau_2(\gamma_{CD} - \gamma_{BCD}) - 2\alpha\tau_3(\gamma_{BCD} - 1)]$ | $g_1 = 0$ | $g_2 = 0$ | $g_3 = 2\alpha$ |
| $d^{-(d_2+d_3+d_5)}$ | $g_0 = exp[-\alpha\tau_2(\gamma_B - \gamma_{BCD}) - \alpha\tau_3(\gamma_A + \gamma_{BCD} - 2)]$ | $g_1 = 0$ | $g_2 = 2\alpha$ | $g_3 = 0$ |
| $d^{-(d_2+d_4+d_5)}$ | $g_0 = exp[-\alpha\tau_1(\gamma_C - \gamma_{CD}) - \alpha\tau_2(\gamma_{CD} - \gamma_{BCD}) - \alpha\tau_3(\gamma_A + \gamma_{BCD} - 2))]$ | $g_1 = 0$ | $g_2 = 2\alpha$ | $g_3 = 0$ |
| $d^{-(d_1+d_2+d_3+d_4)}$ | $g_0 = exp[-\alpha\tau_1(\gamma_C + \gamma_D - 2\gamma_{CD}) - \alpha\tau_2(\gamma_B + 2\gamma_{CD} - 3\gamma_{BCD}) - \alpha\tau_3(\gamma_A + 3\gamma_{BCD} - 4)]$ | $g_1 = 2\alpha$ | $g_2 = 0$ | $g_3 = 2\alpha$ |
| $d^{-(d_1+d_2+d_3+d_5)}$ | $g_0 = exp[-\alpha\tau_1(\gamma_D - \gamma_{CD}) - \alpha\tau_2(\gamma_B + \gamma_{CD} - 2\gamma_{BCD}) - \alpha\tau_3(\gamma_A + 2\gamma_{BCD} - 3)]$ | $g_1 = 0$ | $g_2 = \alpha$ | $g_3 = 2\alpha$ |
| $d^{-(d_1+d_2+d_4+d_5)}$ | $g_0 = exp[-\alpha\tau_1(\gamma_C + \gamma_D - 2\gamma_{CD}) - 2\alpha\tau_2(\gamma_{CD} - \gamma_{BCD}) - \alpha\tau_3(\gamma_A + 2\gamma_{BCD} - 3)]$ | $g_1 = 0$ | $g_2 = \alpha$ | $g_3 = 2\alpha$ |
| $d^{-(d_1+d_3+d_4+d_5)}$ | $g_0 = exp[-\alpha\tau_1(\gamma_C + \gamma_D - 2\gamma_{CD}) - \alpha\tau_2(\gamma_B + 2\gamma_{CD} - 3\gamma_{BCD}) - 3\alpha\tau_3(\gamma_{BCD} - 1)]$ | $g_1 = \alpha$ | $g_2 = 0$ | $g_3 = 2\alpha$ |
| $d^{-(d_2+d_3+d_4+d_5)}$ | $g_0 = exp[-\alpha\tau_1(\gamma_C - \gamma_{CD}) - \alpha\tau_2(\gamma_B + \gamma_{CD} - 2\gamma_{BCD}) - \alpha\tau_3(\gamma_A + 2\gamma_{BCD} - 3)]$ | $g_1 = \alpha$ | $g_2 = 2\alpha$ | $g_3 = 0$ |
| $d^{-(d_1+d_2+d_3+d_4+d_5)}$ | $g_0 = exp[-\alpha\tau_1(\gamma_C + \gamma_D - 2\gamma_{CD}) - \alpha\tau_2(\gamma_B + 2\gamma_{CD} - 3\gamma_{BCD}) - \alpha\tau_3(\gamma_A + 3\gamma_{BCD} - 4)]$ | $g_1 = \alpha$ | $g_2 = \alpha$ | $g_3 = 2\alpha$ |



| | **History 33** | | | |
|---|---|---|---|---|
| $d^{-(d_1+d_2)}$ | $g_0 = exp[-\alpha\tau_1(\gamma_D - \gamma_{CD}) - \alpha\tau_2(\gamma_B + \gamma_{CD} - 2\gamma_{BCD}) - 2\alpha\tau_3(\gamma_{BCD} - 1)]$ | $g_1 = 0$ | $g_2 = 0$ | $g_3 = 2\alpha$ |
| $d^{-(d_3+d_4)}$ | $g_0 = exp[-\alpha\tau_1(\gamma_C - \gamma_{CD}) - \alpha\tau_2(\gamma_{CD} - \gamma_{BCD}) - \alpha\tau_3(\gamma_A + \gamma_{BCD} - 2)]$ | $g_1 = 2\alpha$ | $g_2 = 0$ | $g_3 = 0$ |
| $d^{-(d_1+d_3+d_5)}$ | $g_0 = exp[-\alpha\tau_1(\gamma_D - \gamma_{CD}) - \alpha\tau_2(\gamma_{CD} - \gamma_{BCD}) - \alpha\tau_3(\gamma_A + \gamma_{BCD} - 2)]$ | $g_1 = 0$ | $g_2 = 0$ | $g_3 = 2\alpha$ |
| $d^{-(d_1+d_4+d_5)}$ | $g_0 = exp[-\alpha\tau_1(\gamma_C + \gamma_D - 2\gamma_{CD}) - 2\alpha\tau_2(\gamma_{CD} - \gamma_{BCD}) - 2\alpha\tau_3(\gamma_{BCD} - 1)]$ | $g_1 = 0$ | $g_2 = 0$ | $g_3 = 2\alpha$ |
| $d^{-(d_2+d_3+d_5)}$ | $g_0 = exp[-\alpha\tau_2(\gamma_B - \gamma_{BCD}) - \alpha\tau_3(\gamma_A + \gamma_{BCD} - 2)]$ | $g_1 = 0$ | $g_2 = 2\alpha$ | $g_3 = 0$ |
| $d^{-(d_2+d_4+d_5)}$ | $g_0 = exp[-\alpha\tau_1(\gamma_C - \gamma_{CD}) - \alpha\tau_2(\gamma_B + \gamma_{CD} - 2\gamma_{BCD}) - 2\alpha\tau_3(\gamma_{BCD} - 1))]$ | $g_1 = 0$ | $g_2 = 2\alpha$ | $g_3 = 0$ |
| $d^{-(d_1+d_2+d_3+d_4)}$ | $g_0 = exp[-\alpha\tau_1(\gamma_C + \gamma_D - 2\gamma_{CD}) - \alpha\tau_2(\gamma_B + 2\gamma_{CD} - 3\gamma_{BCD}) - \alpha\tau_3(\gamma_A + 3\gamma_{BCD} - 4)]$ | $g_1 = 2\alpha$ | $g_2 = 0$ | $g_3 = 2\alpha$ |
| $d^{-(d_1+d_2+d_3+d_5)}$ | $g_0 = exp[-\alpha\tau_1(\gamma_D - \gamma_{CD}) - \alpha\tau_2(\gamma_B + \gamma_{CD} - 2\gamma_{BCD}) - \alpha\tau_3(\gamma_A + 2\gamma_{BCD} - 3)]$ | $g_1 = 0$ | $g_2 = \alpha$ | $g_3 = 2\alpha$ |
| $d^{-(d_1+d_2+d_4+d_5)}$ | $g_0 = exp[-\alpha\tau_1(\gamma_C + \gamma_D - 2\gamma_{CD}) - \alpha\tau_2(\gamma_B + 2\gamma_{CD} - 3\gamma_{BCD}) - 3\alpha\tau_3(\gamma_{BCD} - 1)]$ | $g_1 = 0$ | $g_2 = \alpha$ | $g_3 = 2\alpha$ |
| $d^{-(d_1+d_3+d_4+d_5)}$ | $g_0 = exp[-\alpha\tau_1(\gamma_C + \gamma_D - 2\gamma_{CD}) - 2\alpha\tau_2(\gamma_{CD} - \gamma_{BCD}) - \alpha\tau_3(\gamma_A + 2\gamma_{BCD} - 3)]$ | $g_1 = \alpha$ | $g_2 = 0$ | $g_3 = 2\alpha$ |
| $d^{-(d_2+d_3+d_4+d_5)}$ | $g_0 = exp[-\alpha\tau_1(\gamma_C - \gamma_{CD}) - \alpha\tau_2(\gamma_B + \gamma_{CD} - 2\gamma_{BCD}) - \alpha\tau_3(\gamma_A + 2\gamma_{BCD} - 3)]$ | $g_1 = \alpha$ | $g_2 = 2\alpha$ | $g_3 = 0$ |
| $d^{-(d_1+d_2+d_3+d_4+d_5)}$ | $g_0 = exp[-\alpha\tau_1(\gamma_C + \gamma_D - 2\gamma_{CD}) - \alpha\tau_2(\gamma_B + 2\gamma_{CD} - 3\gamma_{BCD}) - \alpha\tau_3(\gamma_A + 3\gamma_{BCD} - 4)]$ | $g_1 = \alpha$ | $g_2 = \alpha$ | $g_3 = 2\alpha$ |

| | **History 34** | | | |
|---|---|---|---|---|
| $d^{-(d_1+d_2)}$ | $g_0 = exp[-\alpha\tau_1(\gamma_C + \gamma_D - 2\gamma_{CD}) - 2\alpha\tau_2(\gamma_{CD} - \gamma_{BCD}) - 2\alpha\tau_3(\gamma_{BCD} - 1)]$ | $g_1 = 0$ | $g_2 = 0$ | $g_3 = 2\alpha$ |
| $d^{-(d_3+d_4)}$ | $g_0 = exp[-\alpha\tau_2(\gamma_B - \gamma_{BCD}) - \alpha\tau_3(\gamma_A + \gamma_{BCD} - 2)]$ | $g_1 = 2\alpha$ | $g_2 = 0$ | $g_3 = 0$ |
| $d^{-(d_1+d_3+d_5)}$ | $g_0 = exp[-\alpha\tau_1(\gamma_D - \gamma_{CD}) - \alpha\tau_2(\gamma_{CD} - \gamma_{BCD}) - \alpha\tau_3(\gamma_A + \gamma_{BCD} - 2)]$ | $g_1 = 0$ | $g_2 = 0$ | $g_3 = 2\alpha$ |
| $d^{-(d_1+d_4+d_5)}$ | $g_0 = exp[-\alpha\tau_1(\gamma_D - \gamma_{CD}) - \alpha\tau_2(\gamma_B + \gamma_{CD} - 2\gamma_{BCD}) - 2\alpha\tau_3(\gamma_{BCD} - 1)]$ | $g_1 = 0$ | $g_2 = 0$ | $g_3 = 2\alpha$ |
| $d^{-(d_2+d_3+d_5)}$ | $g_0 = exp[-\alpha\tau_1(\gamma_C - \gamma_{CD}) - \alpha\tau_2(\gamma_{CD} - \gamma_{BCD}) - \alpha\tau_3(\gamma_A + \gamma_{BCD} - 2)]$ | $g_1 = 0$ | $g_2 = 2\alpha$ | $g_3 = 0$ |
| $d^{-(d_2+d_4+d_5)}$ | $g_0 = exp[-\alpha\tau_1(\gamma_C - \gamma_{CD}) - \alpha\tau_2(\gamma_B + \gamma_{CD} - 2\gamma_{BCD}) - 2\alpha\tau_3(\gamma_{BCD} - 1))]$ | $g_1 = 0$ | $g_2 = 2\alpha$ | $g_3 = 0$ |
| $d^{-(d_1+d_2+d_3+d_4)}$ | $g_0 = exp[-\alpha\tau_1(\gamma_C + \gamma_D - 2\gamma_{CD}) - \alpha\tau_2(\gamma_B + 2\gamma_{CD} - 3\gamma_{BCD}) - \alpha\tau_3(\gamma_A + 3\gamma_{BCD} - 4)]$ | $g_1 = 2\alpha$ | $g_2 = 0$ | $g_3 = 2\alpha$ |
| $d^{-(d_1+d_2+d_3+d_5)}$ | $g_0 = exp[-\alpha\tau_1(\gamma_C + \gamma_D - 2\gamma_{CD}) - 2\alpha\tau_2(\gamma_{CD} - \gamma_{BCD}) - \alpha\tau_3(\gamma_A + 2\gamma_{BCD} - 3)]$ | $g_1 = 0$ | $g_2 = \alpha$ | $g_3 = 2\alpha$ |
| $d^{-(d_1+d_2+d_4+d_5)}$ | $g_0 = exp[-\alpha\tau_1(\gamma_C + \gamma_D - 2\gamma_{CD}) - \alpha\tau_2(\gamma_B + 2\gamma_{CD} - 3\gamma_{BCD}) - 3\alpha\tau_3(\gamma_{BCD} - 1)]$ | $g_1 = 0$ | $g_2 = \alpha$ | $g_3 = 2\alpha$ |
| $d^{-(d_1+d_3+d_4+d_5)}$ | $g_0 = exp[-\alpha\tau_1(\gamma_D - \gamma_{CD}) - \alpha\tau_2(\gamma_B + \gamma_{CD} - 2\gamma_{BCD}) - \alpha\tau_3(\gamma_A + 2\gamma_{BCD} - 3)]$ | $g_1 = \alpha$ | $g_2 = 0$ | $g_3 = 2\alpha$ |
| $d^{-(d_2+d_3+d_4+d_5)}$ | $g_0 = exp[-\alpha\tau_1(\gamma_C - \gamma_{CD}) - \alpha\tau_2(\gamma_B + \gamma_{CD} - 2\gamma_{BCD}) - \alpha\tau_3(\gamma_A + 2\gamma_{BCD} - 3)]$ | $g_1 = \alpha$ | $g_2 = 2\alpha$ | $g_3 = 0$ |
| $d^{-(d_1+d_2+d_3+d_4+d_5)}$ | $g_0 = exp[-\alpha\tau_1(\gamma_C + \gamma_D - 2\gamma_{CD}) - \alpha\tau_2(\gamma_B + 2\gamma_{CD} - 3\gamma_{BCD}) - \alpha\tau_3(\gamma_A + 3\gamma_{BCD} - 4)]$ | $g_1 = \alpha$ | $g_2 = \alpha$ | $g_3 = 2\alpha$ |
